# ECPAS/Ecm29-Mediated 26S Proteasome Disassembly is an Adaptive Response to Glucose Starvation

**DOI:** 10.1101/2022.06.06.495046

**Authors:** Won Hoon Choi, Yejin Yun, Insuk Byun, Seho Lee, Jiho Sim, Shahar Levi, Sumin Kim, Seo Hyeong Park, Jeongmoo Jun, Kwang Pyo Kim, Yong Tae Kwon, Dohyun Han, Tomoki Chiba, Chaok Seok, Michael H. Glickman, Min Jae Lee

## Abstract

The 26S proteasome consists of loosely associated 20S catalytic and 19S regulatory complexes. Approximately half of the proteasomes in eukaryotic cells exist as free 20S complexes; however, our mechanistic and physiological understanding of what determines the ratio of 26S to 20S species remains incomplete. Here, we show that glucose starvation in mammalian cells results in the uncoupling of 26S holoenzymes into intact 20S and 19S subcomplexes. Subcomplex affinity-purification and quantitative mass spectrometry revealed that Ecm29 proteasome adaptor and scaffold (ECPAS) is a crucial mediator of this structural remodeling. The loss of ECPAS abrogated 26S dissociation, leading to decreased degradation of 20S proteasome substrates such as puromycylated polypeptides and lysine-less cyclin B. *In silico* modeling analysis suggested that the conformational changes of ECPAS may commence the disassembly process. ECPAS was also essential for proper endoplasmic reticulum stress response and cell survival during glucose starvation. In addition, we evaluated the role of ECPAS *in vivo* using the mouse xenograft model and observed that glucose-deprived tumor tissues had significantly elevated 20S proteasome levels. Collectively, our results indicate that the 20S-19S disassembly mediated by ECPAS is a novel mechanism adapting global proteolysis to physiological needs and an effective cellular strategy against proteotoxic stress.

## Introduction

The rate of proteolysis in a cell is a function of its physiological state, and the flux continuously changes in response to nutritional and environmental status. The ubiquitin (Ub)-proteasome system (UPS) is the primary cellular process responsible for degrading cytosolic, nuclear, and integral/peripheral membrane proteins under normal conditions. The UPS also controls protein quality by selectively proteolyzing misfolded and oxidatively damaged proteins [1]. Recent evidence suggests that aberrations in the UPS lead to the accumulation of potentially harmful proteins and, eventually, protein aggregates, a hallmark of aging and proteopathies [2]. If the proteasomal activity is compromised, nuclear factor (erythroid-derived 2)-like 1 (NFE2L1/Nrf1) escapes from endoplasmic reticulum (ER)-associated protein degradation and promotes *de novo* proteasome biogenesis, inducing a “bounce-back response” [3]. Unlike the UPS, autophagy, another cellular degradation process, usually operates at the basal level; however, during starvation or other types of stress, bulk autophagy is induced and indiscriminately breaks down cytoplasmic constituents to replenish anabolic intermediates [4]. The UPS and autophagy are also connected via intricate regulation mechanisms and coordinate their functions to maintain global proteolytic capacity [5, 6]. As a result, identifying molecular etiologies contributing to abnormal proteostasis and developing pharmacological strategies modulating targeted or global proteolysis are active research areas.

The 26S proteasome is the primary ATP-dependent proteolytic holoenzyme in eukaryotes [7]. The conventional 26S proteasome consists of a catalytic 20S proteasome (hereafter referred to as 20S; molecular weight ∼730 kDa) and either one or two loosely associated, but dissociable, 19S regulatory complexes (19S; ∼930 kDa) [8]. The doubly capped proteasome with a 19S-20S-19S configuration is also known as the 30S proteasome, distinguishing it from the singly capped 26S proteasome. Upon coming together in the 26S holoenzyme, the heptameric PSMA/α ring at the 20S surface and the hexameric AAA+ ATPase PMSC/Rpt ring at the base of the 19S form an asymmetric interface around a contiguous substrate translocation channel from the 19S base into the 20S lumen (please note that we will be using the official names of proteasome subunits and others as established by the HUGO consortium). The axial alignment between the 20S and 19S constantly fluctuates in response to ATP hydrolysis, contributing to the proteolytic potency of 26S proteasomes [9, 10]. Although more than half of proteasomes exist in their free 20S form [11], the 20S proteasome was regarded as a latent enzyme requiring activation. However, the physiological relevance of Ub-independent, 20S-mediated proteolysis in eukaryotes has been increasingly recognized, especially for degradation of disordered or oxidized proteins [12, 13]. Furthermore, it was recently demonstrated that, when engaged with substrates, the 20S undergoes a considerable conformational change, including the “gate” opening [14]. In contrast to our advanced understanding of 26S proteasome assembly, substrate selection, and proteolytic processes, the exact mechanisms controlling the cellular levels of free 20S proteasomes as well as their contribution to cellular proteolysis remain largely elusive

Large-scale changes in the proteome in response to environmental stressors require an enhanced proteolytic capacity of proteasomes to maintain protein quality and homeostasis. The inhibition of USP14/Ubp6 or phosphorylation of PSMD11/Rpn6 effectively increased proteasome activity and degradation of toxic proteins [15, 16]; however, numerous endogenous substrates were insensitive to these activation methods. The inhibition of mTOR, which mimics the nutrient-deprivation condition and upregulates cellular autophagic flux, appeared to affect the transcription and activity of 26S proteasomes, although the exact mechanism remains controversial [17]. In response to severe stress conditions, 26S proteasomes are regulated in a spatiotemporal manner (instead of undergoing energy-costly degradation and synthesis) and packaged into membrane-less organelles, such as proteasome storage granules (PSGs; for example, during energy depletion), aggresomes (catalytic inhibition), or nuclear foci (hypertonic stress) [18–20]. These proteasome inclusions are formed via reversible liquid-liquid phase separation (LLPS), allowing these condensates to re-dissipate into the nucleus or cytoplasm when the stress is removed [21]. This spatiotemporal regulation allows cells to rapidly adapt to proteostatic challenges, especially considering that the proteasome is a long-lived and abundant constituent of the cell [22, 23].

Nutrient deprivation triggers a transition of the cellular proteome from a proliferative state to a quiescent one. In yeast and plants, in response to glucose or carbon depletion, cellular proteasomes are condensed into PSGs near ER membranes [24]. In mammalian cells, however, the formation of PSGs is rarely reported and little is known about cellular proteasomal responses to metabolic challenges. In the current study, we discovered an unexpected adaptive response of 26S proteasomes to prolonged glucose depletion: the disassembly of 26S proteasomes into intact 20S and 19S subcomplexes. Using biochemical and proteomic approaches, we found that, during this conformational transition, the interaction between the 26S proteasome and a proteasome adaptor protein ECPAS/Ecm29 [25–27] was significantly shifted from the 20S to the 19S, triggering the disassembly. Consequently, deletion of *ECPAS* effectively abrogated the 20S-19S disassembly. Computational prediction analysis using AlphaFold and our *ab initio* docking algorithm (GalaxyTongDock) produced two distinct binding models of ECPAS to the assembled and disassembled states of proteasomes. We also demonstrated that free 20S generated under glucose-depleted conditions facilitated the degradation of misfolded proteins in a Ub-independent manner and prevented ER stress-induced cellular dysfunction and apoptosis. Finally, using a mouse xenograft model, we found that elevated levels of 20S proteasomes correlated with the reduced levels of glucose in tumor tissues. These findings suggest an essential pro-survival role for 26S proteasome disassembly that allows cells to cope with excess amounts of misfolded and potentially proteotoxic proteins in response to glucose starvation conditions.

## Results

### Dissociation of the 26S/30S proteasome into subcomplexes in response to glucose depletion

To elucidate the effect of different stress conditions on proteasomes, we tagged a 20S subunit PSMB2/μ4 and a 19S subunit PSMD14/Rpn11 with TEV protease-cleavable hexahistidine-biotin (HB), which allowed a rapid affinity-purification of human proteasomes using streptavidin beads (Figure 1a) [28]. This method enabled the efficient isolation not only of proteasome holoenzymes (26S and 30S) but also of their subcomplexes (20S and 19S, respectively) from HEK293 cells stably expressing one of these tagged subunits. We observed that, in cells cultured in glucose-depleted media for 12 h (final glucose concentration < 0.2 mg/dL with serum supplementation, compared with regular media containing > 450 mg/dL glucose), the population of purified PSMB2-HB proteasomes was markedly shifted from the 26S/30S composition to the 20S proteasome, using non-denaturing (native) electrophoresis and subsequent in-gel fluorogenic substrate (suc-LLVY-AMC) hydrolysis (Figure 1b). Silver staining and immunoblotting analysis following SDS-PAGE of proteasomes isolated using this tagged PSMB2-HB subunit showed that the protein levels of 19S base and lid components (PSMC2/Rpt1 and PSMD14, respectively) and 19S-interacting proteins, such as VCP/p97 and USP14/Ubp6, were significantly reduced during glucose starvation, while those of 20S proteasomes were comparable (Figure 1c). A similar compositional change in proteasomes was observed in multiple cell lines with different HB-tagged 20S subunits (e.g., A549-PSMB5-HB cell line; Supplementary Figures 1a and 1b). These results suggest that the disassembly of 26S proteasomes into the 20S and 19S is a prevalent response mechanism in glucose-starved mammalian cells.

**Figure 1.**
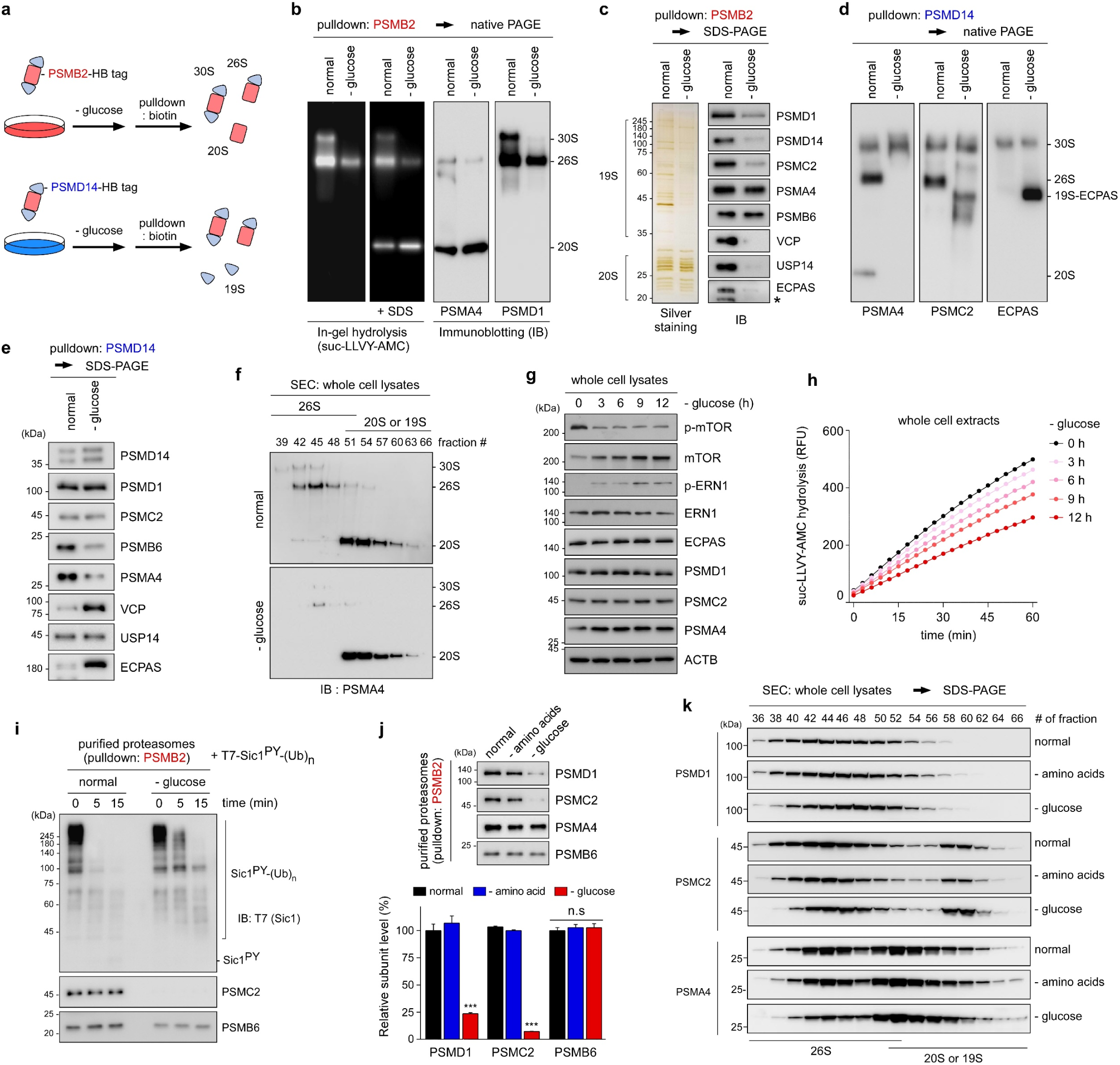
Glucose depletion leads to the disassociation of 20S and 19S subcomplexes. (**a**) Schematic representation of 26S proteasome-affinity tags and purification schemes used in this study. HEK293 cells stably expressing hexahistidine-biotin (HB)-tagged PSMB2/β4 or PSMD14/Rpn11 were treated with glucose-depleted media for subsequent proteasome affinity-purification. (**b**) Glucose depletion for 12 h triggered the dissociation of 26S proteasomes. Purified PSMB2-HB proteasomes were resolved by non-denaturing (native) polyacrylamide gel electrophoresis (PAGE), followed by visualization using in-gel suc-LLVY-AMC hydrolysis (*left* panels) and immunoblotting (IB; *right* panels). The addition of 0.02% SDS activated the 20S proteasome. (**c**) As in (b), except that the proteasomes were analyzed using denaturing SDS-PAGE followed by silver staining (*left*) and IB using various antibodies against the 20S, 19S subunits, and 19S-interacting proteins (*right*). *, nonspecific signals. (**d** and **e**) As in (b) and (c), respectively, except that 19S-tagged PSMD14-HB proteasomes were analyzed. (**f**) Whole-cell lysates (WCLs) were analyzed using size-exclusion chromatography (SEC), and each fraction was subjected to native PAGE/IB. (**g**) Time-dependent changes of the endogenous proteins in HEK293 cells during glucose depletion, analyzed by SDS-PAGE/IB. (**h**) Proteasome activity was monitored through hydrolysis of the suc-LLVY-AMC using WCLs under normal and glucose-depleted conditions. A representative result from these independent experiments is shown. (**i**) Polyubiquitinated Sic1 (Sic1^PY^-Ub_n_) protein was incubated with purified PSMB2-HB proteasomes, and its degradation was examined using SDS–PAGE/IB. (**j**) The subunits of purified proteasomes were compared after depriving of amino acids or glucose from the media for 12 h using SDS-PAGE/IB (*upper*). For quantification, signals from PSMD1, PSMC2, and PSMB6 were normalized to those of PSMA4 (*lower*). Bars represent the mean ± s.d. of three independent experiments (N = 3), *** p < 0.001 (one-way ANOVA followed by Bonferroni’s *post hoc* test). n.s., not significant. (**k**) SEC analysis after 12 h of depletion of amino acids or glucose was performed and each fraction was subjected to SDS-PAGE/IB for proteasome subunits.

We next performed parallel experiments using 19S-tagged PSMD14-HB proteasomes. This approach also showed a significant loss of interaction between the 20S and 19S after glucose deprivation (Figures 1d and 1e), consistent with the previous observations using PSMB2-HB proteasomes. In both cases, the disassembled 20S and 19S subcomplexes appeared to retain their structural integrity (Figures 1b and 1d). To monitor the 26S disassembly in whole-cell lysates (WCLs), the samples were gently fractionated by size-exclusion chromatography. Using subsequent native PAGE, we observed that upon glucose depletion, a significant portion of 19S was disassembled from the 26S/30S species (Figure 1f). The glucose starvation-mediated 26S disassembly occurred in a time-dependent manner over a 12-h time period (Supplementary Figures 1c and 1d), possibly reflecting the gradual reduction of intracellular energy states. At the same time, the abundance of individual 26S proteasome subunits in WCLs and their mRNA levels were not affected during the starvation period (Figure 1g and Supplementary Figure 1e). We also observed a rapid decrease in mTOR phosphorylation (starting at ∼3 h post-glucose depletion) and a gradual increase in ER to nucleus signaling 1 (ERN1/IRE1) phosphorylation (peaking at ∼ 9 h) (Figure 1g), corresponding to the inhibition of the mTOR signaling pathway and activation of unfolded protein responses (UPR), respectively. Proteasomes in the starved cells were largely dispersed throughout the cells without forming noticeable foci such as PSGs (Supplementary Figures 1f and 1g). Upon re-incubation of starved cells with a glucose-containing medium, the isolated PSMB2-HB and PSMB5-HB-proteasomes from HEK293 and A549 cells, respectively, were retrieved as the 26S form (Supplementary Figure 1h), indicating that this conformational change is a reversible process influenced by the intracellular glucose concentration.

Next, we evaluated the total activity of proteasomes in WCLs using suc-LLVY-AMC, a substrate considered to be more specific toward the chymotrypsin-like PSMB5 activity but generally accepted to represent overall proteasome activity. We observed that total cellular proteasome activity progressively decreased with glucose deprivation (Figure 1h). Generally, the specific peptidase activity of the 20S is lower than that of the 26S proteasome [14], consistent with a transition from 26S to 20S species during glucose deprivation. Furthermore, the purified PSMB2-HB proteasomes also showed considerably lower peptidyl substrate hydrolytic activity in response to glucose deprivation (Supplementary Figure 1i). When we incubated these purified proteasomes with polyubiquitinated Sic1 containing PY motif (Ub-Sic1^PY^), which is a more physiological substrate than fluorogenic peptides [29, 30], the degradation of Ub-Sic1^PY^ was significantly delayed in the PSMB2-HB proteasomes purified during glucose starvation compared to that in control proteasomes (Figure 1i). We note that the 26S proteasome disassembly was not observed in response to amino acid deprivation during the same time points (Figure 1j and 1k), consistent with the current consensus that amino acid depletion induces bulk autophagy without affecting the levels of proteasomes [6]. Therefore, our data indicate that the mechanism underlying the glucose-sensitive 26S proteasome assembly/disassembly is different from the cellular adaptation responding to amino acid depletion.

### Identification and quantification of changes in proteasome composition and interacting proteins during glucose depletion

To explore the molecular mechanism underlying 26S/30S proteasome disassembly during glucose depletion, we performed a mass spectrometry (MS) analysis with an intensity-based absolute quantification (iBAQ) algorithm, which allows precise protein quantification without labeling [31]. HEK293-PSMB2-HB stable cells were cultured under normal and glucose-starvation conditions for 12 h, and proteasomes and their interacting proteins were purified using streptavidin pulldowns for the iBAQ-MS analysis (Figure 2a). Using our cutoff criteria (threshold at 2-fold change in absolute quantity and adjusted *p*-value < 0.05; see *Methods* for details), we identified a total of 206 differentially associated proteins (DAPs). Among them, 141 proteins had significantly reduced binding affinity towards PSMB2-HB proteasomes, while 65 had increased affinity after 12 h of glucose deprivation (Figure 2b and Supplementary Tables 1–4). Our quantitative DAP profile indicated that, under glucose-depleted conditions, the levels of constituent 19S subunits were markedly decreased to ∼ 20 % of the normal levels (Figures 2c and 2d), substantiating our biochemical findings (Figure 1). The levels of proteasome activators (PSME1–4), were unchanged (Supplementary Tables 2), suggesting that hybrid proteasomes such as the 11S-20S complex are not significantly involved in proteasome dynamics during glucose depletion.

**Figure 2.**
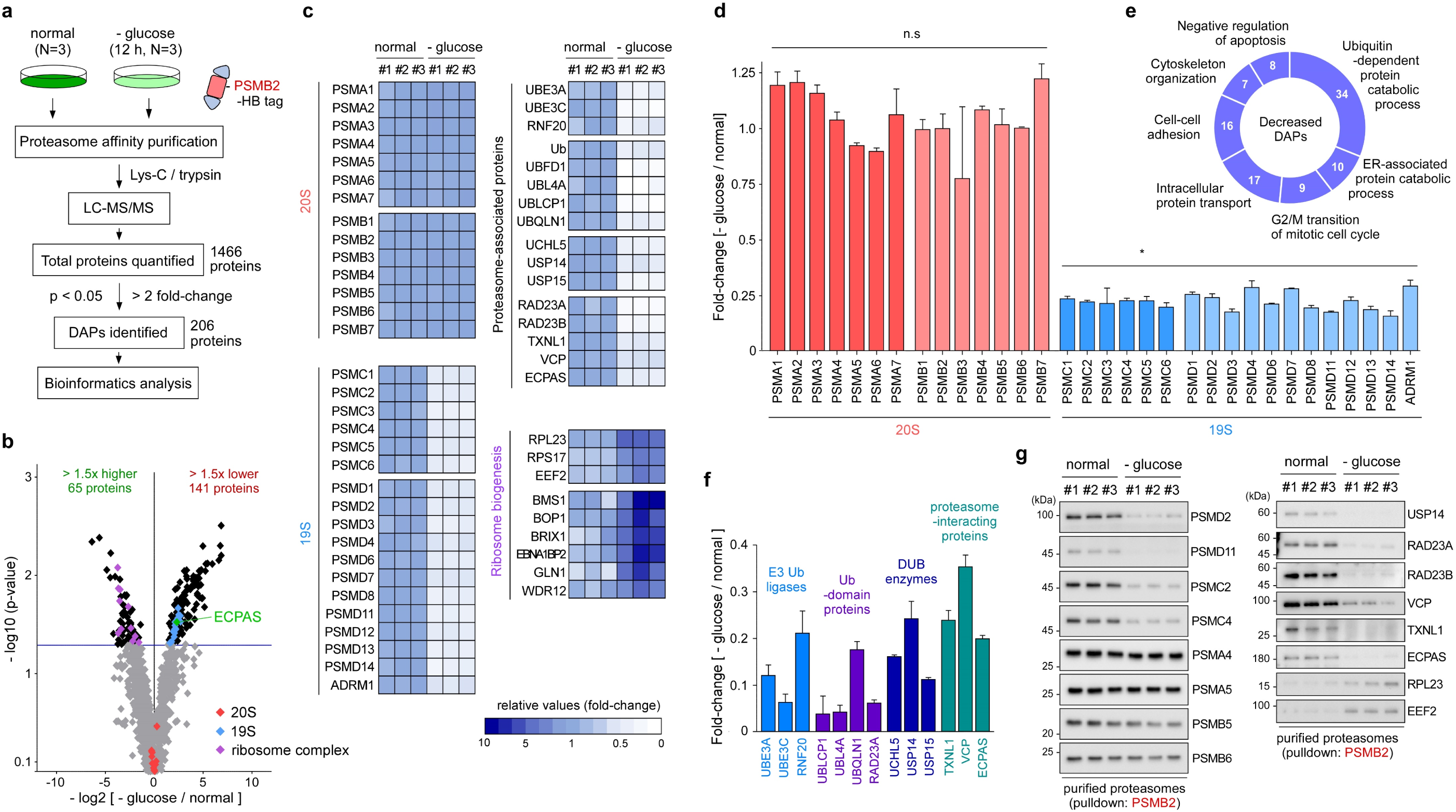
Analysis of proteasome disassembly using quantitative mass spectrometry (MS) (**a**) Overview of quantitative MS analysis with the label-free, intensity-based absolute quantification (iBAQ) method, using affinity-purified PSMB2-HB proteasomes. A total of 206 differentially associated proteins (DAPs) were identified in the PSMB2-HB proteasomes after 12 h of glucose depletion. (**b**) Volcano plot of the total identified proteins. The blue line shows the threshold *p*-value (*p* < 0.05). Non-DAPs are shown in gray (*p* > 0.05 or < two-fold change in log2 ratios of glucose starvation to normal conditions). Red, blue, and black circles depict unchanged 20S subunits, upregulated 19S subunits, and downregulated ribosome complex subunits, respectively. (**c**) The heat map shows distinct changes in the relative abundance of proteasome subunits, proteasome-associated proteins, and ribosome biogenesis-related proteins. Degrees of upregulation (darker blue) and downregulation (lighter blue to white) are explained in the color key. (**d**) Relative abundance changes of the constitutional subunits of 26S proteasomes. Data represented include the mean ± s.d. of three independent experiments (N = 3), \**p* < 0.05 (one-way ANOVA with Bonferroni’s multiple comparison test). n.s., not significant. (**e**) Classification of downregulated DAPs into different biological pathways using gene ontology analysis. Of the 141 DAPs, 34 proteins were enriched in Ub-dependent proteolysis, 17 in intracellular protein transport, and 10 in ER-associated proteolysis. (**f**) Representative decrease in the quantity of proteasome-associated proteins, including E3 ubiquitin (Ub) ligases and deubiquitinating (DUB) enzymes, following glucose depletion. (**g**) Validation of DAPs using IB analysis against the 20S, 19S, and proteasome-associated proteins.

Gene ontology analysis revealed that DAPs with decreased affinity were clustered within several biological processes, such as Ub-dependent protein catabolism, ER-associated protein catabolism, intracellular protein transport, and negative regulation of apoptosis (Figure 2e). As expected, in response to stress, the majority of 19S-interacting proteins were also lost from affinity-isolated proteasome complexes, including E3 Ub ligases (UBE3A, UBE3C, and RNF20), Ub and Ub-domain proteins (RAD23A, UBFD1, UBL4A, UBLCP1, and UBQLN2), deubiquitinating enzymes (UCHL5, USP14, and USP15), and other known proteasome-associated proteins (TXNL1, VCP, and ECPAS) [32] (Figure 2f and Supplementary Table 3). In sharp contrast, another set of DAPs had increased association with PSMB2-HB proteasomes in response to glucose starvation (Figures 2b and 2d). Notably, we found that many RNA-binding and ribosomal proteins associated directly with the 20S proteasome were significantly higher in the glucose-depleted group than in the control groups (Figures 2b, 2c, and Supplementary Figures 2a – 2c). These findings suggested that under stress conditions, proteasome disassembly may affect not only the degradation of existing proteins but also *de novo* protein synthesis, reshaping the global proteome [33]. Next, we performed immunoblotting to validate our proteomics data, using PSMB2-HB proteasome and found that the level changes in co-purified proteins, such as downregulated RAD23A/B, VCP, and ECPAS, and upregulated RPL23 and EEF2) were consistent with the results of iBAQ-MS analysis (Figure 2g and Supplementary Figure 2d). Taken together, our label-free quantitative MS analysis demonstrated that in response to glucose depletion, the 26S proteasome holoenzyme effectively dissociates into the 20S and 19S, together with a significant interactome change in numerous proteasome-associated proteins.

### ECPAS-dependent regulation of human 26S/30S proteasomes upon glucose depletion

One possible explanation for 26S proteasome dissociation under glucose-depleted conditions could be insufficient ATP hydrolysis by the PSMC AAA+ ATPase ring, which is required for strengthening the 20S-19S interaction. However, we observed that, even after glucose starvation for 12 h, cellular ATP levels decreased only to ∼70–80 % of control levels (Supplementary Figure 3a). Furthermore, the treatment of cells with proteasome inhibitors known to stabilize the interaction between the 20S and 19S without requiring ATP hydrolysis [34] potently delayed proteasome disassembly (Supplementary Figures 3b and 3c). These results suggested that the observed proteasome remodeling was mediated by a regulated process rather than ATP deficiency. Based on our proteomics data (Figures 2b, 2c, and 2f), we postulated that the proteasome adaptor protein ECPAS may play a role in 26S disassembly. Ecm29, the yeast homolog of ECPAS, was initially identified as a salt-sensitive proteasome component that stabilizes the 26S/30S proteasomes in yeast [25]. However, in mammalian cells, ECPAS was reported to induce the dissociation of the holoenzymes in response to oxidative stress [35], although its physiological function remains largely obscure. Our purification system showed that ECPAS was co-purified with both PSMB2-HB and PSMD14-HB proteasomes under normal conditions, while it was bound only to PSMD14-HB complexes but not to PSMB2-HB under glucose-depleted conditions (Figures 1c, 1d, 1e, and 3a). Native PAGE analysis showed that ECPAS co-migrated with intact 19S complexes under glucose-depleted conditions (Figure 1d), further demonstrating that ECPAS-proteasome binding shifted from the 26S to the 19S, depending on glucose availability.

**Figure 3.**
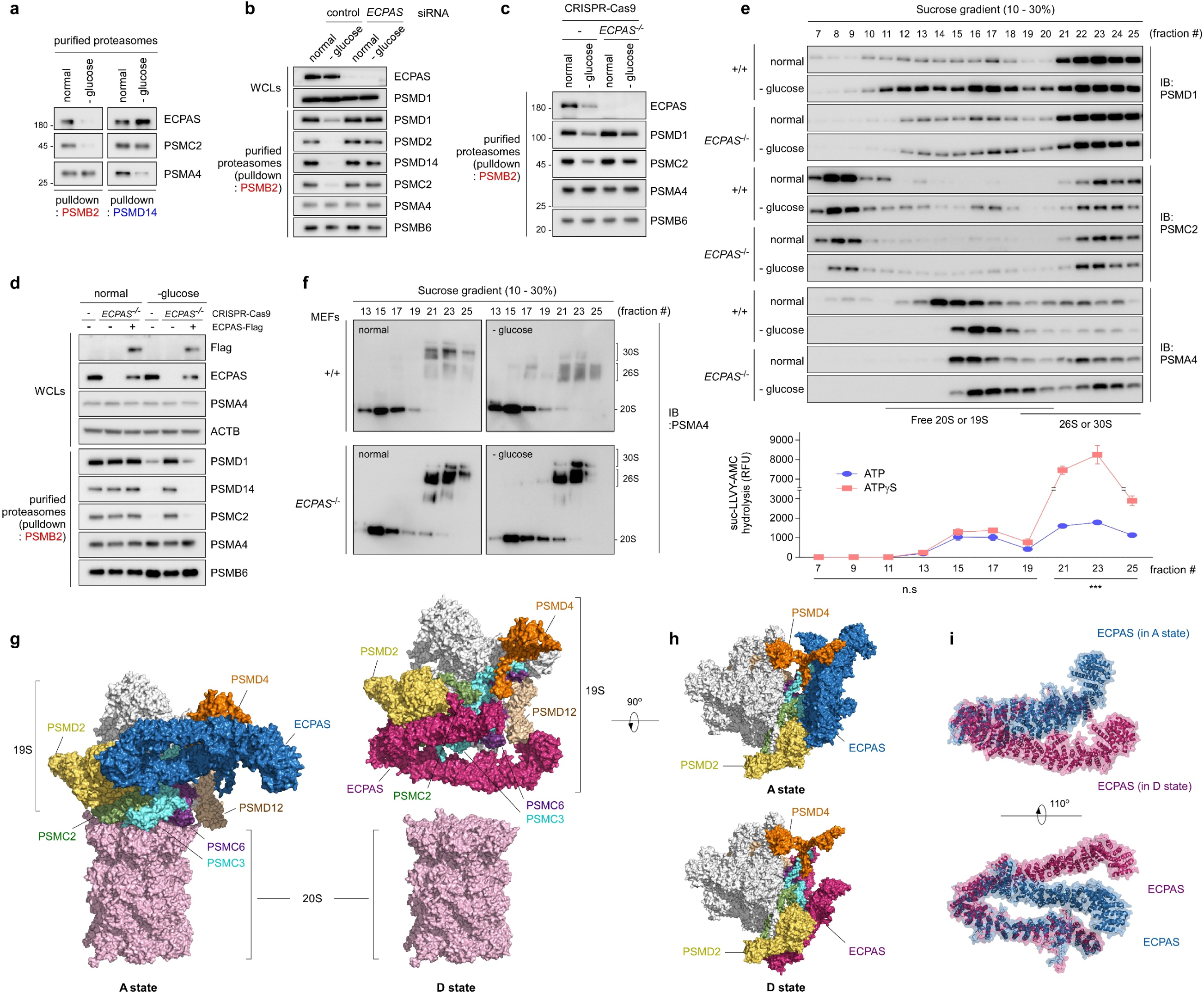
ECPAS/Ecm29 mediates 26S/30S proteasome disassembly in response to glucose depletion. (**a**) Direct interactions between endogenous ECPAS and purified PSMB2-HB (*left*) or PSMD14 (*right*) proteasomes under normal conditions are shown. Cellular proteasomes were pulled down using streptavidin beads under normal and 12 h glucose-depleted conditions and analyzed by SDS-PAGE/IB using anti-ECPAS antibodies. Glucose depletion induced a strong dissociation of ECPAS from the 20S proteasomes but resulted in a tighter binding to the 19S complex than did normal conditions. (**b**) HEK293-PSMB2-HB cells were transfected with scrambled (control) or *ECPAS* siRNA for 48 h and then cultured in glucose-deprived media for an additional 12 h. Purified PSMB2-HB proteasomes were subjected to SDS-PAGE/IB using indicated antibodies. (**c**) As in (b), except that the experiments were conducted with *ECPAS* knockout (HEK293-PSMB2-HB*^ECPAS-/-^*) cells generated by the CRISPR/Cas9 system. (**d**) Adding back ECPAS to the knockout cells. HEK293-PSMB2-HB*^ECPAS-/-^* cells were transfected with ECPAS-expressing plasmids for 36 h and then incubated in glucose-depleted media for 12 h. PSMB2-HB proteasomes were affinity-purified, separated using SDS-PAGE, and then probed with antibodies against the 20S or 19S subunits. (**e**) Sucrose-gradient ultracentrifugation of WCLs from wild-type and *ECPAS*^-/-^ mouse embryonic fibroblasts (MEFs) grown in standard or glucose-depleted media for 12 h. (*Upper*) Fractionated samples were analyzed by SDS-PAGE/IB. (*Lower*) The fractions containing 26S/30S proteasomes showed significantly elevated (∼8-fold) suc-LLVY-AMC hydrolysis in the presence of ATPγS. (**f**) As in (e), except the samples were analyzed by native PAGE/IB. (**g** and **h**) Modeled conformations of ECPAS-26S proteasome complex predicted by AlphaFold and GalaxyTongDock (an *ab initio* docking algorithm), viewed from the side (g) and top (h). Two distinct conformations were generated and designated as the “assembled” (A) and “disassembled” (D) states, in the latter of which ECPAS sterically obstructs the 19S-20S interface. (**i**) Structural comparison of the ECPAS structures in the A and D states (shown in cobalt and magenta, respectively). The modeled structures are superimposed with the coincided N-terminal portions and are shown from different perspectives

The results presented above indicated that ECPAS was potentially involved in 20S-19S disassembly; therefore, to confirm these findings, we investigated the role of ECPAS using loss-of-function experiments. The knockdown and knockout of *ECPAS* in HEK293-PSMB2-HB cells using siRNA and the CRISPR/Cas9 system, respectively, stabilized PSMB2-isolated 26S/30S proteasomes even under glucose-depleted conditions (Figures 3b and 3c). To further verify the involvement of ECPAS in 26S proteasome remodeling, HEK293-PSMB2-HB*^ECPAS^*^-/-^ cells were transfected with ECPAS, which rescued the biochemical phenotype of 26S proteasomes during glucose starvation (Figure 3d). The association/dissociation between the 20S and 19S subcomplexes was also analyzed in WCLs from wild-type (+/+) and *ECPAS*^-/-^ mouse embryonic fibroblasts (MEFs) using sucrose-gradient ultracentrifugation. In contrast to +/+ MEFs, glucose starvation in *ECPAS*^-/-^ MEFs did not significantly change the levels of proteasome subunits in the earlier fractions containing free 20S or 19S complex (Figure 3e). The fractionated samples were further analyzed using native PAGE; the results showed that, after glucose deprivation, the proportion of 26S/30S proteasomes in *ECPAS*^-/-^ MEFs was significantly increased compared with that in +/+ MEFs (Figure 3f). In addition, we found that following carbon source starvation or during the stationary phase, yeast proteasomes in an Ecm29Δ strain consistently had a higher ratio of 26S/30S configuration to free 20S than did those from the control strain (Supplementary Figure 3d). The phenotypic similarity between yeast and mammalian proteasomes suggests that the 26S proteasome dissociation mediated by ECPAS/Ecm29 is evolutionarily conserved in eukaryotes, possibly as a global protein quality control mechanism during glucose starvation.

To gain mechanistic insight into the ECPAS-mediated disassembly of 26S proteasomes, we performed *in silico* modeling to determine the structure of the ECPAS-26S proteasome complex. The free ECPAS structure (UniProt ID: Q5VYK3) was predicted by AlphaFold to have a horseshoe-shaped fold with a high confidence score (median pLLDDT = 88.4 from residues 1 to 1,845) (Supplementary Figure 3e). The overall morphology was similar to yeast Ecm29 structures obtained by the Finley group using electron microscopy ∼20 years ago [25]. The AlphaFold-predicted structures of yeast Ecm29 and human ECPAS were highly analogous, even though the amino acid identity between them is only 25.5%. The two segments showing high sequence similarity between yeast Ecm29 and human ECPAS were located in the two stretched branches with many HEAT repeats. On the contrary, the angled region between them, located at the residues 1,048–1,148, had low confidence scores (median pLDDT = 65.89). To predict the interaction between ECPAS and the 26S proteasomes (PDB: 6MSB), we employed an *ab initio* protein-protein docking algorithm termed GalaxyTongDock (see *Methods* for details) and obtained two distinct complex models (Figures 3g and 3h), which may represent the “assembled” and “disassembled” conformations between the 20S and 19S subcomplexes.

In the “assembled” states, the N-terminal shank of ECPAS directly interacted with the proteasome-residing Ub receptors, such as PSMD2/Rpn1 and PSMD4/Rpn10, on the face of the 19S (Figures 3g and 3h). The Ub-interacting motifs of PSMD2 and PSMD4 were excluded from the ECPAS-19S interaction, ruling out the potential inhibitory effect of ECPAS binding on normal proteasome functions. In the “disassembled” state, the horseshoe curvature at the low pLDDT region of ECPAS was noticeably altered while the overall conformational landscape of ECPAS was only slightly changed (Figure 3i). Simultaneously, the C-terminal region of ECPAS in this conformation protruded toward the hexameric PSMC ring/heptameric PSMA ring interface, potentially obstructing their association. Therefore, the conformational change of ECPAS might be the primary regulatory checkpoint for the disassembly of the 26S upon glucose starvation. These results imply that our integrative *in silico* modeling algorithm with a predictive power of AlphaFold can predict a highly plausible biochemical mechanism (e.g., the dissociative role of ECPAS in the 20S and 19S), complement to conventional approaches. However, how ECPAS structures are changed and what signaling pathways regulate this process remain to be elucidated.

### Facilitated degradation of misfolded proteins by 20S proteasomes during glucose depletion

The 20S proteasome proteolyzes fully or partially disordered substrates in a Ub- and ATP-independent manner. To investigate the contribution of disassembled 20S proteasomes under glucose starvation conditions to the removal of misfolded proteins, we first used puromycin, which incorporates nascent polypeptides into their C-termini, generating inherently disordered puromycylated polypeptides [36]. In +/+ MEFs and A549 cells, we observed that puromycylated polypeptides were degraded more rapidly in glucose-depleted media than in control media (Figures 4a, 4b, Supplementary Figures 4a, and 4b). This preferential removal of the polypeptidyl-puro species was effectively abrogated by treatment with the proteasome inhibitor MG132, but was virtually unaffected by other inhibitors, such as bafilomycin A1 (autophagy/lysosomal inhibitor) or MLN7243 (E1 Ub-activating enzyme inhibitor) (Figure 4a and Supplementary Figure 4a). Independently, we observed the induction of autophagy-related factors during glucose starvation (Supplementary Figures 1e and 4d); however, bafilomycin A1 treatment had little effect on the degradation of polypeptidyl-puro species (Figure 4a and Supplementary Figure 4a). These results indicated that lysosomal targeting or Ub modification is not the major clearance route for nascent misfolded proteins at least during glucose-deficient stress. In contrast to the results above, the steady-state levels of puromycylated polypeptides in *ECPAS^-/-^* MEFs were largely comparable between normal and glucose-depleted conditions. At the same time, their levels in the knockout cells substantially increased in response to MG132 and MLN7243, but not bafilomycin A1 treatment (Figure 4a). These observations are in line with a role of free 20S in the removal of misfolded proteins under stress conditions. Cycloheximide chase experiments, conducted after a 2 h puromycin labeling, also showed that the degradation of puromycylated polypeptides in *ECPAS^-/-^* MEFs was significantly delayed, probably due to the impaired 20S-19S disassembly (Figure 4b and Supplementary Figure 4b). These results indicate that free 20S proteasomes play a key role in the clearance of nascent misfolded proteins under glucose-deficient conditions.

**Figure 4.**
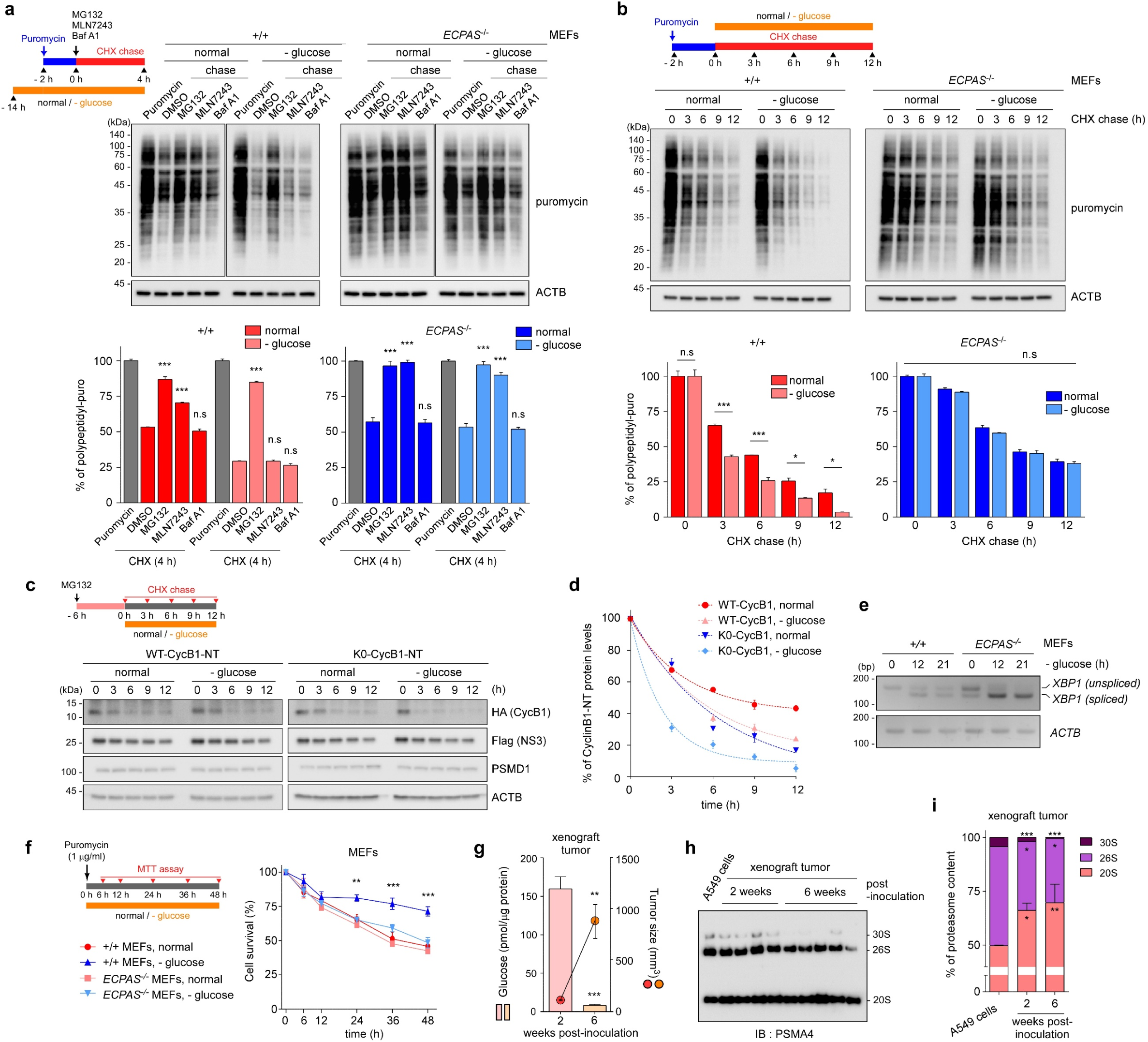
Free 20S proteasomes facilitate Ub-independent substrate degradation and contribute to cell survival under glucose-depletion. (**a**) Wild-type (+/+) and *ECPAS*^-/-^ MEFs were cultured for 12 h under normal and glucose-depletion conditions, followed by a 2 h puromycin (5 μg/mL) treatment and a subsequent 4 h cycloheximide (CHX; 80 μg/mL) chase experiment. The CHX chase was performed in the presence of DMSO, the proteasome inhibitor MG1312 (10 μM), the E1 Ub-activating enzyme inhibitor MLN7243 (1 μM), or the autophagic flux inhibitor bafilomycin A1 (BafA1; 100 nM). SDS-PAGE/IB for puromycylated polypeptides (*upper*); their quantity was normalized to that of endogenous ACTB/β-actin (l*ower*). Shown are the average percentage of remaining polypeptidyl-puro (mean ± s.d) from three independent experiments (N = 3), *** *p* < 0.001 (one-way ANOVA followed by the Bonferroni *post hoc* test). (**b**) As in (a), except that the chase experiment was performed for the various time periods. * *p* < 0.05, *** *p* < 0.001 (one-way ANOVA followed by the Bonferroni *post hoc* test; N = 3). (**c**) HEK293 cells were transfected with the wild-type and lysine-less mutant of cyclin B1 N-terminal segment (WT-CycB1-NT and K0-CycB1-NT, respectively) for 24 h, cultured in normal and glucose-depleted media for indicated periods, and prepared for IB analysis to examine CycB1-NT degradation. (**d**) Quantification of HA-CycB1-NT signals in (c), normalized to those of co-expressed flag-NS3. (**e**) mRNA levels of unspliced and spliced *XBP1* in +/+ and *ECPAS*^-/-^ MEFs after glucose starvation (12 h). Semi-quantitative RT-PCR (25 cycles) and quantitative normalization were performed using ACTB gene expression. (**f**) Cell survival upon puromycin exposure and glucose starvation. Cells were cultured in normal or glucose-depleted media in the absence or presence of puromycin (250 ng/mL) for up to 48 h, and their survival was quantified using an MTT assay. ** *p* < 0.01, *** *p* < 0.001 (one-way ANOVA followed by the Bonferroni *post hoc* test; N = 3). (**g**) Glucose was significantly depleted in late-stage tumors derived from mouse xenograft models. A549 cells (5 × 10^5^) in 0.1 mL PBS/matrigel were injected into the s.c. dorsa of NOD/SCID mice at the right-hind flank. Tumor tissues were harvested 2- and 6-week post-inoculation (N = 4 and 5, respectively). Glucose levels normalized to the total protein concentration were compared to tumor sizes. Data are presented as the mean ± s.e.m.; \*\**p* < 0.01 and \*\*\**p* < 0.001 (unpaired Student’s *t*-test). (**h** and **i**) Elevated levels of the 20S proteasomes under more glucose-depleted conditions. WCLs from cultured A549 cells and tumor tissues with different xenograft periods were subjected to native PAGE followed by IB against PSMA4 (h). The relative amounts of the 20S, 26S, and 30S proteasomes were quantified from (i). Data are presented as the mean ± s.d.; \**p* < 0.05, \*\**p* < 0.01, and \*\*\**p* < 0.001 (one-way ANOVA followed by the Bonferroni *post hoc* test).

Since both 26S and 20S proteasomes have common catalytic sites and mechanisms, the remaining 26S may affect the degradation of polypeptidyl-puro species. To confirm the contribution of 26S proteasome disassembly to Ub-independent proteolysis, we utilized lysine-less mutants (K0) of the intrinsically disordered N-terminal segment of cyclin B1 (CycB1-NT), which is shorter-lived than full-length CycB1 and directly proteolyzed by 20S proteasomes [14]. Transiently overexpressed K0-CycB1-NT was more rapidly turned over than wild-type counterparts (WT-CycB1-NT), while co-expressed NS3 levels were essentially unchanged (Figure 4c). Similar to the polypeptidyl-puro species, K0-CycB1-NT degradation was significantly accelerated under glucose-depleted conditions (5.2 h vs 2.4 h half-lives), while WT-CycB1-NT half-lives were modestly decreased (8.7 h vs 5.9 h half-lives; Figures 4c and 4d). mRNA levels of K0-CycB1-NT were little changed during glucose starvation (Supplementary Figure 4c), indicating that the reduction of this model 20S substrate was indeed post-translational. Collectively, these data strongly indicate that disassembled 20S proteasomes were more efficient at proteolyzing disordered proteins in a Ub-independent manner than 26S proteasomes. This property is expected to be a cell survival strategy in response to glucose starvation that restrains cellular protein folding capacity and produces more misfolded and unstructured proteins.

We further investigated whether the rapid removal of misfolded proteins by the 20S proteasome is advantageous for maintaining proteostasis and subsequently cell survival. Previous studies demonstrated that glucose depletion induced ER stress and the UPR [37, 38]. We observed that our stress condition (glucose deprivation for 12 h) also strongly increased the levels of ER stress markers and UPR signaling (Supplementary Figures 4d and 4e). To evaluate the role of disassembled 20S proteasome in response to ER stress, we first examined the ERN1-mediated splicing of X-box binding protein 1 (*XBP1)* mRNA, an evolutionarily conserved transcription factor required for the activation of the UPR [39]. RT-PCR analysis showed that glucose-starved *ECPAS*^-/-^ MEFs produced significantly more spliced *XBP1* mRNA than +/+ MEFs (Figure 4e). Additional quantitative RT-PCR analysis revealed the upregulation of numerous genes involved in the ATF4-CHOP axis in response to glucose depletion (Supplementary Figure 4f). Notably, the levels of these genes, which are involved in UPR-mediated apoptosis, were significantly higher in *ECPAS*^-/-^ MEF than in +/+ MEFs (Supplementary Figure 4f), indicating that the loss of *ECPAS* resulted in hypersensitivity to glucose depletion-mediated ER stress. When puromycylated polypeptides were overexpressed, we found that *ECPAS*^-/-^ MEFs had significantly higher cytotoxicity than did +/+ MEFs after prolonged glucose deprivation (Figure 4f). These results collectively indicate that proteasome disassembly upon glucose depletion is an effective compensatory strategy to degrade structurally disordered proteins and to provide a survival benefit to cells under ER stress.

Tumorigenesis is a pathological condition characterized by energy depletion and, in human tumor tissues, it is known that glucose levels are significantly lower than those in normal tissues [40]. When we examined mouse tumor tissues from various xenograft models in different stages, we found that glucose concentrations were drastically reduced in late-stage tumors than in early-stage tumors (159.6 ± 54.6 pmol/μg glucose concentration and 111.9 ± 5.6 mm^3^ tumor volume in 2-week post-inoculation samples vs. 8.1 ± 5.8 pmol/μg and 877.8 ± 368.9 mm^3^ in 6-week samples: Figure 4g). In addition, the 20S proteasome/total proteasome ratios were significantly higher in tumors (66.1 ± 3.4 % and 69.7 ± 8.6 % in 2- and 6-week samples, respectively) than in parental A549 cells (49.7 ± 0.3 %) and they were negatively correlated with the glucose concentration (Figure 4h and 4i). These data indicate that the 26S proteasome disassembly may confer glucose-deficient cells with the ability to degrade various substrates in a Ub-independent manner during tumorigenesis and, therefore, present novel therapeutic strategies for cancer. Tumor suppressor proteins such as p53 and retinoblastoma (Rb) are known to be degraded by this manner, increasing cell death resistance and cell cycle arrest [41]. Depletion of 26S or 30S holoenzymes accompanied by a concomitant increase in free 20S proteasomesmay be a mechanism to shift cellular metabolism to a less regulated but more essential form of intracellular proteolysis, potentially serving as a hallmark of cancer.

## Discussion

Intracellular proteolytic mechanisms have evolved to allow cell survival in unfavorable environments. To maintain protein quality control, as well as the cellular amino acid pool, it is essential to promptly adjust the overall output of cellular proteolysis in response to the excess amounts of to-be-degraded substrates. Here, we report that the 26S and 30S proteasomes are dissociated into the 20S and 19S subcomplexes during glucose deprivation and are reversibly re-associated when the stress is relieved (Figure 5). During the stress, proteolytic activities attributed to 20S proteasomes increased, suggesting that 26S disassembly is not necessarily a mechanism to attenuate proteolysis but rather to alter substrate specificity. ECPAS, more commonly known as Ecm29, showed increased affinity toward the 19S in the disassembly process and its deletion resulted in a delayed 26S disassembly process in mammals. Moreover, as a mechanistically similar process was observed in budding yeast—which is more sensitive to variations in glucose availability and even more tolerant to drastic reduction in its metabolic activity upon extreme starvation—the disassembly of 26S proteasome under glucose-starved conditions transpired to be an evolutionarily conserved mechanism necessary for global proteostasis and cell survival. Using *in silico* analysis, we propose a mechanism of conformational rearrangement of ECPAS, which may function as a necessary process for commencing the 26S/30S disassembly by allowing the disengagement of the PSMC ring from the PSMA ring.

**Figure 5.**
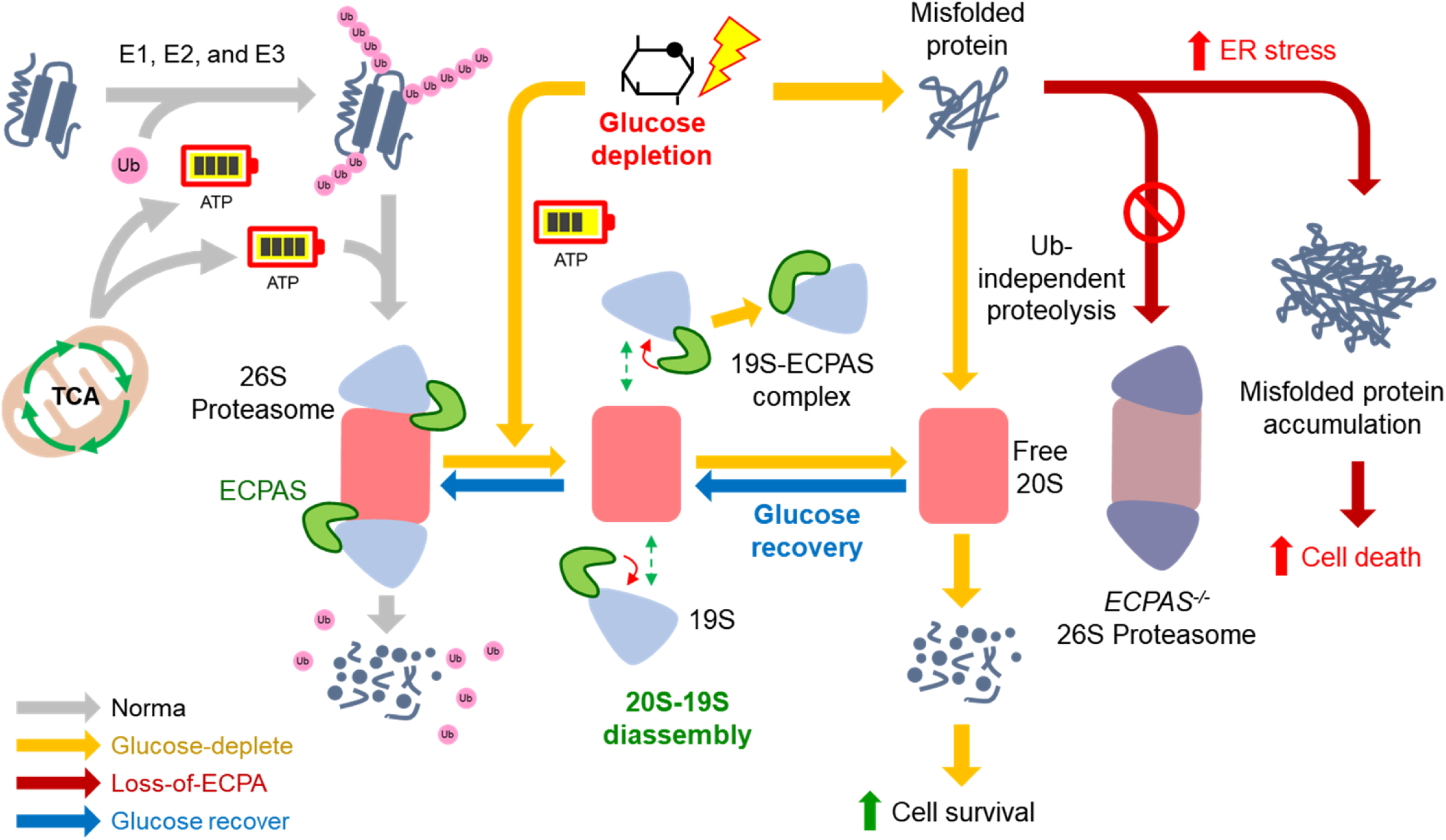
A Model of proteasome disassembly upon glucose starvation as an adaptive survival strategy. Under normal conditions, the 26S-30S proteasome degrades polyubiquitinated protein in a Ub- and ATP-dependent manner. When a cell experiences low levels of available glucose, the 26S-30S proteasome is disassembled into the 20S and 19S subcomplexes, mediated by the conformational changes of ECPAS. Elevated levels of the free 20S proteasome facilitate the Ub-independent proteolysis of misfolded proteins, a strategy that may improve cell survival by alleviating ER stress and proteopathic stress.

When cellular proteostasis is substantially perturbed by chronic stressors, such as starvation, oxidation, and proteasome inhibition, 26S proteasomes are often sequestered into various areas of the cell, such as the cytoplasm (e.g., PSGs), nucleus (starvation-induced proteasome assembly in the nucleus, or SIPAN), and perinuclear regions (aggresomes). These LLPS-mediated proteasome inclusion bodies are generally believed to be cytoprotective: either the impaired proteasomes are temporarily reserved or the concentrated proteasomes are more efficient at eliminating excess aberrant proteins. Intriguingly, the formation of PSGs is hardly observed in mammals upon glucose depletion unlike in yeast or plants (our observations and [24]). Considering the high concentration of proteasomes in cells and the energy-costly *de novo* proteasome synthesis, it seems plausible that a large share of cellular proteasomes cannot be entirely sequestered in response to various stress conditions. Thus, 26S disassembly into the 20S and 19S proteasomes can provide a more rapid and pervasive adaptive cellular mechanism. It is possible that the disassembly process precedes autophagy induction under starvation conditions and the free 20S and autophagosomes largely have exclusive substrates. The two mechanisms are expected to complement each other at the later stage of stress responses: when total 26S proteasome activity is suppressed through sequestration or disassembly, induced autophagy will provide cells with energy and anabolic intermediates. When the stress is relieved, the 20S and 19S subcomplexes will efficiently re-associate for a highly controlled ATP-dependent and Ub-dependent proteolysis.

Eukaryotic cells contain a mix of 20S, 26S, and 30S proteasomes, while free 20S accounts for more than half of the total proteasomes [11, 42]. In addition, more ∼20% of cellular proteins are consistently degraded by the 20S proteasome [43]. However, it was originally believed that the 20S was an intermediate of the 26S proteasome assembling process or a latent enzymatic complex, which was released from the holoenzyme when ATP levels decreased [44–47]. The lack of an ATPase adaptor or Ub receptor has also concealed the proteolytic role of free 20S proteasomes; however, recent studies clearly demonstrated the functional importance of the 20S as a stand-alone protease through biochemical, structural, and mass-spectrometric analysis [48]. Archaea and certain bacteria contain a primordial form of the 20S-like protease complex that is only loosely associated with ATPase regulators and may also function independently [49–51]. We postulate that substrate hydrolysis at the proteolytic chamber may lead to conformational changes as observed in the ClpP proteases (by the local pH drop in the lumen) [52]. This cycle confers a critical positive feedback for processive substrate degradation under stress conditions. Our current study supports the previous observations that such primordial proteolytic activity may be carried over into eukaryotic 20S proteasomes under certain stress conditions.

It is conceivable that the structural remodeling of 26S proteasomes under glucose starvation conditions has strong pathological implications. During glucose depletion, it was mainly the free 20S proteasome, rather than the 26S or autophagy, that was responsible for the clearance of translationally misfolded proteins. The 20S proteasome more readily processes excessively produced misfolded proteins in a Ub-independent manner. In this context, one may consider the 20S as a “first-responder” or “emergency proteasome” to elevated levels of aggregate-prone misfolded polypeptides. Therefore, our data, together with those of previous findings, provide a framework for a novel and universal proteasome quality control mechanism, suggesting that maintaining a functional 20S proteasome pool in the cell confers pro-survival properties in response to numerous proteopathic challenges. Chronic stress, which impairs the efficient 20S-26S proteasome conversion, may gradually lead to the development of proteopathies over time. It has been known for decades that bortezomib and other proteasome-targeting anticancer drugs initiate severe UPR or ER stress; however, the mechanisms underlying the anti-tumor effects are not completely understood. Based on our observations, we propose that a significant portion of these therapeutic effects may result from stabilizing the 26S holoenzyme complex and, subsequently augmenting ER stress-mediated apoptosis.

## Methods

### Cell culture and treatments

Mammalian cells used in this study, including HEK293, HEK293-PSMB2-HB, HEK293-PSMD14-HB, HEK293-PSMB2-HB-ΔECPAS, HeLa, +/+ MEFs, and *ECPAS*^-/-^ MEFs [53] were cultured in DMEM, whereas A549 and A549-PSMB5-HB cells were grown in RPMI-1640 media. All culture media supplemented with heat-inactivated 10% fetal bovine serum, 100 units/mL penicillin/streptomycin, and 2 mM L-glutamine. Cells were routinely tested for mycoplasma contamination. For glucose depletion, cells were washed twice with glucose-free DMEM and cultured in glucose-free DMEM supplemented with 10% dialyzed FBS (A3382001, Gibco), 100 units/mL penicillin/streptomycin, and 2 mM L-glutamine. All the reagents were treated to the cells as below unless otherwise described: MG132 (10 μM; M-1157, AG scientific), bafilomycin A1 (100 nM; 11038, Cayman Chemical), MLN7243 (1 μM, 30108, Cayman Chemical), Cycloheximide (80 μg/mL; 14126, Cayman Chemical), and puromycin (250 ng/mL to 10 μg/mL; 13884, Cayman Chemical).

### Generation of retrovirus for protein expression and stable cell lines

Stable cell lines such as A549-PSMB5-HB for proteasome purification were generated by retrovirus transduction and antibiotics selection, largely as previously described [54]. Briefly, plasmids expressing pQCXIP-PSMB5-HB and VSV-G were co-transfected into a 293 GP2 cell line. After transfection for 48 h, the medium containing the retrovirus was collected and mixed with 5 μg/mL polybrene to transduce A549 cells and then gradually selected with puromycin (1 to 3 μg/mL) to establish stable cell lines.

### Establishment of HEK293-PSMB2-HB*^ECPAS^*^-/-^ cell lines

sgRNA sequences targeting *ECPAS* (5′-GCACAGCGAACTCTCATGGAGG-3′) were identified using the CRISPR RGEN Tools (http://www.rgenome.net/) by screening exonic regions of *ECPAS* (RefSeq: NC_000009.12). The sgRNAs with overhangs for BbsI sites were annealed using two primers, 5′-caccgGCACAGCGAACTCTCATGG-3′ and 5′-aaacCCATGAGAGTTCGCTGTGCc-3′, and ligated into digested pSpCas9(BB)-2A-GFP with BbsI by T4 DNA ligase. HEK293-PSMB2-HB cells were transfected with 5 μg of the cloned plasmid using Lipofectamine 3000 (Invitrogen). After 48 h incubation, cells were trypsinized, and GFP-positive cells were sorted by fluorescence-assisted cell sorting (FACS, BD AriaIII). To isolate single clones, FACS-sorted cells were seeded in 96-well culture plates with low density (∼ 0.5 cells per well), and the knockout efficiency of each clone was later confirmed by genotyping with Sanger sequencing and immunoblotting against ECPAS.

### Purification of human proteasomes

Human proteasomes were isolated by affinity purification from a stable HEK293 and A549 cell lines expressing biotin-tagged human PSMB2, PSMB5, and PSMD14 as previously described [29]. The cells were harvested with buffer A (proteasome lysis buffer; 25 mM Tris-HCl pH 7.5, 10% glycerol, 5 μM MgCl_2_, 1 mM ATP, 1 mM DTT, and protease inhibitor cocktail) and homogenized with Dounce homogenizer. After that, the lysates were centrifuged at 16,000 × g for 15 min at 4 °C. Nest, the supernatants were incubated with streptavidin magnetic beads for 16 h at 4 °C. The beads were washed twice with buffer A and once with buffer B (TEV cleavage buffer; 50 mM Tris-HCl pH 7.5 containing 1 mM ATP and 10% glycerol). Next, the proteasomes were eluted from the magnetic resin by incubating with TEV protease (Invitrogen) in buffer B containing 1 mM ATP for 1 h 30 min at 30 °C. Then, the eluted human proteasomes were concentrated with Amicon Ultra-0.5 centrifugal filter units (Millipore).

### Yeast strains, growth conditions, and yeast proteasome analysis

SUB62 WT (ura3-52, his3-200, leu2-3,112, lys2-801, trp1-1, gal), SUB62 Δecm29::TRP1. Cultures were grown in Standard YEP medium (1% yeast extract, 2% Bacto Peptone) supplemented with 2% dextrose (YPD). Yeast cells were grown exponentially in 5 mL YPD medium for 24h (the logarithmic phase) or for 9 days (the stationary phase and starvation period). Yeast cells were centrifuged at 3000 rpm for 2 min, and the pellet was resuspended in 2-fold volume of ATP-depleted buffer A (1 M Tris, pH 7.4, 10% glycerol, 1 M MgCl_2_). Cell were lysed using buffer A (0.5 M ATP and 1 M DTT added), 2-fold volume of glass beads was added to the tubes and the lysates underwent 5-6 intervals of 1 min vortex followed by 1 min in ice. The resultant supernatant was clarified by further centrifugation at 13,000 rpm for 30 min at 4 °C. The final supernatant was used immediately as the crude native extract. The protein concentration was determined by the method of Bradford and all gels were normalized to protein content. For nondenaturing PAGE resolving of 30S\26S\20S proteasomes, protein crude extract samples were resolved by 4% nondenaturing Tris-borate (pH 8.0) PAGE. The gels were then incubated in 10 ml of 0.1 mM suc-LLVY-AMC in buffer A for 10 min at 30 °C. Proteasome bands were visualized upon exposure to UV light (360 nm). For immunochemical identification of the 19S, protein samples were resolved on 10% SDS-PAGE followed by immunoblotting analysis using antibodies against Rpn1 (1:5,000) and Rpn5 (1:5,000).

### SDS-PAGE and gel staining

Cultured cells were washed twice with PBS, and whole-cell lysates were prepared in RIPA buffer supplemented with a protease inhibitor cocktail. For SDS-PAGE, lysates mixed with sample buffer (final 6.3% glycerol, 80 mM Tris-HCl pH 6.8, 62.5 μg/mL bromophenol blue, 10 mg/mL SDS, and 5% 2-mercaptoethanol) and denatured at 85°C for 10 min. After the running step, relatively small amounts of proteins were visualized by silver staining (K14040D, Koma Biotech) as the manufacturer’s guideline. Briefly, gels were fixed for 20 min or overnight in a fixing solution (40% (v/v) ethanol, 10% acetic acid, and 40% deionized water). They were then rinsed with a second fixing solution (50% ethanol in deionized water) and incubated with a sensitizing solution. After several washing steps with ultrapure water, gels were incubated in staining solution for 20 min and then in developing solution until the desired intensity was obtained. Image development was terminated by adding acetic acid (final 2%) and gentle agitation for 10 min.

### Immunoblotting

For immunoblotting, separated proteins by SDS-PAGE were transferred to polyvinylidene difluoride (PVDF) membrane. The membranes were blocked with 5% nonfat milk in TBS-T (20 mM Tris-HCl pH 7.5, 150 mM NaCl, and 0.1% (w/v) Tween 20) solution and incubated with primary antibody. Antibody sources and dilution factors were as follows: anti-β-actin/ACTB (A1978, Sigma, 1/10,000), anti-PSMA4 (PW8115, Enzo Life Science, 1/5,000), anti-PSMA5 (PA5-17295, Invitrogen, 1/1,000), anti-PSMB5 (PA1-977, Invitrogen, 1/1,000), anti-PSMB6 (PA1-978, Invitrogen, 1/1,000), anti-PSMC2 (sc-166972, Santa Cruz Biotechnology, 1/1,000), anti-PSMC4 (AT3466a, Abgent, 1/1,000), anti-PSMD1 (sc-514809, Santa Cruz Biotechnology, 1/1,000), anti-PSMD2 (PA527663, Invitrogen, 1/1,000), anti-PSMD11 (A15306, Abclonal, 1/1,000), anti-PSMD14 (ab109123, Abcam, 1/1,000), anti-LC3 (L7543, Sigma, 1/2,000), anti-USP14 (PA5-30300, Invitrogen, 1/1,000), anti-RAD23A (A19884, Abclonal, 1/1,000), anti-RAD23B (A20000, Abclonal, 1/1,000), anti-VCP (MA3-004, Invitrogen, 1/250), anti-TXNL1 (A6322, Abclonal, 1/1,000), anti-ECPAS (PA3-035, Invitrogen, 1/1,000), anti-RPL23 (A4292, Abclonal, 1/1,000), anti-EEF2 (A2068, Abclonal, 1/1,000), anti-eEF1A1 (05-235, Millipore, 1/1,000), anti-RPL11 (ab79352, Abcam, 1/1,000), anti-RPS6 (2217S, Cell Signaling, 1/1,000), anti-mTOR (2983S, Cell Signaling, 1/1,000), anti-p-mTOR (5536S, Cell Signaling, 1/1,000), anti-p-AKT (4060S, Cell Signaling, 1/1,000), anti-IRE1/ERN1 (3294s, Cell Signaling, 1/1,000), anti-p-IRE1/p-ERN1 (PA1-16927, Invitrogen, 1/1,000), anti-T7 (A190-117A, Bethyl Laboratories, 1/1,000), anti-HSC70 (ADI-SPA-815-D, Enzo Life Science, 1/1,000), anti-HSP70 (ADI-SPA-810-D, Enzo Life Science, 1/1,000), anti-RPS14 (A6727, Abclonal, 1/1,000), anti-RPL8 (A10042, Abclonal, 1/1,000), anti-eIF3e (ab36766, Abcam, 1/1,000), anti-puromycin (MABE343, Millipore, 1/3,000), anti-HA (901501, Biolegend, 1/1,000), and anti-flag (PA1-984B, Invitrogen, 1/1,000).. The membranes were then washed with TBS-T solution several times and followed by a horseradish peroxidase-conjugated anti-rabbit IgG or anti-mouse IgG antibody. After antibody incubation, bands were visualized via chemiluminescence, imaged, and quantified. Image J software was used to quantify bands intensity from independently replicated experiments more than three times, and statistical analysis was performed using GraphPad Prism V5 software (GraphPad Software). Results were evaluated to be significant with 95% confidence (*p*-value less than 0.05).

### Native gel electrophoresis and in-gel activity assay

Native gel analysis was performed as previously described [18]. WCLs and chromatography fractions were prepared with buffer A and buffer C (proteasome SEC buffer; 50 mM NaH_2_PO_4_ pH 7.5, 10 mM NaCl, 5 mM MgCl_2_, 5 mM ATP, 1 mM DTT, and protease inhibitor cocktails), respectively. Samples were resolved by NuPAGE™ 3–8% Tris-Acetate Protein Gels (Thermo-Fisher) at 150 V for 3–4 h. Gels were incubated in buffer D (in-gel activity assay buffer; 20 mM Tris, 1 mM ATP, 5 mM MgCl_2_) with proteasome fluorogenic substrate (100 μM suc-LLVY-AMC) for visualizing proteasome complexes. The addition of 0.02% SDS during in-gel hydrolysis assay activated and visualized the 20S proteasome. After the in-gel hydrolysis assay, separated protein complexes were transferred to PVDF membranes for subsequent immunoblotting analysis.

### Monitoring proteasomal activity via fluorogenic peptide substrates

Hydrolysis of the fluorogenic-peptide substrates suc-LLVY-AMC (7-amino-4-methylcoumarin; Bachem) was quantified to determine the proteolytic activity of proteasomes, as previously described [29]. Briefly, the fluorogenic-peptide substrate hydrolysis assay was carried out with 0.5 nM purified proteasome and 12.5 μM of suc-LLVY-AMC in buffer E (proteasome activity assay buffer; 50 mM Tris-HCl pH 7.5, 1 mg/mL BSA, 1 mM EDTA, 1 mM ATP, and 1 mM DTT). Proteasome activity was monitored by measuring free AMC fluorescence in a black 96-well plate on a TECAN infinite m200 fluorometer.

### *In vitro* ubiquitination of Sic1^PY^ and its degradation assay with purified proteasomes

Polyubiquitinated Sic1 with PY motif (Ub-Sic1^PY^) was prepared as previously described [29]. Briefly, the Ub conjugation reaction was conducted with 10 pmol Sic1^PY^, 2 pmol Uba1, 5 pmol Ubc4, 5 pmol Rps5, and 1.2 nmol ubiquitin in buffer F (ubiquitination reconstitution buffer; 50 mM Tris-HCl pH 7.4, 100 mM NaCl, 1 mM DTT, 2 mM ATP, and 10 mM MgCl2) for 4 h and 25 °C. For Ub-Sic1 degradation assay, purified human proteasomes (5 nM) under normal or glucose-depleted conditions were incubated with 20 nM of Ub-Sic1^PY^ in buffer E. Ub-Sic1^PY^ degradation was monitored by SDS-PAGE/immunoblotting using anti-T7 antibodies (Millipore).

### Sucrose density gradient fractionation and size exclusion chromatography (SEC)

For sucrose-gradient ultracentrifugation, +/+ and *ECPAS*^-/-^ MEFs extracts were homogenized with 1 mL of buffer C and centrifuged. After harvesting, the soluble fractions were loaded on top of a pre-established sucrose gradient (12 mL, 10–30%) and centrifuged at 312,000 × g in a Beckman SW-41 Ti rotor for 16 h at 4 °C. After that, the gradients were manually fractionated into 300 μL fractions from top to bottom. For SEC, HeLa WCLs were centrifuged twice at 18,000 × g for 30 min at 4 °C, loaded (11 mg by protein) onto a column, and eluted with buffer C. SEC was carried out on a Superose 6 Increase 10/300 GL column by fast protein liquid chromatography (ÄKTA; GE Healthcare); 0.25 mM fractions were collected, and 10% glycerol was added to each fraction.

### Assessment of cell viability and intracellular ATP levels

Cell viability assays were performed using a modified 3-(4, 5-dimethylthiazol-2-yl)-2, 5-diphenyltetrazolium bromide (MTT) assay. Briefly, MTT solution was added to the cell culture media and incubated for 3 h at 37 °C to form blue MTT-formazan products, which were assessed by measuring absorbance measurement (570 nm for test and 630 nm for reference wavelength). Cellular ATP concentration was measured using a coupled luciferin-luciferase reaction with the ATP determination kit (A22066, Invitrogen). Briefly, cells were plated in 6-well plates in equal densities. After treatment, cells were washed twice with PBS and then scrapped with buffer G (ATP assessment buffer; 25 mM Tris-HCl pH 7.5, 0.1% Triton X-100). Cells were homogenized with a 26G 1/2 needle attached to a 1 mL syringe and incubated for 15 min on ice. The samples were centrifuged at 13,000 rpm for 10 min at 4 °C in a tabletop centrifuge. Samples (10 μL) were mixed with 90 μL of ATP reaction solution and rapidly assessed using a luminometer (TECAN Infinite 200 PRO)

### RNA isolation and RT-PCR

Total RNA was isolated from either A549 or MEF cells using TRIzol-reagents and additional purification using RNeasy mini-columns (Qiagen) with on-column DNase I treatment. cDNA samples were prepared using two micrograms of total RNA by reverse transcription (RT)-PCR (Accupower RT-pre-mix, Bioneer). Quantitative RT-PCR reactions were conducted with 1/20 diluted cDNA, SYBR qPCR master mixture (Kapa Biosystems, USA) as the reporter dye, and 10 pmol of primers to detect mRNA expression of specific genes. Primer sequences for human gene were as follows: for *PSMB5*, forward 5′-ACTACGTGGACAGTGAAGGG-3′ and reverse 5′-GGCATCTCTGTAGGTGGCTT-3′; for *PSMD4*, forward 5′- CTTCGTGTATCTATGGAA-3′ and reverse 5′-TCATACTGCTTAGGTCAGG-3′; for *PSMA7* forward 5′-AAGCAGCGTTATACGCAGAG-3′ and reverse 5′- CATCAAAGTCGAAACCCACGAT-3′; for *PSMA4*, forward 5′- AGAAGTGGAGCAGTTGATCA-3′ and reverse 5′-TCTCTGATTCTATTTATCCTTTTCT-3′; for *PSMC6*, forward 5′- GGAGGGCTATCAGAACAGATCC-3′ and reverse 5′- GGCTCGTGCAAGAGTGTTT-3′; for *NFE2L1*, forward 5′- CATTCTGCTGAGTTTGATTGGGG-3′ and reverse 5′-TTGTGGAACTGGGTCTGAGTAT-3′; for *LC3*, forward 5′-AACATGAGCGAGTTGGTCAAG-3′ and reverse 5′- GCTCGTAGATGTCCGCGAT-3′; for SQSTM1, 5′-AAGCCGGGTGGGAATGTTG -3′ and reverse 5′-CCTGAACAGTTATCCGACTCCAT-3′; and for *GAPDH*, forward 5′- GAGTCAACGGATTTGGTCGT-3′ and reverse 5′-GACAAGCTTCCCGTTCTCAG-3′. And primer sequences for mouse gene were as follows: for *mXBP1* (spliced), forward 5′- AGCACCTCTAAGCTCTTCAG-3′ and reverse 5′-GAGCCCTCATATCCACAGTCAC-3′; for *mATF4*, forward 5′-ATGGCCGGCTATGGATGAT-3′ and reverse 5′- CGAAGTCAAACTCTTTCAGATCCA-3′; for *mCHOP*, forward 5′-AAGCCTGGTATGAGGATCTGC-3′ and reverse 5′-TTCCTGGGGATGAGATATAGGTG-3′; for *mBAX*, forward 5′-GCAGAGGATGATTGCTGACG-3′ and reverse 5′- GCAAAGTAGAAGAGGGCAACC-3′; for *mBIM*, forward 5′- GGAGATACGGATTGCACAGGAG-3′ and reverse 5′- CTCCATACCAGACGGAAGATAAAG-3′; for *mDR5*, forward 5′- TGTGTCGATGCAAACCAGGCAC-3′ and reverse 5′-GCCGTTTTGGAGACACACTTCC-3′; for *mGADD34*, forward 5′-ACACAGAAGAGGAAGAGGACAG-3′ and reverse 5′- TCCCTTGGTCTTCTCTCCTG-3′ and reverse 5′-TCCCTTGGTCTTCTCTCCTG-3′; for *mTRB3*, forward 5′-CAGGAAGAAACCGTTGGAGTT-3′ and reverse 5′- CCAAAAGGATATAAGGCCCCAGT-3′; and for *mGAPDH*, forward 5′- AGGTCGGTGTGAACGGATTTG-3′ and reverse 5′-GGGGTCGTTGATGGCAACA-3′. Each mRNA level was normalized to that of *GAPDH*. Unpaired Student’s t-tests were performed to evaluate the results, where *p*-values of less than 0.05 were considered statistically significant.

### RNA interference

The siRNAs were synthesized by Genolution. The sequences of siRNA are as follows: control siRNA sense 5′- CCUCGUGCCGUUCCAUCAGGUAGUU-3′; control siRNA antisense 5′- CUACCUGAUGGAACGGCACGAGGUU-3′; *ECPAS* siRNA-B sense 5′- CUAGGAGUCCUGCAAUCAAUU-3′; *ECPAS* siRNA-B antisense 5′- UUGAUUGCAGGACUCCUAGUU-3′; *ECPAS* siRNA-C sense 5′- GCUAAGUACCCACAAAGAAUU-3′; *ECPAS* siRNA-C antisense 5′-UUCUUUGUGGGUACUUAGCUU-3′. Both siRNA-B and siRNA-C target the sequences within the 3′ untranslated regions (3′-UTR) of human *ECPAS* mRNA. siRNA duplexes (siRNA B and C) were transfected into HEK293-PSMB2-HB cells using RNAiMAX (Invitrogen) dissolved in Opti-MEM (Invitrogen), following the manufacturer’s protocols. The final concentration of siRNA for transfection was 20 nM. After 24 h, cells were trypsinized, suspended in the transfection mixture, and re-plated. On the following day, cells were transfected again with the siRNA duplexes and grown for an additional 48 h for an effective knockdown.

### LC-MS/MS analysis with the iBAQ algorithm

The label-free, quantitative LC-MS/MS analysis with the iBAQ algorithm was performed, mainly as previously described [18], with modifications in data analysis. Briefly, the iBAQ values obtained from biological triplicates in each condition were normalized to the average values of PSMB2 subunits from biological replicates. To identify proteasome-associating proteins whose interactions were significantly altered upon glucose starvation, we first selected preliminary differentially associated proteins (DAPs; >1.5-fold changes and <0.05 absolute p-values) between glucose-deprived and control conditions. Next, using the log2-iBAQ values, we estimated the empirical null distribution (i.e., a protein is not differentially expressed between two conditions) of t-statistic values between the two conditions by performing random permutations of all six samples (three normal and three glucose-depleted conditions) 1,000 times. We then computed adjusted p-values using the estimated empirical distribution. We finally selected the DAPs as the proteins with adjusted *p*-values < 0.05 and absolute log2-fold-changes > 1 (2-fold). In addition, we included the proteins detected more than once in one sample group alone as the DAPs.

### Computational modeling construction

*In silico* structure modeling of the ECPAS-26S proteasome complex was performed using AlphaFold [55] and GalaxyTongDock [56], based on the available structure of the 26S proteasome in the substrate-accepting state (PDB: 6MSB [57]) and the predicted structure of ECPAS (https://alphafold.ebi.ac.uk/entry/Q5VYK3). The AlphaFold model has a high confidence level in terms of the overall structure, with a median pLDDT of 88.41. For the initial-stage construction of ECPAS-bound 19S structures, the interface option of GalaxyTongDock was used to favor previously proposed interaction partners of ECPAS, including subunits PSMD2/Rpn1, PSMD4/Rpn10, PSMC2/Rpt1, and PSMC3/Rpt5 [25, 35]. Two different modeling with PSMD and PSMC subunits resulted in two different conformational states, termed “assembled” and “disassembled” states, respectively. The undefined loop structure in the middle of PSMD4 (amino acids 192-323) was filled with a predicted structure generated by AlphaFold-Multimer and GalaxyLoop [58]. Energy minimization, segment connection, side-chain repacking, and additional linker modeling were performed by in-house GALAXY tools for the predicted complexes to retain no disaccorded structure, such as steric clashes at the interface and mismatches from the experimentally identified proteasome structures. For further refinement, the models were equilibrated by the GalaxyRefineComplex program which uses a linear combination of energy functions including physics-based, knowledge-based, and restraint energy terms [59].

### Tumor xenograft models and assessment of intracellular glucose levels

Mice were housed in a facility fully accredited by the Institutional Animal Care and Use Committee (IACUC) of Seoul National University. For the assessment of 26S/20S proteasome ratios in a tumor, A549 cells (5 × 10^6^ cells) were suspended in 1 mL of 1:1 mixture of PBS and matrigel, and 100 μL aliquot was subcutaneously injected into the right flank region of 7-week-old NOD/SCID mice (The Jackson Laboratory) with a 26-gauge syringe. The mice were sacrificed after 2 or 6 weeks post-implantation, and the tumor tissues, lungs, and other organs were removed for subsequent biochemical analyses.

Cellular glucose concentration was measured using the Picoprobe Glucose Assay Kit (ab169559, Abcam) according to the manufacturer’s protocol. Briefly, xenograft mouse tumor tissues were homogenized with 100 μL of Glucose Assay Buffer and centrifuged at 13,000 rpm for 5 min in a tabletop centrifuge. The supernatant of lysates was assessed for protein content using the Coomassie Protein Assay Reagent (23200, Thermo Fisher Scientific) and spin filtered through Amicon Ultra-0.5 centrifugal filter units (10 kDa MWCO, Millipore) to remove proteins and various enzymes. The filtered lysates were loaded at 2 μg protein per well in a 96 black-well plate, and a reaction mixture was added. After incubation for 30 min at 37 °C, the fluorescence signal was measured at Ex/Em 535/587 nm.

### Statistical analysis

Statistical significance of differences between various groups was determined by two-tailed Student’s *t*-test or one-way ANOVA followed by the Bonferroni *post hoc* test for most data (GraphPad Prism, ver. 9.3.1). Differences were considered significant at *p* < 0.05.

## Acknowledgment

This work was supported by grants from the National Research Foundation (NRF; 2021R1A2C2008023 to M.J.L., 2020R1A5A1019023 to D.H., M.J.L., and Y.T.K., and 2020M3A9G7103933 to C.S.), the Korea Health Industry Development Institute and Korea Dementia Research Center (HU21C0071 to M.J.L.), the Creative-Pioneering Researchers Program through Seoul National University (to M.J.L.), the Gyeonggi-do Regional Research Center program (GRRC-Kyung Hee 2018[B03] to K.P.K.), the BK21 FOUR program (to W.H.C), and Israel Science Foundation grants (755/19 and 2512/18 to M.H.G.). We gratefully acknowledge H. K. Song (Korea University) and K. Nam (UT Arlington) for technical assistance on structural analysis.

## Author Contributions

Most of the biochemical and cell-based experiments were performed by W.H.C. and Y.Y; CRISPR/Cas9-mediated ECPAS knockout was conducted by I.B. and S.H.P; gel filtration chromatography assays were conducted by S.K.; structural modeling was conducted by S.L., J.S., I.B., J.J., and C.S.; quantitative LC-MS/MS was performed by K.P.K. and D.H.; *ECPAS*^-/-^ MEFs were generated by T.C.; xenograft tumor tissues were analyzed by Y.T.K.; yeast data and CycB degradation data were produced by S.L. and M.H.G.; the paper was written by M.J.L., W.H.C. Y.Y., and M.H.G.; and was critically reviewed by all authors.

## Competing financial interests

The authors declare no competing financial interests.

**Note**: All supplementary information and source data files are available in the online version of the paper.

