## Supplementary Information for "ECPAS/Ecm29-Mediated 26S Proteasome Disassembly is an Adaptive Response to Glucose Starvation"

This PDF file includes:

Supplementary Figures 1 to 4

Supplementary Table 1

### SUPPLEMENTARY FIGURES

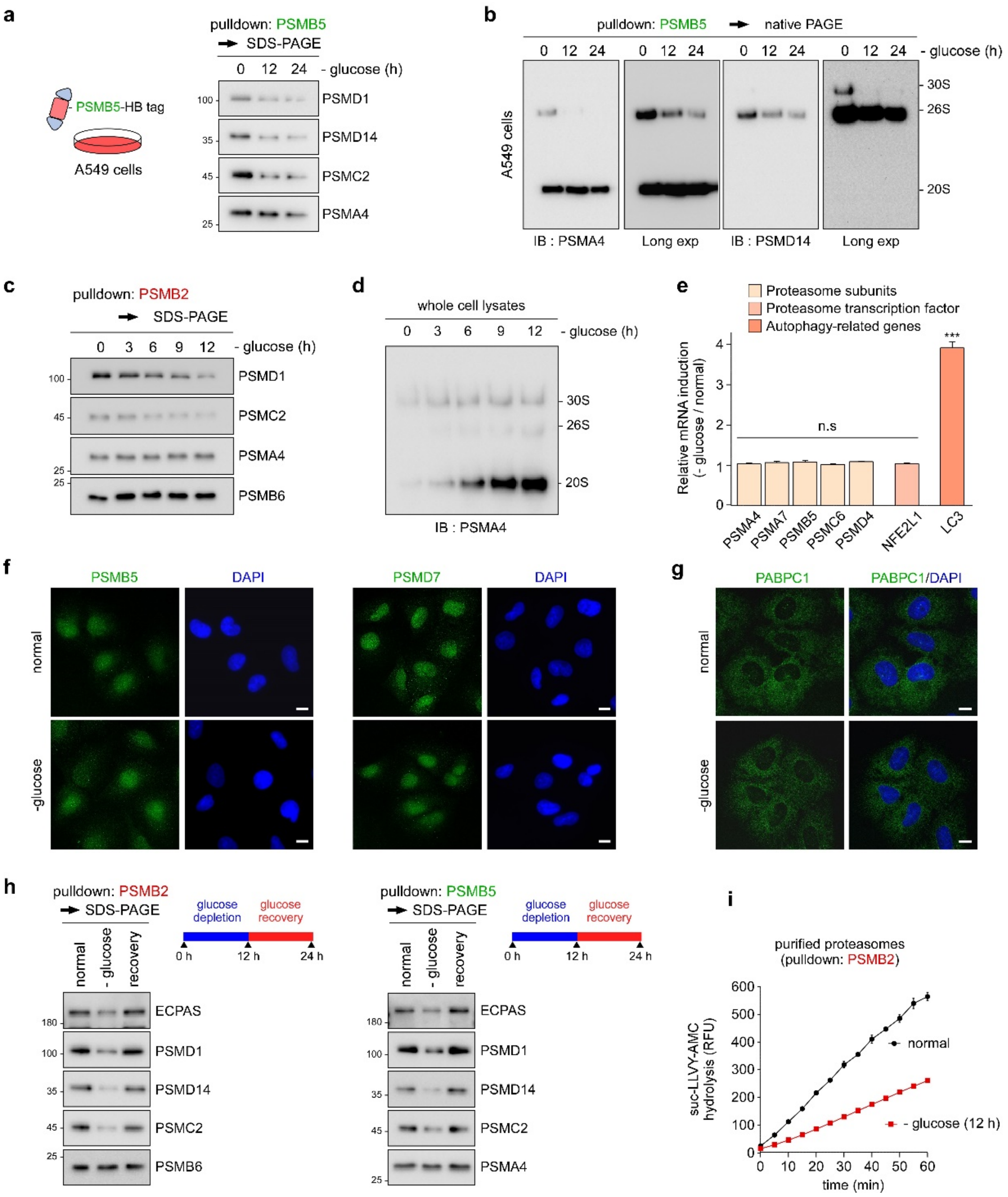

**Supplementary figure 1. Glucose depletion leads to the dissociation of intracellular 26S/30S proteasomes into 20S and 19S subcomplexes**

(**a**) PSMB5-HB proteasomes were affinity-purified from A549-PSMB5-HB stable cells, which were glucose-deprived for the indicated periods (0, 12, and 24 h), and were subjected to SDS-PAGE and immunoblotting (IB) using antibodies against 20S and 19S subunits. (**b**) As in (a), except that the purified proteasomes were resolved using non-denaturing (native) PAGE with subsequent IB; blots were probed for PSMA4 or PSMD14. (**c**) HEK293-PSMB2-HB cells were cultured in glucose-depleted media for indicated time periods (0 to 12 h), and the proteasome subunit levels in purified proteasomes were analyzed by SDS-PAGE/IB. (**d**) As in (c), except that the whole-cell lysates (WCLs) were analyzed using native PAGE/IB. (**e**) A549 cells were cultured in glucose-depleted media for 12 h, total RNA was isolated, and proteasome subunit genes (*PSMA4, PSMA7, PSMB5, PSMC6,* and *PSMD4*), a transcription factor of proteasome biogenesis (*NFE2L1*), and an autophagy-related gene (*LC3*) were analyzed by quantitative reverse transcription-polymerase chain reaction (qRT-PCR). *GAPDH* mRNA levels were used to normalize the results. Values are represented as the mean ± s.d. of three independent experiments (N = 3); $^{***}p < 0.001$ (one-way analysis of variance (ANOVA) with Bonferroni's multiple comparison test). n.s., not significant. (**f**) Representative images of A549 cells treated with glucose-depleted media for 12 h. Immunofluorescence staining with anti-PSMB5 (*left*) or anti-PSMD7 (*right*) antibodies (green). Nuclei were counter-stained with DAPI (blue). (**g**) As in (f), except that poly(A) binding protein cytoplasmic 1 (PABPC1/PABP1) was immunostained as a marker of stress granules (**h**) (*left*) HEK293-PSMB2-HB cells were cultured in glucose-depleted media for 12 h and re-incubated with normal (glucose-containing) media for an additional 12 h. Purified PSMB2-HB proteasomes were subjected to native PAGE/IB. (*right*) The same experiments were performed with A549-PSMB5-HB stable cells. Considering that there was no induction of proteasome subunits and NFE2L1/Nrf1 genes during glucose starvation, as shown in panel (e), these results strongly suggest that the reconstitution of 26S proteasomes might be the reassembly of pre-existing 20S and 19S subcomplexes, instead of *de novo* synthesized 26S proteasomes. (**i**) Proteasome activity was measured using suc-LLVY-AMC and purified PSMB2-HB proteasomes under normal and glucose-depleted (for 12 h) conditions. These data supplement Figure 1.

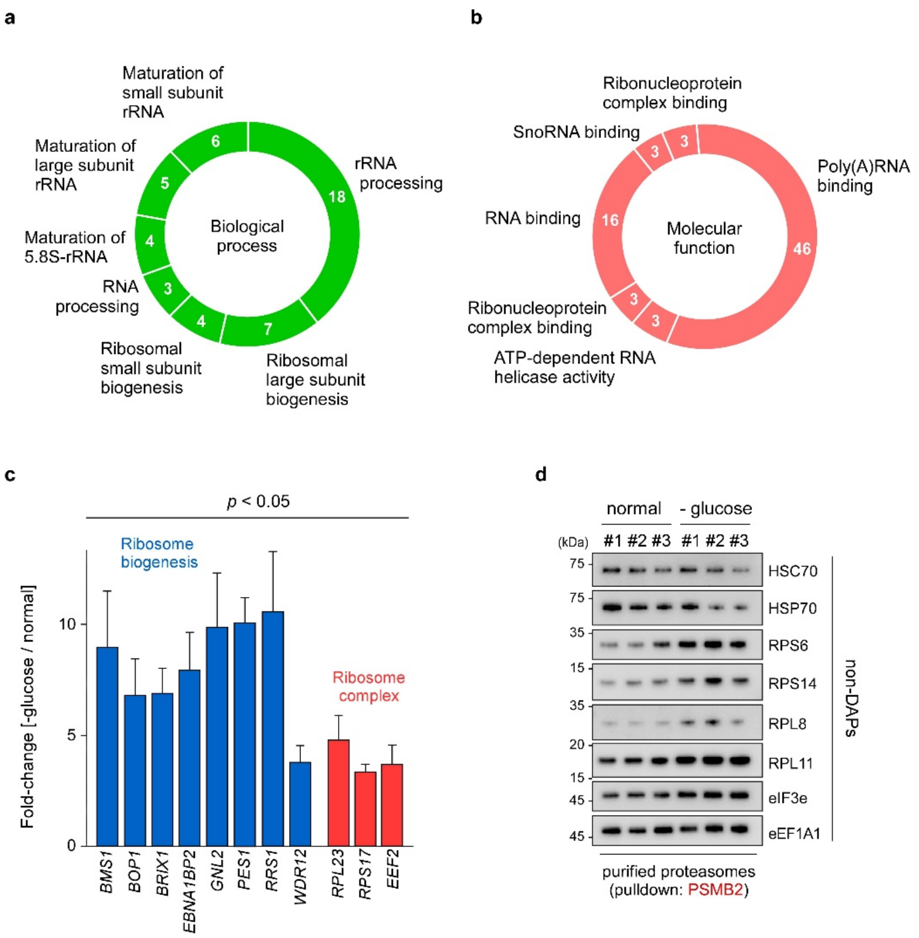

**Supplementary figure 2. Gene ontology (GO) enrichment of the differentially associated proteins (DAPs), which were more strongly bound to the purified proteasome during glucose starvation**

(**a** and **b**) Classification of increased DAPs by GO analysis by DAVID functional annotation tool. The important biological processes (**a**) and molecular functions (**b**) were enriched, and the count numbers are displayed. Note that the majority of increased DAPs are involved in ribosome biogenesis and rRNA processing. (**c**) Quantification of fold-changes in the increased DAPs, which are ribosome subunits or involved in ribosome biogenesis. Data represent mean ± s.d (N = 3). $p < 0.05$ (one-way ANOVA with Bonferroni's multiple comparison test). (**d**) Examination of non-DAPs, whose levels were changed less than 2-fold from the iBAQ-MS analysis using IB analysis. These data supplement Figure 2.

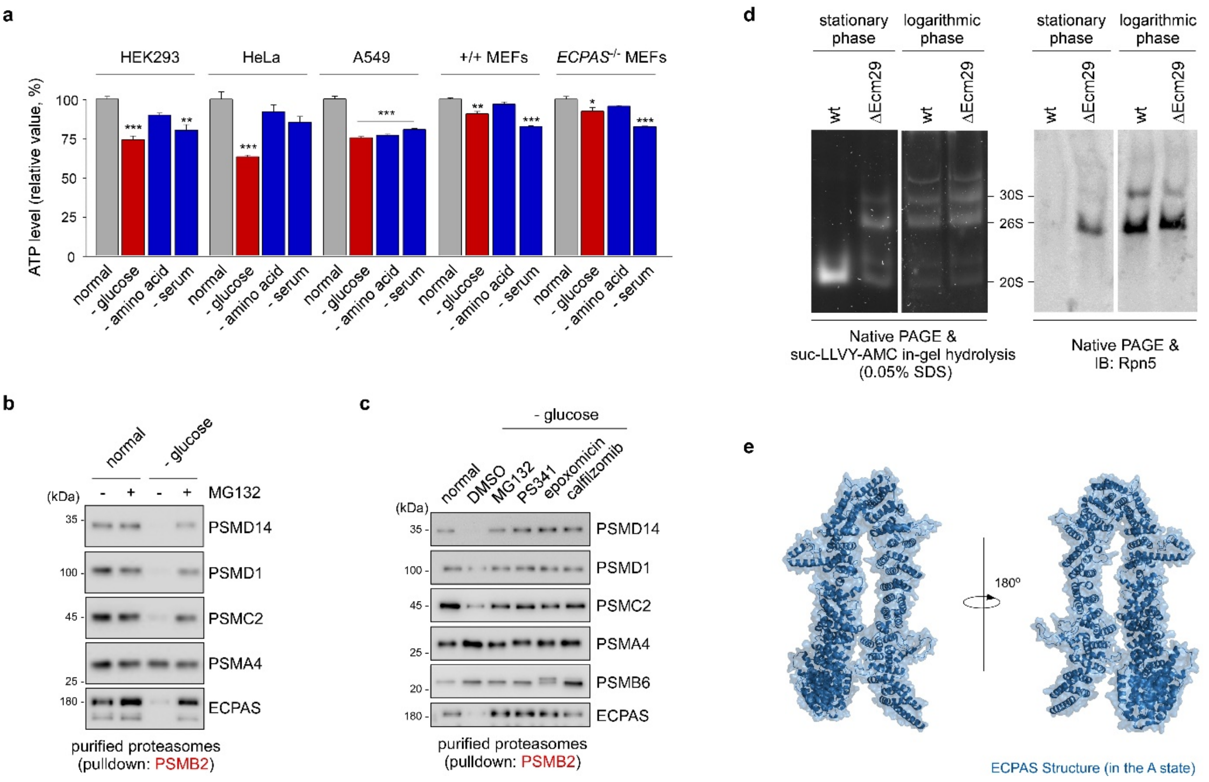

**Supplementary figure 3. Changes in cellular ATP levels and yeast 26S proteasomes during nutrient deprivation**

(**a**) Nutrient deprivation had modest effects on intracellular ATP levels. HEK293, HeLa, A549, wild-type (+/+), and *ECPAS*$^{-/-}$ mouse embryonic fibroblasts (MEFs) were cultured in nutrient-restricted (glucose-free, amino acid-free, and serum-free) media for 12 h, and cellular ATP concentrations were measured using the bioluminescent assay (based on ATP-dependent luciferin-luciferase reactions). Data are presented as the mean ± s.d. (N = 3); $*p < 0.05$, $**p < 0.01$, $***p < 0.001$ (one-way ANOVA with Bonferroni's multiple comparison test). (**b**) Inhibition of 20S-19S disassembly by MG132 (5 μM) in HEK293-PSMB2-HB stable cells. Affinity purified PSMB2-HB proteasomes were analyzed by SDS-PAGE/IB. (**c**) Cells treated with both reversible (MG132 and PS-341) and irreversible (epoxomicin and carfilzomib) proteasome inhibitors were analyzed as described in (b). (**d**) Yeast proteasomes were purified in stationary or logarithmic phase cultures, separated by native PAGE, and visualized by suc-

LLVY-AMC in-gel hydrolysis in the presence of 0.05% SDS (*left*). After the in-gel activity assay, the gel was transferred to a PVDF membrane and subjected to IB against yeast Rpn5 (*right*). (**e**) A predicted structure (ribbon diagram) of ECPAS by AlphaFold (UniProt ID: Q5VYK3). These data supplement Figure 3.

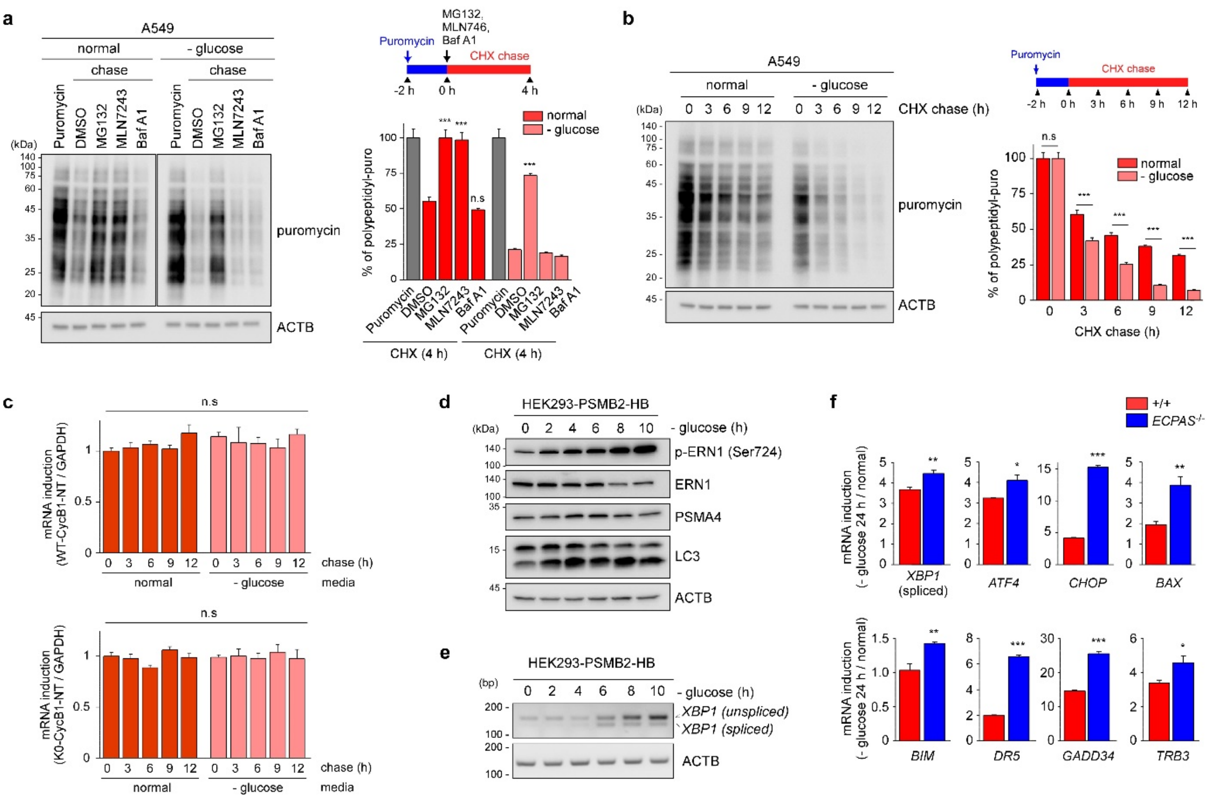

**Supplementary figure 4. Misfolded protein degradation and ER stress in glucose-depleted cells**

(**a** and **b**) As in Figures 4a and 4b, except that the puromycin-labeling and monitoring of polypeptidyl-puro species under various conditions were conducted with A549 cells. (*right panels*) Quantification of polypeptidyl-puro levels, which were normalized to endogenous ACTB/β-actin levels. Data represent the average percentage of remaining polypeptidyl-puro (mean ± s.d) of three independent experiments (N = 3), *** $p < 0.001$ (one-way ANOVA followed by the Bonferroni *post hoc* test). (**c**) Total RNA was isolated from the WT- and K0-CycB1-NT-transfected HEK293 cells after glucose starvation for 12 h. Quantitative RT-PCR was conducted using primers specific for either WT or K0-CycB-NT. mRNA levels were normalized to levels of the *GAPDH* gene. The plotted values are means ± s.d. of three independent experiments. n.s., not significant. (**d**) HEK293-PSMB2-HB cells were cultured in glucose-depleted media for indicated time periods (0 to 10 h). WCLs were separated using

SDS-PAGE with subsequent IB; blots were probed for ERN1/IRE1, p-ERN1/p-IRE1 (Ser 724), α3, LC3, and ACTB/β-actin. (**e**) As in (a), except that mRNA levels of endogenous unspliced *XBP1*, spliced *XBP1,* and *ACTB* from HEK-2930PSMB-HB cells were measured by RT-PCR analysis. (**f**) Total RNA was isolated from +/+ and $ECPAS^{-/-}$ MEFs after 24 h of glucose starvation and subjected to quantitative RT-PCR for various unfolded protein response (UPR)-target genes. Data are presented as the means ± s.d. of three independent experiments (N = 3). *, $p < 0.01$ (one-way ANOVA with Bonferroni's multiple comparison test). These data supplement Figure 4.

**Supplementary Table S1**. (**a**) All the proteins identified by iBAQ-MS. (**b**) Normalized quantities of the identified proteins based on the levels of the PSMB2 subunit. (**c**) Decreased DAPs after glucose depletion. (**d**) Increased DAPs after glucose depletion.

**Table S1. Total identified proteins_raw value**

| Protein IDs | Protein names | Gene names | Normal #1 | Normal #2 | Normal #3 | glucose-free #1 | glucose-free #2 | glucose-free #3 |
|---|---|---|---|---|---|---|---|---|
| Protein IDs | Protein names | Gene names | iBAQ normal_1 | iBAQ normal_2 | iBAQ normal_3 | iBAQ glucose_1 | iBAQ glucose_2 | iBAQ glucose_3 |
| A00006;P49721 | Proteasome subunit beta type-2 | PSMB2 | 1786200000 | 955300000 | 1094900000 | 930150000 | 746610000 | 513550000 |
| A00007;O00487 | 26S proteasome non-ATPase regulatory subunit 14 | PSMD14 | 167660000 | 109360000 | 157580000 | 12167000 | 15719000 | 11199000 |
| A0A024R7W5;A0A087X0Q1;A0A087WY31;Q7Z739;Q9BYJ9 | YTH domain-containing family protein 3;YTH domain-containing family protein 1 | YTHDF3;YTHDF1 | 0 | 97333 | 729690 | 638420 | 1127300 | 130560 |
| A0A075B6E5;Q8N8S7-3;Q8N8S7-2;Q8N8S7;Q9UI08 | Protein enabled homolog;Ena/VASP-like protein | ENAH;EVL | 837600 | 419790 | 915900 | 0 | 0 | 0 |
| H0YN88;A0A075B716;P0CW22;P08708;H0YN73;H3BNC9;REV__F8WFC6;REV__Q9H089 | 40S ribosomal protein S17-like;40S ribosomal protein S17 | RPS17;RPS17L | 143830000 | 41132000 | 92972000 | 178520000 | 203500000 | 122940000 |
| A0A087WSW7;F6XU50;Q9NRZ9-6;Q9NRZ9-5;Q9NRZ9-4;Q9NRZ9-3;Q9NRZ9-2;Q9NRZ9;Q9NW36 | Lymphoid-specific helicase | HELLS | 281370 | 200590 | 133060 | 47713 | 85911 | 112330 |
| E2QRB9;E9PIR7;F8W809;A0A087WSW9;E9PNQ6;A0A087WSY9;Q16881-7;Q16881-5;Q16881-2;Q16881-4;Q16881-3;Q16881-6;Q16881 | Thioredoxin reductase 1, cytoplasmic | TXNRD1 | 0 | 112980 | 213270 | 0 | 0 | 0 |
| A0A087WT58 | | MROH2A | 110190 | 61609 | 51313 | 0 | 0 | 0 |
| A0A087WTP3;Q92945;M0R0I5;M0QXW7;M0QYH3;M0QYG1;M0R0C6 | Far upstream element-binding protein 2 | KHSRP | 1590900 | 878140 | 1682500 | 814690 | 1273500 | 604500 |
| A0A087WTV6;A0A087WZR9;Q96C36 | Pyrroline-5-carboxylate reductase;Pyrroline-5-carboxylate reductase 2 | PYCR2 | 0 | 0 | 220640 | 0 | 167390 | 0 |
| A0A087WTZ5;E9PJ81;Q04323;Q04323-2;E9PS08;E9PRQ7 | UBX domain-containing protein 1 | UBXN1 | 296030 | 659830 | 907330 | 0 | 144550 | 0 |
| A0A087WUK2;O14979-3;O14979-2;O14979 | Heterogeneous nuclear ribonucleoprotein D-like | HNRNPDL | 1098700 | 760790 | 2053100 | 53310 | 877910 | 409220 |
| E7ETK0;A0A087WUS0;P62847-2;P62847-3;P62847;P62847-4 | 40S ribosomal protein S24 | RPS24 | 20117000 | 5333000 | 22247000 | 11886000 | 32216000 | 20565000 |
| A0A087WUT6;O60841 | Eukaryotic translation initiation factor 5B | EIF5B | 1717500 | 1920700 | 3384000 | 486840 | 1083500 | 2014400 |
| A0A087WUZ3;Q01082;Q01082-3;Q01082-2;F8W6C1;P11277-3;P11277;P11277-2;K7EIL4;K7ERK8;K7ELR7;K7EP41;K7ERC6;K7ENA0;K7EPY4 | Spectrin beta chain, non-erythrocytic 1 | SPTBN1 | 8972700 | 3438200 | 7767300 | 502290 | 817920 | 531950 |
| A0A087WV05;Q99417 | C-Myc-binding protein | MYCBP | 12604000 | 6083900 | 10000000 | 3867200 | 6548100 | 3519400 |
| A0A087WV66;P46013-2;P46013 | Antigen KI-67 | MKI67 | 226180 | 234700 | 292850 | 259950 | 106830 | 222640 |
| A0A087WW61;A0A087WX67;A0A087WXV3;A0A087WXQ5;K7EJM5;A0A087WV68;K7EN94;B1AKP7;G3V162;A0A087X260;A0A087WYY0;Q13148-4;Q13148;A0A087WZC9;A0A087WX29 | TAR DNA-binding protein 43 | TARDBP | 5778100 | 1998700 | 4904400 | 4322500 | 5299700 | 2344200 |
| A0A087WVC1;A0A087WVB0;Q9BQ39 | ATP-dependent RNA helicase DDX50 | DDX50 | 607210 | 620020 | 1746400 | 2204700 | 5032900 | 2550500 |
| A0A087WVQ6;Q00610-2;Q00610;J3KS13;A0A087WX41;P53675-2;P53675;K7EJJ5;J3KSQ2;J3QL20;J3KRF5;F5H5N6 | Clathrin heavy chain;Clathrin heavy chain 1 | CLTC | 57104000 | 16897000 | 29570000 | 801070 | 2472200 | 1322300 |
| A0A087WVV2;Q9P2E9-3;Q9P2E9-2;Q9P2E9;F8W7S5;A0A087WU26;V9GY78;H0YDJ4;H9KV85;E9PN67;C9J066;Q8N4C6-6;Q8N4C6-11;Q8N4C6-4;Q8N4C6-9;Q8N4C6-2;Q8N4C6;Q8N4C6-5;Q8N4C6-10;Q8N4C6-7 | Ribosome-binding protein 1 | RRBP1 | 186510 | 278450 | 910750 | 1210200 | 4178800 | 1658200 |
| A0A087WVZ9;P19388;A0A087WWX0;A0A087WVN5 | DNA-directed RNA polymerases I, II, and III subunit RPABC1 | POLR2E | 13245000 | 13362000 | 23084000 | 9113600 | 12553000 | 6639000 |
| A0A087WW40;Q9Y371;Q9Y371-2 | Endophilin-B1 | SH3GLB1 | 100400 | 0 | 0 | 0 | 0 | 0 |
| A0A087WW66;H7BZR6;H7C378 | | PSMD1 | 2754300 | 920120 | 827650 | 136900 | 66586 | 0 |
| A0A087WWP8;Q5VWQ0;H7C2D3;C9JM20;Q5VWQ0-4 | Round spermatid basic protein 1 | RSBN1;RSBN1L | 0 | 0 | 69593 | 58515 | 286540 | 0 |
| A0A087WX58;K7EPS6;Q7Z4V5-2;Q7Z4V5 | Hepatoma-derived growth factor-related protein 2 | HDGFRP2 | 4389600 | 2138900 | 1425400 | 1703400 | 1862000 | 527940 |
| A0A087WXM6;J3QQT2;J3KRX5;P18621;P18621-3;J3KRB3;A0A087WWH0;J3QS96;J3QLC8;P18621-2;A0A087WY81;J3KSJ0;K7EQ71;K7EKI8;O60437 | 60S ribosomal protein L17 | RPL17 | 198590000 | 125750000 | 247910000 | 195360000 | 359540000 | 204420000 |
| A0A087WXZ3;Q9Y5A9;Q9Y5A9-2;S4R3V3;S4R3J8 | YTH domain-containing family protein 2 | YTHDF2 | 7177500 | 3982000 | 12949000 | 8910600 | 14730000 | 7593000 |
| A0A087WY00;G3V394;F8WE88;F8W6H6;Q9Y4I1-2;Q9Y4I1;Q9Y4I1-3;Q9UES5;O95317;E7ERV5 | Unconventional myosin-Va | MYO5A | 45445 | 19209 | 67913 | 8636.2 | 62563 | 23246 |
| A0A087WY43;P06493;E5RIU6;A0A087WZZ9;P06493-2;K7ELV5;Q00526 | Cyclin-dependent kinase 1 | CDK1 | 8342400 | 4896600 | 8745800 | 1524500 | 3667500 | 1841300 |
| A0A087WY71;E9PFW3;Q96CW1-2;Q96CW1;C9JTK4;C9JGT8;C9JJ47;C9JJD3;C9JPV8;H7C4C3 | AP-2 complex subunit mu | AP2M1 | 4939000 | 1911300 | 4793200 | 2119000 | 1941000 | 1475700 |
| A0A087WYN9;Q7Z478 | ATP-dependent RNA helicase DHX29 | DHX29 | 556580 | 242200 | 1309700 | 591700 | 1833800 | 971680 |
| A0A087WYP2;Q8IZQ5;H0YE28 | Selenoprotein H | C11orf31;SELH | 1333100 | 1259900 | 1341700 | 1213600 | 2872200 | 2282000 |
| A0A087WYR0;P09132 | Signal recognition particle 19 kDa protein | SRP19 | 5485000 | 1763100 | 0 | 1121800 | 3236900 | 1289900 |
| B4DDC6;A0A087WYT3;Q15185-3;Q15185-4;Q15185;Q15185-2 | Prostaglandin E synthase 3 | PTGES3 | 6841500 | 9466500 | 3612200 | 3059200 | 4917300 | 0 |
| A0A087WYU1;Q9Y5X1 | Sorting nexin;Sorting nexin-9 | SNX9 | 835420 | 527670 | 661100 | 252270 | 300950 | 228760 |
| A0A087WZH7;P29966 | Myristoylated alanine-rich C-kinase substrate | MARCKS | 6384900 | 3413000 | 9614900 | 353490 | 836000 | 396330 |
| D6R9P3;D6RD18;D6RBZ0;A0A087WZV1;Q99729-3;Q99729-2;Q99729-4;Q99729 | Heterogeneous nuclear ribonucleoprotein A/B | HNRNPAB | 14099000 | 10600000 | 28489000 | 18566000 | 35176000 | 18354000 |
| H7C2F6;J3KPF9;A0A087X011;E9PES4;Q9Y496 | Kinesin-like protein;Kinesin-like protein KIF3A | KIF3A | 0 | 0 | 0 | 0 | 0 | 0 |
| A0A087X020;Q9Y3A5;F8WE72 | Ribosome maturation protein SBDS | SBDS | 16851000 | 2685900 | 12385000 | 700900 | 2874200 | 1514500 |
| C9J8Z2;H7C4F1;C9J975;A0A087X044;E7ESL9;A0A087X1U1;F5GXM9;E7EUN2;Q96P47-3;Q96P47-6;Q9UPQ3-3;Q9UPQ3-2;Q9UPQ3;Q96P47;Q96P47-2;Q96P47-4 | Arf-GAP with GTPase, ANK repeat and PH domain-containing protein 3;Arf-GAP with GTPase, ANK repeat and PH domain-containing protein 1 | AGAP1;AGAP3 | 188790 | 158980 | 1035600 | 0 | 0 | 0 |
| A0A087X054;Q9Y4L1;E9PJ21;A0A087WWI4;K7EQK2;Q9Y4L1-2 | Hypoxia up-regulated protein 1 | HYOU1 | 322070 | 151540 | 433710 | 101820 | 194440 | 95679 |
| A0A087X0H9;Q5T8P6-4;Q5T8P6;Q5T8P6-5;Q5T8P6-3;Q5T8P6-2;E2PSN0 | RNA-binding protein 26 | RBM26 | 0 | 27877 | 0 | 0 | 0 | 0 |
| A0A087X0K1;Q9Y376;B7ZBJ4;Q5TAW7;Q9H9S4 | Calcium-binding protein 39;Calcium-binding protein 39-like | CAB39;CAB39L | 1251400 | 134240 | 367400 | 80070 | 0 | 0 |
| A0A087X0M4;Q9BWU0;A0A087WWF4 | Kanadaptin | SLC4A1AP | 143000 | 42832 | 109230 | 0 | 158280 | 0 |
| A0A087X1B7;X6R700;Q9Y3Y2-4;Q9Y3Y2;Q9Y3Y2-3 | Chromatin target of PRMT1 protein | CHTOP | 1270900 | 0 | 0 | 652350 | 2350100 | 1515600 |
| A0A087X1N3;O75818-2;O75818;X6RLL4;F8WC38 | Ribonuclease P protein subunit p40 | RPP40 | 5597700 | 879800 | 2952400 | 1987900 | 2855300 | 1502100 |
| A0A087X1R3;Q8N2N9;A6QL64 | Ankyrin repeat domain-containing protein 36B;Ankyrin repeat domain-containing protein 36A | ANKRD36B;ANKRD36 | 218950 | 139400 | 274930 | 44038 | 17504 | 0 |
| A0A087X1Z3;Q9UL46;H0YM70;H0YKU2;H0YLG1;H0YLD2 | Proteasome activator complex subunit 2 | PSME2 | 254020000 | 134030000 | 228380000 | 126420000 | 119250000 | 83514000 |
| E9PDU5;A0A087X295;Q9NNW5;C9K020;E9PBK6 | WD repeat-containing protein 6 | WDR6 | 0 | 78518 | 152780 | 36323 | 151950 | 263750 |
| A0A087X2D8;O60271-9;O60271-4;O60271-2;O60271;O60271-5;O60271-3 | C-Jun-amino-terminal kinase-interacting protein 4 | SPAG9 | 34891 | 0 | 23436 | 25920 | 0 | 0 |
| A0A087X2I1;P62333;H0YJC0;H0YJS8;H0YJX2;H0YJT1;H0YJE9;H0YJD2;H0YJY8 | 26S protease regulatory subunit 10B | PSMC6 | 894850000 | 330400000 | 431160000 | 84065000 | 67662000 | 33935000 |
| Q5JXX1;Q5JXX2;A0A088AWL8;Q15014;Q5JXX6 | Mortality factor 4-like protein 2 | MORF4L2 | 2179300 | 650380 | 3262800 | 515800 | 981810 | 326120 |
| J3KP36;F8W7U3;E7ESD2;A0A096LPC5;Q5SNT6;Q9Y4E1-3;Q9Y4E1-6;Q9Y4E1-2;Q9Y4E1;Q9Y4E1-4;Q641Q2-2;Q641Q2;A0A087WYF6;Q9Y4E1-5 | WASH complex subunit FAM21B;WASH complex subunit FAM21C;WASH complex subunit FAM21A | FAM21C;FAM21A;FAM21B | 49658 | 0 | 47332 | 0 | 0 | 0 |
| A2A2Q9;Q9Y312 | Protein AAR2 homolog | AAR2 | 1568900 | 603660 | 1353000 | 170900 | 296440 | 113830 |
| A2A3N6;A6PW57;Q99755-2;Q99755-4;Q99755-3;Q99755 | Putative PIP5K1A and PSMD4-like protein | PIPSL | 91721 | 0 | 123220 | 0 | 0 | 0 |
| A2A3R5 | 40S ribosomal protein S6 | RPS6 | 2688800 | 2006000 | 3820200 | 0 | 4007000 | 2037000 |
| A2AB88;O15212 | Prefoldin subunit 6 | PFDN6 | 326260 | 1401200 | 2655500 | 811540 | 587850 | 538730 |
| A2ABK4;P18615;P18615-3;E9PD43;P18615-4 | Negative elongation factor E | NELFE | 0 | 148850 | 219180 | 266100 | 268620 | 197360 |
| A2AE51;A2AE48;Q12899 | Tripartite motif-containing protein 26 | TRIM26 | 741710 | 152550 | 135240 | 0 | 0 | 0 |
| A2IDA3;P29372-5;P29372-4;P29372-2;P29372 | DNA-3-methyladenine glycosylase | MPG | 113050 | 0 | 232000 | 0 | 0 | 0 |
| B7ZM87;A2RUF3;O75044;A0A075B7B5;Q5VZB4;A0A075B7E6;P0DMP2;P0DJJ0;A0A075B7C8;A0A075B7E9;G5EA48;Q7Z6B7-2;Q7Z6B7;A0A075B743 | SLIT-ROBO Rho GTPase-activating protein 2 | SRGAP2 | 2500500 | 1452700 | 1620800 | 97401 | 201100 | 156330 |
| A3KFL5;A3KFL2;A3KFL4;Q13868-2;Q13868;A3KFL1 | Exosome complex component RRP4 | EXOSC2 | 0 | 444320 | 153020 | 1578500 | 352410 | 282760 |
| A5YKK6;A5YKK6-4;A5YKK6-3;A5YKK6-2;H3BR89;H3BMH0;H3BVC9;H3BMZ2 | CCR4-NOT transcription complex subunit 1 | CNOT1 | 126090 | 87857 | 264480 | 62262 | 826760 | 70909 |
| H0YEY4;A6NCQ0;A6NJU6;A6NFX8;Q9UKK9 | ADP-sugar pyrophosphatase | NUDT5 | 0 | 0 | 390680 | 0 | 0 | 0 |
| A6NDF3;Q9BVG4 | Protein PBDC1 | PBDC1 | 426730 | 170720 | 837670 | 0 | 0 | 0 |
| C9J9K3;A6NE09;P08865;C9JQR9;G3V2D8;Q9UBC7;Q68DK2-4;Q68DK2-2;Q68DK2 | 40S ribosomal protein SA | RPSA;RPSAP58 | 287020000 | 194740000 | 352790000 | 227280000 | 417550000 | 224290000 |
| A6NEM2;P51610-2;P51610;P51610-4;H7C1C4 | Host cell factor 1;HCF N-terminal chain 1;HCF N-terminal chain 2;HCF N-terminal chain 3;HCF N-terminal chain 4;HCF N-terminal chain 5;HCF N-terminal chain 6;HCF C-terminal chain 1;HCF C-terminal chain 2;HCF C-terminal chain 3;HCF C-terminal chain 4;HCF C-terminal chain 5;HCF C-terminal chain 6 | HCFC1 | 40091 | 57704 | 25645 | 124250 | 209560 | 50014 |
| A6NHK2;P62304 | Small nuclear ribonucleoprotein E | SNRPE | 22689000 | 18866000 | 28101000 | 13770000 | 35470000 | 20964000 |
| A6NHR9;A6NHR9-2;J3KTL8;A6NHR9-3;J3KRK8 | Structural maintenance of chromosomes flexible hinge domain-containing protein 1 | SMCHD1 | 155600 | 51975 | 150550 | 598980 | 868740 | 290040 |
| A6NN50;O75147-2;O75147-1;O75147-4;O75147;A8MSZ8 | Obscurin-like protein 1 | OBSL1 | 296570 | 169970 | 373180 | 53087 | 96626 | 17134 |
| A8MQ02;J3KN01;P55196-3;P55196;P55196-5;Q5TIG5;P55196-2;P55196-6;P55196-1;H0Y7R8;H0Y8L4;H0Y8U8;H0Y9I0;H0Y948 | Afadin | MLLT4 | 802220 | 286690 | 259880 | 88579 | 138730 | 30835 |
| J3QLE5;A8MT02;P14678-2;P63162;P14678;P63162-2;P14678-3;J3KRY3;S4R3P3 | Small nuclear ribonucleoprotein-associated proteins B and B;Small nuclear ribonucleoprotein-associated protein N | SNRPN;SNRPB | 6036200 | 18540000 | 6415500 | 1670400 | 6180600 | 469190 |
| A8MT37;P49840 | Glycogen synthase kinase-3 alpha | GSK3A | 1089600 | 739760 | 613760 | 135070 | 218230 | 0 |
| E9PQ63;A8MTM1;P16152-2;P16152;O75828 | Carbonyl reductase [NADPH] 1;Carbonyl reductase [NADPH] 3 | CBR1;CBR3 | 0 | 0 | 243250 | 0 | 0 | 0 |
| A8MUA9;A8MU27;P55854;P55854-2 | Small ubiquitin-related modifier 3 | SUMO3 | 4651800 | 682950 | 1810000 | 0 | 681230 | 143790000 |
| F8WDX0;F8WB12;F8WDS8;C9J0U9;D3YTH5;A8MUM1;Q53HC9 | Protein TSSC1 | TSSC1 | 2081800 | 1345100 | 3261600 | 867600 | 1259400 | 680530 |
| H7BY10;K7EJV9;K7ERT8;A8MUS3;P62750;K7EMA7 | 60S ribosomal protein L23a | RPL23A | 306520000 | 165970000 | 412010000 | 230990000 | 566390000 | 255830000 |
| A8MV53;Q9NQ55-2;Q9NQ55;Q9NQ55-3;H7C446;C9J3W3 | Suppressor of SWI4 1 homolog | PPAN | 3231000 | 1418700 | 2847400 | 5748900 | 10519000 | 4881900 |
| E7EVE9;A8MX75;P18074;K7EIT8;P18074-2 | TFIIH basal transcription factor complex helicase XPD subunit | ERCC2 | 480290 | 298750 | 411130 | 117990 | 135680 | 80050 |
| A8MXL6;P55735-2;P55735;P55735-3 | Protein SEC13 homolog | SEC13 | 3476300 | 321410 | 1072300 | 0 | 0 | 0 |
| D6R991;D6REM6;A8MXP9;P43243;H0Y8T4 | Matrin-3 | MATR3 | 1501900 | 116890 | 594410 | 528680 | 336310 | 655900 |
| B8ZZ10;A8MYV2;Q9NQ29-2;Q9NQ29-3;Q9NQ29;B8ZZ09 | Putative RNA-binding protein Luc7-like 1 | LUC7L | 0 | 0 | 813530 | 0 | 0 | 0 |
| B0QY89;Q9Y262;Q9Y262-2;B0QY90;C9JHP4;C9K0Q7;H7C3A0;H0Y7E6 | Eukaryotic translation initiation factor 3 subunit L | EIF3L | 42255000 | 23920000 | 56168000 | 58493000 | 86805000 | 49091000 |
| H7BY36;B0QYK0;C9JGE3;Q01844-2;Q01844-6;Q01844-3;Q01844;Q01844-5 | RNA-binding protein EWS | EWSR1 | 10207000 | 5562800 | 11526000 | 8373800 | 15166000 | 7721600 |
| B0YIW6;P48444;Q6P1Q5;P48444-2 | Coatomer subunit delta | ARCN1 | 1177600 | 439280 | 913560 | 182260 | 436330 | 93726 |
| B1AJQ6;Q86Y82 | Syntaxin-12 | STX12 | 894360 | 180320 | 524420 | 0 | 0 | 0 |
| B1AJZ9;B1AJZ9-3;H0YCR1;Q5JYW6;H0Y5P0;H0YE38;B1AJZ9-4 | Forkhead-associated domain-containing protein 1 | FHAD1 | 633790 | 0 | 0 | 0 | 60634 | 0 |
| B1AK87;B1AK88;P47756-2;P47756;B1AK85 | F-actin-capping protein subunit beta | CAPZB | 8182700 | 2973100 | 4018600 | 0 | 2359900 | 995630 |
| B1AM48;B1AM49;Q12926-2;Q12926;C9JB16;B1APY9;B1APY8;Q14576-2;P26378-4;P26378-2;Q14576;P26378-5;P26378;P26378-3 | ELAV-like protein;ELAV-like protein 2;ELAV-like protein 3;ELAV-like protein 4 | ELAVL2;ELAVL4;ELAVL3 | 0 | 325900 | 475380 | 0 | 557820 | 259690 |
| B1AMN7;Q9P2B4 | CTTNBP2 N-terminal-like protein | CTTNBP2NL | 231080 | 59503 | 0 | 0 | 0 | 0 |
| F2Z2Y2;B1AMN9;Q9H7E2-2;Q9H7E2;Q9H7E2-3 | Tudor domain-containing protein 3 | TDRD3 | 15451000 | 19415000 | 23617000 | 14890000 | 24334000 | 19564000 |
| R4GNH9;R4GMQ7;B1AMU7;B1AMU4;Q9Y3B2;B1AMU3 | Exosome complex component CSL4 | EXOSC1 | 1241500 | 166280 | 0 | 0 | 311640 | 0 |
| B1ANW7;Q86VF7-4;Q86VF7-3;Q86VF7;Q86VF7-2 | Nebulin-related-anchoring protein | NRAP | 17392 | 30910 | 55632 | 0 | 0 | 0 |
| B1AUU8;P42566;P42566-2;S4R3U1 | Epidermal growth factor receptor substrate 15 | EPS15 | 458950 | 114010 | 306390 | 0 | 50211 | 0 |
| Q5T7C4;Q5T7C6;B2RPK0;P09429 | Putative high mobility group protein B1-like 1;High mobility group protein B1 | HMGB1;HMGB1P1 | 1418300 | 626940 | 1544700 | 486800 | 328550 | 77213 |
| B2WTI3;B2WTI4;K7EJU9;Q6NYC1-2;Q6NYC1;Q6NYC1-3 | Bifunctional arginine demethylase and lysyl-hydroxylase JMJD6 | JMJD6 | 0 | 250510 | 0 | 0 | 0 | 0 |
| B3EWF7 | Laforin, isoform 9 | EPM2A | 0 | 0 | 0 | 383600 | 669250 | 275660 |
| B3KS98;O15372;A0A087WZK9;E5RJT0;E5RGU4;E5RFW7;E5RFH0;E5RHC7;E5RH59 | Eukaryotic translation initiation factor 3 subunit H | EIF3H | 23338000 | 12602000 | 32443000 | 31412000 | 58718000 | 42356000 |
| P49959;B3KTC7;F8W7U8;P49959-2;F5GXT0;F5H742;F5H256 | Double-strand break repair protein MRE11A | MRE11A | 424640 | 430290 | 956850 | 255380 | 596380 | 133510 |
| H0YHC3;F8W020;F8W118;F8VXI6;F8VUX1;F8VY35;F8VV59;B7Z9C2;F8W0J6;F5H4R6;H0YIV4;B3KV44;P55209-2;P55209;F8VRJ2;H0YH88;F8VVB5 | Nucleosome assembly protein 1-like 1 | NAP1L1 | 0 | 0 | 0 | 618390 | 0 | 0 |
| H0Y5J4;B4DDD6;Q9UJU6-5;Q9UJU6-4;Q9UJU6;Q9UJU6-2;Q9UJU6-3;B4DDP6 | Drebrin-like protein | DBNL | 696250 | 1032900 | 1356100 | 0 | 0 | 0 |
| B4DEM9;Q9Y2S7 | Polymerase delta-interacting protein 2 | POLDIP2 | 162140 | 138250 | 181670 | 0 | 44889 | 20116 |
| B4DJ07;Q9UPT5-2;Q9UPT5-5;Q9UPT5-1;Q9UPT5-6;Q9UPT5;C9JME6;C9JKC2;B5MCY9;K7ERQ5;Q9UPT5-4 | Exocyst complex component 7 | EXOC7 | 148380 | 60411 | 268220 | 0 | 26564 | 0 |
| B4DJV2;O75390;H0YIC4;F8VWQ5;F8VRI6;F8VPF9;F8VPA1;F8VR34;F8VTT8;F8W1S4;F8W4S1;H0YH82;F8W642;F8W0J2;F8VZK9;F8VP03;F8VX07;F8VRP1;F8VX68;F8VU34 | Citrate synthase;Citrate synthase, mitochondrial | CS | 6616300 | 2425500 | 2328400 | 2020500 | 1690800 | 536030 |
| B4DJV5;Q13610 | Periodic tryptophan protein 1 homolog | PWP1 | 2107700 | 418130 | 3041700 | 1623100 | 3353900 | 371040 |
| B4DKY1;P49589-2;P49589;P49589-3 | Cysteine--tRNA ligase, cytoplasmic | CARS | 397120 | 193280 | 499370 | 79613 | 237040 | 124750 |
| B4DQP0;Q9H9L7;H0YEQ5 | Akirin-1 | AKIRIN1 | 1770800 | 785370 | 1023100 | 0 | 0 | 0 |
| H0Y2Y8;B4DQX7;Q15942;C9JJK5;H7C3D3 | Zyxin | ZYX | 3998000 | 1165200 | 983930 | 0 | 0 | 0 |
| B4DUR8;P49368;P49368-2;Q5SZX6;Q5SZX9;E9PRC8 | T-complex protein 1 subunit gamma | CCT3 | 8715700 | 3307500 | 5001200 | 3456900 | 4060600 | 1653000 |
| E9PNU1;B4DXC8;Q9Y5B8-2;Q9Y5B8 | Nucleoside diphosphate kinase 7 | NME7 | 1068500 | 357850 | 868490 | 80612 | 276490 | 67085 |
| B4DXZ6;P51114;P51114-2;P51114-3;C9JAJ4;E7EU85;E9PFF5;C9J5B4;C9IZ22;C9JZE0;C9JY20;H7C4S4 | Fragile X mental retardation syndrome-related protein 1 | FXR1 | 4398900 | 2677100 | 6554900 | 2687400 | 8072700 | 4007900 |
| G3V576;G3V4W0;B4DY08;G3V4C1;G3V2Q1;P07910-2;P07910;P07910-3;P07910-4;G3V555;G3V5V7;G3V575;G3V2D6;P0DMR1;O60812;B7ZW38;B2RXH8;G3V2H6;G3V4M8;G3V5X6;G3V3K6;G3V251;B4DSU6 | Heterogeneous nuclear ribonucleoproteins C1/C2;Heterogeneous nuclear ribonucleoprotein C-like 4;Heterogeneous nuclear ribonucleoprotein C-like 1;Heterogeneous nuclear ribonucleoprotein C-like 3;Heterogeneous nuclear ribonucleoprotein C-like 2 | HNRNPC;HNRNPCL4;HNRNPCL1;HNRNPCL3;HNRNPCL2 | 18056000 | 12278000 | 18307000 | 15902000 | 29636000 | 12246000 |
| B4DY09;Q12905;X6R6Z1 | Interleukin enhancer-binding factor 2 | ILF2 | 14476000 | 6794800 | 21030000 | 39819000 | 21499000 | 13016000 |
| Q8NBY1;B4E0Y9;Q9P289;Q9P289-2;Q9P289-3;C9JJV0;C9JDH9;C9J6L2;C9JCC0;C9J232;H7C279;O00506-3;O00506-2;O00506 | Serine/threonine-protein kinase 26 | STK26 | 3442900 | 1534600 | 2571400 | 215700 | 393390 | 252960 |
| G3V1C3;Q9BZZ5-1;Q9BZZ5-3;Q9BZZ5-5;Q9BZZ5-2;B4E283;Q9BZZ5;H0YER7 | Apoptosis inhibitor 5 | API5 | 0 | 0 | 304250 | 0 | 129150 | 100960 |
| B5MCF9;O00541-2;O00541;H7C267;B3KXD6 | Pescadillo homolog | PES1 | 0 | 0 | 171520 | 1943300 | 1652900 | 626670 |
| B5MDF5;P62826;J3KQE5;H0YFC6;F5H018;A0A087X0W0;B4DV51 | GTP-binding nuclear protein Ran | RAN | 18076000 | 7533800 | 21809000 | 4940600 | 10914000 | 3607500 |
| B6VEX4;Q8IZP0-10;Q8IZP0-8;Q8IZP0-7;Q8IZP0-2;Q8IZP0-4;Q8IZP0-3;Q8IZP0-5;Q8IZP0-6;Q8IZP0-9;Q8IZP0;E9PEZ7;E7EW77;E7EP65;F8WAL6;Q8IZP0-11;Q9NYB9-3;Q9NYB9-2;Q8IZP0-12;Q9NYB9-4;Q9NYB9 | Abl interactor 1;Abl interactor 2 | ABI1;ABI2 | 598400 | 280760 | 451020 | 54815 | 210750 | 39235 |
| B7WPG3;C9IYN3;D6W592;Q8WVV9-5;Q8WVV9-4;Q8WVV9;Q8WVV9-3;Q8WVV9-2;H7C464;C9JJZ7;H7BXH8 | Heterogeneous nuclear ribonucleoprotein L-like | HNRNPLL;HNRPLL | 544330 | 586850 | 597970 | 122340 | 109760 | 87759 |
| B7Z4B8;Q9BUJ2-4;Q9BUJ2-2;Q9BUJ2;M0R3F1;Q9BUJ2-3;M0QYZ0;M0QYM5;M0QYI8;M0R203;M0QZV6;M0R0K8 | Heterogeneous nuclear ribonucleoprotein U-like protein 1 | HNRNPUL1 | 6865100 | 3779200 | 7063300 | 6519800 | 8975700 | 5564500 |
| B7Z4J8;D6RJ07;Q9UL40-3;Q9UL40;Q9UL40-2 | Zinc finger protein 346 | ZNF346 | 889880 | 540770 | 1230800 | 416020 | 1195200 | 726010 |
| E7EQ12;H0Y7F0;H0Y9H6;E9PDE4;E7ES10;E9PCH5;E7EVY3;B7Z574;E7ESM9;P20810-3;P20810-4;P20810-8;P20810-2;P20810;P20810-9;P20810-5;P20810-10;P20810-7;P20810-6;D6RAA8;D6RC54;F8W7E0;E7EQA0;H0YD33 | Calpastatin | CAST | 614080 | 214770 | 535740 | 1433600 | 431870 | 318090 |
| B7Z6D5;Q96GQ7;A0A087X059 | Probable ATP-dependent RNA helicase DDX27 | DDX27 | 0 | 0 | 0 | 1306000 | 2433200 | 1510700 |
| B7Z7P8;P62495-2;P62495 | Eukaryotic peptide chain release factor subunit 1 | ETF1 | 2054900 | 836940 | 1881200 | 590310 | 1879000 | 916280 |
| F5H8L0;Q5R372-7;Q5R372-8;Q5R372-6;Q5R372-5;B7ZAP0 | Rab GTPase-activating protein 1-like, isoform 10 | RABGAP1L | 0 | 0 | 0 | 0 | 0 | 0 |
| E7ENZ3;B7ZAR1;E9PCA1;P48643-2;P48643 | T-complex protein 1 subunit epsilon | CCT5 | 5144900 | 2126700 | 4726100 | 5736400 | 3435500 | 1618100 |

| | | | | | | | | |
|---|---|---|---|---|---|---|---|---|
| B7ZKM0;Q15020;F8VV04;Q15020-2;F8W667;H0YHU8;Q15020-3 | Squamous cell carcinoma antigen recognized by T-cells 3 | SART3 | 2990000 | 1694600 | 2983200 | 2339600 | 2996000 | 1321200 |
| C9JMM0;B8ZZ43;S4R2Y4;Q13185 | Chromobox protein homolog 3 | CBX3 | 3712300 | 2351400 | 5258100 | 1357900 | 3711500 | 1387800 |
| B8ZZ87;Q6P582;Q6NZ67;H7BZ50;H7C173;H7C202 | Mitotic-spindle organizing protein 2A;Mitotic-spindle organizing protein 2B | MZT2B;MZT2A | 476180 | 294040 | 575560 | 0 | 0 | 0 |
| B8ZZF8;Q9Y4Y9 | U6 snRNA-associated Sm-like protein LSm5 | LSM5 | 41236000 | 18851000 | 29510000 | 21447000 | 21180000 | 5522200 |
| B8ZZN6;P63165;B8ZZJ0;B8ZZ67;B9A032;P63165-2 | Small ubiquitin-related modifier 1 | SUMO1 | 9398700 | 2370600 | 4061100 | 824560 | 1152800 | 765370 |
| B8ZZQ6;P06454-2;P06454 | Prothymosin alpha;Prothymosin alpha, N-terminally processed;Thymosin alpha-1 | PTMA | 11946000 | 4716600 | 8293600 | 12021000 | 11368000 | 7141100 |
| B8ZZS4;Q9H8Y5 | Ankyrin repeat and zinc finger domain-containing protein 1 | ANKZF1 | 247810 | 173820 | 476360 | 32946 | 127550 | 23830 |
| B8ZZU8;Q15370;Q15370-2;I3L0M9 | Transcription elongation factor B polypeptide 2 | TCEB2 | 28864000 | 12802000 | 24684000 | 14465000 | 16144000 | 12014000 |
| B9ZVN9;O95602 | DNA-directed RNA polymerase;DNA-directed RNA polymerase I subunit RPA1 | POLR1A | 533960 | 520050 | 1142200 | 419570 | 746650 | 187740 |
| C9J0D1;Q71UI9-2;Q71UI9;P0C0S5;C9J386;Q71UI9-5;Q71UI9-3;Q71UI9-4;Q99878;Q96KK5;Q9BTM1;Q16777;Q93077;Q8IUE6;Q7L7L0;Q6FI13;P20671;P0C0S8;P04908;Q96QV6;P16104;Q9BTM1-2 | Histone H2A;Histone H2A.V;Histone H2A.Z;Histone H2A type 1-J;Histone H2A type 1-H;Histone H2A.J;Histone H2A type 2-C;Histone H2A type 1-C;Histone H2A type 2-B;Histone H2A type 3;Histone H2A type 2-A;Histone H2A type 1-D;Histone H2A type 1;Histone H2A type 1-B/E;Histone H2A type 1-A;Histone H2AX | H2AFV;H2AFZ;HIST1H2AJ;HIST1H2AH;H2AFJ;HIST2H2AC;HIST1H2AC;HIST2H2AB;HIST3H2A;HIST2H2AA3;HIST1H2AD;HIST1H2AG;HIST1H2AB;HIST1H2AA;H2AFX | 9486200 | 9915100 | 13674000 | 3979600 | 6887500 | 4751300 |
| C9J0E4;P01040 | Cystatin-A;Cystatin-A, N-terminally processed | CSTA | 3280100 | 8307000 | 5131600 | 1627500 | 5146500 | 2544600 |
| C9J712;C9J0J7;C9JQ45;G5E9Q6;P35080-2;P35080 | Profilin;Profilin-2 | PFN2 | 7572600 | 3130900 | 0 | 0 | 0 | 0 |
| C9J0K6;P30626-3;P30626-2;P30626 | Sorcin | SRI | 334490 | 1081800 | 1278400 | 614550 | 867480 | 111390 |
| C9JB42;C9JDR7;C9JDX6;F8WCZ4;C9J2P2;E9PH18;C9J2C4;Q8WWF6;O75190-2;O75190-3;O75190 | DnaJ homolog subfamily B member 3;DnaJ homolog subfamily B member 6 | DNAJB6;DNAJB3 | 1746000 | 1938600 | 3046100 | 0 | 0 | 0 |
| C9J4M6;C9J2Y9;P30876 | DNA-directed RNA polymerase;DNA-directed RNA polymerase II subunit RPB2 | POLR2B | 3264800 | 2034600 | 3626500 | 978250 | 1170100 | 577110 |
| C9K0R9;C9J381;P20839-2;P20839-4;P20839;P20839-3;P20839-7;P20839-5;P20839-6;C9J029;H7C5T1;H7C511;E7ETK5;H0Y4R1;P12268 | Inosine-5-monophosphate dehydrogenase;Inosine-5-monophosphate dehydrogenase 1;Inosine-5-monophosphate dehydrogenase 2 | IMPDH1;IMPDH2 | 18578000 | 10228000 | 15968000 | 759830 | 776040 | 535650 |
| C9J384;Q9BQ75-2;Q9BQ75 | Protein CMSS1 | CMSS1 | 2045800 | 1469900 | 2434300 | 3105000 | 5866000 | 2818900 |
| C9J3N8;P04792 | Heat shock protein beta-1 | HSPB1 | 5782900 | 7409500 | 11345000 | 0 | 1274200 | 1108700 |
| G5E9R3;M0R0A1;C9J4Z3;Q6P4E4;E9PEL3;M0R2L6;P61513;A6NKH3 | 60S ribosomal protein L37a | RPL37A | 48223000 | 24168000 | 83974000 | 56501000 | 112650000 | 77687000 |
| C9J5E5;C9JIZ1;C9JHS7;C9JHZ8;C9JES8;C9JZI8;C9JBS3;C9JEJ8;A0A088AWM1;C9JHN6;C9JT62;C9JK79 | | HDLBP | 759190 | 0 | 0 | 0 | 0 | 0 |
| E9PKT8;C9J6D1;C9JZI7;Q99733;Q99733-2;E9PNW0;E9PJJ2;E9PS34;E9PNJ7;E9PKI2;C9J1B1;E9PP22;A8MXH2;H0YCI4 | Nucleosome assembly protein 1-like 4 | NAP1L4 | 2520200 | 1216700 | 2742900 | 0 | 632430 | 0 |
| C9J6N5;C9JQ40;C9JW51;Q9H098;Q9H098-2;C9J3Q3 | Protein FAM107B | FAM107B | 1044600 | 509970 | 606560 | 0 | 0 | 0 |
| C9J6P4;Q7Z2W4-3;Q7Z2W4-2;Q7Z2W4 | Zinc finger CCCH-type antiviral protein 1 | ZC3HAV1 | 439580 | 263980 | 482330 | 288030 | 494420 | 212530 |
| C9JTX4;C9J826;C9JK18;C9JKY1;P14923 | Junction plakoglobin | JUP | 124450 | 393810 | 146850 | 144040 | 260280 | 143450 |
| C9J8R4;C9JVE2;Q96GG9;C9JRU6;C9J0B2;C9JUW4 | DCN1-like protein;DCN1-like protein 1 | DCUN1D1 | 7237900 | 1231300 | 3012600 | 1035000 | 1081600 | 202020 |
| C9JA08;Q96D46;C9IY70;C9IZW9;C9K0C2;C9J0B9 | 60S ribosomal export protein NMD3 | NMD3 | 917430 | 507820 | 769890 | 0 | 923340 | 553860 |
| C9JZG2;C9JKQ7;C9JBJ1;C9JTV6;F8WD69;P22102-2;P22102 | Trifunctional purine biosynthetic protein adenosine-3;Phosphoribosylamine--glycine ligase;Phosphoribosylformylglycinamidine cyclo-ligase;Phosphoribosylglycinamide formyltransferase | GART | 640280 | 0 | 0 | 0 | 0 | 0 |
| C9JF79;P40925-2;P40925;P40925-3;C9JRL4;B9A041;B8ZZ51 | Malate dehydrogenase;Malate dehydrogenase, cytoplasmic | MDH1 | 103440 | 77871 | 464690 | 188470 | 0 | 0 |
| C9JFE4;Q13098-5;Q13098;Q13098-7;A0A096LP07;A0A096LPJ3;A8K070;Q13098-6;J3QQX0 | COP9 signalosome complex subunit 1 | GPS1 | 1299000 | 371210 | 1242000 | 336400 | 560330 | 330710 |
| C9JFV4;Q8IZL8;E7EV54 | Proline-, glutamic acid- and leucine-rich protein 1 | PELP1 | 87540 | 0 | 101660 | 84682 | 356250 | 125240 |
| C9JPA8;C9JHN8;Q7L5D6-2;Q7L5D6 | Golgi to ER traffic protein 4 homolog | GET4 | 8277300 | 0 | 1046800 | 1279400 | 0 | 0 |
| C9JI97;Q5XPI4;Q5XPI4-2;H7C3U1 | E3 ubiquitin-protein ligase RNF123 | RNF123 | 4852700 | 2322700 | 6016100 | 3420200 | 7433000 | 3088900 |
| C9JJ54;Q8IWB7 | WD repeat and FYVE domain-containing protein 1 | WDFY1 | 172560 | 0 | 0 | 0 | 0 | 0 |
| C9JLU1;P52434;P52434-4;P52434-3;P52434-2 | DNA-directed RNA polymerases I, II, and III subunit RPABC3 | POLR2H | 8633400 | 5953000 | 13916000 | 5170200 | 6401600 | 2971800 |
| C9JXB8;C9JNW5;P83731 | 60S ribosomal protein L24 | RPL24 | 71152000 | 44259000 | 104270000 | 76577000 | 85564000 | 105780000 |
| C9JPS1;Q9Y6D9-3;Q9Y6D9;C9JP81 | Mitotic spindle assembly checkpoint protein MAD1 | MAD1L1 | 295590 | 0 | 131880 | 0 | 0 | 0 |
| C9JQD4;O43447;H0YEL5;O43447-2;A6NM32;R4GN99;H0Y548;P30405-2;P30405;Q08752 | Peptidyl-prolyl cis-trans isomerase;Peptidyl-prolyl cis-trans isomerase H | PPIH | 1189900 | 1459200 | 1850800 | 922010 | 836940 | 0 |
| C9JQV0;Q9BRJ6;H7C0T1;H7C2R9 | Uncharacterized protein C7orf50 | C7orf50 | 11712000 | 5923300 | 16045000 | 12337000 | 25279000 | 10297000 |
| C9JUJ0;Q9UFW8 | CGG triplet repeat-binding protein 1 | CGGBP1 | 796430 | 1039200 | 1692400 | 1440700 | 2910500 | 976680 |
| H0Y3P2;D3DQV9;P78344-2;P78344;E9PKF8;H0YEC5;H0YD77;H0YCH5;H0YE22;H0YD99;H0YEN8;H0YDC0;H0YCF8 | Eukaryotic translation initiation factor 4 gamma 2 | EIF4G2 | 1804100 | 864020 | 2334300 | 941480 | 2501800 | 733100 |
| D3DTX6;Q96SB3 | Neurabin-2 | PPP1R9B | 106670 | 130630 | 231640 | 0 | 0 | 0 |
| D3YTB1;F8W727;P62910 | 60S ribosomal protein L32 | RPL32 | 185910000 | 76133000 | 139290000 | 140130000 | 226060000 | 150640000 |
| D6R907;D6RBI9;D6RBR6;D6RFG3;D6REL6;I6L9I8;Q9H201-2;Q9Y6I3-3;Q9Y6I3;Q9H201;Q9Y6I3-1 | Epsin-3;Epsin-1 | EPN3;EPN1 | 1071500 | 0 | 901720 | 0 | 0 | 171000 |
| D6R938;E9PBG7;E9PF82;Q13557-8;Q13557-12;Q13557-9;Q13557-10;Q13557-5;Q13557;Q13557-3;Q13557-6;Q13557-4;Q13557-11 | Calcium/calmodulin-dependent protein kinase type II subunit delta | CAMK2D | 1741700 | 466300 | 883530 | 222230 | 213660 | 81110 |
| D6RAD4;P50613;D6R9G1;D6RFL0;D6RGG9;D6RI01;D6R9Z2;D6RF14;D6RIG9;D6REC6;Q9C098 | Cyclin-dependent kinase 7 | CDK7 | 2107100 | 1402300 | 2084400 | 325840 | 522100 | 72838 |
| D6RAN4;P32969;H0Y9V9;E7ESE0;H0Y9R4;REV__O75648-5;REV__O75648-2;REV__O75648 | 60S ribosomal protein L9 | RPL9 | 47603000 | 38904000 | 73029000 | 60294000 | 127200000 | 69402000 |
| D6RD63;D6RAX7;Q9BT78-2;Q9BT78;D6RFN0 | COP9 signalosome complex subunit 4 | COPS4 | 698640 | 229040 | 657890 | 53724 | 0 | 92509 |
| D6RD47;P62266;D6RDJ2;D6RIX0;D6R9I7 | 40S ribosomal protein S23 | RPS23 | 209230000 | 90281000 | 185310000 | 80688000 | 192420000 | 60271000 |
| D6RDM7;P61086-3;P61086 | Ubiquitin-conjugating enzyme E2 K | UBE2K | 0 | 0 | 0 | 0 | 0 | 0 |
| D6REB0;Q13137-5;Q13137-2;Q13137;Q13137-3;Q13137-4 | Calcium-binding and coiled-coil domain-containing protein 2 | CALCOCO2 | 107750 | 138510 | 231040 | 0 | 0 | 0 |
| E5RGS4;O60925 | Prefoldin subunit 1 | PFDN1 | 515650 | 1489300 | 2727100 | 0 | 136040 | 336510 |
| E5RGW4 | | NPM1 | 0 | 0 | 0 | 0 | 0 | 0 |
| E5RHG8;Q15369;R4GMY8;Q15369-2 | Transcription elongation factor B polypeptide 1 | TCEB1 | 6789500 | 1940500 | 4092100 | 869470 | 1517500 | 627610 |
| E5RIM3;Q9Y263;H0YBW4 | Phospholipase A-2-activating protein | PLAA | 409630 | 307890 | 591480 | 30117 | 0 | 0 |
| E5RJ93;E5RIS7;P23193-2;P23193 | Transcription elongation factor A protein 1 | TCEA1 | 1447800 | 768370 | 1155200 | 976040 | 1127200 | 616790 |
| E5RJN7;E7EVI1;F8VPI7;E7EPX0;E7EVJ4;E9PH62;Q9NUL3-6;Q9NUL3-8;Q9NUL3-3;Q9NUL3-7;Q9NUL3-2;Q9NUL3;G5EA18;Q9NUL3-4;E5RJ67;E5RFK1 | Double-stranded RNA-binding protein Staufen homolog 2 | STAU2 | 1080700 | 805430 | 2219200 | 682320 | 2155100 | 874960 |
| E5RJR5;P63208;E7ERH2;F8W8N3;P63208-2;E5RGM3 | S-phase kinase-associated protein 1 | SKP1 | 23828000 | 8526800 | 10573000 | 13058000 | 5789500 | 2970600 |
| E5RJU9;Q86UE4;H0YBJ8 | Protein LYRIC | MTDH | 210540 | 478930 | 982610 | 673640 | 1662900 | 963780 |
| E5RK69 | Annexin | ANXA6 | 1700200 | 238540 | 574450 | 0 | 0 | 0 |
| E7EMB1;Q9UH65;E9PJM7 | Switch-associated protein 70 | SWAP70 | 737960 | 397750 | 625770 | 541170 | 422190 | 243480 |
| E7EMC7;Q13501-2;Q13501;E9PFW8 | Sequestosome-1 | SQSTM1 | 706270 | 393290 | 535180 | 245000 | 169200 | 51365 |
| E7ENQ1;G3XAA2;H7C360;G5E948;E7EN19;E7ESS2;O95819-4;O95819-2;O95819;O95819-5;O95819-3;C9J840;E7EX83;H7C0P6;C9J338;Q9UKE5-8;Q9UKE5-5;Q9UKE5-7;Q9UKE5-3;Q9UKE5-6;Q9UKE5-2;Q9UKE5-4;Q9UKE5 | Mitogen-activated protein kinase kinase kinase kinase 4 | MAP4K4 | 226030 | 110190 | 75598 | 0 | 66989 | 42050 |
| E7END7;P62820;P62820-2;Q9H0U4;E9PLD0;Q92928;A0A087WTI1;H0YLJ8;H0YL94;H0YMN7;H0YNE9;P62820-3;P61026;P51153;Q92930;P61006-2;P61006;P59190-2;P59190 | Ras-related protein Rab-1A;Ras-related protein Rab-1B;Putative Ras-related protein Rab-1C | RAB1A;RAB1B;RAB1C | 7022200 | 5360900 | 6603100 | 504960 | 1279900 | 444070 |
| E7EV99;E7ENY0;P35611-2;P35611-6;P35611;P35611-3;P35611-4;P35611-5;H0Y9H2;D6RF25;D6RJE2;H0YG19;D6RAH3 | Alpha-adducin | ADD1 | 3080100 | 1293700 | 3533500 | 0 | 0 | 0 |
| E7EPB3;P50914 | 60S ribosomal protein L14 | RPL14 | 145580000 | 54855000 | 105010000 | 70658000 | 145170000 | 83488000 |
| H0Y3Y4;G3V1Q4;E7ES33;E7EPK1;Q16181-2;Q16181 | Septin-7 | Sep-07 | 445800 | 0 | 0 | 0 | 0 | 0 |
| E7EPN9;Q9Y520-3;Q9Y520-6;Q9Y520-4;Q9Y520-5;Q9Y520;Q9Y520-7;Q9Y520-2;H7C5N8 | Protein PRRC2C | PRRC2C | 643360 | 443340 | 1208400 | 1467600 | 3208200 | 1563900 |
| E7EQL8;Q96RT7-2;Q96RT7-3;Q96RT7 | Gamma-tubulin complex component 6 | TUBGCP6 | 0 | 0 | 0 | 0 | 0 | 0 |
| E7EQR4;P15311 | Ezrin | EZR | 0 | 85219 | 93651 | 96111 | 51947 | 52423 |
| E7EQV3;P11940;A0A087WTT1;E7ERJ7;P11940-2;H0YAR2;Q9H361;H0YBN4;H0YAP2;E5RGH3;E5RH24;E5RJB9;H0YAS7;H0YAS6;H0YB75;H0YC10;H0YB86;H0YAW6;Q96DU9-2;Q96DU9 | Polyadenylate-binding protein;Polyadenylate-binding protein 1;Polyadenylate-binding protein 3 | PABPC1;PABPC3 | 16007000 | 17250000 | 27134000 | 12511000 | 27949000 | 15403000 |
| E7EVX8;E7ESA8;Q8WWY3;Q8WWY3-3;Q8WWY3-2;E7EU94;E7ESX0;E7EN72 | U4/U6 small nuclear ribonucleoprotein Prp31 | PRPF31 | 507930 | 197500 | 956640 | 0 | 0 | 0 |
| E7ESP9;P07197;E7EMV2;P07197-2 | Neurofilament medium polypeptide | NEFM | 72399000 | 54631000 | 137600000 | 12253000 | 8505300 | 5516700 |
| E7ETC2;Q5F2F8;Q08209-5;Q08209-3;Q08209-2;P16298-2;P16298-3;Q08209;P16298;P16298-4;Q5F2G0;Q08209-4 | Serine/threonine-protein phosphatase;Serine/threonine-protein phosphatase 2B catalytic subunit alpha isoform;Serine/threonine-protein phosphatase 2B catalytic subunit beta isoform | PPP3CA;PPP3CB | 394910 | 0 | 121920 | 0 | 119440 | 0 |
| E7ETU9;O00469;O00469-2;O00469-3 | Procollagen-lysine,2-oxoglutarate 5-dioxygenase 2 | PLOD2 | 568910 | 239940 | 558550 | 0 | 38315 | 0 |
| J3KQ96;E7ETY2;Q13428-6;Q13428-7;Q13428;Q13428-3;Q13428-4;Q13428-2;Q13428-8;Q13428-5 | Treacle protein | TCOF1 | 29247 | 137650 | 459970 | 118860 | 368370 | 180780 |
| E7ETY4;E9PC69;Q7KZI7-10;Q7KZI7-13;Q7KZI7-7;Q7KZI7-2;Q7KZI7-15;Q7KZI7-5;Q7KZI7-16;Q7KZI7-4;Q7KZI7-9;Q7KZI7-12;Q7KZI7-14;Q7KZI7-3;Q7KZI7-6;Q7KZI7-11;Q7KZI7-8;Q7KZI7;K7EK17;K7EKG8;F5H6N2;H0YKJ3;F5H4F6;H0YNV4;J3KNR0;A0A087X0I6;P27448-8;Q96L34-2;P27448-6;P27448-3;P27448-4;Q96L34;P27448-2;P27448;P27448-7;Q9P0L2 | Serine/threonine-protein kinase MARK2;MAP/microtubule affinity-regulating kinase 3;MAP/microtubule affinity-regulating kinase 4;Serine/threonine-protein kinase MARK1 | MARK2;MARK4;MARK3;MARK1 | 77421 | 29044 | 41970 | 0 | 29318 | 18406 |
| E7ETZ4;Q9Y6E2;Q75MG1;B5MCH7;C9JF98;E7EMS9;B5MCE7;Q9Y6E2-2;E9PFE3;F8WDX8;E9PFD4 | Basic leucine zipper and W2 domain-containing protein 2 | BZW2 | 5228300 | 3265200 | 6867200 | 1119800 | 2559900 | 1085800 |
| H0Y867;E7EU81;F1T0J2;Q14789;Q14789-2 | Golgin subfamily B member 1 | GOLGB1 | 1346100 | 379250 | 587240 | 165230 | 170740 | 0 |
| E7EU96;Q5U5J2;Q8NEV1;P68400;A0A087WY74;P68400-2 | Casein kinase II subunit alpha 3;Casein kinase II subunit alpha | CSNK2A1;CSNK2A3 | 1778500 | 1228900 | 2188600 | 1355900 | 3369400 | 1731900 |
| E7EUR8;REV__Q86UP2-2;REV__Q86UP2-3;REV__Q86UP2-4;REV__Q86UP2;P78312-4;P78312-6;P78312-2;P78312-5;P78312 | Protein FAM193A | FAM193A | 1228200 | 313830 | 1337200 | 756380 | 2386300 | 362400 |
| E7EVA0;P27816-6;P27816;P27816-2;P27816-4;P27816-5;H7C4C5;B5MEG9;H0Y2V1;F8W9U4;H7C456 | Microtubule-associated protein;Microtubule-associated protein 4 | MAP4 | 3208900 | 1866200 | 5619300 | 6547900 | 9687600 | 4796900 |
| E7EVJ5;Q96F07-2;Q96F07;H7C229;E7EW33;A0A087WWZ1;A0A087WTQ3;Q7L576-2;A0A087WU52;A0A087WWL1;H0YL50;A0A087WVE1;A0A087X0D6;REV__Q8IUG5-2;REV__Q8IUG5;REV__A0A075B6F5;REV__F5GXR6;Q7L576-3 | Cytoplasmic FMR1-interacting protein 2 | CYFIP2 | 1399600 | 662610 | 1339200 | 79259 | 314200 | 60217 |
| E7EW05;Q9NVU7-2;Q9NVU7 | Protein SDA1 homolog | SDAD1 | 505990 | 456910 | 1576200 | 608030 | 1690400 | 927980 |
| E9PID8;E7EWR4;P33240-2;P33240 | Cleavage stimulation factor subunit 2 | CSTF2 | 615940 | 450930 | 685840 | 321070 | 379780 | 112340 |
| E7EX17;P23588;B4DS13;F8VP89;F8W0K0;F8VSC7;F8VX11;F8VRU1;F8VYE9 | Eukaryotic translation initiation factor 4B | EIF4B | 24428000 | 14397000 | 19571000 | 8279600 | 15425000 | 6400300 |
| E7EX90;Q14203-5;Q14203-2;Q14203-3;Q14203-4;Q14203-6;Q14203;Q6AWB1;REV__H0Y8C5;REV__Q5VTR2 | Dynactin subunit 1 | DCTN1;DKFZp686E0752 | 806630 | 464230 | 579900 | 1606000 | 1556500 | 196910 |
| E7EXA6;Q8WVB6-3;Q8WVB6;Q8WVB6-2;Q15813;Q15813-2 | Chromosome transmission fidelity protein 18 homolog;Tubulin-specific chaperone E | CHTF18;TBCE | 336250 | 45763 | 221500 | 40068 | 50947 | 38326 |
| F8VZJ2;F8W0W4;H0YHX9;Q13765;E9PAV3-2;E9PAV3;F8W1N5;F8VNW4 | Nascent polypeptide-associated complex subunit alpha;Nascent polypeptide-associated complex subunit alpha, muscle-specific form | NACA | 51064000 | 46951000 | 127780000 | 17789000 | 80256000 | 28901000 |
| E9PB61;Q86V81 | THO complex subunit 4 | ALYREF | 0 | 0 | 1271800 | 216460 | 3828100 | 0 |
| E9PB93;O15514;E9PHV4 | DNA-directed RNA polymerase II subunit RPB4 | POLR2D | 4816800 | 882320 | 4580300 | 0 | 1491000 | 721760 |
| E9PBS1;P22234;P22234-2;D6RF62 | Multifunctional protein ADE2;Phosphoribosylaminoimidazole-succinocarboxamide synthase;Phosphoribosylaminoimidazole carboxylase | PAICS | 746060 | 247960 | 519020 | 119820 | 236120 | 121550 |
| E9PC74;Q13144;C9JRD9;H7C2X0 | Translation initiation factor eIF-2B subunit epsilon | EIF2B5 | 1019700 | 1081800 | 1690700 | 1470600 | 3659400 | 1572000 |
| E9PCY7;G8JLB6;P31943;D6RBM0;D6RIU0;D6RAM1;D6R9T0;D6RFM3;D6RIT2;D6RDU3;D6RJ04;D6RIH9;E7EQJ0;E5RGV0;E7EN40;P55795;E5RGH4;D6RDL0;H0YAQ2;H0YB39;D6RF17;H0YBD7;D6R9D3;H0YBG7;E5RJ94;H7C2J6;Q9NQA5 | Heterogeneous nuclear ribonucleoprotein H;Heterogeneous nuclear ribonucleoprotein H, N-terminally processed;Heterogeneous nuclear ribonucleoprotein H2 | HNRNPH1;HNRNPH2 | 40544000 | 15840000 | 36381000 | 27139000 | 22796000 | 9772200 |
| E9PD53;Q9NTJ3;Q9NTJ3-2;C9JR83;C9JVD8;C9J578;C9J9E4 | Structural maintenance of chromosomes protein;Structural maintenance of chromosomes protein 4 | SMC4 | 3009700 | 1659600 | 4604900 | 1612700 | 3754800 | 1198600 |
| E9PDE8;O95757;D6RJ96 | Heat shock 70 kDa protein 4L | HSPA4L | 105110 | 39154 | 97299 | 20689 | 22937 | 25465 |
| E9PDP5;Q8IWZ3;Q8IWZ3-4;Q8IWZ3-6;H7C2F5;H3BLS9;O75179-3;D6RHC4;Q8IWZ3-5;Q8IWZ3-3;Q8IWZ3-2;O75179-5 | Ankyrin repeat and KH domain-containing protein 1 | ANKHD1 | 0 | 0 | 0 | 0 | 0 | 0 |
| E9PE20;P49458-2;P49458 | Signal recognition particle 9 kDa protein | SRP9 | 0 | 0 | 11210000 | 0 | 2026500 | 0 |
| E9PEB5;Q96AE4;Q96AE4-2;C9JSZ1 | Far upstream element-binding protein 1 | FUBP1 | 1307400 | 644240 | 1009700 | 510770 | 519320 | 247010 |
| E9PF58;Q92747-2;Q92747 | Actin-related protein 2/3 complex subunit 1A | ARPC1A | 927570 | 423290 | 485050 | 0 | 0 | 0 |
| E9PFK5;P78316-2;P78316 | Nucleolar protein 14 | NOP14 | 8487.4 | 0 | 103330 | 21372 | 273130 | 141160 |
| E9PH82;Q8NCA5-2;Q8NCA5 | Protein FAM98A | FAM98A | 779400 | 709420 | 1580300 | 60991 | 1155900 | 621270 |
| E9PHA2;Q15003-2;Q15003;H7C415;C9J470 | Condensin complex subunit 2 | NCAPH | 54535 | 22980 | 297670 | 0 | 0 | 0 |
| E9PHH3;H7C1K4;P18827 | Syndecan-1 | SDC1 | 0 | 0 | 448810 | 0 | 0 | 0 |
| E9PHY5;O43491-3;O43491-4;O43491;I6L9B1;E9PII3;E9PK52;O43491-2;Q6ZSX4;E9PN54;Q6R5J7 | Band 4.1-like protein 2 | EPB41L2 | 409840 | 107510 | 446490 | 0 | 25160 | 26603 |
| E9PSE5;E9PIA8;P50897-2;P50897 | Palmitoyl-protein thioesterase 1 | PPT1 | 1877900 | 1885600 | 3644200 | 0 | 0 | 0 |
| E9PID5;F8VU88;F8VWR9;E9PPF6;E9PJZ7;G3V200;Q13136-2;Q13136;O75334-2;O75334-4;O75334-3;O75334 | Liprin-alpha-1;Liprin-alpha-2 | PPFIA1;PPFIA2 | 462050000 | 259800000 | 363940000 | 52112000 | 49802000 | 38355000 |
| E9PIF2;Q13206 | Probable ATP-dependent RNA helicase DDX10 | DDX10 | 38423 | 24176 | 45502 | 56994 | 242840 | 89183 |
| E9PJD7 | | CYHR1 | 0 | 0 | 0 | 0 | 0 | 0 |
| E9PJF0;E9PQW4;P27361-3;P27361;H0YEX6;E9PRH7;B3KR49;E9PBK7;P27361-2 | Mitogen-activated protein kinase;Mitogen-activated protein kinase 3 | MAPK3 | 1183700 | 491670 | 619320 | 157280 | 122410 | 243380 |
| E9PMI6;E9PJF4;J3KN38;P54105 | Methylosome subunit pICln | CLNS1A | 0 | 482040 | 1016300 | 0 | 114330 | 0 |
| E9PK85;P62070-2;P62070-3;P62070;P62070-4 | Ras-related protein R-Ras2 | RRAS2 | 291620 | 0 | 841770 | 0 | 0 | 0 |
| E9PKG1;H7C2I1;Q99873-3;Q99873-2;Q99873-4;Q99873;E9PQ98;E9PIX6;E9PNR9;H0YDE4;A0A087X1W2;Q9NR22-2;Q9NR22 | Protein arginine N-methyltransferase 1 | PRMT1 | 6038200 | 3901500 | 5752800 | 3968600 | 4604200 | 2682800 |
| E9PKH2;P50454;H0YEP8 | Serpin H1 | SERPINH1 | 913990 | 289320 | 817920 | 0 | 209910 | 0 |
| E9PKP7;P17480-2;P17480;E9PLT2 | Nucleolar transcription factor 1 | UBTF | 25127 | 0 | 184100 | 28551 | 0 | 0 |
| F8VSA6;E9PS38;S4R3E9;E9PL57;Q15843 | NEDD8 | NEDD8;NEDD8-MDP1 | 7728700 | 7912900 | 10125000 | 4615600 | 6358400 | 3616700 |
| E9PL69;P23921 | Ribonucleoside-diphosphate reductase large subunit | RRM1 | 0 | 0 | 0 | 234000 | 1057700 | 606950 |
| E9PNH5;E9PLI6;E9PS76;Q8N9N8 | Probable RNA-binding protein EIF1AD | EIF1AD | 5052300 | 600560 | 950690 | 1049300 | 1005100 | 0 |
| E9PLL6;P46776;E9PJD9;E9PLX7 | 60S ribosomal protein L27a | RPL27A | 174380000 | 124350000 | 224610000 | 16471000 | 175180000 | 92615000 |
| E9PRZ9;E9PM92;O00193 | Small acidic protein | C11orf58;SMAP | 28930000 | 8997000 | 16508000 | 394320 | 6109200 | 1620600 |
| E9PN89 | | HSPA8 | 3400300 | 1451200 | 102310 | 1991700 | 1715000 | 1880400 |
| E9PS42;E9PND2;E9PP21;P21291 | Cysteine and glycine-rich protein 1 | CSRP1 | 1461100 | 431420 | 0 | 0 | 0 | 0 |
| E9PNR8;Q9NWT6 | Hypoxia-inducible factor 1-alpha inhibitor | HIF1AN | 0 | 609160 | 1004300 | 0 | 0 | 0 |
| E9PPY3;O43159 | Ribosomal RNA-processing protein 8 | RRP8 | 0 | 488710 | 113150 | 1239100 | 2162400 | 927280 |
| E9PQY2;Q9NQP4 | Prefoldin subunit 4 | PFDN4 | 0 | 0 | 341970 | 266780 | 283430 | 326400 |

| | | | | | | | | |
|---|---|---|---|---|---|---|---|---|
| E9PR30;P62861 | 40S ribosomal protein S30 | FAU | 5925900 | 10762000 | 10978000 | 115860 | 4587400 | 2896300 |
| H0YFP2;E9PS41;Q6RFH5-2;Q6RFH5 | WD repeat-containing protein 74 | WDR74 | 1347900 | 0 | 1083300 | 1483800 | 3656000 | 1690900 |
| F2Z2B9;Q9BSJ2;Q9BSJ2-4;Q9BSJ2-3;B3KTU7 | Gamma-tubulin complex component 2 | TUBGCP2 | 550780 | 121870 | 719860 | 26857 | 0 | 0 |
| F2Z2E2;Q86VI3;E9PDT6;E7EWC2;F5H7S7;Q13576-3;Q13576-2;Q13576 | Ras GTPase-activating-like protein IQGAP3 | IQGAP3 | 0 | 0 | 0 | 0 | 0 | 0 |
| F2Z2K0;Q9UNZ2-6;Q9UNZ2;Q9UNZ2-5 | NSFL1 cofactor p47 | NSFL1C | 1087000 | 384080 | 802440 | 0 | 233940 | 0 |
| F2Z2W6;F2Z2Y5;H7BXJ5;P05114 | Non-histone chromosomal protein HMG-14 | HMGN1 | 0 | 0 | 0 | 0 | 0 | 0 |
| F2Z2W7;Q8IZ69-2;Q8IZ69;C9K041;H7C100 | tRNA (uracil-5-)-methyltransferase homolog A | TRMT2A | 1635400 | 436100 | 1508100 | 425680 | 1877300 | 1054900 |
| F2Z388;P42766;REV__Q9P209 | 60S ribosomal protein L35 | RPL35 | 426500000 | 263210000 | 615250000 | 263490000 | 748650000 | 538900000 |
| F5GWA4;Q9H3F6-2;Q9H3F6 | BTB/POZ domain-containing adapter for CUL3-mediated RhoA degradation protein 3 | KCTD10 | 1050500 | 816390 | 942980 | 0 | 129040 | 186470 |
| F5GWX5;Q14839;Q14839-2;O75052 | Chromodomain-helicase-DNA-binding protein 4 | CHD4 | 61650 | 8428.8 | 174190 | 0 | 62363 | 15643 |
| F5H5V4;F5GX23;J3KN29;O00233-2;O00233;F5H7X1;F5H169 | 26S proteasome non-ATPase regulatory subunit 9 | PSMD9 | 776650 | 209660 | 654650 | 0 | 0 | 0 |
| F5GX64;P38935;F5H5K3;H3BRR1 | DNA-binding protein SMUBP-2 | IGHMBP2 | 0 | 204670 | 213420 | 0 | 110370 | 98130 |
| F5GX77;Q9UI30-2;Q9UI30;F5GYQ2 | Multifunctional methyltransferase subunit TRM112-like protein | TRMT112 | 11499000 | 6920800 | 10811000 | 6893400 | 6034700 | 4471800 |
| F5GXT8;F5H796;Q9NR28-2;Q9NR28-3;Q9NR28 | Diablo homolog, mitochondrial | DIABLO | 595000 | 857660 | 1783100 | 0 | 501190 | 0 |
| F5GXV8;F5GY08;J3KPD3;Q9Y580;F5H674;F5H606;G3V1T9;I3L521 | RNA-binding protein 7 | RBM7 | 1386200 | 598140 | 959710 | 2389100 | 2859900 | 1486300 |
| F5GXW1;H0Y7H7;H0Y599;Q8N1I0-2;Q8N1I0;Q8N1I0-3 | Dedicator of cytokinesis protein 4 | DOCK4 | 0 | 0 | 17258 | 0 | 0 | 0 |
| F5GXW5;Q99622;U3KQ85;U3KQ07 | Protein C10 | C12orf57 | 2552400 | 430470 | 2082600 | 0 | 508990 | 0 |
| F5GY05;Q96EK7-2;Q96EK7 | Constitutive coactivator of peroxisome proliferator-activated receptor gamma | FAM120B | 813140 | 396170 | 565840 | 394060 | 543280 | 161370 |
| F5GYK2;Q9NRL3;Q9NRL3-3;M0QXN2;M0R2A7;M0QYS2;M0R0P4;M0R0W8;M0R317;R4GN16;M0QZM0;Q9NRL3-2 | Striatin-4 | STRN4 | 1069200 | 230860 | 366180 | 0 | 116750 | 0 |
| F5H867;F5H0E2;H0YFS2;F5GZS6;J3KPF3;P08195-2;P08195-3;P08195;P08195-4 | 4F2 cell-surface antigen heavy chain | SLC3A2 | 343720 | 124160 | 755650 | 0 | 127460 | 0 |
| H0YJ30;G3V582;F5H039;Q9NQX3;Q9NQX3-2 | Gephyrin;Molybdopterin adenylyltransferase;Molybdopterin molybdenumtransferase | GPHN | 0 | 0 | 200330 | 0 | 0 | 0 |
| F5H1D6;Q07864;F5H7E4 | DNA polymerase;DNA polymerase epsilon catalytic subunit A | POLE | 65249 | 30047 | 207180 | 8517.3 | 33184 | 27452 |
| F5H1S9;G8JLB3;Q9Y606-2;Q9Y606;F5GY32 | tRNA pseudouridine synthase;tRNA pseudouridine synthase A, mitochondrial | PUS1 | 187510 | 90176 | 1078300 | 250330 | 656180 | 413150 |
| F5H2F4;P11586 | C-1-tetrahydrofolate synthase, cytoplasmic;Methylenetetrahydrofolate dehydrogenase;Methenyltetrahydrofolate cyclohydrolase;Formyltetrahydrofolate synthetase;C-1-tetrahydrofolate synthase, cytoplasmic, N-terminally processed | MTHFD1 | 1710000 | 732770 | 1798000 | 831040 | 1893900 | 618020 |
| F5H2T0;O95163 | Elongator complex protein 1 | IKBKAP | 811810 | 340050 | 854860 | 316290 | 437110 | 117140 |
| F5H5U2;Q8NHQ9;E7EX41;Q8NHQ9-2 | ATP-dependent RNA helicase DDX55 | DDX55 | 733680 | 331510 | 1347900 | 874640 | 1617000 | 595770 |
| F5H669;Q8N684-2;Q8N684;Q8N684-3;J3QT54;F5H6M0;F5H047;F5H5B7;F5H1W4;F5H2K8;F5H6P5;F5H6A8;F5GXA3;C9JM38;C9J286;C9J323 | Cleavage and polyadenylation specificity factor subunit 7 | CPSF7 | 1845700 | 1812800 | 2367400 | 950410 | 1195300 | 478500 |
| F5H6T1;P61160;P61160-2 | Actin-related protein 2 | ACTR2 | 214560 | 0 | 0 | 0 | 0 | 0 |
| F5H8J2;H7BZ94;P07237;H0Y3Z3;I3NI03;I3L398;I3L0S0;I3L3U6;I3L312;I3L514;I3L4M2;I3L1Y5;I3L3P5 | Protein disulfide-isomerase | P4HB | 6675100 | 2417500 | 4788500 | 2398000 | 2838400 | 999490 |
| H3BTB3;H3BMQ5;H3BN59;I3L0H8;F6QDS0;H3BQK0;Q9UMR2-2;Q9UMR2-4;Q9NUU7;Q9UMR2 | ATP-dependent RNA helicase DDX19B;ATP-dependent RNA helicase DDX19A | DDX19A;DDX19B | 222740 | 71822 | 851860 | 0 | 0 | 0 |
| F6S8N6;H7BY58;P22061;P22061-2 | Protein-L-isoaspartate O-methyltransferase;Protein-L-isoaspartate(D-aspartate) O-methyltransferase | PCMT1 | 334990 | 512960 | 776120 | 519300 | 589330 | 354060 |
| F6WQW2;P43487-2;P43487;C9JXG8;C9JJ34;C9JDM3;C9JGV6;C9JIC6;Q5T0Z6;Q9C010;Q9C010-2 | Ran-specific GTPase-activating protein | RANBP1 | 9087900 | 3737800 | 8823600 | 3178500 | 4253700 | 2442800 |
| F8VNY3;P45973 | Chromobox protein homolog 5 | CBX5 | 1829800 | 0 | 0 | 0 | 0 | 0 |
| F8VQF4;F8VRS8;F8W145;F8VP97;F8VV52;Q9NZN8-2;Q9NZN8-4;Q9NZN8 | CCR4-NOT transcription complex subunit 2 | CNOT2 | 0 | 443990 | 773000 | 0 | 145610 | 0 |
| F8VPD4;P27708;H7C2E4 | CAD protein;Glutamine-dependent carbamoyl-phosphate synthase;Aspartate carbamoyltransferase;Dihydroorotase | CAD | 261880 | 90403 | 222990 | 153320 | 261960 | 145900 |
| H0YH87;F8WB06;F8VQP2;V9GY86;Q99700-2;Q24JQ7;Q99700-4;Q99700;F8VVY6;F8W0B5 | Ataxin-2 | ATXN2 | 0 | 0 | 0 | 84705 | 86428 | 38493 |
| F8VR82;F8VYE8;P36873;P36873-2;A0A087WYY5;F8W0W8 | Serine/threonine-protein phosphatase;Serine/threonine-protein phosphatase PP1-gamma catalytic subunit | PPP1CC | 4366700 | 1036000 | 5094600 | 2440800 | 3117000 | 1744400 |
| F8VTY2;Q8NFH4 | Nucleoporin Nup37 | NUP37 | 1089300 | 463910 | 1193500 | 0 | 54105 | 0 |
| F8VV32;P61626 | Lysozyme;Lysozyme C | LYZ | 975800 | 1274300 | 1844200 | 400550 | 928960 | 452300 |
| F8VVL1;O43583 | Density-regulated protein | DENR | 4336100 | 2710700 | 4918800 | 1819900 | 1469500 | 0 |
| F8VW28 | | PPP1R12A | 0 | 0 | 385420 | 0 | 201420 | 0 |
| F8VW50;Q2TAC6;Q2TAC6-3;Q2TAC6-2 | Kinesin-like protein;Kinesin-like protein KIF19 | KIF19 | 753660 | 450370 | 850290 | 532460 | 1107500 | 368490 |
| F8VXG7;Q99590-2;Q99590 | Protein SCAF11 | SCAF11 | 44781 | 54002 | 59723 | 24490 | 0 | 0 |
| F8VXU5;Q9UBQ0;Q9UBQ0-2;Q05DG7 | Vacuolar protein sorting-associated protein 29 | VPS29 | 21089000 | 6842700 | 13184000 | 1615500 | 1282100 | 513920 |
| F8VZY9;P05783;CON__H-INV:HIT000015463 | Keratin, type I cytoskeletal 18 | KRT18 | 4446500 | 7823400 | 11641000 | 9852800 | 9213500 | 12445000 |
| F8W031;F8W1K5;F8VXJ7;Q9Y2B0 | Protein canopy homolog 2 | CNPY2 | 103120 | 80186 | 392390 | 0 | 102330 | 46310 |
| F8W1I6;Q13561;Q13561-3;Q13561-2;F8VW18;F8VX93;H0YHL1;F8VRV7;H0YI98;F8W0U6 | Dynactin subunit 2 | DCTN2 | 3862500 | 860590 | 1061500 | 1034700 | 475780 | 430050 |
| F8W1R7;G3V1V0;P60660-2;P60660;J3KND3;G8JLA2;B7Z6Z4;G3V1Y7;F8VPF3;H0YI43;F8VZU9;F8W1I5;F8W180;P14649;F8VXL3 | Myosin light polypeptide 6 | MYL6 | 20976000 | 12882000 | 19931000 | 1705800 | 2170700 | 1134200 |
| F8W543 | | NAP1L1 | 0 | 0 | 0 | 0 | 277290 | 0 |
| F8W6I7;P09651-3;P09651-2;P09651;F8VTQ5;Q32P51;F8VZ49;F8W646;H0YH80;F8VYN5 | Heterogeneous nuclear ribonucleoprotein A1;Heterogeneous nuclear ribonucleoprotein A1, N-terminally processed;Heterogeneous nuclear ribonucleoprotein A1-like 2 | HNRNPA1;HNRNPA1L2 | 41893000 | 22855000 | 59377000 | 45905000 | 71771000 | 41986000 |
| F8W726;Q14157-4;Q14157-1;Q14157-3;Q14157;Q14157-5;H0Y5H6;H7C2T8 | Ubiquitin-associated protein 2-like | UBAP2L | 118340 | 0 | 318290 | 0 | 154910 | 25040 |
| F8W7C6;A0A087WV22;H7C123;H7C2C5 | | RPL10 | 65723000 | 30676000 | 81460000 | 47512000 | 65891000 | 24320000 |
| J3QSX6;Q5JVV3;H0Y3K7;F8W8M4;Q5T6N4;O14639-4;O14639-3;O14639-5;O14639-2;O14639-6;O14639 | Actin-binding LIM protein 1 | ABLIM1 | 0 | 0 | 0 | 0 | 0 | 0 |
| F8W914;Q9NQC3-2;Q9NQC3-5;Q9NQC3;H7C106;Q9NQC3-3;Q9NQC3-4;Q9NQC3-6 | Reticulon;Reticulon-4 | RTN4 | 3953700 | 3360900 | 4696900 | 637880 | 1253200 | 609500 |
| F8W930;Q9Y6M1-5;Q9Y6M1-6;Q9Y6M1-3;Q9Y6M1-4;Q9Y6M1-1;Q9Y6M1 | Insulin-like growth factor 2 mRNA-binding protein 2 | IGF2BP2 | 6053000 | 2655000 | 9634900 | 6948900 | 14136000 | 6350400 |
| F8WAJ0;Q9H8H2-4;Q9H8H2-2;Q9H8H2 | Probable ATP-dependent RNA helicase DDX31 | DDX31 | 0 | 27968 | 135810 | 117950 | 431430 | 202650 |
| F8WAS9;Q6P1Q9 | Methyltransferase-like protein 2B | METTL2B | 3355400 | 0 | 3530300 | 2302700 | 1285800 | 1916300 |
| F8WB69;O94888 | UBX domain-containing protein 7 | UBXN7 | 645110 | 542250 | 1555800 | 0 | 0 | 0 |
| F8WCI3;Q96SN8-4;Q96SN8-3;Q96SN8-2;Q96SN8;F8WF55;B1AMJ5 | CDK5 regulatory subunit-associated protein 2 | CDK5RAP2 | 0 | 0 | 0 | 0 | 0 | 0 |
| F8WCR5;H7C2U6;Q9BPW8 | Protein NipSnap homolog 1 | NIPSNAP1 | 1109100 | 977130 | 2933200 | 0 | 862570 | 0 |
| F8WD66;P86791;P86790 | Vacuolar fusion protein CCZ1 homolog;Vacuolar fusion protein CCZ1 homolog B | CCZ1B;CCZ1 | 958780 | 0 | 359300 | 0 | 0 | 0 |
| F8WDT8;G3V0G3;Q9NY93-2;Q9NY93;F8WEI3;H7C3E9 | Probable ATP-dependent RNA helicase DDX56 | DDX56 | 448710 | 505190 | 1028400 | 308260 | 1137300 | 686790 |
| F8WJN3;Q16630-3;Q16630;Q16630-2;F8W084 | Cleavage and polyadenylation specificity factor subunit 6 | CPSF6 | 5789300 | 4165700 | 6280300 | 274460 | 492680 | 395580 |
| G3V0E5;P02786 | Transferrin receptor protein 1;Transferrin receptor protein 1, serum form | TFRC | 1567200 | 206240 | 949360 | 0 | 0 | 0 |
| G3V1D4;H0YIA8;Q9NUP9;O14910;J3KN23;H0YI92;Q9HAP6 | Protein lin-7 homolog C;Protein lin-7 homolog A;Protein lin-7 homolog B | LIN7C;LIN7A;LIN7B | 2145700 | 974130 | 2253500 | 0 | 971650 | 745860 |
| G3V203;J3QQ67;Q07020;H0YHA7;Q07020-2;F8VUA6;F8VYV2;A0A075B7A0;F8VXR6 | 60S ribosomal protein L18 | RPL18 | 229060000 | 161790000 | 301890000 | 297160000 | 320390000 | 242050000 |
| V9GY54;G3V3G9;Q5TAQ9 | DDB1- and CUL4-associated factor 8 | DCAF8 | 306220 | 122380 | 236690 | 0 | 0 | 0 |
| G3V3M6;P27695;G3V3C7;G3V5Q1;G3V5D9;G3V5M0;G3V359;G3V2D9;H7C4A8;G3V3Y6 | DNA-(apurinic or apyrimidinic site) lyase;DNA-(apurinic or apyrimidinic site) lyase, mitochondrial | APEX1 | 23034000 | 11515000 | 26906000 | 28605000 | 46671000 | 27256000 |
| G3V4F7;P61011-2;P61011 | Signal recognition particle 54 kDa protein | SRP54 | 1015800 | 405520 | 467490 | 549570 | 1211700 | 386820 |
| G3V4P8;P60983;G3V3X4;M0R1D2;M0R0C1;M0QYJ8;O60234 | Glia maturation factor beta | GMFB | 23428000 | 6373900 | 10990000 | 121680 | 0 | 104610 |
| G3V5E4;G3V4W4;Q96EK6 | Glucosamine 6-phosphate N-acetyltransferase | GNPNAT1 | 0 | 0 | 1138600 | 0 | 0 | 0 |
| G3V4X1 | | PSMC1 | 9427100 | 4158500 | 6643500 | 155450 | 0 | 0 |
| G3V4X5 | | PSMA3 | 85523000 | 56508000 | 64123000 | 35545000 | 33421000 | 32731000 |
| G3V529;Q9GZR7;Q9GZR7-2;F5GYL3;A0A087WXU8 | ATP-dependent RNA helicase DDX24 | DDX24 | 71051 | 156720 | 498070 | 595010 | 1400100 | 502980 |
| H0YIZ6;G3V5A5;G3V541;Q9H6D7;H0YJ09;G3V5Z2;G3V4X7;Q9H6D7-3;Q9H6D7-4;G3V2I5;Q9H6D7-2 | HAUS augmin-like complex subunit 4 | HAUS4 | 468060 | 632750 | 355300 | 88936 | 0 | 0 |
| G3V5T9;P24941-2;P24941;E7ESI2;G3V317 | Cyclin-dependent kinase 2 | CDK2 | 1317500 | 475250 | 1192200 | 148050 | 462160 | 352240 |
| G3V5Z7;P60900;G3V295;P60900-2;G3V3I1;P60900-3;G3V3U4;H0YJC4;G3V4S5;G3V2S7 | Proteasome subunit alpha type;Proteasome subunit alpha type-6 | PSMA6 | 2123600000 | 1304600000 | 1535600000 | 1063900000 | 893990000 | 621240000 |
| G3XAG3;J3QRG6;P42771-3;P42771;P42771-4 | Cyclin-dependent kinase inhibitor 2A | CDKN2A | 0 | 388660 | 381470 | 0 | 0 | 0 |
| G3XAM7;P35221;P35221-2;P35221-3;E5RJ41;E5RIB1;P26232-4;P26232-6;P26232-3;P26232-2;P26232-5;P26232 | Catenin alpha-1 | CTNNA1 | 271690 | 33879 | 102140 | 0 | 84422 | 18781 |
| G5E933;O95248;O95248-4 | Myotubularin-related protein 5 | SBF1 | 0 | 0 | 0 | 0 | 47536 | 34588 |
| G5E972;P42167-3;P42167-2;P42167;P42166 | Lamina-associated polypeptide 2, isoforms beta/gamma;Thymopoietin;Thymopentin;Lamina-associated polypeptide 2, isoform alpha;Thymopoietin;Thymopentin | TMPO | 50908 | 0 | 220610 | 0 | 177150 | 28810 |
| G5E9D5;Q9BQ52-4;Q9BQ52;E7ES68;Q9BQ52-3;J3QRS2;V9GYU5;J3QL08;J3QLK4;V9GYS7;Q9BQ52-2;V9GZ72 | Zinc phosphodiesterase ELAC protein 2 | ELAC2 | 634050 | 367350 | 743250 | 213090 | 614930 | 336910 |
| G5EA31;P53992-2;P53992 | Protein transport protein Sec24C | SEC24C | 306620 | 162470 | 146300 | 82809 | 57884 | 0 |
| H0Y3X6;Q9H3P2;H7C2C7;Q9H3P2-7;C9JEM7;C9JHL4 | Negative elongation factor A | NELFA | 308830 | 235550 | 349480 | 45470 | 116140 | 0 |
| H0Y4S9;O00505 | Importin subunit alpha-4 | KPNA3 | 6969100 | 488030 | 982290 | 3682500 | 1854500 | 410420 |
| H0Y4T6;J3KMW3;H0Y8P6;H0Y610;Q9Y237;Q9Y237-3;Q9Y237-2 | Peptidyl-prolyl cis-trans isomerase NIMA-interacting 4 | PIN4 | 2133300 | 890450 | 2801400 | 905440 | 1987800 | 1171300 |
| H0Y5S9;H0Y682;P49674 | Casein kinase I isoform epsilon | CSNK1E | 2386200 | 867870 | 2122500 | 1408200 | 1809800 | 405180 |
| H0Y6V3;H0Y7V8;H0Y5V1;Q5VT25-3;Q5VT25-4;Q5VT25-5;Q5VT25;Q5VT25-2;Q5VT25-6;H0Y6R0 | Serine/threonine-protein kinase MRCK alpha | CDC42BPA | 197990 | 92153 | 120480 | 0 | 202660 | 77301 |
| H0Y650;Q86Y56-2;Q86Y56;E9PGY2 | Dynein assembly factor 5, axonemal | DNAAF5 | 187570 | 101160 | 203030 | 0 | 0 | 0 |
| H0Y6U5;Q86XI2;Q86XI2-2;F8WE06 | Condensin-2 complex subunit G2 | NCAPG2 | 856060 | 370050 | 864570 | 366650 | 506250 | 229050 |
| H0Y7A7;P62158;E7ETZ0;E7EMB3;Q96HY3;M0QZ52;G3V361;F8WBR5;G3V479;G3V226;P27482;C9J7T9;P02585 | Calmodulin | CALM2;CALM1;CALM3 | 78433000 | 50037000 | 79503000 | 10726000 | 14457000 | 8204100 |
| H0Y8E6;P49736 | DNA replication licensing factor MCM2 | MCM2 | 275300 | 186940 | 973150 | 95614 | 37201 | 27721 |
| H0YA96;H0Y8G5;Q14103-4;Q14103-3;Q14103-2;Q14103;D6RAF8;D6RF44;D6RD83;D6RBQ9 | Heterogeneous nuclear ribonucleoprotein D0 | HNRNPD | 56589000 | 26670000 | 80099000 | 33211000 | 64686000 | 26819000 |
| H0Y9J6;O95486-2;O95486 | Protein transport protein Sec24A | SEC24A | 1166700 | 0 | 0 | 0 | 0 | 0 |
| H0Y9Y2;Q8NFC6 | Biorientation of chromosomes in cell division protein 1-like 1 | BOD1L1 | 0 | 0 | 0 | 132080 | 0 | 0 |
| H0YE76;H0YCB3;Q8N9N2-2;Q8N9N2;H0YER2;E9PM82;E9PR40 | Activating signal cointegrator 1 complex subunit 1 | ASCC1 | 8817400 | 595750 | 10504000 | 1445700 | 1720100 | 1788400 |
| H0YCK3;P55265-5;P55265-3;P55265-2;P55265;P55265-4 | Double-stranded RNA-specific adenosine deaminase | ADAR | 43615 | 51177 | 296710 | 147180 | 337400 | 171320 |
| H0YCM7;Q5JVZ5;Q96JJ3-3;Q96JJ3;Q5JVZ8;Q5JW01;Q92556-2;Q92556-3;Q92556 | Engulfment and cell motility protein 2;Engulfment and cell motility protein 1 | ELMO2;ELMO1 | 285440 | 104360 | 0 | 210840 | 268340 | 27694 |
| H0YCN4;Q9BTE7;E9PQV9;E9PM04 | DCN1-like protein;DCN1-like protein 5 | DCUN1D5 | 2759200 | 1027600 | 2174900 | 288180 | 1172500 | 854530 |
| H0YDP6;H0YEL0;O14893-3;O14893-2;O14893;G5EA29 | Gem-associated protein 2 | GEMIN2 | 3229400 | 1656700 | 4206300 | 2986700 | 6683800 | 1865700 |
| H0YI37;H0YH81;P06576 | ATP synthase subunit beta;ATP synthase subunit beta, mitochondrial | ATP5B | 5069800 | 7931900 | 6258200 | 4374600 | 2491400 | 1854500 |
| H0YHF6;O94952-1;O94952;H0YIE9 | F-box only protein 21 | FBXO21 | 934080 | 1604400 | 3824500 | 917920 | 533310 | 97680 |
| H0YIZ0;P34897-3;P34897-2;P34897;G3V2Y4 | Serine hydroxymethyltransferase;Serine hydroxymethyltransferase, mitochondrial | SHMT2 | 661090 | 564490 | 1168200 | 1014600 | 1234800 | 967460 |
| H0YJ03 | | PSMA3 | 3024000 | 4143000 | 2655400 | 577350 | 1377000 | 740550 |
| H0YKD8;P46779;P46779-2;P46779-3;H0YLP6;H0YMF4;P46779-4;P46779-5 | 60S ribosomal protein L28 | RPL28 | 198130000 | 122250000 | 267150000 | 163840000 | 316620000 | 215710000 |
| H0YLW7;H0YMW8;H0YNF3;H0YMR6;H0YLE6;H0YL55;H7BXE3;Q9NWH9 | SAFB-like transcription modulator | SLTM | 499000 | 746690 | 1790400 | 1650200 | 3103400 | 1668500 |
| H0YLA2;P37108 | Signal recognition particle 14 kDa protein | SRP14 | 50387000 | 30098000 | 58281000 | 46469000 | 107830000 | 49564000 |
| H0YM23;O75179-6;O75179-7;O75179-2;O75179;H0YLQ3;O75179-4 | Ankyrin repeat domain-containing protein 17 | ANKRD17 | 607330 | 190740 | 463010 | 265980 | 418070 | 133260 |
| H0YM91;J3KP65;O15178-2;O15178 | Brachyury protein | T | 0 | 0 | 716210 | 0 | 0 | 0 |
| H0YMV8;Q71UM5;C9JLI6;C9J1C5 | 40S ribosomal protein S27;40S ribosomal protein S27-like | RPS27L | 4079800 | 3025100 | 9021900 | 704270 | 7995000 | 2716200 |
| H0YN26;P39687;O95626;O43423;H7BZ09;Q5T6W8 | Acidic leucine-rich nuclear phosphoprotein 32 family member A | ANP32A | 33091000 | 14230000 | 33502000 | 13549000 | 9848300 | 6879400 |
| H0YN81;Q9GZS3;H0YL19;H0YMF9;H0YM76;H3BQA8;H0YLA1 | WD repeat-containing protein 61;WD repeat-containing protein 61, N-terminally processed | WDR61 | 36846000 | 17858000 | 27003000 | 14592000 | 18801000 | 14066000 |
| H3BLZ8;Q92841-1;Q92841-3;Q92841;Q92841-2 | Probable ATP-dependent RNA helicase DDX17 | DDX17 | 15702000 | 11797000 | 29098000 | 15763000 | 30585000 | 14945000 |
| H3BM45;Q8N957 | Ankyrin repeat and fibronectin type-III domain-containing protein 1 | ANKFN1 | 751670 | 379460 | 412250 | 218580 | 182770 | 59792 |
| H3BMH2;H3BSC1;P62491-2;Q15907-2;P62491;Q15907;B4DQU5 | Ras-related protein Rab-11A;Ras-related protein Rab-11B | RAB11A;RAB11B | 8485600 | 5666000 | 7575300 | 1958400 | 1087100 | 861460 |
| H3BMZ1;H3BRL8;H3BPZ1;H3BS72;Q9P035-2;Q9P035 | Very-long-chain (3R)-3-hydroxyacyl-CoA dehydratase 3 | HACD3 | 294580 | 0 | 332720 | 0 | 0 | 0 |
| H3BP76;H3BUK7;H3BNF6;H3BRL2;H3BQK6;Q14764 | Major vault protein | MVP | 7086100 | 5564700 | 0 | 8008800 | 146170 | 1017300 |
| H3BSB3;H3BNT4;Q99547 | M-phase phosphoprotein 6 | MPHOSPH6 | 579310 | 366530 | 769440 | 8229300 | 11742000 | 5716700 |
| H3BP71;O75150-4;O75150;A0A087WTK2;O75150-3;REV__F5GXW3;REV__F5H4F8;REV__Q8TBY8-3;REV__Q8TBY8-2;REV__Q8TBY8;REV__G3V1Q7;REV__A6NE52 | E3 ubiquitin-protein ligase BRE1B | RNF40 | 1573500 | 760960 | 1210300 | 742680 | 1675900 | 708580 |
| H3BPE1;H3BQK9;Q9UPN3-3;Q9UPN3-2;Q9UPN3-4;Q9UPN3;H0Y390;Q9UPN3-5;E9PNZ4;E9PLY5;H0YD69;E9PLY0;H0Y314 | Microtubule-actin cross-linking factor 1, isoforms 1/2/3/5 | MACF1 | 381210 | 137000 | 343890 | 2893.8 | 14620 | 6504.6 |
| H3BPE7;P35637-2;P35637 | RNA-binding protein FUS | FUS | 21140000 | 17847000 | 24486000 | 35414000 | 57553000 | 30456000 |
| H3BPK3;H3BPQ4;Q16775-2;Q16775;H3BQW8;Q16775-3 | Hydroxyacylglutathione hydrolase, mitochondrial | HAGH | 902880 | 433370 | 890830 | 658320 | 643250 | 273620 |
| H3BSW3;H3BQF1;H3BQZ9;P07741-2;P07741 | Adenine phosphoribosyltransferase | APRT | 654360 | 0 | 264380 | 220080 | 198060 | 85480 |
| H3BRL3;O14562;H3BUM8 | Ubiquitin domain-containing protein UBFD1 | UBFD1 | 3212500 | 1768100 | 2959000 | 0 | 134120 | 58312 |
| H3BRS3;Q9Y4R8 | Telomere length regulation protein TEL2 homolog | TELO2 | 659000 | 372140 | 995290 | 139880 | 211320 | 58746 |
| H3BS42;Q9H5H4 | Zinc finger protein 768 | ZNF768 | 559650 | 287720 | 667100 | 86064 | 281410 | 41630 |
| H3BSM5;P60520;H3BQ50 | Gamma-aminobutyric acid receptor-associated protein-like 2 | GABARAPL2 | 3360800 | 324250 | 1288500 | 0 | 0 | 0 |
| H3BTN5;P14618-2;P14618;P14618-3;H3BT25;H3BUW1;H3BTJ2;H3BQ34;H3BR70;P30613-2;P30613;H3BN34;H3BQZ3;H3BU13 | Pyruvate kinase;Pyruvate kinase PKM | PKM | 1018000 | 605930 | 1283900 | 64601 | 527960 | 275140 |
| H3BUF6;Q8WWM7-6;Q8WWM7-8;Q8WWM7-5;Q8WWM7-4;Q8WWM7-9;Q8WWM7-2;Q8WWM7;Q8WWM7-3;H3BSK9;Q8WWM7-7 | Ataxin-2-like protein | ATXN2L | 154420 | 159330 | 694530 | 747330 | 2023600 | 679760 |
| H7BXY3;Q7L2E3-3;Q7L2E3;Q7L2E3-2 | Putative ATP-dependent RNA helicase DHX30 | DHX30 | 2609800 | 1591400 | 3402300 | 2401100 | 3534200 | 1848600 |
| H7BYY1;F5H7S3;B7Z596;P09493-2;H0YK48;P09493-5;P09493-8;P09493-7;H0YNC7;P09493-6;H0YL42;H0YK20;H0YN06 | Tropomyosin alpha-1 chain | TPM1 | 7309300 | 7216400 | 14140000 | 0 | 310900 | 56199 |
| H7BZE2;Q15654 | Thyroid receptor-interacting protein 6 | TRIP6 | 868740 | 226750 | 0 | 0 | 0 | 0 |
| H7BZU3;P31939-2;P31939 | Bifunctional purine biosynthesis protein PURH;Phosphoribosylaminoimidazolecarboxamide formyltransferase;IMP cyclohydrolase | ATIC | 1178000 | 522480 | 1303600 | 0 | 439420 | 0 |
| H7C2R6;H7BZW1;P57076;C9JX57;D3DSE6;C9J818;F8VZ95 | UPF0769 protein C21orf59 | C21orf59 | 1951900 | 1050500 | 2281100 | 390830 | 1410900 | 873700 |
| H7BZW2;Q6P2H3-3;Q6P2H3-4;Q6P2H3-2;Q6P2H3 | Centrosomal protein of 85 kDa | CEP85 | 193150 | 232750 | 234940 | 0 | 0 | 0 |
| H7C2B1;Q9UMY1-2;Q9UMY1 | Nucleolar protein 7 | NOL7 | 0 | 1169400 | 855050 | 3107700 | 3252400 | 2775600 |
| H7C2Q8;Q99848 | Probable rRNA-processing protein EBP2 | EBNA1BP2 | 4049700 | 2336400 | 5659600 | 14349000 | 25758000 | 14636000 |

| | | | | | | | | |
|---|---|---|---|---|---|---|---|---|
| H7C3P9;Q9UNS2-2;Q9UNS2;K7ES36;C9JLV5 | COP9 signalosome complex subunit 3 | COPS3 | 1414400 | 741700 | 1440600 | 614880 | 844730 | 363470 |
| H7C3T4;Q13162 | Peroxiredoxin-4 | PRDX4 | 244810 | 313690 | 1223900 | 80622 | 395690 | 259490 |
| H7C492;Q9Y276 | Mitochondrial chaperone BCS1 | BCS1L | 0 | 212170 | 634790 | 0 | 0 | 0 |
| H8Y6P7;P0CAP2-2;P0CAP2-3;P0CAP2 | DNA-directed RNA polymerase II subunit GRINL1A | GCOM1;POLR2M | 88984 | 88621 | 100870 | 106150 | 49837 | 38929 |
| I3L0N3;P46459-2;P46459 | Vesicle-fusing ATPase | NSF | 783120 | 90208 | 533130 | 0 | 0 | 0 |
| I3L1P5;O14964-2;O14964 | Hepatocyte growth factor-regulated tyrosine kinase substrate | HGS | 0 | 0 | 0 | 0 | 0 | 0 |
| I3L2B0;I3L318;K7EIG1;O75153 | Clustered mitochondria protein homolog | CLUH | 140140 | 73995 | 462490 | 0 | 87642 | 33794 |
| I3L2C7;P57678;E7EN12;I3L4M4;I3L4A4;I3L399 | Gem-associated protein 4 | GEMIN4 | 866980 | 480870 | 1100600 | 817380 | 1717000 | 734870 |
| I3L397;I3L504;P63241;P63241-2;F8WCJ1;C9J7B5;C9J4W5;Q9GZV4;Q6IS14 | Eukaryotic translation initiation factor 5A;Eukaryotic translation initiation factor 5A-1 | EIF5A | 107110000 | 50340000 | 149450000 | 19273000 | 34337000 | 14951000 |
| I3L3P7;P62244;H3BVC7;I3L246;H3BV27;H3BT37;I3L303 | 40S ribosomal protein S15a | RPS15A | 178370000 | 130010000 | 265430000 | 156720000 | 302270000 | 122790000 |
| I3L4J1;Q9UN37 | Vacuolar protein sorting-associated protein 4A | VPS4A | 1654100 | 230450 | 416670 | 0 | 0 | 105680 |
| J3KN10;P42356;P42356-2 | Phosphatidylinositol 4-kinase alpha | PI4KA | 88625 | 41138 | 94419 | 0 | 0 | 0 |
| J3KN67 | | TPM3 | 6324600 | 403570 | 5370900 | 0 | 0 | 0 |
| J3KNL6;O15027-2;O15027-4;O15027-3;O15027;O15027-5;Q8N9G1;X6RGP5 | Protein transport protein Sec16A | SEC16A | 643450 | 346900 | 554440 | 128320 | 178460 | 88513 |
| J3KNS1;Q9UPW5-2;Q9UPW5;Q9UPW5-3;Q5VZW3;J3KQF5;Q96MI9-2;Q96MI9-3;Q96MI9 | Cytosolic carboxypeptidase 1;Cytosolic carboxypeptidase 4 | AGTPBP1;AGBL1 | 63486 | 0 | 143620 | 37477 | 156480 | 35615 |
| Q5SQL3;J3KP92;Q86Z02-4;Q86Z02-3;Q86Z02-2;Q86Z02 | Homeodomain-interacting protein kinase 1 | HIPK1 | 4755200 | 1043900 | 3021000 | 594700 | 99688 | 300200 |
| J3KPS3;P04075;H3BQN4;P04075-2;H3BPS8;H3BUH7;H3BMQ8;H3BR04;H3BR68;H3BU78;Q5T7D5;A0A087WXX2;P05062 | Fructose-bisphosphate aldolase;Fructose-bisphosphate aldolase A | ALDOA | 43717000 | 17187000 | 45767000 | 18706000 | 21429000 | 14885000 |
| J3KQ32;Q9NTK5;Q9NTK5-2;Q9NTK5-3;C9JTK6;C9JCJ9 | Obg-like ATPase 1 | OLA1 | 8770100 | 2495800 | 9557800 | 11042000 | 17456000 | 7401900 |
| J3KQ34;J3KQ41;Q9H9Q2;Q9H9Q2-3 | COP9 signalosome complex subunit 7b | COPS7B | 755930 | 0 | 229500 | 0 | 0 | 0 |
| J3KR24;P41252;Q5TCC4;Q5TCC5;Q5TCD2;Q5TCD1;Q5TCC6 | Isoleucine--tRNA ligase, cytoplasmic | IARS | 6252800 | 3879500 | 8549400 | 4583500 | 9937400 | 4135700 |
| J3KRA9;Q96Q15-2;Q96Q15;Q96Q15-4;Q96Q15-3;A0A087WXM7;E9PNP6;REV__P51888;I3L400;I3L0W2;H0Y304;A0A087WV90;E9PDN5;A0A087WTU7;A0A075B6G3;P11532-3;P11532-2;P11532-4;P11532 | Serine/threonine-protein kinase SMG1 | SMG1 | 37950 | 20702 | 173920 | 35364 | 170540 | 36021 |
| J3QR85;J3KSR7;Q9Y5J1 | U3 small nucleolar RNA-associated protein 18 homolog | UTP18 | 0 | 227480 | 567810 | 1060400 | 1760900 | 0 |
| J3KSW6;Q8IX18-3;Q8IX18;J3KSX9;J3KS08;J3KTK0;Q8IX18-4;Q8IX18-2 | Probable ATP-dependent RNA helicase DHX40 | DHX40 | 1078600 | 719280 | 1262700 | 933440 | 515460 | 533570 |
| J3KTA4;P17844-2;P17844;J3QRQ7;J3QR02;J3KRX8;J3QSF1;J3KRZ1;X6RLV5;J3QKN9 | Probable ATP-dependent RNA helicase DDX5 | DDX5 | 5375300 | 5129200 | 8688200 | 8275000 | 14070000 | 7410500 |
| J3QR09;J3KTE4;P84098;J3QL15 | Ribosomal protein L19;60S ribosomal protein L19 | RPL19 | 281180000 | 152550000 | 347870000 | 262280000 | 491970000 | 267590000 |
| J3KTI3;O95544-3;O95544;O95544-2 | NAD kinase | NADK | 0 | 0 | 0 | 0 | 0 | 0 |
| J3KTL2;Q07955-3;Q07955;Q07955-2;J3QQV5;J3KSW7;J3KSR8 | Serine/arginine-rich splicing factor 1 | SRSF1 | 311790 | 74427 | 116760 | 43093 | 214750 | 110220 |
| J3QLI9;P62314 | Small nuclear ribonucleoprotein Sm D1 | SNRPD1 | 1190100 | 2198600 | 2586200 | 2362400 | 0 | 0 |
| J3QQW9;Q15022 | Polycomb protein SUZ12 | SUZ12 | 138690 | 114510 | 230250 | 42139 | 0 | 0 |
| J3QRS3;P19105;O14950;P24844;J3KTJ1;P24844-2 | Myosin regulatory light chain 12A;Myosin regulatory light chain 12B;Myosin regulatory light polypeptide 9 | MYL12A;MYL12B;MYL9 | 2555300 | 850850 | 1526200 | 0 | 0 | 0 |
| J3QRS9;X6R4W8;O43670-2;O43670;O43670-4;O43670-3;J3KS31;J3QS27;J3KTL1;J3KRB6;J3KRW6;H0Y3M2 | BUB3-interacting and GLEBS motif-containing protein ZNF207 | ZNF207 | 2644900 | 2259400 | 1601500 | 1697400 | 2046200 | 601100 |
| J3QT28;O43684-2;O43684;J3QSX4 | Mitotic checkpoint protein BUB3 | BUB3 | 19847000 | 10097000 | 11921000 | 6385600 | 4800700 | 3645600 |
| K4DI93;Q13620-3;Q13620-1;Q13620;A6NE76;Q13619-2;Q13619 | Cullin-4B | CUL4B | 93371 | 0 | 0 | 0 | 51732 | 0 |
| K7EM02;K7EIJ8;Q8IYT4-2;Q8IYT4 | Katanin p60 ATPase-containing subunit A-like 2 | KATNAL2 | 6647200 | 6096600 | 6895100 | 0 | 0 | 0 |
| K7EIP6;P51148;P51148-2 | Ras-related protein Rab-5C | RAB5C | 7774800 | 4816100 | 6358900 | 1970200 | 1201200 | 682520 |
| K7EJ19;K7EIU7;K7EPK2;O43776-2;O43776;K7EMQ6;K7EQ35 | Asparagine--tRNA ligase, cytoplasmic | NARS | 881830 | 466040 | 1864600 | 1411700 | 2383400 | 987280 |
| K7EIY6;Q9BV68-2;Q9BV68;U3KQF4;U3KPY8 | E3 ubiquitin-protein ligase RNF126 | RNF126 | 3195600 | 1253400 | 2660600 | 0 | 687750 | 407060 |
| K7EJY5;Q6P1N0-2;Q6P1N0 | Coiled-coil and C2 domain-containing protein 1A | CC2D1A | 328410 | 151730 | 279940 | 208990 | 259540 | 178040 |
| K7EL20;O75821;K7ER90;K7ENA8;K7EP16;K7EL60;K7ENH0 | Eukaryotic translation initiation factor 3 subunit G | EIF3G | 29252000 | 17407000 | 35121000 | 26831000 | 48879000 | 27756000 |
| K7ELC2;S4R456;K7EM56;K7EJ78;S4R417;K7EQJ5;P62841;REV__B4DHD0 | 40S ribosomal protein S15 | RPS15 | 70572000 | 23343000 | 67149000 | 50684000 | 96085000 | 9974900 |
| K7ELG9;Q3MHD2;Q3MHD2-2 | Protein LSM12 homolog | LSM12 | 0 | 0 | 0 | 0 | 620920 | 304570 |
| K7ELL7;P14314-2;P14314 | Glucosidase 2 subunit beta | PRKCSH | 639620 | 92642 | 465970 | 88667 | 267480 | 101560 |
| K7EM13;P10644-2;P10644 | cAMP-dependent protein kinase type I-alpha regulatory subunit;cAMP-dependent protein kinase type I-alpha regulatory subunit, N-terminally processed | PRKAR1A | 767860 | 0 | 253830 | 0 | 160500 | 0 |
| K7EM18;P41567 | Eukaryotic translation initiation factor 1 | EIF1 | 10929000 | 5991500 | 9577900 | 5464500 | 12734000 | 4181000 |
| K7ENR6;Q969U7-2;Q969U7;V9GZ55;K7ER45 | Proteasome assembly chaperone 2 | PSMG2 | 183610000 | 71328000 | 98543000 | 77715000 | 52484000 | 35516000 |
| K7ERF1;Q9UBQ5;K7ES31;K7EMQ9;Q9UBQ5-2;A0A087WVB9;K7EK53;K7EQM4 | Eukaryotic translation initiation factor 3 subunit K | EIF3K | 9778300 | 8680500 | 16304000 | 5585400 | 14576000 | 9495300 |
| K7ERI7;K7EMH1;P35268;K7EJT5;K7EP65;K7EKS7;K7ELC4 | 60S ribosomal protein L22 | RPL22 | 317920000 | 245760000 | 534280000 | 273470000 | 657510000 | 350970000 |
| Q3HM38;X6R2P6;K7ESJ4;O14737;O14737-2 | Programmed cell death protein 5 | PDCD5 | 2388500 | 2031100 | 3160900 | 0 | 578300 | 415200 |
| K7ESL1;P83876 | Thioredoxin-like protein 4A | TXNL4A | 0 | 0 | 0 | 942470 | 0 | 0 |
| M0R0P1;M0R299;M0QXL5;M0R2Q4;P22087;M0R2U2;M0R1H0;M0R2B0 | rRNA 2-O-methyltransferase fibrillarin | FBL | 708690 | 1080100 | 3598100 | 1174900 | 4010000 | 2243800 |
| M0QXS5;P14866;P14866-2;M0QYT0;M0R1W6;M0R076;B4DVF8;M0QYL7 | Heterogeneous nuclear ribonucleoprotein L | HNRNPL | 18610000 | 9912500 | 17986000 | 14591000 | 16377000 | 9181000 |
| M0QXZ5;Q96B54 | Zinc finger protein 428 | ZNF428 | 5720400 | 1315000 | 2470300 | 922900 | 1169300 | 754430 |
| M0R2P8;M0R3A4;M0QYA2;Q9NWS0 | PIH1 domain-containing protein 1 | PIH1D1 | 324270 | 553930 | 1336000 | 58775 | 442580 | 230390 |
| M0QZC5;P62280;M0R1H6;M0R1H5 | 40S ribosomal protein S11 | RPS11 | 447120000 | 223250000 | 551300000 | 375980000 | 677210000 | 318540000 |
| M0QZI4;Q8IV08 | Phospholipase D3 | PLD3 | 637100 | 510190 | 994730 | 0 | 0 | 0 |
| M0R0F0;P46782;M0R0R2;M0QZN2 | 40S ribosomal protein S5;40S ribosomal protein S5, N-terminally processed | RPS5 | 421340000 | 230380000 | 459340000 | 356080000 | 620770000 | 239300000 |
| M0R0Y2;M0R2M1;P54920 | Alpha-soluble NSF attachment protein | NAPA | 0 | 84553 | 133320 | 0 | 0 | 0 |
| M0R0Z3;P35269 | General transcription factor IIF subunit 1 | GTF2F1 | 192750 | 161060 | 389470 | 0 | 190210 | 0 |
| M0R117;Q02543;M0R3D6;M0R1A7;M0R0P7 | 60S ribosomal protein L18a | RPL18A | 157840000 | 76884000 | 152900000 | 127950000 | 184350000 | 104180000 |
| M0R2S2;M0R165;Q9UBC2-4;Q9UBC2-3;Q9UBC2;Q9UBC2-2 | Epidermal growth factor receptor substrate 15-like 1 | EPS15L1 | 955080 | 404830 | 1267600 | 344010 | 367580 | 0 |
| M0R210;P62249;M0R3H0;M0R1M5;A0A087WZ27;M0QX76;Q6IPX4 | 40S ribosomal protein S16 | RPS16;ZNF90 | 337360000 | 103670000 | 219590000 | 254230000 | 402340000 | 214400000 |
| M0R2B7;P28340;M0R2I8;M0QXE6;M0QXQ2;M0R3H8;M0QZR8 | DNA polymerase;DNA polymerase delta catalytic subunit | POLD1 | 68781 | 3630600 | 0 | 8767900 | 2516900 | 0 |
| M0R2N4;S4R353;K7EJH3;S4R443;K7EJM7;M0QYK5;S4R3L8;Q01081-3;Q8WU68-3;Q8WU68-2;Q8WU68;Q01081-2;Q01081 | Splicing factor U2AF 35 kDa subunit;Splicing factor U2AF 26 kDa subunit | U2AF1L4;U2AF1 | 0 | 10861000 | 24689000 | 61396000 | 52074000 | 53266000 |
| M0R2S1;M0R1M6 | | UBA52 | 0 | 252700 | 164000 | 0 | 0 | 0 |
| M0R3F6;M0R2Z9;Q8IX01-4;Q8IX01-3;Q8IX01 | SURP and G-patch domain-containing protein 2 | SUGP2 | 0 | 0 | 79421 | 0 | 0 | 0 |
| M0R366;Q9BTV5;M0R1R0 | Fibronectin type III and SPRY domain-containing protein 1 | FSD1 | 1877600 | 761640 | 2941400 | 1531100 | 2161300 | 723180 |
| O00139;O00139-4;O00139-2;O00139-1;O00139-5;D6R9M0;H0Y5Z9;H0Y8H2 | Kinesin-like protein KIF2A | KIF2A | 6594200 | 4102400 | 6060200 | 4665700 | 6107100 | 3691300 |
| O00151 | PDZ and LIM domain protein 1 | PDLIM1 | 22600000 | 3781000 | 7717200 | 0 | 0 | 0 |
| O00178;F5H716;F5H257;B0QY59 | GTP-binding protein 1 | GTPBP1 | 888780 | 722520 | 850520 | 1223300 | 2110800 | 1249200 |
| O00203-3;O00203;H0YBD0 | AP-3 complex subunit beta-1 | AP3B1 | 43530 | 0 | 0 | 0 | 0 | 0 |
| O00231;O00231-2;J3QRY4;J3QS13;J3KSW3 | 26S proteasome non-ATPase regulatory subunit 11 | PSMD11 | 654400000 | 354930000 | 420390000 | 58463000 | 51939000 | 33599000 |
| O00232;O00232-2;J3KTJ5;J3KSK1;REV__A8MT70-3;REV__A8MT70;REV__A8MT70-2;P0C617 | 26S proteasome non-ATPase regulatory subunit 12 | PSMD12 | 431490000 | 253280000 | 283060000 | 52146000 | 49391000 | 27153000 |
| O00267-2;O00267 | Transcription elongation factor SPT5 | SUPT5H | 3194900 | 1485400 | 3164500 | 970930 | 1231000 | 334270 |
| O00291-3;V5LU97;O00291;C9JMG5 | Huntingtin-interacting protein 1 | HIP1 | 1588300 | 448900 | 1335000 | 0 | 34021 | 153400 |
| O00303;H0YDT6 | Eukaryotic translation initiation factor 3 subunit F | EIF3F | 23117000 | 13110000 | 28346000 | 34322000 | 50642000 | 15930000 |
| O00410;O00410-3;H0Y8C6;O00410-2;E7EQT5;E7EV12;E7ETV3;C9JXE0;C9JZD8;C9J875;C9JZ53;C9JQT6;E7ESA1;E7ESZ1;E7EWK4;E7EX05;E7ETV8;C9JMV5;H0Y3V4 | Importin-5 | IPO5 | 1235700 | 422560 | 1005600 | 1116100 | 1400100 | 567960 |
| O00425;O00425-2;F8WD15 | Insulin-like growth factor 2 mRNA-binding protein 3 | IGF2BP3 | 6439500 | 4146200 | 12843000 | 10639000 | 22498000 | 10687000 |
| O00442;O00442-2 | RNA 3-terminal phosphate cyclase | RTCA | 276120 | 0 | 104690 | 0 | 66098 | 0 |
| O00488 | Zinc finger protein 593 | ZNF593 | 2078200 | 2945800 | 3624400 | 0 | 9046500 | 3474900 |
| O00567;H0Y653;H0YDU4;Q5JXT2 | Nucleolar protein 56 | NOP56 | 331130 | 326410 | 1346400 | 833620 | 2297700 | 1103000 |
| O00571-2;O00571;O15523-2;O15523;A0A087WVZ1;C9J8G5;C9J081;O15523-3;B4DLA0;D6RCM4;Q9NQI0-3;Q9NQI0-2;Q9NQI0-4;Q9NQI0 | ATP-dependent RNA helicase DDX3X;ATP-dependent RNA helicase DDX3Y | DDX3X;DDX3Y | 5795900 | 3490400 | 7341400 | 2552100 | 6531700 | 2408500 |
| O14654 | Insulin receptor substrate 4 | IRS4 | 1811900 | 1340800 | 2658100 | 421250 | 557760 | 328910 |
| O14744-5;O14744;G3V580;C9JSX3;H0YJX6;H0YJY6;H0YJ77;G3V5L5;G3V2L6;G3V2X6;G3V507;G3V2F5;G3V5T6;H7BZ44 | Protein arginine N-methyltransferase 5;Protein arginine N-methyltransferase 5, N-terminally processed | PRMT5 | 3699500 | 1725700 | 2785000 | 2063000 | 3085500 | 1412300 |
| O14776-2;O14776;G3V220 | Transcription elongation regulator 1 | TCERG1 | 4066000 | 877980 | 1745300 | 1847400 | 2138000 | 826430 |
| O14818;H0Y586;O14818-4;A0A087WYS6;Q8TAA3-2;Q8TAA3-5;Q8TAA3 | Proteasome subunit alpha type-7 | PSMA7 | 1581600000 | 1345300000 | 1408400000 | 968650000 | 927180000 | 779440000 |
| O14818-2 | Proteasome subunit alpha type-7 | PSMA7 | 39169000 | 4590000 | 43573000 | 10622000 | 15234000 | 16319000 |
| O14828-2;O14828 | Secretory carrier-associated membrane protein 3 | SCAMP3 | 673720 | 190980 | 2048700 | 0 | 0 | 0 |
| O14974-4;O14974;F8VZN8;O14974-5;O14974-3;O14974-2;H0YIS4;H0YHI8;H0YHL8;H0YIM2;H0YIS3 | Protein phosphatase 1 regulatory subunit 12A | PPP1R12A | 1539700 | 904060 | 697630 | 0 | 130850 | 28894 |
| O15020-2;O15020;A4QPE4 | Spectrin beta chain, non-erythrocytic 2 | SPTBN2 | 1771800 | 301550 | 1406500 | 0 | 34796 | 0 |
| O15144;G5E9S7;C9JTV5;G5E9J0 | Actin-related protein 2/3 complex subunit 2 | ARPC2 | 1012800 | 243740 | 451700 | 0 | 0 | 0 |
| O15145;F8VR50;C9JZD1 | Actin-related protein 2/3 complex subunit 3 | ARPC3 | 6009400 | 2086200 | 2180600 | 0 | 0 | 0 |
| O15160-2;O15160;D6RDJ3;E7EQB9 | DNA-directed RNA polymerases I and III subunit RPAC1 | POLR1C | 969170 | 584240 | 1117200 | 747020 | 1353100 | 713900 |
| O15226;O15226-2;A0A087WXT2 | NF-kappa-B-repressing factor | NKRF | 2122000 | 1200500 | 3733900 | 5255300 | 12064000 | 4724300 |
| O15294;O15294-3;O15294-2;O15294-4;C9JZL3;H7C2T7 | UDP-N-acetylglucosamine--peptide N-acetylglucosaminyltransferase 110 kDa subunit | OGT | 1859900 | 832870 | 1031700 | 408360 | 331210 | 231950 |
| O15344;O15344-2;C9J453;A0A087X255;A0A087X0X0;C9JZJ7 | E3 ubiquitin-protein ligase Midline-1 | MID1 | 1030500 | 667470 | 668880 | 672520 | 1241000 | 405650 |
| O15355 | Protein phosphatase 1G | PPM1G | 3926200 | 1204900 | 4817800 | 390360 | 472460 | 365760 |
| O15371-2;O15371-3;O15371;B0QYA5;B0QYA6;B0QYA8;B0QYA7;B0QYA4 | Eukaryotic translation initiation factor 3 subunit D | EIF3D | 20436000 | 11133000 | 25526000 | 16860000 | 36431000 | 19789000 |
| O15381-3;O15381-4;O15381-2;O15381-5;O15381 | Nuclear valosin-containing protein-like | NVL | 332290 | 123850 | 446800 | 0 | 87649 | 0 |
| O15400-2;O15400 | Syntaxin-7 | STX7 | 2129900 | 436010 | 1412500 | 0 | 38923 | 0 |
| O94900;O15405-2;O15405;O94842-3;O94842-2;O94842 | Thymocyte selection-associated high mobility group box protein TOX;TOX high mobility group box family member 3;TOX high mobility group box family member 4 | TOX;TOX3;TOX4 | 226320 | 299980 | 215580 | 431650 | 196730 | 315720 |
| O15446;O15446-2 | DNA-directed RNA polymerase I subunit RPA34 | CD3EAP | 377150 | 355520 | 736700 | 67153 | 538510 | 94159 |
| O43143 | Pre-mRNA-splicing factor ATP-dependent RNA helicase DHX15 | DHX15 | 13718000 | 6456700 | 9663700 | 2305100 | 3152100 | 1518300 |
| O43148;O43148-2;K7EP06;K7EPP5;K7ERH6 | mRNA cap guanine-N7 methyltransferase | RNMT | 1387500 | 482910 | 633290 | 1497800 | 846270 | 427830 |
| O43172-2;O43172 | U4/U6 small nuclear ribonucleoprotein Prp4 | PRPF4 | 901230 | 438030 | 903350 | 1237100 | 580110 | 287770 |
| O43175;Q5SZU1 | D-3-phosphoglycerate dehydrogenase | PHGDH | 7297300 | 2696700 | 8327300 | 624760 | 1974800 | 907250 |
| O43242;H0YGV8;F5H8K4 | 26S proteasome non-ATPase regulatory subunit 3 | PSMD3 | 506070000 | 361430000 | 455120000 | 54688000 | 44592000 | 37471000 |
| O43242-2 | 26S proteasome non-ATPase regulatory subunit 3 | PSMD3 | 2206700 | 5229200 | 7162800 | 374870 | 488020 | 472770 |
| O43324;D6RBD7;C9J1V9;O43324-2;H0YAL7;H0YAH9;D6RCQ0 | Eukaryotic translation elongation factor 1 epsilon-1 | EEF1E1;EEF1E1-BLOC1S5 | 3680900 | 4971600 | 7799700 | 4784800 | 7634200 | 5383000 |
| O43390;O43390-3;O43390-2;B4DT28;O43390-4 | Heterogeneous nuclear ribonucleoprotein R | HNRNPR | 9392900 | 7059300 | 20070000 | 13615000 | 30015000 | 12467000 |
| O43395;O43395-3 | U4/U6 small nuclear ribonucleoprotein Prp3 | PRPF3 | 1632200 | 1252000 | 2893200 | 1973400 | 2311100 | 1153300 |
| O43396;K7ER96;K7EML9;K7EPB7;K7EKG2;K7EME7 | Thioredoxin-like protein 1 | TXNL1 | 107550000 | 59373000 | 72108000 | 12323000 | 13131000 | 7270200 |
| O43432;O43432-3;O43432-4;Q5SWD0 | Eukaryotic translation initiation factor 4 gamma 3 | EIF4G3 | 125090 | 81721 | 372990 | 305070 | 749840 | 430200 |
| O43660-2;O43660;H0YA24 | Pleiotropic regulator 1 | PLRG1 | 778550 | 82066 | 493490 | 115340 | 464450 | 235470 |
| O43707;O43707-2;F5GXS2;O43707-3;H7C144;K7EJH8;K7EP19;D6RH00 | Alpha-actinin-4 | ACTN4 | 232990000 | 83591000 | 142720000 | 1047600 | 889780 | 196350 |
| O43709;O43709-3;H7C0G4 | Probable 18S rRNA (guanine-N(7))-methyltransferase | WBSCR22 | 822340 | 376830 | 1743800 | 554950 | 2277500 | 258910 |
| O43719;Q5H919;Q5H918 | HIV Tat-specific factor 1 | HTATSF1 | 1743700 | 792420 | 1808300 | 500590 | 638140 | 263930 |
| O43765;K7EMD6 | Small glutamine-rich tetratricopeptide repeat-containing protein alpha | SGTA | 3820100 | 717820 | 1803000 | 0 | 0 | 0 |
| O43809 | Cleavage and polyadenylation specificity factor subunit 5 | NUDT21 | 8769500 | 4424600 | 6494300 | 772590 | 467840 | 470140 |
| O43818 | U3 small nucleolar RNA-interacting protein 2 | RRP9 | 144550 | 0 | 383780 | 1335800 | 2092200 | 825660 |
| O43865-2;O43865;H0Y8B3;Q96HN2-4;Q96HN2-3;Q96HN2-2;Q96HN2 | Putative adenosylhomocysteinase 2;Adenosylhomocysteinase;Putative adenosylhomocysteinase 3 | AHCYL1;AHCYL2 | 1456900 | 697880 | 1189100 | 46697 | 196030 | 0 |
| O43929-2;O43929-3;O43929;CON__Q2YDI2 | Origin recognition complex subunit 4 | ORC4 | 84145 | 120850 | 181020 | 0 | 0 | 0 |
| O60216 | Double-strand-break repair protein rad21 homolog | RAD21 | 199890 | 121870 | 423840 | 117150 | 0 | 121000 |
| O60231;Q5SQH5;A2AB15 | Putative pre-mRNA-splicing factor ATP-dependent RNA helicase DHX16 | DHX16 | 1163300 | 853200 | 1076800 | 116120 | 171800 | 77212 |
| O60264;P28370-2;P28370 | SWI/SNF-related matrix-associated actin-dependent regulator of chromatin subfamily A member 5 | SMARCA5 | 514970 | 150070 | 744400 | 74015 | 144680 | 53387 |
| O60306 | Intron-binding protein aquarius | AQR | 61724 | 56776 | 102880 | 51442 | 162240 | 101960 |
| O60506-4;O60506-3;O60506-2;O60506;B7Z645;O60506-5;F6UXX1;H0YAA0 | Heterogeneous nuclear ribonucleoprotein Q | SYNCRIP | 17655000 | 7707400 | 20105000 | 20620000 | 17865000 | 9075600 |
| O60524-4;O60524-3;O60524;G3V5V3;O60524-5;G3V4D9;G3V547;O60524-2 | Nuclear export mediator factor NEMF | NEMF | 2588600 | 1391900 | 3162100 | 1882700 | 3602500 | 2054600 |
| O60568;H7C2V1;H7C2S8;H7C0B8 | Procollagen-lysine,2-oxoglutarate 5-dioxygenase 3 | PLOD3 | 258680 | 167580 | 232500 | 0 | 0 | 0 |
| S4R3E5;O60684;Q5TFJ7 | Importin subunit alpha-7 | KPNA6 | 824220 | 1047600 | 886480 | 0 | 324860 | 0 |
| O60739 | Eukaryotic translation initiation factor 1b | EIF1B | 12310000 | 0 | 4453300 | 0 | 2231000 | 0 |
| O60749-2;O60749;D6RC15;H0YK42;Q13596-2;Q13596;Q13596-3 | Sorting nexin-2 | SNX2 | 651510 | 555950 | 692040 | 27285 | 67112 | 0 |
| O60832;O60832-2;C9IYT0;H7C2Q2;H7C0M1 | H/ACA ribonucleoprotein complex subunit 4 | DKC1 | 1357300 | 1159100 | 3152500 | 1711600 | 3712400 | 1374500 |
| O60869-2;O60869-3;O60869 | Endothelial differentiation-related factor 1 | EDF1 | 13128000 | 14017000 | 29303000 | 10862000 | 21343000 | 12017000 |
| S4R357;O60870-2;O60870 | DNA/RNA-binding protein KIN17 | KIN | 0 | 288660 | 0 | 444260 | 1060700 | 300740 |
| O60884;I3L320;A0A087WT48 | DnaJ homolog subfamily A member 2 | DNAJA2 | 10961000 | 3897400 | 9237500 | 2203900 | 1993000 | 839230 |
| Q5TCW7;O60942-3;O60942-4;O60942-2;O60942 | mRNA-capping enzyme;Polynucleotide 5-triphosphatase;mRNA guanylyltransferase | RNGTT | 712020 | 285160 | 468750 | 84196 | 202670 | 285840 |
| O75083;D6RD66;O75083-3 | WD repeat-containing protein 1 | WDR1 | 15942000 | 5908300 | 8775000 | 210020 | 777260 | 2545300 |
| O75131;A0A087WYQ3;A0A087WXR6;E5RHZ0;H0YB26;A0A087WUS8;E5RFT7;E7ENV7;Q86YQ8-2;Q9HCH3-2;Q96FN4-2;Q96FN4;Q8IYJ1;Q96A23;O95741;Q9UBL6-2;Q86YQ8;Q96A23-2;Q9HCH3;O95741-2;Q9UBL6 | Copine-3 | CPNE3 | 14568000 | 4594200 | 10663000 | 0 | 326640 | 167960 |
| O75152;E9PQ61;E9PBY7 | Zinc finger CCCH domain-containing protein 11A | ZC3H11A | 0 | 0 | 20473 | 0 | 207480 | 83681 |
| O75323;O75323-2;C9J7B1;C9K068;H7C333;F8WBI5 | Protein NipSnap homolog 2 | GBAS | 1827900 | 559980 | 1727200 | 0 | 607930 | 192620 |
| O75368 | SH3 domain-binding glutamic acid-rich-like protein | SH3BGRL | 2502800 | 546300 | 1309400 | 0 | 0 | 0 |
| O75369-6;O75369-3;O75369-2;O75369-9;O75369;O75369-8;E7EN95;O75369-5;O75369-4;O75369-7;H7C5L4 | Filamin-B | FLNB | 5701400 | 2114800 | 2910300 | 173480 | 231610 | 181670 |

| | | | | | | | | |
|---|---|---|---|---|---|---|---|---|
| O75436;S4R3Q6;O75436-2;Q4G0F5 | Vacuolar protein sorting-associated protein 26A | VPS26A | 4899800 | 2304100 | 3059300 | 283820 | 493940 | 202020 |
| O75531 | Barrier-to-autointegration factor;Barrier-to-autointegration factor, N-terminally processed | BANF1 | 3458100 | 1289100 | 10296000 | 453820 | 7402800 | 0 |
| O75533;H7C341;F8WC19;B4DGZ4;O75533-2 | Splicing factor 3B subunit 1 | SF3B1 | 1158600 | 776770 | 953960 | 1113400 | 1214200 | 703600 |
| O75534-2;O75534;O75534-3;O75534-4;E9PLT0;E9PNG3;E9PJK7 | Cold shock domain-containing protein E1 | CSDE1 | 3828300 | 1383200 | 5681500 | 635920 | 1172100 | 303560 |
| O75569;G5E9Q4;O75569-3;O75569-2;F8WEG8;C9JMM3 | Interferon-inducible double-stranded RNA-dependent protein kinase activator A | PRKRA | 2720900 | 1704100 | 2875200 | 2242300 | 6946200 | 2467000 |
| O75592-2;O75592;H7C3U4 | E3 ubiquitin-protein ligase MYCBP2 | MYCBP2 | 0 | 0 | 5417.9 | 29885 | 7420.7 | 0 |
| O75607 | Nucleoplasmin-3 | NPM3 | 675800 | 5477100 | 4551900 | 0 | 2814900 | 5082300 |
| O75643;O75643-2;B4E0P5;Q86VH2-3;Q86VH2-2;Q86VH2 | U5 small nuclear ribonucleoprotein 200 kDa helicase | SNRNP200 | 1740300 | 1265800 | 2477200 | 1135000 | 1144000 | 784440 |
| O75683 | Surfeit locus protein 6 | SURF6 | 386000 | 410430 | 1063000 | 2525100 | 3904000 | 2066300 |
| O75691 | Small subunit processome component 20 homolog | UTP20 | 0 | 0 | 0 | 0 | 70159 | 32857 |
| X6RKY7;X6RF82;X6RA30;O75794 | Cell division cycle protein 123 homolog | CDC123 | 972850 | 546740 | 1334700 | 0 | 0 | 0 |
| O75822;O75822-2;O75822-3;H0YLP3;H0YGJ7 | Eukaryotic translation initiation factor 3 subunit J | EIF3J | 29960000 | 19515000 | 32541000 | 46726000 | 61317000 | 36324000 |
| O75934 | Pre-mRNA-splicing factor SPF27 | BCAS2 | 496340 | 274220 | 299700 | 1438000 | 172400 | 387700 |
| X6RA56;X6RLR1;O75935-3;O75935-2;O75935;X6RCK5 | Dynactin subunit 3 | DCTN3 | 4694800 | 2352500 | 2893000 | 1178400 | 1269600 | 1037200 |
| O75937 | DnaJ homolog subfamily C member 8 | DNAJC8 | 1694500 | 2377200 | 713240 | 3090300 | 2860600 | 748580 |
| O75940 | Survival of motor neuron-related-splicing factor 30 | SMNDC1 | 525740 | 211580 | 294180 | 0 | 198270 | 110450 |
| O75976 | Carboxypeptidase D | CPD | 60741 | 24320 | 75128 | 0 | 0 | 0 |
| O76003 | Glutaredoxin-3 | GLRX3 | 1882600 | 1282200 | 1727500 | 648980 | 918190 | 512380 |
| O76021;J3QSV6;O76021-2;I3L3C4;I3L3U9;I3L234;I3L1A3 | Ribosomal L1 domain-containing protein 1 | RSL1D1 | 6869100 | 4114300 | 14385000 | 8260300 | 15839000 | 8901200 |
| O76041-2 | | | 163450 | 0 | 0 | 0 | 0 | 0 |
| O76071 | Probable cytosolic iron-sulfur protein assembly protein CIAO1 | CIAO1 | 326810 | 230330 | 754210 | 248230 | 435120 | 184620 |
| O76094;O76094-2;R4GNC1;D6RDY6 | Signal recognition particle subunit SRP72 | SRP72 | 1023700 | 719220 | 2052200 | 1246900 | 2054200 | 1019700 |
| O94763-2;O94763-3;O94763;O94763-4;I3NI51 | Unconventional prefoldin RPB5 interactor 1 | URI1 | 0 | 269350 | 535250 | 213090 | 622990 | 294900 |
| O94822;O94822-3;S4R3T2;H7BYG8;O94822-2 | E3 ubiquitin-protein ligase listerin | LTN1 | 245900 | 133730 | 348820 | 206660 | 270400 | 192760 |
| O94906;O94906-2 | Pre-mRNA-processing factor 6 | PRPF6 | 188330 | 136140 | 460590 | 374540 | 528310 | 241260 |
| O94927;K7EQ61 | HAUS augmin-like complex subunit 5 | HAUS5 | 121990 | 178440 | 228150 | 0 | 0 | 0 |
| O94973-3;O94973;O94973-2;H0YEG0;H0YDE9;E9PR62;E9PPZ3;E9PS94;E9PQP4;E9PNC4;E9PPY8 | AP-2 complex subunit alpha-2 | AP2A2 | 97793 | 0 | 93605 | 0 | 0 | 0 |
| O94986-1;O94986-2;O94986-3;O94986 | Centrosomal protein of 152 kDa | CEP152 | 0 | 0 | 0 | 0 | 0 | 0 |
| O94992 | Protein HEXIM1 | HEXIM1 | 1246600 | 783070 | 1589600 | 0 | 0 | 0 |
| O95218-2;O95218 | Zinc finger Ran-binding domain-containing protein 2 | ZRANB2 | 2169800 | 1359000 | 1561900 | 104050 | 606440 | 0 |
| O95235-2;O95235 | Kinesin-like protein KIF20A | KIF20A | 0 | 0 | 88895 | 0 | 0 | 0 |
| O95239-2;O95239;Q2VIQ3 | Chromosome-associated kinesin KIF4A | KIF4A | 364160 | 90781 | 299130 | 213580 | 223860 | 154190 |
| O95295 | SNARE-associated protein Snapin | SNAPIN | 1969200 | 1197500 | 2266100 | 507890 | 1105300 | 456620 |
| O95347;O95347-2;Q5T821 | Structural maintenance of chromosomes protein 2 | SMC2 | 3472400 | 1601100 | 4502900 | 1263300 | 2540200 | 1332100 |
| O95373 | Importin-7 | IPO7 | 77132 | 22135 | 34284 | 0 | 0 | 0 |
| Q5H8X8;O95399;O95399-2 | Urotensin-2 | UTS2 | 44227000 | 49159000 | 74770000 | 34393000 | 35756000 | 21013000 |
| O95400 | CD2 antigen cytoplasmic tail-binding protein 2 | CD2BP2 | 4955900 | 3917800 | 5272200 | 2389900 | 2546000 | 1523900 |
| O95456;O95456-2;F8WBH7 | Proteasome assembly chaperone 1 | PSMG1 | 102700000 | 65504000 | 64521000 | 15756000 | 16277000 | 13969000 |
| O95714 | E3 ubiquitin-protein ligase HERC2 | HERC2 | 137190 | 34791 | 135790 | 61715 | 94836 | 52709 |
| O95777;F2Z2Y6;C9JIZ0;C9JNV3 | U6 snRNA-associated Sm-like protein LSm8 | LSM8 | 8124000 | 4769300 | 7630700 | 6398200 | 7751100 | 5171100 |
| O95782-2;O95782 | AP-2 complex subunit alpha-1 | AP2A1 | 5652900 | 1353800 | 3676800 | 1907400 | 2469500 | 1300400 |
| Q5JW30;O95793-2;O95793;A0A087X1A5;O95793-3;Q5JW28;F6UDC6 | Double-stranded RNA-binding protein Staufen homolog 1 | STAU1 | 6937600 | 3623800 | 8796100 | 6518500 | 9934800 | 5038800 |
| O95816;O95816-2;A2A296 | BAG family molecular chaperone regulator 2 | BAG2 | 4682600 | 1553900 | 2286300 | 690090 | 996300 | 457260 |
| O95985-3;O95985-2;O95985;C9JKE2;C9JT00;F2Z2A5;C9JEI7;H0Y6W0 | DNA topoisomerase 3-beta-1;DNA topoisomerase | TOP3B | 0 | 24548 | 70416 | 118470 | 124320 | 0 |
| P00338;P00338-3;P00338-4;P00338-5;P00338-2;F5GXH2;F5GYU2;F5GXY2;F5GZQ4;F5H5J4;F5H6W8;F5H8H6;F5GXC7;F5GWW2;F5GXU1 | L-lactate dehydrogenase A chain | LDHA | 15885000 | 5050800 | 9241500 | 1442700 | 1649100 | 823370 |
| P00367;P00367-3;P00367-2;P49448;REV__E7EV89;REV__P29375-2;REV__P29375;REV__F5H1F7;A6NF36 | Glutamate dehydrogenase 1, mitochondrial;Glutamate dehydrogenase 2, mitochondrial | GLUD1;GLUD2 | 6897500 | 3779600 | 13379000 | 504030 | 691160 | 197770 |
| P02545-2;P02545-6;P02545-3;P02545;H0YAB0;Q5TCI8;P02545-5;P02545-4 | Prelamin-A/C;Lamin-A/C | LMNA | 0 | 97179 | 264570 | 0 | 83286 | 23910 |
| P04259 | Keratin, type II cytoskeletal 6B | KRT6B | 255320 | 1276700 | 53980 | 65018 | 145390 | 291400 |
| P04406;E7EUT5;P04406-2 | Glyceraldehyde-3-phosphate dehydrogenase | GAPDH | 5227900 | 2568100 | 5315500 | 701310 | 1072600 | 642320 |
| P04637-4;P04637;E7EMR6;A0A087X1Q1;E7EQX7;J3KP33;A0A087WZU8;P04637-7;P04637-5;P04637-6;P04637-2;P04637-3 | Cellular tumor antigen p53 | TP53 | 177980 | 71534 | 0 | 40641 | 0 | 128620 |
| P04899;P04899-4;P04899-6;P04899-3;P04899-5;P04899-2 | Guanine nucleotide-binding protein G(i) subunit alpha-2 | GNAI2 | 402450 | 185730 | 644000 | 0 | 145140 | 86949 |
| P05023-4;P05023;P05023-2;P05023-3;M0R116;P13637;P13637-2;P13637-3;M0QXF2;P20648 | Sodium/potassium-transporting ATPase subunit alpha-1 | ATP1A1 | 563830 | 203870 | 644580 | 21885 | 171760 | 39690 |
| P05109 | Protein S100-A8;Protein S100-A8, N-terminally processed | S100A8 | 548960 | 0 | 0 | 0 | 0 | 0 |
| P05198;H0YJS4;G3V4T5 | Eukaryotic translation initiation factor 2 subunit 1 | EIF2S1 | 28948000 | 15730000 | 31656000 | 15163000 | 28847000 | 15577000 |
| P05387;H0YDD8 | 60S acidic ribosomal protein P2 | RPLP2 | 262230000 | 209820000 | 333690000 | 341760000 | 556590000 | 315890000 |
| P05388;F8VWS0;F8VU65;P05388-2;F8VZS0;F8VQY6;F8VRK7;G3V210;F8VW21;F8VS58;F8VPE8;Q8NHW5;F8VWV4;F8W1K8;F8VYN4 | 60S acidic ribosomal protein P0;60S acidic ribosomal protein P0-like | RPLP0;RPLP0P6 | 149030000 | 80049000 | 169790000 | 107580000 | 196500000 | 91812000 |
| P05455;E7ERC4;E9PGX9;E9PFL9 | Lupus La protein | SSB | 9301400 | 4328600 | 9321700 | 7850300 | 16365000 | 6244000 |
| P06730;P06730-2;D6RFJ3;D6RBW1;P06730-3;H0Y8J7 | Eukaryotic translation initiation factor 4E | EIF4E | 346840 | 219980 | 659820 | 310420 | 1210100 | 349730 |
| P06733;P06733-2;K7EM90;E5RGZ4;P13929-3;P13929-2;P13929;E5RI09;E5RG95;K7EPM1;K7EKN2;F5H0C8;P09104-2;P09104;U3KQP4;F5H1C3 | Alpha-enolase | ENO1 | 34041000 | 9698500 | 26971000 | 7328200 | 10962000 | 3690800 |
| P06748-2;P06748;P06748-3 | Nucleophosmin | NPM1 | 564840000 | 346750000 | 804230000 | 679630000 | 1273100000 | 696250000 |
| Q5VU59;P06753-2;A0A087WWU8;P06753-3;P06753-6;Q5VU61;D6R904;K7EP68;D6RFM2;H0YL80;K7EQW8 | | DKFZp686J1372;TPM3 | 140660000 | 138820000 | 178810000 | 3473800 | 3523700 | 998450 |
| Q16778;P33778;P23527;P06899;Q8N257;Q6DRA6;Q6DN03 | Histone H2B type 2-E;Histone H2B type 1-B;Histone H2B type 1-O;Histone H2B type 1-J;Histone H2B type 3-B | HIST2H2BE;HIST1H2BB;HIST1H2BO;HIST1H2BJ;HIST3H2BB | 11019000 | 10089000 | 13510000 | 10126000 | 10713000 | 9245100 |
| P07195;A8MW50;C9J7H8;F5H793;F5H5G7;F5H155;A0A087WUM2;G3XAP5;F5H245;Q6ZMR3;P07864;Q9BYZ2 | L-lactate dehydrogenase B chain;L-lactate dehydrogenase | LDHB | 7114400 | 4306000 | 7212200 | 1262000 | 2972700 | 1479200 |
| P07196;A0A087X0W2 | Neurofilament light polypeptide | NEFL | 29485000 | 28427000 | 60150000 | 4676900 | 2870900 | 2319900 |
| P07355;H0YKS4;H0YMD0;H0YMU9;H0YN42;P07355-2;A6NMY6;H0YMM1;H0YKZ7;H0YLV6;H0YMT9;H0YKX9;H0YNP5;H0YM50;H0YKV8;H0YKL9;H0YMW4;H0YN52;H0YMD9;H0YNA0;H0YN28;H0YL33;H0YNB8;H0YKN4;H0YLE2 | Annexin A2;Annexin;Putative annexin A2-like protein | ANXA2;ANXA2P2 | 16770000 | 8217400 | 17137000 | 522540 | 1939000 | 764630 |
| Q5JP53;P07437;Q5ST81;A0A075B736;Q5SQY0;I3L2F9;A6NNZ2;G3V2A3;CON__ENSEMBL:ENSBTAP00000025008;Q9H4B7;G3V5W4;G3V2R8;G3V3R4;G3V2N6 | Tubulin beta chain | TUBB | 25841000 | 16553000 | 24762000 | 27665000 | 18904000 | 12290000 |
| P07814;V9GYZ6;V9GZ76 | Bifunctional glutamate/proline--tRNA ligase;Glutamate--tRNA ligase;Proline--tRNA ligase | EPRS | 7281600 | 5046300 | 7781700 | 8461100 | 10935000 | 5033300 |
| P07900;P07900-2;G3V2J8;Q58FF6;Q58FG0;Q14568;REV__H3BMH3;Q58FG1;Q92538 | Heat shock protein HSP 90-alpha | HSP90AA1 | 30736000 | 12457000 | 29025000 | 19878000 | 24571000 | 12051000 |
| P08107;P08107-2;V9GZ37 | Heat shock 70 kDa protein 1A | HSPA1A | 260740000 | 134750000 | 200260000 | 121080000 | 150770000 | 89463000 |
| P08133-2;P08133;E7EMC6;E5RJF5;E5RK63;E5RI05;E5RFF0;H0YC77;E5RIU8;E5RJR0 | Annexin A6;Annexin | ANXA6 | 75479000 | 20030000 | 36104000 | 1623600 | 2443000 | 899520 |
| P08237;P08237-3;P08237-2;F8VP00;F8VX13;F8VNX2;F8VZQ1;F8W1J8;F8VSF7;F8VW30;F8VSL1;F8VUB8;F8VTQ3;F8VYK8;F8VZI0;F8VVE3;H0YHB8 | ATP-dependent 6-phosphofructokinase, muscle type | PFKM | 7228100 | 2008000 | 3392600 | 0 | 79808 | 32670 |
| P08238;Q58FF7;Q58FF8;Q5T9W8 | Heat shock protein HSP 90-beta;Putative heat shock protein HSP 90-beta-3 | HSP90AB1;HSP90AB3P | 125750000 | 58350000 | 116430000 | 70435000 | 85319000 | 43212000 |
| P08579 | U2 small nuclear ribonucleoprotein B | SNRPB2 | 16966000 | 6145500 | 21393000 | 16923000 | 23691000 | 12113000 |
| P08621-2;P08621;P08621-3;M0QYR1;P08621-4;M0QX04 | U1 small nuclear ribonucleoprotein 70 kDa | SNRNP70 | 18349000 | 12992000 | 23091000 | 16767000 | 30373000 | 10884000 |
| P08670;B0YJC4;B0YJC5;Q5JVS8;U3KPR1;B4DIR1;K7ELP4;H7C5W5;F8W835;K7EJU1;K7EKH9;CON__Q61726;A0A087X106;CON__O43790;CON__Q6NT21;CON__P78385;CON__Q14533;CON__P78386;P14136-3;P14136;P14136-2;P41219;P41219-2;O43790;P78385;Q14533;P78386 | Vimentin | VIM | 42683000 | 38239000 | 66811000 | 13756000 | 7618200 | 6437400 |
| P08758;E9PHT9;D6RBL5;D6RBE9;D6RCN3 | Annexin A5;Annexin | ANXA5 | 4323800 | 4436600 | 5683600 | 989530 | 655710 | 450960 |
| P09012;M0QXK2;M0R221;M0R2B8;M0QZG7;M0R268;M0R0G9 | U1 small nuclear ribonucleoprotein A | SNRPA | 13427000 | 9039400 | 14022000 | 17089000 | 24850000 | 9388600 |
| Q5TAL2;P09234 | U1 small nuclear ribonucleoprotein C | SNRPC | 6126200 | 4364600 | 6481100 | 7565500 | 4068500 | 4734200 |
| P09493-3;P09493-4;P09493-10;P09493-9;H0YL52;Q6ZN40;P09493;H0YKX5;H0YKP3;H0YKZ6;A0A087WTJ7 | Tropomyosin alpha-1 chain | TPM1 | 633690 | 177320 | 575690 | 0 | 0 | 0 |
| P09496-2;P09496-4;P09496-3;P09496;F8WF69;P09496-5;C9J8P9 | Clathrin light chain A | CLTA | 162930000 | 56758000 | 116970000 | 3936300 | 4358200 | 1775600 |
| P09497-2;P09497;H0Y9Q6;D6RJD1 | Clathrin light chain B | CLTB | 163020000 | 50188000 | 108650000 | 3516500 | 5134000 | 1276200 |
| P09525;Q6P452;P09525-2 | Annexin A4;Annexin | ANXA4 | 6467600 | 6721400 | 7446400 | 0 | 0 | 0 |
| P09661;H0YLR3;H0YMA0;H0YKK0 | U2 small nuclear ribonucleoprotein A | SNRPA1 | 11205000 | 11746000 | 15089000 | 8058700 | 15242000 | 9903900 |
| P09874;Q5VX84;Q5VX85 | Poly [ADP-ribose] polymerase 1 | PARP1 | 6342600 | 2633600 | 5581800 | 4645400 | 5759600 | 3226000 |
| P09972;K7EKH5;C9J8F3;J3KSV6;J3QKP5;A8MVZ9 | Fructose-bisphosphate aldolase C | ALDOC | 208150 | 186580 | 455760 | 0 | 25409 | 0 |
| P10155-3;P10155-5;P10155-4;P10155;H0Y9N5;P10155-2;D6RE09;G5E9R9 | 60 kDa SS-A/Ro ribonucleoprotein | TROVE2 | 4132300 | 2193900 | 3384600 | 937070 | 1488700 | 961220 |
| P10599-2;P10599 | Thioredoxin | TXN | 32061000 | 21682000 | 38678000 | 22250000 | 27592000 | 14385000 |
| P10809 | 60 kDa heat shock protein, mitochondrial | HSPD1 | 0 | 20471 | 70224 | 34875 | 126410 | 67346 |
| P11021;REV__H3BUZ0;REV__Q9NXD2-2;REV__Q9NXD2 | 78 kDa glucose-regulated protein | HSPA5 | 102120000 | 59431000 | 83230000 | 36556000 | 39980000 | 22926000 |
| P11142;E9PKE3;E9PNE6;P11142-2;A8K7Q2;E9PS65;E9PLF4;E9PQQ4;E9PQK7;E9PK54;E9PM13;E9PPY6;E9PN25;E9PI65 | Heat shock cognate 71 kDa protein | HSPA8 | 273430000 | 126790000 | 204500000 | 94769000 | 131170000 | 81407000 |
| P11171-5;P11171-2;P11171;P11171-6;P11171-4;P11171-3;P11171-7 | Protein 4.1 | EPB41 | 309330 | 94357 | 450210 | 0 | 0 | 0 |
| P11279;P11279-2 | Lysosome-associated membrane glycoprotein 1 | LAMP1 | 1504200 | 512150 | 1140400 | 0 | 0 | 0 |
| P11387 | DNA topoisomerase 1 | TOP1 | 1272100 | 659020 | 1834300 | 73981 | 338990 | 172600 |
| P11388;P11388-2;P11388-3;P11388-4;J3KTB7 | DNA topoisomerase 2-alpha | TOP2A | 1727500 | 929440 | 2276100 | 53544 | 205820 | 51887 |
| Q5HY81;P11441 | Ubiquitin-like protein 4A | UBL4A | 4617500 | 4191800 | 5071000 | 0 | 0 | 229360 |
| P12004 | Proliferating cell nuclear antigen | PCNA | 30901000 | 13416000 | 46752000 | 13806000 | 27252000 | 11047000 |
| P12081-3;P12081-2;P12081-4;P12081;B4E1C5;B4DDD8;B3KWE1;E7ETE2;D6RF05;D6RJE6 | Histidine--tRNA ligase, cytoplasmic | HARS | 14989000 | 9280900 | 20862000 | 6458600 | 16585000 | 7649700 |
| P12270 | Nucleoprotein TPR | TPR | 0 | 0 | 0 | 0 | 0 | 0 |
| P12277;G3V4N7;G3V461;H0YJG0 | Creatine kinase B-type | CKB | 122420 | 275810 | 593290 | 111510 | 71948 | 40892 |
| P12532;C9JSQ1;F8WCN3;P12532-2;C9JJX8;C9J995;C9J8F6 | Creatine kinase U-type, mitochondrial | CKMT1A;CKMT1B | 485580 | 225020 | 455270 | 0 | 39273 | 0 |
| P12814;P12814-3;P12814-2;P12814-4;H9KV75;H0YJW3;P35609-2;P35609;H7C5W8;Q08043;G3V2N5;A0A087WSZ2;G3V2W4;H0YJ11;F6THM6;G3V5M4;G3V2X9;G3V2E8;G3V380 | Alpha-actinin-1 | ACTN1 | 8689500 | 1489700 | 2486000 | 19489 | 0 | 0 |
| P12956;B1AHC9;P12956-2;P51449-2;P51449 | X-ray repair cross-complementing protein 6 | XRCC6 | 19250000 | 11814000 | 37886000 | 20955000 | 31155000 | 18228000 |
| P13010;C9JZ81 | X-ray repair cross-complementing protein 5 | XRCC5 | 18662000 | 9674200 | 28017000 | 21149000 | 34145000 | 17941000 |
| P13489;E9PMA9;E9PMN0;E9PIK5;E9PMJ3;E9PLZ3;E9PIM9;H0YCR7 | Ribonuclease inhibitor | RNH1 | 40182 | 0 | 0 | 1217900 | 612420 | 565350 |
| P13639;M0R0I6 | Elongation factor 2 | EEF2 | 61764000 | 28981000 | 63145000 | 79634000 | 156940000 | 81418000 |
| P13667;C9JMN9 | Protein disulfide-isomerase A4 | PDIA4 | 4492000 | 1859800 | 5375400 | 3105000 | 3603100 | 2227500 |
| P13674-3;P13674-2;P13674 | Prolyl 4-hydroxylase subunit alpha-1 | P4HA1 | 667930 | 451510 | 698550 | 35763 | 115350 | 24978 |
| P13797;P13797-3;P13797-2;F8W8D8;U3KQI3;H7C4N2;F2Z2Z9;Q5TBN3;Q14651;C9JAM8 | Plastin-3 | PLS3 | 35121000 | 4146200 | 6450000 | 227720 | 615920 | 247020 |
| P13861-2;P13861;H7C330;H7C4A9;C9J830 | cAMP-dependent protein kinase type II-alpha regulatory subunit | PRKAR2A | 1911400 | 1121300 | 1968900 | 361420 | 676810 | 248140 |
| P14625;Q96GW1;H0YIV0;Q58FF3;F8W026;F8VPC7 | Endoplasmin | HSP90B1 | 7858400 | 3865800 | 9060000 | 3006500 | 5897900 | 3032400 |
| P14868;P14868-2;H7BZ35;C9J7S3;C9JLC1;C9JQM9;H7C278 | Aspartate--tRNA ligase, cytoplasmic | DARS | 13175000 | 8888900 | 18449000 | 12471000 | 19709000 | 9297600 |
| P15880;H0YEN5;E9PQD7;I3L404;E9PMM9;E9PM36;E9PPT0;H3BNG3 | 40S ribosomal protein S2 | RPS2 | 193090000 | 136010000 | 215270000 | 162490000 | 319470000 | 160840000 |
| P15924;P15924-3;P15924-2 | Desmoplakin | DSP | 446180 | 386070 | 479700 | 65049 | 100520 | 35898 |
| P16402;P10412;Q02539;P22492;P16401 | Histone H1.3;Histone H1.4 | HIST1H1D;HIST1H1E | 0 | 0 | 0 | 110580 | 0 | 0 |
| P16403 | Histone H1.2 | HIST1H1C | 39980000 | 19316000 | 31268000 | 16351000 | 22477000 | 11145000 |
| P16989;P16989-2;P16989-3;H0YG22 | Y-box-binding protein 3 | YBX3 | 40743000 | 33532000 | 63850000 | 33673000 | 58010000 | 36275000 |
| P17066;P48741 | Heat shock 70 kDa protein 6 | HSPA6 | 63603000 | 32322000 | 49820000 | 29419000 | 36523000 | 20686000 |
| P17252;J3KRN5 | Protein kinase C alpha type | PRKCA | 133220 | 128250 | 378590 | 0 | 0 | 0 |
| P17812;P17812-2 | CTP synthase 1 | CTPS1 | 2937500 | 1419400 | 2278800 | 168930 | 922710 | 336820 |
| P17858;P17858-2;F8WEU2;V9GYV7;Q5VSR5 | ATP-dependent 6-phosphofructokinase, liver type | PFKL | 27171000 | 17261000 | 20131000 | 1156000 | 1922600 | 471030 |
| P17980;E9PM69;E9PKD5;E9PMD8 | 26S protease regulatory subunit 6A | PSMC3 | 1087800 | 902300 | 731440 | 0 | 0 | 0 |
| P17987;E7EQR6;F5H282;E7ERF2;F5H136;F5H7Y1;F5H726;F5GZI8;F5H676 | T-complex protein 1 subunit alpha | TCP1 | 5028900 | 2333700 | 5544700 | 2109400 | 2891500 | 793760 |
| P18077;C9K025;F8WBS5;F8WB72 | 60S ribosomal protein L35a | RPL35A | 194870000 | 158290000 | 345610000 | 286970000 | 480860000 | 263350000 |
| P18124;A8MUD9;C9JIJ5;C9JZ88 | 60S ribosomal protein L7 | RPL7 | 192470000 | 149590000 | 294950000 | 141390000 | 307720000 | 183300000 |
| P18206-2;P18206;Q5JQ13;P18206-3 | Vinculin | VCL | 1131500 | 460960 | 838700 | 298290 | 162170 | 98217 |
| P18669 | Phosphoglycerate mutase 1 | PGAM1 | 6125500 | 10848000 | 14263000 | 0 | 7058600 | 4660100 |
| P19338;C9JYW2;C9JLB1;C9J1H7;C9JWL1 | Nucleolin | NCL | 161150000 | 96437000 | 237510000 | 359670000 | 660350000 | 341330000 |
| P19387;H3BRR2 | DNA-directed RNA polymerase II subunit RPB3 | POLR2C | 6552400 | 2912500 | 6129300 | 1945700 | 2197100 | 729260 |
| P19525-2;P19525;F8WBH4;C9JZT2 | Interferon-induced, double-stranded RNA-activated protein kinase | EIF2AK2 | 432950 | 221810 | 1012100 | 902490 | 2824200 | 1328900 |
| P19784;H3BNI9;H3BSA1;H3BV19;P17540 | Casein kinase II subunit alpha | CSNK2A2 | 3157400 | 1064500 | 1942300 | 2543900 | 7620200 | 2332000 |
| P20042 | Eukaryotic translation initiation factor 2 subunit 2 | EIF2S2 | 28117000 | 15488000 | 31641000 | 17807000 | 29909000 | 13444000 |
| P20073-2;P20073 | Annexin A7 | ANXA7 | 1514000 | 1333700 | 3934300 | 62043 | 0 | 0 |
| P20290;P20290-2;D6RDG3;H0Y9Y1 | Transcription factor BTF3 | BTF3 | 47300000 | 29060000 | 61395000 | 29456000 | 51128000 | 27960000 |
| P20618 | Proteasome subunit beta type-1 | PSMB1 | 1301200000 | 860530000 | 1144700000 | 817440000 | 613810000 | 493430000 |
| Q5HY54;P21333-2;P21333;Q60FE5;A0A087WWY3;F8WE98;H0Y5F3;H0Y5C6 | Filamin-A | FLNA | 11819000 | 4039200 | 5040600 | 243080 | 335170 | 23307 |
| P21359-2;P21359-6;P21359;H0Y465;J3KSB5;P21359-4;J3KRT8;J3QQN8;P21359-3;P21359-5 | Neurofibromin;Neurofibromin truncated | NF1 | 123240 | 39374 | 278520 | 44730 | 69541 | 128770 |
| P22059 | Oxysterol-binding protein 1 | OSBP | 146620 | 0 | 116100 | 0 | 0 | 0 |
| P22314-2;P22314;Q5JRR6;Q5JRR9;Q5JRS0;Q5JRS1;Q5JRS3;Q5JRS2 | Ubiquitin-like modifier-activating enzyme 1 | UBA1 | 1522200 | 590470 | 1173000 | 567420 | 384840 | 151790 |

| | | | | | | | | |
|---|---|---|---|---|---|---|---|---|
| Q32Q12;P22392;P22392-2;C9K028;F6XY72;E5RHP0;E7ERL0;J3KPD9;O60361 | Nucleoside diphosphate kinase;Nucleoside diphosphate kinase B;Putative nucleoside diphosphate kinase | NME1-NME2;NME2;NME1;NME2P1 | 461910 | 0 | 593930 | 0 | 90536 | 0 |
| P22626;P22626-2;A0A087WUI2 | Heterogeneous nuclear ribonucleoproteins A2/B1 | HNRNPA2B1 | 48296000 | 36018000 | 83671000 | 65110000 | 101210000 | 76979000 |
| P23246;P23246-2 | Splicing factor, proline- and glutamine-rich | SFPQ | 7934300 | 4682700 | 9239100 | 6447200 | 13716000 | 6566800 |
| P23284 | Peptidyl-prolyl cis-trans isomerase B | PPIB | 12504000 | 5840900 | 10393000 | 5641300 | 7090600 | 4145400 |
| P23396;P23396-2;E9PL09;E9PPU1;F2Z2S8;H0YCJ7;H0YF32;H0YEU2;E9PQ96;E9PJH4;E9PK82;E9PSF4;E9PQX2;H0YES8;E9PJN9 | 40S ribosomal protein S3 | RPS3 | 397850000 | 221080000 | 424660000 | 296250000 | 604260000 | 295230000 |
| P23526-2;P23526 | Adenosylhomocysteinase | AHCY | 516090 | 221480 | 677900 | 0 | 0 | 0 |
| P23528;E9PP50;E9PK25;G3V1A4;E9PQB7;E9PLJ3;E9PS23 | Cofilin-1 | CFL1 | 1283400000 | 596350000 | 702670000 | 8703700 | 10921000 | 12582000 |
| P24534;C9JZW3;F8WF65;F2Z2G2 | Elongation factor 1-beta | EEF1B2 | 68036000 | 34442000 | 59003000 | 13489000 | 35846000 | 12066000 |
| P24666;G5E9R5;P24666-4;F2Z2Q9;P24666-3;P24666-2;D3YTI2 | Low molecular weight phosphotyrosine protein phosphatase | ACP1 | 15699000 | 12679000 | 19088000 | 73805 | 385780 | 233070 |
| P24928;A0A087WWE2;P24928-2;Q9NY15-2;Q9NY15 | DNA-directed RNA polymerase II subunit RPB1;DNA-directed RNA polymerase | POLR2A | 1245100 | 990280 | 1946100 | 633460 | 1111800 | 555780 |
| P25205;P25205-2;J3KQ69 | DNA replication licensing factor MCM3 | MCM3 | 666190 | 430630 | 779590 | 180120 | 320520 | 123850 |
| P25311;C9JEV0 | Zinc-alpha-2-glycoprotein | AZGP1 | 572680 | 346320 | 514200 | 0 | 102820 | 76451 |
| P25398 | 40S ribosomal protein S12 | RPS12 | 338970000 | 215500000 | 247340000 | 254180000 | 403770000 | 86646000 |
| P25786;P25786-2;F5GX11;B4DEV8;F5H112 | Proteasome subunit alpha type-1 | PSMA1 | 2734200000 | 1912100000 | 2311000000 | 2226000000 | 1512000000 | 1191500000 |
| P25787;C9JCK5 | Proteasome subunit alpha type-2 | PSMA2 | 1309000000 | 810400000 | 1004100000 | 996750000 | 691220000 | 522380000 |
| P25788-2;P25788;G3V3W4;G3V5N4 | Proteasome subunit alpha type-3 | PSMA3 | 3953100000 | 2093200000 | 2807900000 | 2662000000 | 1931800000 | 1346900000 |
| P25789;H0YN18;H0YMZ1;H0YL69;H0YMA1;H0YKT8;H0YMI6;H0YKS0;H0YLC2;H0YLS6;H0YMV3 | Proteasome subunit alpha type-4;Proteasome subunit alpha type;Proteasome subunit beta type | PSMA4 | 2957300000 | 1745000000 | 2231800000 | 1878500000 | 1337700000 | 985790000 |
| P26038 | Moesin | MSN | 0 | 62855 | 213020 | 0 | 137730 | 217100 |
| P26196;Q8IV96 | Probable ATP-dependent RNA helicase DDX6 | DDX6 | 453810 | 84453 | 0 | 185080 | 680570 | 0 |
| P26358-2;P26358;P26358-3;K7ENW7;K7ELB1;K7ENQ6;K7EJL0;K7ERQ1;K7ER10 | DNA (cytosine-5)-methyltransferase 1 | DNMT1 | 2171200 | 1440700 | 2384800 | 477620 | 1318500 | 707140 |
| P26373;P26373-2;J3QSB4;J3KS98;H3BUK8;H3BTH3 | 60S ribosomal protein L13 | RPL13 | 225080000 | 186050000 | 360310000 | 333220000 | 606890000 | 424040000 |
| P26583;D6R9A6 | High mobility group protein B2 | HMGB2 | 7311900 | 1481600 | 8351100 | 1038900 | 2122000 | 1552700 |
| P26599;P26599-2;P26599-3;A6NLN1;A0A087WUW5;K7EKJ7;K7EK45;K7ELW5 | Polypyrimidine tract-binding protein 1 | PTBP1 | 49321000 | 14879000 | 75656000 | 24141000 | 90604000 | 26367000 |
| P26639;P26639-2 | Threonine--tRNA ligase, cytoplasmic | TARS | 610150 | 395640 | 1117600 | 131600 | 380500 | 105490 |
| P26640;P26640-2;A2ABF4;H0Y426 | Valine--tRNA ligase | VARS | 8499700 | 4574300 | 11843000 | 3060500 | 7718900 | 3144000 |
| P26641;P26641-2 | Elongation factor 1-gamma | EEF1G | 144370000 | 89855000 | 125260000 | 62159000 | 131640000 | 69113000 |
| P27348;E9PG15 | 14-3-3 protein theta | YWHAQ | 78576000 | 30207000 | 43454000 | 11536000 | 13240000 | 7395300 |
| X1WI28;P27635;Q96L21;B8A6G2;A6QRI9 | 60S ribosomal protein L10 | RPL10 | 8793700 | 9601400 | 17882000 | 6246000 | 14484000 | 9034300 |
| P27694;I3L4R8 | Replication protein A 70 kDa DNA-binding subunit;Replication protein A 70 kDa DNA-binding subunit, N-terminally processed | RPA1 | 1390100 | 1190600 | 1550700 | 2394800 | 2669300 | 926190 |
| P27797;K7EJB9;K7EL50 | Calreticulin | CALR | 5668100 | 2112400 | 4638500 | 3195200 | 3537700 | 2758200 |
| P28065;A2ACR1;P28065-2;A2ACR0;A2ACQ9 | Proteasome subunit beta type-9;Proteasome subunit beta type | PSMB9 | 6049500 | 4001500 | 4937200 | 3144400 | 2618800 | 1226100 |
| P28066;P28066-2 | Proteasome subunit alpha type-5 | PSMA5 | 1247600000 | 756560000 | 953320000 | 658220000 | 554120000 | 374810000 |
| P28070 | Proteasome subunit beta type-4 | PSMB4 | 2370700000 | 1328800000 | 1842100000 | 1481500000 | 1153800000 | 837730000 |
| P28072;A0A087X2I4;I3L3X7 | Proteasome subunit beta type-6 | PSMB6 | 2324900000 | 1596000000 | 2089100000 | 1499600000 | 1223900000 | 841030000 |
| P28074;P28074-2;H0YJM8;G8JLC2;G3V3K3 | Proteasome subunit beta type-5 | PSMB5 | 1466300000 | 888920000 | 1005500000 | 730140000 | 744120000 | 480100000 |
| P28074-3 | Proteasome subunit beta type-5 | PSMB5 | 7043100 | 0 | 0 | 0 | 0 | 0 |
| P28482-2;P28482 | Mitogen-activated protein kinase 1 | MAPK1 | 157430 | 0 | 0 | 0 | 0 | 0 |
| P29083;C9J329;E9PER7 | General transcription factor IIE subunit 1 | GTF2E1 | 55115 | 44986 | 331300 | 0 | 0 | 0 |
| P29401;P29401-2;E9PFF2;F8W888 | Transketolase | TKT | 1359300 | 1281300 | 1812700 | 209460 | 248200 | 77231 |
| P29692;E9PRY8;P29692-2;E9PK01;P29692-4;E9PQ49;E9PI39;E9PQZ1;A0A087X1X7;E9PPR1;E9PL12;E9PMW7;E9PIZ1;H0YCK7;E9PK06;H0YE72;H0YE58;E9PK72;E9PKK3;E9PNW6;E9PJD0 | Elongation factor 1-delta | EEF1D | 55651000 | 39152000 | 69951000 | 18265000 | 44317000 | 24349000 |
| P29692-3;E9PL71;E9PN91 | Elongation factor 1-delta | EEF1D | 1039600 | 267870 | 321920 | 0 | 139350 | 189610 |
| P30040;F8VY02;F8W1G0 | Endoplasmic reticulum resident protein 29 | ERP29 | 4408700 | 2753900 | 4029800 | 391990 | 1639900 | 1385900 |
| P30050;P30050-2 | 60S ribosomal protein L12 | RPL12 | 268380000 | 154830000 | 272080000 | 232260000 | 448680000 | 215900000 |
| P30086 | Phosphatidylethanolamine-binding protein 1;Hippocampal cholinergic neurostimulating peptide | PEBP1 | 2718000 | 184000 | 1698200 | 11669000 | 127160 | 0 |
| P30101 | Protein disulfide-isomerase A3 | PDIA3 | 7031000 | 2961500 | 7068100 | 5220700 | 3562300 | 2232600 |
| P30153;B3KQV6;C9J9C1;E9PH38;E9PPI5;E9PHZ6;P30154-5;P30154-4;P30154;P30154-2 | Serine/threonine-protein phosphatase 2A 65 kDa regulatory subunit A alpha isoform | PPP2R1A | 13586000 | 5295000 | 8974100 | 4487200 | 3384800 | 2475400 |
| P30260;P30260-2;G5EA36 | Cell division cycle protein 27 homolog | CDC27 | 82724 | 39929 | 115060 | 265080 | 287880 | 184770 |
| P30419;B7Z8J4;P30419-2;K7EN82;K7EN42;Q5VUC6;Q5VUC7;O60551 | Glycylpeptide N-tetradecanoyltransferase 1 | NMT1 | 1354000 | 1197700 | 3067800 | 1375300 | 3976000 | 1866100 |
| P30533;D6REW6 | Alpha-2-macroglobulin receptor-associated protein | LRPAP1 | 2044500 | 1290500 | 1400300 | 195070 | 558780 | 115160 |
| P30622-2;P30622-1;P30622;F5H0N7;J3KP58 | CAP-Gly domain-containing linker protein 1 | CLIP1 | 93983 | 8603.6 | 0 | 0 | 0 | 12073 |
| P31689;P31689-2 | DnaJ homolog subfamily A member 1 | DNAJA1 | 17368000 | 8825500 | 11293000 | 276220 | 1909300 | 1054900 |
| P31942-2;P31942;P31942-3 | Heterogeneous nuclear ribonucleoprotein H3 | HNRNPH3 | 730790 | 824720 | 2636400 | 799050 | 1651600 | 945320 |
| P31944 | Caspase-14;Caspase-14 subunit p17, mature form;Caspase-14 subunit p10, mature form;Caspase-14 subunit p20, intermediate form;Caspase-14 subunit p8, intermediate form | CASP14 | 0 | 0 | 0 | 0 | 0 | 0 |
| P31946-2;P31946;Q4VY20;Q4VY19 | 14-3-3 protein beta/alpha;14-3-3 protein beta/alpha, N-terminally processed | YWHAB | 78587000 | 33653000 | 48098000 | 13690000 | 15397000 | 7958900 |
| P31948;P31948-2;P31948-3;F5H783;F5GXD8;H0YGI8 | Stress-induced-phosphoprotein 1 | STIP1 | 15753000 | 4499500 | 8791800 | 3122100 | 6969000 | 2005900 |
| P31949 | Protein S100-A11;Protein S100-A11, N-terminally processed | S100A11 | 55252000 | 103940000 | 146190000 | 10252000 | 14755000 | 8706100 |
| P32119;A6NIW5;P32119-2 | Peroxiredoxin-2 | PRDX2 | 8706800 | 3630200 | 9080000 | 2299000 | 7244300 | 3286700 |
| P33176;J3KNA1;O60282-2;O60282;Q12840 | Kinesin-1 heavy chain | KIF5B | 120290 | 31931 | 264060 | 70495 | 0 | 0 |
| P33991;E5RFR3;E5RFJ8 | DNA replication licensing factor MCM4 | MCM4 | 205630 | 46113 | 232590 | 56549 | 0 | 17156 |
| P33993;C9J8M6;P33993-2;P33993-3 | DNA replication licensing factor MCM7 | MCM7 | 77634 | 0 | 35251 | 0 | 0 | 0 |
| P34931 | Heat shock 70 kDa protein 1-like | HSPA1L | 150680 | 0 | 0 | 0 | 0 | 0 |
| P35249;C9JZI1;P35249-2;C9J8M3;C9JXZ7;C9JTT7;C9JGY5;F8WE44;H7C1P0;C9JW34 | Replication factor C subunit 4 | RFC4 | 5151400 | 3563900 | 7041500 | 2481800 | 4606500 | 2266100 |
| P35250-2;P35250;H7C5P4;A0A087WVY3;H7C5Q7;F8WC37;H7C5A0;H7C5G4;H7C5S7;H7C5P1;REV__Q96KP1 | Replication factor C subunit 2 | RFC2 | 8034800 | 3760800 | 9330000 | 3772600 | 6347100 | 2977100 |
| P35251-2;P35251;H0Y8U4 | Replication factor C subunit 1 | RFC1 | 1314400 | 139090 | 391760 | 215060 | 721440 | 342460 |
| P35579;P35579-2;Q5BKV1;B1AH99;E7ERA5;A1L2Z2;REV__S4R3H4;REV__E7EQT4;REV__Q9UKV3-5;REV__Q9UKV3 | Myosin-9 | MYH9 | 3730800 | 1725500 | 2616000 | 109660 | 121980 | 66886 |
| P35580;P35580-5;P35580-2;P35580-3;P35580-4 | Myosin-10 | MYH10 | 1403800 | 389280 | 946500 | 124220 | 111150 | 69968 |
| P35606-2;P35606;H0YAC7;D6R997;D6RBZ7;D6RCL6;D6RBG7;D6RBT6 | Coatomer subunit beta | COPB2 | 4046000 | 1056700 | 2411900 | 408630 | 691770 | 190280 |
| P35612-2;P35612-8;P35612-9;P35612-4;P35612-3;P35612 | Beta-adducin | ADD2 | 402890 | 245320 | 213640 | 0 | 43935 | 33673 |
| P35998;P35998-2;C9JLS9 | 26S protease regulatory subunit 7 | PSMC2 | 1063400000 | 545540000 | 650200000 | 114180000 | 102360000 | 67779000 |
| P36021 | Monocarboxylate transporter 8 | SLC16A2 | 14436000 | 10025000 | 19206000 | 21665000 | 11275000 | 8377200 |
| P36578;H3BM89;H3BTP7;H3BU31 | 60S ribosomal protein L4 | RPL4 | 58220000 | 34552000 | 89035000 | 60583000 | 143310000 | 73577000 |
| P36915-2;P36915 | Guanine nucleotide-binding protein-like 1 | GNL1 | 404460 | 278910 | 1099000 | 359320 | 915550 | 416120 |
| X6RJP6;P37802;P37802-2 | Transgelin-2 | TAGLN2 | 28914000 | 11647000 | 15517000 | 433830 | 377410 | 261000 |
| P38159;P38159-2;H0Y6E7;H3BT71;Q96E39;H3BUY5;H3BR27;P38159-3;H3BNC1;O75526;Q8N7X1 | RNA-binding motif protein, X chromosome;RNA-binding motif protein, X chromosome, N-terminally processed;RNA binding motif protein, X-linked-like-1 | RBMX;RBMXL1 | 4488400 | 4410700 | 7650300 | 4481600 | 7966500 | 3536500 |
| P38646;D6RJI2;D6RA73;H0YBG6 | Stress-70 protein, mitochondrial | HSPA9 | 3467300 | 857520 | 3766500 | 104670 | 598380 | 58732 |
| P38919;I3L3H2;C9JUF0;J3QL52;J3QQP0;E9PBH4;F8WE11 | Eukaryotic initiation factor 4A-III;Eukaryotic initiation factor 4A-III, N-terminally processed | EIF4A3 | 6032700 | 4367200 | 5442900 | 2891200 | 5986500 | 2637600 |
| P39019;M0R2L9;M0QXK4;M0QYF7;M0R140;A0A075B6E2 | 40S ribosomal protein S19 | RPS19 | 321730000 | 249220000 | 434940000 | 451400000 | 578500000 | 417410000 |
| P39023;G5E9G0;B5MCW2;H7C422;H7C3M2;F8WCR1;Q92901 | 60S ribosomal protein L3 | RPL3 | 233810000 | 151660000 | 291720000 | 189930000 | 403000000 | 174540000 |
| P39748;P39748-2;F5H1Y3;I3L3E9;CON__P01030;CON__ENSEMBL:ENSBTAP00000007350 | Flap endonuclease 1 | FEN1 | 3008100 | 1934000 | 4721000 | 2791000 | 6804000 | 2609100 |
| P40222 | Alpha-taxilin | TXLNA | 2392400 | 1611000 | 5293500 | 618180 | 2950300 | 1510700 |
| P40227;P40227-2;Q92526-2;Q92526;Q92526-3;J3KRI6 | T-complex protein 1 subunit zeta | CCT6A | 9269900 | 6001000 | 10446000 | 9488100 | 5377500 | 4330300 |
| P40429;M0QYS1;Q8J015;Q6NVV1;M0QZU1;A0A096LPE0 | 60S ribosomal protein L13a | RPL13A;RPL13a | 135600000 | 96171000 | 204720000 | 139340000 | 241640000 | 154730000 |
| P40926;G3XAL0;P40926-2 | Malate dehydrogenase, mitochondrial;Malate dehydrogenase | MDH2 | 1208200 | 282550 | 1863700 | 109090 | 339000 | 173490 |
| P40937-2;P40937;F5H304;C9JH72;F5H0U6 | Replication factor C subunit 5 | RFC5 | 2506800 | 802660 | 2933300 | 658990 | 1562300 | 516320 |
| P40938-2;P40938;A0A087X270 | Replication factor C subunit 3 | RFC3 | 6460700 | 3873700 | 6717300 | 4366100 | 7256300 | 3380900 |
| P40939;H0YFD6 | Trifunctional enzyme subunit alpha, mitochondrial;Long-chain enoyl-CoA hydratase;Long chain 3-hydroxyacyl-CoA dehydrogenase | HADHA | 339850 | 39267 | 115250 | 0 | 0 | 0 |
| P41091;Q2VIR3;F8W810;Q2VIR3-2;H7BZU1 | Eukaryotic translation initiation factor 2 subunit 3;Putative eukaryotic translation initiation factor 2 subunit 3-like protein | EIF2S3;EIF2S3L | 35058000 | 24601000 | 31634000 | 31289000 | 25360000 | 32362000 |
| P41214;P41214-2;Q5SY38 | Eukaryotic translation initiation factor 2D | EIF2D | 662420 | 306430 | 840750 | 638260 | 1424400 | 521850 |
| P41227-2;P41227 | N-alpha-acetyltransferase 10 | NAA10 | 513940 | 0 | 580280 | 0 | 474690 | 0 |
| P41240;H3BUM9;H3BN15 | Tyrosine-protein kinase CSK | CSK | 1728700 | 650700 | 920020 | 535760 | 616810 | 358010 |
| P41250;H7C443 | Glycine--tRNA ligase | GARS | 90876 | 264220 | 292760 | 109810 | 83794 | 23597 |
| P42285;H0YAC4;H0Y8U3;D6REC7 | Superkiller viralicidic activity 2-like 2 | SKIV2L2 | 4210600 | 2650800 | 3616500 | 4967200 | 7367600 | 3314600 |
| P42677;Q5T4L4 | 40S ribosomal protein S27 | RPS27 | 345080000 | 144640000 | 364790000 | 129720000 | 323210000 | 37638000 |
| P42695;G3V1A9;E9PKK4;E9PQA3;E9PLE0 | Condensin-2 complex subunit D3 | NCAPD3 | 910780 | 287980 | 919290 | 331590 | 606040 | 264510 |
| P42696;A2A2V2;P42696-2;Q5TCT4 | RNA-binding protein 34 | RBM34 | 1422300 | 839190 | 2601200 | 2539800 | 4285900 | 2777300 |
| P42704 | Leucine-rich PPR motif-containing protein, mitochondrial | LRPPRC | 86336 | 45509 | 144440 | 41110 | 42735 | 10274 |
| P43246-2;P43246;V9H019;E9PHA6;A0A087X078;A0A087X1E7;V9H015;V9H0B2;C9J809 | DNA mismatch repair protein Msh2 | MSH2 | 4977100 | 2528300 | 6357800 | 1564000 | 3215500 | 1100200 |
| P43686;P43686-2 | 26S protease regulatory subunit 6B | PSMC4 | 636800000 | 355460000 | 445060000 | 82283000 | 68806000 | 40170000 |
| P46060;B0QYT5;B0QYT4;B0QYT6;H0Y4Q3 | Ran GTPase-activating protein 1 | RANGAP1 | 3635900 | 1442800 | 3000200 | 1259700 | 1029600 | 642450 |
| P46063;F5GYB7;F5H4P4;F5H3W0;F5H2L2;F8WA66;F8WD97 | ATP-dependent DNA helicase Q1 | RECQL | 1139400 | 975510 | 2027500 | 829270 | 2592300 | 1324500 |
| P46087;P46087-4;A0A087WV73;P46087-2;P46087-3;F5GWB7;F5H709;F5H5X6;F5H359;F5GYR3;F5H8G6 | Probable 28S rRNA (cytosine(4447)-C(5))-methyltransferase | NOP2 | 1576100 | 1969300 | 4815100 | 5233800 | 11105000 | 6350500 |
| P46379-2;P46379;P46379-3;P46379-4;P46379-5;F6S6P2;X6REW1;F6UR09;F6U1F2;F6XTU0;F6RG75;F6WML8;F6VEM6;F6X9W3;F6U341;F6TC96;H0Y4L1;K7EKW3;H0Y710;E7ESS3;O95164;Q71RG4-2;Q71RG4 | Large proline-rich protein BAG6 | BAG6 | 4943600 | 2489200 | 3235200 | 187490 | 68203 | 169630 |
| P46777;R4GNJ2;Q5T7N0 | 60S ribosomal protein L5 | RPL5 | 149750000 | 115660000 | 279010000 | 217450000 | 409970000 | 222630000 |
| P46778;G3V1B3;M0R181 | 60S ribosomal protein L21 | RPL21 | 64379000 | 57766000 | 154310000 | 86837000 | 175810000 | 78568000 |
| P46781;B5MCT8;C9JM19;F2Z3C0;A8MXK4 | 40S ribosomal protein S9 | RPS9 | 320850000 | 190270000 | 447630000 | 212750000 | 453950000 | 243260000 |
| P46783;F6U211;S4R435;Q9NQ39 | 40S ribosomal protein S10;Putative 40S ribosomal protein S10-like | RPS10;RPS10-NUDT3;RPS10P5 | 99738000 | 78723000 | 136650000 | 104520000 | 191160000 | 128650000 |
| P46821;D6RA40;D6RA32;E9PGC8;P78559;P78559-2 | Microtubule-associated protein 1B;MAP1B heavy chain;MAP1 light chain LC1 | MAP1B | 167170 | 30221 | 229780 | 152300 | 396710 | 144020 |
| P46939;P46939-2 | Utrophin | UTRN | 38123 | 5668.3 | 26536 | 0 | 0 | 0 |
| P46940;H0YLE8;A0A087WWP1;A0A087X225 | Ras GTPase-activating-like protein IQGAP1 | IQGAP1 | 3046800 | 10214000 | 5156200 | 414360 | 721350 | 300910 |
| P47813;A6NJH9;O14602;X6RAC9 | Eukaryotic translation initiation factor 1A, X-chromosomal;Eukaryotic translation initiation factor 1A, Y-chromosomal | EIF1AX;EIF1AY | 28072000 | 16175000 | 19230000 | 17396000 | 24683000 | 13548000 |
| P47897-2;P47897;H7C0R3;C9JSG9;C9J165;F2Z2V6 | Glutamine--tRNA ligase | QARS | 5802400 | 2345600 | 4739900 | 2525800 | 5484500 | 2383400 |
| P47914 | 60S ribosomal protein L29 | RPL29 | 291950000 | 169940000 | 393570000 | 295540000 | 597570000 | 368240000 |
| P48426;Q5JUT3;P48426-2 | Phosphatidylinositol 5-phosphate 4-kinase type-2 alpha | PIP4K2A | 1482500 | 288590 | 962200 | 501900 | 924030 | 327780 |
| R4GMR5;P48556;K7EJR3;K7EJC1;K7ERW6;K7ENY6 | 26S proteasome non-ATPase regulatory subunit 8 | PSMD8 | 483060000 | 263940000 | 294820000 | 44068000 | 41645000 | 29073000 |
| P48634-3;P48634;P48634-2;P48634-4 | Protein PRRC2A | PRRC2A | 348050 | 207820 | 449770 | 337330 | 1051800 | 463060 |
| P49006 | MARCKS-related protein | MARCKSL1 | 9366200 | 6187500 | 13976000 | 0 | 1119500 | 263050 |
| P49207 | 60S ribosomal protein L34 | RPL34 | 247030000 | 194800000 | 409110000 | 245180000 | 573820000 | 374460000 |
| P49321;P49321-4;Q5T624;P49321-2;P49321-3;E9PRH9;H0YDS9;E9PI86;H0YF33;E9PPR5;E9PQG5;E9PAU3;E9PKR5;E9PNB5 | Nuclear autoantigenic sperm protein | NASP | 1067900 | 259970 | 782340 | 581100 | 754760 | 420180 |
| P49327 | Fatty acid synthase;[Acyl-carrier-protein] S-acetyltransferase;[Acyl-carrier-protein] S-malonyltransferase;3-oxoacyl-[acyl-carrier-protein] synthase;3-oxoacyl-[acyl-carrier-protein] reductase;3-hydroxyacyl-[acyl-carrier-protein] dehydratase;Enoyl-[acyl-carrier-protein] reductase;Oleoyl-[acyl-carrier-protein] hydrolase | FASN | 887950 | 332800 | 1097200 | 111710 | 134970 | 49047 |
| P49590-2;P49590;B4DQ67 | Probable histidine--tRNA ligase, mitochondrial | HARS2 | 0 | 0 | 0 | 0 | 78739 | 0 |
| P49720;A0A087WXQ8;A0A087WY10 | Proteasome subunit beta type-3 | PSMB3 | 891810000 | 516510000 | 592790000 | 692020000 | 215990000 | 99449000 |
| P49770;H0YJJ8;G3V5E5 | Translation initiation factor eIF-2B subunit beta | EIF2B2 | 3235900 | 884330 | 1852200 | 1641900 | 3231500 | 1569300 |
| P49792;F8WBP7;J3KQ37;F8VYC4;J3KNE0;P0DJD0;P0DJD1;Q7Z3J3;A6NKT7;Q99666;O14715 | E3 SUMO-protein ligase RanBP2 | RANBP2 | 0 | 10697 | 17426 | 0 | 0 | 0 |
| P49841;P49841-2 | Glycogen synthase kinase-3 beta | GSK3B | 1524600 | 939200 | 635700 | 279550 | 444230 | 0 |
| P49915-2;P49915 | GMP synthase [glutamine-hydrolyzing] | GMPS | 2338100 | 1090800 | 1742300 | 451790 | 1021400 | 531710 |
| Q5SX87;P50395;V9GYJ7;V9GYF8;Q5SX90;Q5SX86;Q5SX91;P50395-2;P31150 | Rab GDP dissociation inhibitor beta;Rab GDP dissociation inhibitor alpha | GDI2;GDI1 | 320710 | 221190 | 2383000 | 337570 | 442970 | 61673 |
| Q3KNR6;P50502;H7C3I1;F6VDH7;A0A087X1H6;Q8NFI4;F8WAQ7 | Hsc70-interacting protein;Putative protein FAM10A5 | ST13;ST13P5 | 7436000 | 3328300 | 8897100 | 1691400 | 5030300 | 2081400 |
| P50570-3;P50570-2;P50570-5;P50570-4;P50570;K7EPK9;E5RIK2;E5RHK8;Q9UQ16-5;Q9UQ16-2;Q9UQ16-3;Q9UQ16;Q9UQ16-4;K7EMQ3 | Dynamin-2 | DNM2 | 773380 | 410710 | 536020 | 185600 | 253660 | 239020 |
| P50750;P50750-2;F8VXD2;F8VYH9;F8VZZ0;F8VTV8;F8VWX7;F8VZ51;K7EJ83;H0YAZ9;Q9BVE2;E7EUK8;E5RGN0;A0A087X209;A0A087WZU2;F5H6Z0;J3QSD7;P11802-2;Q00535-2;Q00535;P11802;Q00534;Q96Q40-2;Q96Q40-4;Q96Q40-3;O94921-3;Q96Q40-5;Q96Q40;O94921-2;O94921;Q07002;Q07002-2;Q00536;Q07002-3;Q00537-2;Q00537;Q00536-2;Q9NYV4-3;Q14004-2;Q9NYV4-2;Q9NYV4;Q14004 | Cyclin-dependent kinase 9 | CDK9 | 1975300 | 1358900 | 1685600 | 1626800 | 734120 | 274380 |
| P50990-2;P50990;P50990-3;H7C4C8;H7C2U0 | T-complex protein 1 subunit theta | CCT8 | 16103000 | 5590700 | 17027000 | 10427000 | 4348800 | 2462000 |
| P50991;P50991-2 | T-complex protein 1 subunit delta | CCT4 | 4893200 | 1953200 | 4303200 | 5844100 | 4011000 | 2880700 |

| | | | | | | | | |
|---|---|---|---|---|---|---|---|---|
| P50995-2;P50995;H0Y6E1;E5RIN3;P27216;P27216-2 | Annexin A11 | ANXA11 | 5057700 | 2607100 | 4072400 | 40449 | 146120 | 0 |
| P51116;I3L1Z2 | Fragile X mental retardation syndrome-related protein 2 | FXR2 | 0 | 0 | 869440 | 224050 | 855660 | 319960 |
| P51149;C9J4V0;C9J592;C9J8S3;C9J4S4;C9IZZ0;C9J7D1 | Ras-related protein Rab-7a | RAB7A | 4366600 | 1633300 | 2146200 | 533200 | 347430 | 203650 |
| P51572;P51572-2;C9JMD7;C9JSP1;C9JQ75;C9J0M4 | B-cell receptor-associated protein 31 | BCAP31 | 684400 | 413420 | 1121300 | 0 | 358170 | 0 |
| P51665;H3BNT7;H3BTM8;H3BQV2 | 26S proteasome non-ATPase regulatory subunit 7 | PSMD7 | 373030000 | 209710000 | 273910000 | 57927000 | 47741000 | 33195000 |
| P51991-2;P51991;H7C1J8 | Heterogeneous nuclear ribonucleoprotein A3 | HNRNPA3 | 6968400 | 3744600 | 7403500 | 6615900 | 12813000 | 4918300 |
| P52209-2;P52209;K7EMN2;K7EM49;K7EPF6 | 6-phosphogluconate dehydrogenase, decarboxylating | PGD | 108220 | 37993 | 312490 | 0 | 0 | 30683 |
| P52272-2;P52272;A0A087X0X3;M0R019;M0R2T0;M0R0N3;M0QYQ7;M0QZM1;M0R0Y6;M0R2I7;M0QYL3 | Heterogeneous nuclear ribonucleoprotein M | HNRNPM | 7697400 | 4782400 | 6590700 | 5996200 | 8756200 | 5558400 |
| P52292;J3QLL0;J3KS65;J3QL07 | Importin subunit alpha-1 | KPNA2 | 11869000 | 8134100 | 16426000 | 1504400 | 3244400 | 1742100 |
| P52294;C9JYI4;C9J352;C9JWD9;F2Z3G4;C9J4U1;O15131 | Importin subunit alpha-5;Importin subunit alpha-5, N-terminally processed | KPNA1 | 5322000 | 2980900 | 5356700 | 2740300 | 3092200 | 1416300 |
| P52298-3;P52298;C9JQX9;B3KSB0;P52298-2;F8WE41 | Nuclear cap-binding protein subunit 2 | NCBP2 | 12745000 | 9411500 | 17308000 | 2841800 | 7194600 | 3042700 |
| P52306-6;P52306-2;P52306-3;P52306-4;P52306;P52306-5 | Rap1 GTPase-GDP dissociation stimulator 1 | RAP1GDS1 | 0 | 46807 | 246450 | 47406 | 0 | 0 |
| P52597 | Heterogeneous nuclear ribonucleoprotein F;Heterogeneous nuclear ribonucleoprotein F, N-terminally processed | HNRNPF | 33183000 | 22833000 | 44347000 | 10419000 | 26839000 | 7720700 |
| P52701-4;P52701-3;P52701;A0A087WWJ1;P52701-2;A0A087WYT6 | DNA mismatch repair protein Msh6 | MSH6 | 3573900 | 1820200 | 5114800 | 831090 | 2177000 | 777210 |
| P52907 | F-actin-capping protein subunit alpha-1 | CAPZA1 | 3024000 | 1078600 | 2534400 | 282800 | 506340 | 433290 |
| P53396-2;P53396;P53396-3;K7ESG8;K7EIE7 | ATP-citrate synthase | ACLY | 3444500 | 2175400 | 4619300 | 1404700 | 2877800 | 955470 |
| P53582;H0Y9L0;D6RF24;H0Y955 | Methionine aminopeptidase 1 | METAP1 | 1079600 | 481480 | 1066500 | 0 | 551960 | 0 |
| P53618;E9PP73 | Coatomer subunit beta | COPB1 | 388580 | 89538 | 367480 | 0 | 92402 | 45830 |
| P53621;P53621-2 | Coatomer subunit alpha;Xenin;Proxenin | COPA | 1069600 | 468710 | 1432900 | 101730 | 214420 | 84695 |
| P54136;P54136-2;E5RJM9;E5RH09 | Arginine--tRNA ligase, cytoplasmic | RARS | 5823200 | 4024800 | 9256900 | 5407600 | 10654000 | 4465900 |
| P54577 | Tyrosine--tRNA ligase, cytoplasmic;Tyrosine--tRNA ligase, cytoplasmic, N-terminally processed | YARS | 3842800 | 2303300 | 5054200 | 6510000 | 12779000 | 7717900 |
| P54578-2;P54578;A6NJA2;B7Z4N8;J3QQT6;J3KS55;H0YA47 | Ubiquitin carboxyl-terminal hydrolase 14;Ubiquitin carboxyl-terminal hydrolase | USP14 | 60957000 | 25881000 | 30626000 | 5621100 | 7031400 | 3159400 |
| P54619-2;P54619;P54619-3;H0YHF8;F8VYY9;H0YIC9;E9PGP6;Q9UGJ0-2;Q9UGJ0-3;Q9UGJ0;F8VSH3;F8W046;F8VRY2;F8VSL2;F8VVA3;F8VZX1 | 5-AMP-activated protein kinase subunit gamma-1 | PRKAG1 | 3728400 | 2482100 | 6926900 | 747520 | 265910 | 101250 |
| P54652 | Heat shock-related 70 kDa protein 2 | HSPA2 | 0 | 0 | 0 | 0 | 0 | 0 |
| P54725-2;P54725-3;P54725;K7ENJ0;K7ELW1;K7EQ16 | UV excision repair protein RAD23 homolog A | RAD23A | 12518000 | 3659400 | 7207200 | 403300 | 224380 | 177370 |
| P54727;P54727-2;Q5W0S5;Q5W0S4;H0Y579 | UV excision repair protein RAD23 homolog B | RAD23B | 316910000 | 144420000 | 225550000 | 12828000 | 14800000 | 8444000 |
| P55010;H0YK11;H0YMJ8;H0YM54;H0YLZ1;H0YN40;H0YK29 | Eukaryotic translation initiation factor 5 | EIF5 | 2101800 | 635800 | 1162600 | 426300 | 667110 | 325240 |
| P55036;A6PVX3;H0Y3Y9;H0Y561 | 26S proteasome non-ATPase regulatory subunit 4 | PSMD4 | 387340000 | 163990000 | 168190000 | 51591000 | 30772000 | 30304000 |
| P55060-3;P55060;P55060-4 | Exportin-2 | CSE1L | 6445600 | 1802200 | 2130700 | 658470 | 1260800 | 489910 |
| P55072 | Transitional endoplasmic reticulum ATPase | VCP | 93575000 | 39708000 | 56528000 | 13698000 | 13492000 | 9570800 |
| P55145 | Mesencephalic astrocyte-derived neurotrophic factor | MANF | 1863700 | 1415200 | 1565700 | 158100 | 619810 | 310150 |
| P55769;B1AHD1 | NHP2-like protein 1;NHP2-like protein 1, N-terminally processed | NHP2L1 | 3055100 | 3197700 | 7516600 | 2664800 | 6178600 | 4277000 |
| P55884;P55884-2;C9JZG1;C9JQN7 | Eukaryotic translation initiation factor 3 subunit B | EIF3B | 10801000 | 6167600 | 18615000 | 11408000 | 23100000 | 15250000 |
| P55957;P55957-2 | BH3-interacting domain death agonist;BH3-interacting domain death agonist p15;BH3-interacting domain death agonist p13;BH3-interacting domain death agonist p11 | BID | 350650 | 0 | 338720 | 176590 | 232360 | 296770 |
| P56182;D6RE82 | Ribosomal RNA processing protein 1 homolog A | RRP1 | 266200 | 193910 | 357510 | 1073800 | 1800800 | 1118600 |
| P56192;H0YIP0;P56192-2;H0YI94;H0YHV5;H0YHL6;H0YI27;F5H2V6 | Methionine--tRNA ligase, cytoplasmic | MARS | 4516000 | 2790600 | 6006600 | 3509200 | 6948200 | 3443100 |
| P56537;P56537-2;B7ZBH1;F8WD20;F8WDS6 | Eukaryotic translation initiation factor 6 | EIF6 | 12249000 | 7816800 | 15248000 | 7782000 | 27943000 | 10371000 |
| P57735 | Ras-related protein Rab-25 | RAB25 | 4904000 | 2420700 | 1153300 | 2405600 | 2171600 | 1856700 |
| P57737-2;P57737-4;P57737;P57737-3;E7EP81;I3L167 | Coronin-7;Coronin | CORO7;CORO7-PAM16 | 319410 | 181420 | 575070 | 0 | 222040 | 0 |
| P57772;C9J8T0;P57772-2 | Selenocysteine-specific elongation factor | EEFSEC | 761120 | 516860 | 1265700 | 674220 | 2030800 | 1140400 |
| P59998-2;F8WDD7;H7C0A3;F8WCF6;C9JSD3;H3BLT7;H0Y5E3;J3KQB2;P59998;P59998-3;Q9Y4R7-6;Q9Y4R7-2;Q9Y4R7-5;Q9Y4R7 | Actin-related protein 2/3 complex subunit 4;Tubulin monoglycylase TTLL3 | ARPC4;ARPC4-TTLL3;TTLL3 | 114600 | 141730 | 130350 | 0 | 172920 | 69444 |
| P60174-1;P60174;P60174-4;U3KQF3;U3KPZ0 | Triosephosphate isomerase | TPI1 | 5120500 | 2414400 | 4474600 | 948920 | 662740 | 622010 |
| P60228;H0YBR5;E5RIT4;E5RHS5;E5RIP5;E5RJ25;E5RII3 | Eukaryotic translation initiation factor 3 subunit E | EIF3E | 33424000 | 17994000 | 37956000 | 34166000 | 57306000 | 28908000 |
| P60709;P63261;Q5T8M7;I3L3I0;I3L1U9;G5E9R0;E7EVS6;C9JTX5;C9JZR7;K7EM38;I3L3R2;J3KT65;I3L4N8;P63267-2;Q6S8J3;C9JUM1;A5A3E0;P0CG38;Q9BYX7;P0CG39;F8WCH0 | Actin, cytoplasmic 1;Actin, cytoplasmic 1, N-terminally processed;Actin, cytoplasmic 2;Actin, cytoplasmic 2, N-terminally processed | ACTB;ACTG1 | 2304800000 | 995990000 | 1807600000 | 45403000 | 52022000 | 25189000 |
| P60842;J3KT12;P60842-2;J3KTB5;J3QS69;J3QL43;J3QR64;J3KSZ0;J3KTN0;J3KS25 | Eukaryotic initiation factor 4A-I | EIF4A1 | 93229000 | 51163000 | 112450000 | 56029000 | 111770000 | 66062000 |
| P60866;P60866-2;E5RJX2;E5RIP1;G3XAN0 | 40S ribosomal protein S20 | RPS20 | 407350000 | 192940000 | 428160000 | 148000000 | 334850000 | 111160000 |
| P60953;P60953-1;F8WET9;B1AH79;Q5JYX0;G3V4H1;B1AH78;G3V476;P84095;Q9H4E5-2;P15153;P17081;Q9H4E5 | Cell division control protein 42 homolog | CDC42 | 1743900 | 1105100 | 442760 | 0 | 0 | 0 |
| P60981;F6RFD5;P60981-2 | Destrin | DSTN | 54049000 | 21590000 | 29699000 | 0 | 176020 | 0 |
| P61081;M0QX69 | NEDD8-conjugating enzyme Ubc12 | UBE2M | 3069100 | 1207900 | 3179400 | 927990 | 1185700 | 401570 |
| P61088;F8VQQ8;F8VSD4;F8VV71;Q5JXB2;F8VZ29 | Ubiquitin-conjugating enzyme E2 N;Putative ubiquitin-conjugating enzyme E2 N-like | UBE2N;UBE2NL | 5000100 | 2444800 | 4879300 | 2885900 | 2908000 | 1316500 |
| P61106;X6RFL8 | Ras-related protein Rab-14 | RAB14 | 363300 | 507010 | 82972 | 1060900 | 0 | 0 |
| P61158;F8WEW2;F8WE84;B4DXW1 | Actin-related protein 3 | ACTR3 | 1702200 | 424420 | 509340 | 0 | 0 | 0 |
| P61163;R4GMT0;P42025 | Alpha-centractin;Beta-centractin | ACTR1A;ACTR1B | 6339500 | 1604400 | 3723300 | 847450 | 608620 | 522130 |
| P61221;D6R9I9;D6RGF4;H0Y990 | ATP-binding cassette sub-family E member 1 | ABCE1 | 6140800 | 3331300 | 5285900 | 3429200 | 9582800 | 4534900 |
| P61247;H0Y9Y4;D6RG13;D6RAT0;E9PFI5;H0Y8L7;D6RB09;D6RI02;D6RAS7;D6RED7;D6RGE0 | 40S ribosomal protein S3a | RPS3A | 401990000 | 221550000 | 465800000 | 423910000 | 672090000 | 421740000 |
| P61254;J3KTJ8;J3QRI7;J3QQQ9;J3QQV1;J3QRC4;E5RIT6;J3KSS0;E5RHH1;J3KS10 | 60S ribosomal protein L26 | RPL26;RPL26L1 | 233280000 | 179390000 | 335030000 | 307850000 | 618690000 | 315190000 |
| P61289;P61289-2;K9J957;P61289-3;K7ESG5;B3KQ25;A0A087WTV2;K7ENH2;K7EPX6;K7EKR3;K7EMD0;B7Z8D3;K7EJF8 | Proteasome activator complex subunit 3 | PSME3 | 314630000 | 234150000 | 366370000 | 191670000 | 121830000 | 85685000 |
| P61313;E7EQV9;E7EX53;E7ENU7;P61313-2;E7ERA2 | 60S ribosomal protein L15;Ribosomal protein L15 | RPL15 | 144630000 | 90711000 | 205700000 | 139560000 | 258430000 | 126670000 |
| P61353;K7EQQ9;K7ELC7;K7ERY7 | 60S ribosomal protein L27 | RPL27 | 165020000 | 156220000 | 333680000 | 82119000 | 343420000 | 179600000 |
| P61764;P61764-2 | Syntaxin-binding protein 1 | STXBP1 | 119920 | 25711 | 45703 | 0 | 0 | 0 |
| P61956-2;P61956;Q6EEV6 | Small ubiquitin-related modifier 2;Small ubiquitin-related modifier 4 | SUMO2;SUMO4 | 692800000 | 217970000 | 250690000 | 166320000 | 198070000 | 140450000 |
| P61962;A0A087WWI6;P61962-2 | DDB1- and CUL4-associated factor 7 | DCAF7 | 19470000 | 11258000 | 8703200 | 1100900 | 1458700 | 93749 |
| P61978-3;P61978-2;P61978;Q5T6W5;Q5T6W2;Q5T6W1;S4R457;S4R359 | Heterogeneous nuclear ribonucleoprotein K | HNRNPK | 31802000 | 14987000 | 40400000 | 18116000 | 45494000 | 22149000 |
| P61981 | 14-3-3 protein gamma;14-3-3 protein gamma, N-terminally processed | YWHAG | 25650000 | 16518000 | 28743000 | 6312000 | 5551500 | 5716600 |
| P62081;B5MCP9 | 40S ribosomal protein S7 | RPS7 | 333450000 | 217500000 | 571840000 | 310020000 | 650580000 | 356910000 |
| P62136;P62136-2;H0Y3Y6;E7ETD8;F5H1L6;F5H037;E9PMD7;P62136-3 | Serine/threonine-protein phosphatase PP1-alpha catalytic subunit;Serine/threonine-protein phosphatase | PPP1CA;PPP1CB | 489700 | 218010 | 1115000 | 536220 | 0 | 216290 |
| P62140;C9J9S3;C9JP48 | Serine/threonine-protein phosphatase PP1-beta catalytic subunit;Serine/threonine-protein phosphatase | PPP1CB | 3381000 | 888690 | 1017800 | 240770 | 586750 | 301400 |
| P62191 | 26S protease regulatory subunit 4 | PSMC1 | 651770000 | 343820000 | 473090000 | 76380000 | 72361000 | 48449000 |
| P62195;P62195-2;J3QSA9;J3QQM1;J3KRP2;J3QLH6;J3QRW1;J3QSE0;J3QRR3;J3KTQ9;Q8NB90-3;Q8NB90-2;Q8NB90 | 26S protease regulatory subunit 8 | PSMC5 | 903540000 | 496880000 | 623360000 | 101690000 | 105380000 | 58013000 |
| Q5JR95;P62241 | 40S ribosomal protein S8 | RPS8 | 215750000 | 170570000 | 333030000 | 191260000 | 424080000 | 239190000 |
| P62258;P62258-2;B4DJF2;I3L3T1;K7EM20;I3L0W5;K7EIT4 | 14-3-3 protein epsilon | YWHAE | 483410000 | 217320000 | 268210000 | 64821000 | 86678000 | 55033000 |
| P62263;H0YB22;E5RH77 | 40S ribosomal protein S14 | RPS14 | 552690000 | 307280000 | 692500000 | 757660000 | 1204900000 | 679950000 |
| P62269;J3JS69 | 40S ribosomal protein S18 | RPS18 | 359850000 | 227900000 | 511170000 | 446540000 | 710610000 | 377640000 |
| P62277;E9PS50;J3KMX5 | 40S ribosomal protein S13 | RPS13 | 170240000 | 128900000 | 213650000 | 162350000 | 246350000 | 182760000 |
| P62310;REV__Q9Y4P9 | U6 snRNA-associated Sm-like protein LSm3 | LSM3 | 42071000 | 17969000 | 47756000 | 13496000 | 24036000 | 9186500 |
| P62312 | U6 snRNA-associated Sm-like protein LSm6 | LSM6 | 10043000 | 4745100 | 10784000 | 5617400 | 9973100 | 4019500 |
| P62316;P62316-2;K7ERG4;K7EJB5 | Small nuclear ribonucleoprotein Sm D2 | SNRPD2 | 10855000 | 10419000 | 16785000 | 6074100 | 11301000 | 3966200 |
| P62318-2;P62318;H3BT13 | Small nuclear ribonucleoprotein Sm D3 | SNRPD3 | 31289000 | 22331000 | 33141000 | 18215000 | 34918000 | 17790000 |
| P62424;Q5T8U3;Q5T8U2 | 60S ribosomal protein L7a | RPL7A | 266290000 | 155940000 | 334590000 | 216350000 | 472340000 | 263540000 |
| P62487;E9PIU7;H0YEE4;E9PKH3 | DNA-directed RNA polymerase II subunit RPB7 | POLR2G | 1321500 | 1315100 | 2639800 | 0 | 326850 | 0 |
| P62633-7;P62633-3;P62633-2;P62633;P62633-8;P62633-5;P62633-4;P62633-6 | Cellular nucleic acid-binding protein | CNBP | 1118400 | 220770 | 1460200 | 0 | 0 | 191140 |
| P62701;Q8TD47;A6NH36;C9JEH7;P22090 | 40S ribosomal protein S4, X isoform | RPS4X | 364320000 | 217090000 | 499070000 | 453230000 | 683780000 | 334300000 |
| P63267;P68133;P68032;P62736;Q5T8M8;A6NL76;F8WB63;B8ZZJ2;C9JFL5;F6UVQ4;F6QUT6 | Actin, gamma-enteric smooth muscle;Actin, alpha skeletal muscle;Actin, alpha cardiac muscle 1;Actin, aortic smooth muscle | ACTG2;ACTA1;ACTC1;ACTA2 | 2115600 | 2961300 | 3345200 | 200670 | 114040 | 28872 |
| P62753;A2A3R7 | 40S ribosomal protein S6 | RPS6 | 224810000 | 151440000 | 348220000 | 180230000 | 456900000 | 181990000 |
| P62805 | Histone H4 | HIST1H4A | 20977000 | 14916000 | 31420000 | 10580000 | 15594000 | 9720400 |
| P62829;C9JD32;B9ZVP7;J3KT29;J3KTJ3 | 60S ribosomal protein L23 | RPL23 | 87768000 | 53026000 | 62884000 | 133570000 | 256860000 | 148790000 |
| P62851 | 40S ribosomal protein S25 | RPS25 | 532980000 | 342860000 | 685750000 | 536780000 | 924620000 | 504230000 |
| P62854;Q5JNZ5 | 40S ribosomal protein S26;Putative 40S ribosomal protein S26-like 1 | RPS26;RPS26P11 | 442530000 | 170830000 | 343130000 | 628090000 | 673080000 | 257350000 |
| P62857 | 40S ribosomal protein S28 | RPS28 | 287020000 | 268060000 | 557970000 | 1047900000 | 545130000 | 639680000 |
| P62873-2;P62873;C9JD14;B1AKQ8;C9JZN1;F6X3N5;F6UT28;F5H8J8;F5H100;C9JIS1;B3KVK2;C9JXA5;F5H0S8;E7EP32;E9PCP0;P16520-2;Q9HAV0;P62879;P16520 | Guanine nucleotide-binding protein G(I)/G(S)/G(T) subunit beta-1;Guanine nucleotide-binding protein G(I)/G(S)/G(T) subunit beta-3;Guanine nucleotide-binding protein subunit beta-4;Guanine nucleotide-binding protein G(I)/G(S)/G(T) subunit beta-2 | GNB1;GNB4;GNB2;GNB3 | 2002900 | 906920 | 2535100 | 0 | 399210 | 204460 |
| P62877 | E3 ubiquitin-protein ligase RBX1;E3 ubiquitin-protein ligase RBX1, N-terminally processed | RBX1 | 21725000 | 9775100 | 8248000 | 5458400 | 4733700 | 0 |
| P62888;E5RI99;E5RJH3 | 60S ribosomal protein L30 | RPL30 | 157680000 | 92020000 | 187350000 | 219600000 | 390930000 | 224510000 |
| P62899;B8ZZK4;H7C2W9;C9JU56;B7Z4E3;B7Z4C8;P62899-3;P62899-2 | 60S ribosomal protein L31 | RPL31 | 404600000 | 187140000 | 441150000 | 344610000 | 623880000 | 321690000 |
| P62906 | 60S ribosomal protein L10a | RPL10A | 158640000 | 129890000 | 237590000 | 222490000 | 357550000 | 181850000 |
| P62913;Q5VVC8;Q5VVC9 | 60S ribosomal protein L11 | RPL11 | 241970000 | 130840000 | 313880000 | 218340000 | 419060000 | 209670000 |
| P62917;E9PKZ0;E9PKU4;G3V1A1;E9PP36 | 60S ribosomal protein L8 | RPL8 | 238220000 | 129290000 | 333190000 | 243600000 | 483400000 | 214050000 |
| P62937;F8WE65;C9J5S7;P62937-2;A0A075B767;A0A075B759;Q9Y536;F5H284;A2BFH1 | Peptidyl-prolyl cis-trans isomerase A;Peptidyl-prolyl cis-trans isomerase A, N-terminally processed;Peptidyl-prolyl cis-trans isomerase | PPIA | 6635500 | 5173700 | 15014000 | 1288400 | 5276100 | 1338100 |
| P62979;J3QS39;J3QTR3;F5H6Q2;F5GYU3;F5H2Z3;F5H265;B4DV12;F5H388;F5H747;F5GXK7;J3QKN0;Q5PY61;Q96C32;P62987;P0CG47;P0CG48;M0R1V7;A0A087WV77;F5GZ39;J3QSA3 | Ubiquitin-40S ribosomal protein S27a;Ubiquitin;40S ribosomal protein S27a;Ubiquitin-60S ribosomal protein L40;Ubiquitin;60S ribosomal protein L40;Polyubiquitin-B;Ubiquitin;Polyubiquitin-C;Ubiquitin | RPS27A;UBB;UBC;UBA52 | 1146700000 | 768780000 | 940910000 | 126210000 | 157700000 | 111300000 |
| P62993;J3KT38;P62993-2;J3QLF6 | Growth factor receptor-bound protein 2 | GRB2 | 6627700 | 1839100 | 3594600 | 79236 | 294940 | 156830 |
| P63010-3;P63010;P63010-2;A0A087X253;A0A087WU93;A0A087WZQ6;A0A087WYD1;K7ERB2;K7EN71 | AP-2 complex subunit beta | AP2B1 | 2518800 | 889550 | 2490600 | 529530 | 759240 | 420440 |
| P63104;E7EX29;B0AZS6;P63104-2;B7Z2E6;H0YB80;E7ESK7;E9PD24;E7EVZ2;E5RIR4;E5RGE1;REV__P53355-4;REV__P53355;REV__P53355-3 | 14-3-3 protein zeta/delta | YWHAZ | 151820000 | 56546000 | 107410000 | 30203000 | 33631000 | 18679000 |
| P63151;P63151-2;E5RIY1;Q9Y2T4-3;Q9Y2T4-4;Q9Y2T4-2;Q9Y2T4;H0YB06;E5RFR9;Q00005-6;Q00005;Q00005-2;Q00005-3;Q66LE6;Q00005-4;Q00005-5;Q00005-7 | Serine/threonine-protein phosphatase 2A 55 kDa regulatory subunit B alpha isoform;Serine/threonine-protein phosphatase 2A 55 kDa regulatory subunit B gamma isoform | PPP2R2A;PPP2R2C | 1150500 | 474220 | 1068200 | 180660 | 168450 | 229420 |
| Q96FJ2;P63167 | Dynein light chain 2, cytoplasmic;Dynein light chain 1, cytoplasmic | DYNLL2;DYNLL1 | 25656000 | 761970 | 9149900 | 0 | 1221600 | 0 |
| P63173;J3KT73;J3QL01;J3KSP2 | 60S ribosomal protein L38 | RPL38 | 79474000 | 61632000 | 141370000 | 75737000 | 228000000 | 133720000 |
| Q8WVC2;P63220;Q9BYK1;Q9BYK2 | 40S ribosomal protein S21 | RPS21 | 61953000 | 28242000 | 58491000 | 77029000 | 76816000 | 32091000 |
| P63244;J3KPE3;H0YAF8;H0Y8W2;D6RFX4;D6RHH4;D6R9Z1;H0YAM7;D6RAC2;D6R9L0;D6RFZ9;D6REE5;D6RAU2;E9PD14;D6RBD0;D6RGK8;D6RHJ5;D6R909;H0Y8R5;D6RDI0 | Guanine nucleotide-binding protein subunit beta-2-like 1;Guanine nucleotide-binding protein subunit beta-2-like 1, N-terminally processed | GNB2L1 | 217730000 | 141770000 | 266000000 | 217210000 | 359540000 | 214260000 |
| P67775-2;P67775;E5RI56;E5RFI3;E5RHP4;E7ESG8;E5RHC1;P62714 | Serine/threonine-protein phosphatase 2A catalytic subunit alpha isoform;Serine/threonine-protein phosphatase;Serine/threonine-protein phosphatase 2A catalytic subunit beta isoform | PPP2CA;PPP2CB | 1552600 | 1239100 | 3463900 | 759180 | 1530900 | 569300 |
| P67809;A0A087X1S2;H0Y449;C9J5V9;Q9Y2T7 | Nuclease-sensitive element-binding protein 1 | YBX1 | 140000000 | 65374000 | 189170000 | 136160000 | 198490000 | 122070000 |
| Q5SRQ3;Q5SRQ6;P67870;N0E472 | Casein kinase II subunit beta | CSNK2B-LY6G5B-1181;CSNK2B;CSNK2B-LY6G5B--991 | 9453900 | 4999000 | 7944400 | 5671000 | 9161800 | 3886900 |
| P67936;P67936-2;K7ENT6;K7EMU5;K7EPV9;K7EPB9;K7ERG3 | Tropomyosin alpha-4 chain | TPM4 | 36608000 | 24783000 | 31900000 | 182090 | 270370 | 0 |
| Q5VTE0;P68104;A0A087WVQ9;P68104-2;A0A087WV01;A6PW80;Q5JR01 | Putative elongation factor 1-alpha-like 3;Elongation factor 1-alpha 1;Elongation factor 1-alpha | EEF1A1P5;EEF1A1 | 428920000 | 244870000 | 513560000 | 135100000 | 316070000 | 138280000 |
| P68363;Q71U36-2;Q71U36;F5H5D3;Q9BQE3;P68363-2;P68366-2;P68366;Q13748;F8VQQ4;Q13748-2;F8VVB9;Q6PEY2;F8VRZ4;F8VS66;F8VWV9;F8VX09;C9JEV8;C9JQ00;C9JJQ8;C9JDL2;C9JDS9;F8VRK0;F8VXZ7 | Tubulin alpha-1B chain;Tubulin alpha-1A chain;Tubulin alpha-1C chain;Tubulin alpha-4A chain;Tubulin alpha-3C/D chain;Tubulin alpha-3E chain | TUBA1B;TUBA1A;TUBA1C;TUBA4A;TUBA3C;TUBA3E | 203970000 | 78461000 | 107460000 | 193970000 | 94148000 | 45377000 |
| P68371;P04350;Q13509-2;Q3ZCM7;K7ESM5;M0R2D3;M0R0X0;M0QY85;M0QY37;M0QX14;M0R042;M0QYM7;M0R2T4;A0A075B724 | Tubulin beta-4B chain;Tubulin beta-4A chain | TUBB4B;TUBB4A | 47326000 | 25916000 | 54136000 | 33273000 | 38677000 | 20765000 |
| P78318 | Immunoglobulin-binding protein 1 | IGBP1 | 373030 | 117260 | 0 | 46410 | 0 | 0 |
| P78345;Q5VUC3;Q5VUC9 | Ribonuclease P protein subunit p38 | RPP38 | 714880 | 547600 | 1233100 | 445380 | 1516300 | 766520 |
| P78346;P78346-2;Q5VU10;Q5VU11 | Ribonuclease P protein subunit p30 | RPP30 | 3324100 | 984900 | 2567800 | 4420200 | 5127200 | 2023000 |
| P78347-2;P78347-4;P78347-3;P78347;P78347-5;C9J6M0;A0A087WW90;Q86UP8-3;Q86UP8;Q6EKJ0 | General transcription factor II-I | GTF2I | 6635900 | 3194600 | 7055000 | 2224300 | 2974700 | 1110400 |
| P78362;P78362-2;H7C5L6 | SRSF protein kinase 2;SRSF protein kinase 2 N-terminal;SRSF protein kinase 2 C-terminal | SRPK2 | 1112800 | 678980 | 1244800 | 1487900 | 1917700 | 1160500 |
| P78371-2;P78371;F5GWF6;F8VQ14 | T-complex protein 1 subunit beta | CCT2 | 10735000 | 4565700 | 7969300 | 7862200 | 5971400 | 3567600 |
| P78527;P78527-2;F5GX40;H0YG84 | DNA-dependent protein kinase catalytic subunit | PRKDC | 5014700 | 2640600 | 5421100 | 2989900 | 5950000 | 3828700 |
| P80723-2;P80723 | Brain acid soluble protein 1 | BASP1 | 851600 | 875590 | 2950200 | 138840 | 430840 | 326370 |
| P81605;P81605-2 | Dermcidin;Survival-promoting peptide;DCD-1 | DCD | 11638000 | 7134000 | 13691000 | 15839000 | 7472400 | 8377100 |
| P82979;F8VZQ9;H0YHG0;F8VS12 | SAP domain-containing ribonucleoprotein | SARNP | 884730 | 241160 | 1083300 | 1647900 | 4241700 | 1611300 |
| P84090;G3V279 | Enhancer of rudimentary homolog | ERH | 40994000 | 24395000 | 39776000 | 18072000 | 27365000 | 13895000 |
| P98175-4;P98175-3;P98175-2;P98175 | RNA-binding protein 10 | RBM10 | 131370 | 74084 | 109480 | 102760 | 144610 | 65214 |

| | | | | | | | | |
|---|---|---|---|---|---|---|---|---|
| Q00341;Q00341-2;H0Y394;H7C0A4;H7C2D1;C9JQ82;H7BZC3;C9JMQ6;C9JHS9;C9J739;H7C3D0 | Vigilin | HDLBP | 5554300 | 3119300 | 8372700 | 5387800 | 14821000 | 6000800 |
| Q00577 | Transcriptional activator protein Pur-alpha | PURA | 3711300 | 2931800 | 11577000 | 3199000 | 13831000 | 4502800 |
| Q00688;G3V5F2 | Peptidyl-prolyl cis-trans isomerase FKBP3 | FKBP3 | 14469000 | 9207500 | 24893000 | 28634000 | 55804000 | 28607000 |
| Q00765;E2QRG8 | Receptor expression-enhancing protein 5;Receptor expression-enhancing protein | REEP5 | 4148900 | 3841900 | 3548500 | 371680 | 470060 | 262720 |
| Q00839-2;Q00839;Q5RI18 | Heterogeneous nuclear ribonucleoprotein U | HNRNPU | 15806000 | 8698600 | 30895000 | 25328000 | 61093000 | 25471000 |
| Q01085;Q01085-2;A6NKZ9;E7ETJ9;E7ETC0;E5RGV5;H7BY49;E5RG67;C9JTN7;F8W8I6;Q96B58;P31483-2;P31483 | Nucleolysin TIAR;Nucleolysin TIA-1 isoform p40 | TIAL1;TIA1 | 967080 | 351660 | 652750 | 242780 | 520840 | 42165 |
| Q01105-3;Q01105-4;Q01105-2;Q01105;A0A087X027;P0DME0 | Protein SET;Protein SETSIP | SET;SETSIP | 55951000 | 23507000 | 49344000 | 31027000 | 30119000 | 15294000 |
| Q01469;I6L8B7 | Fatty acid-binding protein, epidermal | FABP5 | 1039500 | 4285500 | 952690 | 746640 | 1228000 | 556980 |
| Q01518-2;Q01518;Q5T0R7;Q5T0R6;Q5T0R5;Q5T0R4;Q5T0R3;Q5T0R2;Q5T0R1;Q5T0R9;Q5T0R8;Q5T0S3;B7Z385;A0A087X0J3;A0A087WZ15;E9PDI2;P40123-3;P40123-2;P40123 | Adenylyl cyclase-associated protein 1 | CAP1 | 1824500 | 570950 | 874440 | 16140 | 37059 | 0 |
| Q01780;Q01780-2;K7EJ37 | Exosome component 10 | EXOSC10 | 5144500 | 2324300 | 5121700 | 8082900 | 13099000 | 5915200 |
| Q01813;Q01813-2;B1APP6;B1APP8;H0Y757;V9GY25;H0Y3Y3 | ATP-dependent 6-phosphofructokinase, platelet type | PFKP | 6302200 | 2860600 | 4246800 | 0 | 549600 | 31524 |
| Q02790;H0YFG2;F5H1U3 | Peptidyl-prolyl cis-trans isomerase FKBP4;Peptidyl-prolyl cis-trans isomerase FKBP4, N-terminally processed | FKBP4 | 2492000 | 1135700 | 3407900 | 1637200 | 1779400 | 760670 |
| Q02809;Q02809-2 | Procollagen-lysine,2-oxoglutarate 5-dioxygenase 1 | PLOD1 | 637720 | 169520 | 168800 | 0 | 0 | 0 |
| Q02878;F8VZ45;U3KQR5;F8VR69;F8VZA3;F8VWR1;F8VU16 | 60S ribosomal protein L6 | RPL6 | 204460000 | 112530000 | 277200000 | 186280000 | 438560000 | 208230000 |
| Q02880-2;Q02880;E9PCY5 | DNA topoisomerase 2-beta;DNA topoisomerase 2 | TOP2B | 502880 | 108010 | 328790 | 74335 | 200970 | 49858 |
| Q04637-5;Q04637-4;Q04637-3;Q04637;Q04637-9;Q04637-6;C9J6B6;C9JF13 | Eukaryotic translation initiation factor 4 gamma 1 | EIF4G1 | 4726700 | 2556200 | 8653100 | 4941600 | 10060000 | 4545400 |
| Q04837;C9K0U8;E7EUY5 | Single-stranded DNA-binding protein, mitochondrial | SSBP1 | 14439000 | 20103000 | 19569000 | 15587000 | 23823000 | 12870000 |
| Q04917;A2IDB2;F8WEB6;A2IDB1;REV__C9JDT0;REV__C9JQI2;REV__Q7Z5H3-4;REV__D6R9V6;REV__A6NJ38;REV__A6NHM7;REV__Q7Z5H3-3;REV__P13521;REV__Q7Z5H3;REV__Q7Z5H3-5;REV__Q7Z5H3-2 | 14-3-3 protein eta | YWHAH | 44834000 | 17672000 | 29451000 | 8928300 | 10129000 | 5202000 |
| Q04941 | Proteolipid protein 2 | PLP2 | 1239900 | 202390 | 1026700 | 0 | 0 | 0 |
| Q05048;A3KFI9 | Cleavage stimulation factor subunit 1 | CSTF1 | 822200 | 98705 | 1598100 | 859930 | 1753500 | 1737600 |
| Q05086-3;Q05086;Q05086-2;S4R306 | Ubiquitin-protein ligase E3A | UBE3A | 12278000 | 6725200 | 9832100 | 560570 | 888430 | 499930 |
| Q05193-5;Q05193-3;Q05193-4;Q05193-2;Q05193 | Dynamin-1 | DNM1 | 168370 | 33015 | 62336 | 0 | 22984 | 34058 |
| Q05639 | Elongation factor 1-alpha 2 | EEF1A2 | 6346400 | 2055500 | 6779200 | 2793700 | 7347700 | 2829000 |
| Q06265;Q06265-2;D6RIY6;D6R905;Q06265-3;Q06265-4;D6RA17;D6RAP4;H0Y9L5 | Exosome complex component RRP45 | EXOSC9 | 5783100 | 4029700 | 6989300 | 11700000 | 12092000 | 4512200 |
| Q06323;Q06323-3;Q06323-2;H0YKK6;H0YLU2 | Proteasome activator complex subunit 1 | PSME1 | 90551000 | 75197000 | 108570000 | 50154000 | 46737000 | 32576000 |
| Q06787-8;Q06787-5;Q06787-9;Q06787-3;Q06787;R9WNI0;A8MQB8;G3V0J0;Q06787-6;Q06787-2;Q06787-4;Q06787-7;A0A087WXI3;G8JLE9;Q8IXW7;A0A087WY29 | Fragile X mental retardation protein 1 | FMR1 | 714930 | 483860 | 1775200 | 164010 | 2363400 | 1218000 |
| Q06830 | Peroxiredoxin-1 | PRDX1 | 24476000 | 14442000 | 23299000 | 6655700 | 20383000 | 10057000 |
| Q07666-2;Q07666 | KH domain-containing, RNA-binding, signal transduction-associated protein 1 | KHDRBS1 | 2735600 | 1243600 | 1691300 | 919800 | 620020 | 683270 |
| Q08211;Q08211-2;REV__H0YA90;REV__Q7Z6J0-3 | ATP-dependent RNA helicase A | DHX9 | 17544000 | 10129000 | 22404000 | 26877000 | 52376000 | 23850000 |
| Q08380;K7EP36;K7EQT9;K7ES75;K7EN99;K7ERZ6;K7EJD3;K7ESM3;K7EKQ5 | Galectin-3-binding protein | LGALS3BP | 1737600 | 552610 | 685730 | 334550 | 0 | 69572 |
| Q08554-2;Q08554 | Desmocollin-1 | DSC1 | 176060 | 75757 | 302270 | 68334 | 76323 | 0 |
| Q08AM6;A0A087WT26;H3BUU8;H3BQD9 | Protein VAC14 homolog | VAC14 | 563000 | 50474 | 470550 | 56987 | 143460 | 59052 |
| Q08J23;Q08J23-2;Q08J23-3 | tRNA (cytosine(34)-C(5))-methyltransferase | NSUN2 | 1776700 | 1228800 | 3942900 | 5941100 | 8633300 | 3668100 |
| Q09028-3;Q09028;Q09028-2;Q09028-4;E9PNS2;E9PIC4;E9PNS6;E9PC52;C9JAJ9;C9JPP3;H0YF10 | Histone-binding protein RBBP4 | RBBP4 | 10237000 | 6014100 | 7937200 | 3540100 | 3363400 | 1986700 |
| Q09161;X6R941 | Nuclear cap-binding protein subunit 1 | NCBP1 | 8444000 | 3669300 | 5579600 | 2028300 | 2390100 | 1477600 |
| Q0VDF9;H7C2A1;B0YIZ1 | Heat shock 70 kDa protein 14 | HSPA14 | 2817300 | 573100 | 1531600 | 2088600 | 2987500 | 1591200 |
| Q10567-4;Q10567-3;Q10567-2;Q10567;C9J1E7 | AP-1 complex subunit beta-1 | AP1B1 | 3019300 | 1084400 | 2484300 | 250940 | 358260 | 402910 |
| Q10570 | Cleavage and polyadenylation specificity factor subunit 1 | CPSF1 | 76935 | 49355 | 48250 | 7447.7 | 17500 | 12867 |
| Q12788;J3KNP2;A0A087WYP7;H3BN88 | Transducin beta-like protein 3 | TBL3 | 507770 | 326180 | 449480 | 1201500 | 2403600 | 1403600 |
| Q12789-3;Q12789 | General transcription factor 3C polypeptide 1 | GTF3C1 | 47963 | 109710 | 296670 | 42183 | 132770 | 0 |
| Q12792;F8VRG3;F8VS81;Q12792-4;Q12792-3 | Twinfilin-1 | TWF1 | 824190 | 307910 | 676640 | 0 | 0 | 0 |
| Q12874 | Splicing factor 3A subunit 3 | SF3A3 | 1621600 | 1078600 | 1395700 | 791260 | 1024300 | 537520 |
| Q12904;Q12904-2;D6R937 | Aminoacyl tRNA synthase complex-interacting multifunctional protein 1;Endothelial monocyte-activating polypeptide 2 | AIMP1 | 2645000 | 1870800 | 4618100 | 2163700 | 5742200 | 2202500 |
| Q12906-7;Q12906-4;Q12906-6;Q12906;Q12906-5;Q12906-2;Q12906-3;K7EKJ9;K7ER69;K7EQR9;K7EKY0;K7ERM6;K7ENK6;K7EM82;K7EJ09;K7EQ75 | Interleukin enhancer-binding factor 3 | ILF3 | 33575000 | 19176000 | 53725000 | 51031000 | 76588000 | 33030000 |
| Q13033;Q13033-2;G3V3G7;G3V340;H0YJZ4 | Striatin-3 | STRN3 | 1256200 | 106700 | 302420 | 0 | 0 | 0 |
| Q5T8C6;Q7Z651;Q13042-3;Q13042-2;Q13042 | Cell division cycle protein 16 homolog | CDC16 | 561380 | 561830 | 1219300 | 280860 | 645290 | 194740 |
| Q13045-2;Q13045-3;Q13045 | Protein flightless-1 homolog | FLII | 20660 | 27537 | 31533 | 0 | 0 | 0 |
| Q13085-3;Q13085-2;Q13085;Q13085-4;Q59FY4;A0A087WVR6;A0A087X0W4;S4R3S7;O00763-3;O00763-2;O00763 | Acetyl-CoA carboxylase 1;Biotin carboxylase | ACACA | 120320 | 78529 | 106230 | 15339 | 24016 | 30373 |
| Q13112 | Chromatin assembly factor 1 subunit B | CHAF1B | 360310 | 273380 | 409040 | 390750 | 681810 | 220100 |
| Q13131;Q13131-2 | 5-AMP-activated protein kinase catalytic subunit alpha-1 | PRKAA1 | 409360 | 160500 | 262870 | 0 | 0 | 0 |
| Q13151 | Heterogeneous nuclear ribonucleoprotein A0 | HNRNPA0 | 44515000 | 20644000 | 46117000 | 25981000 | 63340000 | 31286000 |
| Q13155;A8MU58;F8W950 | Aminoacyl tRNA synthase complex-interacting multifunctional protein 2 | AIMP2 | 1631900 | 610940 | 1115700 | 1984100 | 3178600 | 1511300 |
| Q13200;Q13200-3;Q13200-2;C9JPC0;H7C2Q3;F8WBS8;P11215;P11215-2 | 26S proteasome non-ATPase regulatory subunit 2 | PSMD2 | 387880000 | 175130000 | 242600000 | 40241000 | 41436000 | 26522000 |
| Q13242;H0YIB4;S4R3G0 | Serine/arginine-rich splicing factor 9 | SRSF9 | 1109100 | 820700 | 1588400 | 281850 | 587080 | 245790 |
| Q13263;Q13263-2;M0R0K9;M0R2I3;M0R3C0 | Transcription intermediary factor 1-beta | TRIM28 | 7009500 | 4663900 | 6432000 | 1887200 | 3311500 | 2221200 |
| Q13283 | Ras GTPase-activating protein-binding protein 1 | G3BP1 | 10584000 | 6878200 | 12844000 | 8156900 | 16481000 | 7982700 |
| Q13310-2;Q13310-3;B1ANR0;Q13310;H0Y5F5;H0YEU6;H0YCC8;B1ANR1;H0YEQ8;H0Y6X6;P0CB38;H0YER0;E5RHG7;Q5JQF8 | Polyadenylate-binding protein 4;Polyadenylate-binding protein | PABPC4 | 4550800 | 3828500 | 7645600 | 4102800 | 8726800 | 5763900 |
| Q13315 | Serine-protein kinase ATM | ATM | 0 | 0 | 0 | 0 | 0 | 0 |
| Q13347;Q5TFK1 | Eukaryotic translation initiation factor 3 subunit I | EIF3I | 23977000 | 14346000 | 33027000 | 25165000 | 39012000 | 23351000 |
| Q13395 | Probable methyltransferase TARBP1 | TARBP1 | 166870 | 81281 | 244360 | 28983 | 48032 | 31653 |
| Q13409-6;Q13409-3;Q13409-7;Q13409-2;Q13409-5;Q13409;E9PGG1;E7EUM4;E7ERR6;E7ERH4;E7ET01;E7ETL8;E7EQU2;E7ESD3;E7EMU4;E7EV09;E7EQL5 | Cytoplasmic dynein 1 intermediate chain 2 | DYNC1I2 | 120710 | 0 | 303160 | 0 | 0 | 0 |
| Q13426-3;Q13426-2;Q13426 | DNA repair protein XRCC4 | XRCC4 | 241870 | 121410 | 160470 | 33897 | 165710 | 0 |
| Q13435;E9PJ04;A0A087WZZ5;E9PJT3;H0YCG1;E9PPJ0;E9PIL8;H0YEX5 | Splicing factor 3B subunit 2 | SF3B2 | 2357600 | 1289100 | 2133300 | 1967600 | 1906100 | 973990 |
| Q13442;F8WBW6 | 28 kDa heat- and acid-stable phosphoprotein | PDAP1 | 13193000 | 11531000 | 13489000 | 20313000 | 49096000 | 18894000 |
| Q13451;Q13451-2 | Peptidyl-prolyl cis-trans isomerase FKBP5;Peptidyl-prolyl cis-trans isomerase FKBP5, N-terminally processed | FKBP5 | 1274500 | 722530 | 1335700 | 1825400 | 2281400 | 1170600 |
| Q13535-2;Q13535-3;Q13535;H0Y8R8;H0Y8Y6;H0Y9K2 | Serine/threonine-protein kinase ATR | ATR | 0 | 15415 | 0 | 0 | 15748 | 0 |
| Q13601-2;Q13601;A0A087WZC7 | KRR1 small subunit processome component homolog | KRR1 | 522690 | 550040 | 1079900 | 2755600 | 5025400 | 2856100 |
| Q13616;Q8NI38-2;REV__P15924;Q8IW00-2;Q8IW00 | Cullin-1 | CUL1 | 14773000 | 7289800 | 10241000 | 5923800 | 5269700 | 2205400 |
| Q5T2B5;Q13617;Q13617-2;Q5T2B7;Q5T2B4 | Cullin-2 | CUL2 | 125470 | 66814 | 140580 | 14699 | 0 | 0 |
| Q13618-2;Q13618;A0A087X1R9;H7C1L6;A0A087WTG3;Q13618-3 | Cullin-3 | CUL3 | 515900 | 102260 | 173490 | 101240 | 0 | 63103 |
| Q13627-3;Q13627-4;Q13627-5;Q9Y463-2;Q9Y463-3;Q9Y463;Q13627-2;Q13627;E7EMI5;M0R2X3;M0R131 | Dual specificity tyrosine-phosphorylation-regulated kinase 1A;Dual specificity tyrosine-phosphorylation-regulated kinase 1B | DYRK1A;DYRK1B | 694510 | 0 | 161710 | 0 | 0 | 0 |
| Q5JXI0;Q5JXH9;Q5JXH8;Q5JXH7;Q5JXI3;Q5JXI2;Q5JXI8;Q13642-3;Q13642-1;Q13642-5;Q13642-4;Q13642 | Four and a half LIM domains protein 1 | FHL1 | 2840800 | 1936900 | 3743200 | 1163500 | 1718800 | 1065200 |
| Q13813;Q13813-3;REV__C9J6B6;REV__C9JF13;REV__Q04637-6;REV__Q04637-7;REV__Q04637-5;REV__E7EX73;REV__Q04637-4;REV__E9PGM1;REV__Q04637-3;REV__E7EUU4;REV__Q04637;REV__Q04637-8;REV__Q04637-9 | Spectrin alpha chain, non-erythrocytic 1 | SPTAN1 | 14453000 | 4925900 | 11495000 | 170420 | 248530 | 242130 |
| Q13823;H0YG10 | Nucleolar GTP-binding protein 2 | GNL2 | 134050 | 135970 | 70392 | 889640 | 1308900 | 372450 |
| Q13838;Q13838-2;F6WLT2;Q5STU3;O00148;F6S4E6;F6TRA5;F6UJC5;H0Y400;K7EQN7;B1Q2N1;O00148-2;K7EN69;H0YCC6;F6QYI9;K7EPJ3;F6R6M7 | Spliceosome RNA helicase DDX39B;ATP-dependent RNA helicase DDX39A | DDX39B;DDX39A | 2350400 | 1372200 | 3102000 | 2461900 | 4981900 | 2904200 |
| Q13885 | Tubulin beta-2A chain | TUBB2A | 0 | 76991 | 122560 | 111890 | 340490 | 220380 |
| Q13895;H7BY94;F8WBL2 | Bystin | BYSL | 12173000 | 6374300 | 10171000 | 7371900 | 8468000 | 3967700 |
| Q14008-2;Q14008;Q14008-3;E9PQH5;H0YCF6;H0YEK7 | Cytoskeleton-associated protein 5 | CKAP5 | 843300 | 488520 | 625580 | 1269800 | 2649300 | 1190200 |
| Q14011;K7ENX8;K7ELV6;K7EJV5;K7ELT6;K7EJV1;K7EMY9;K7EPM4;K7ER40;K7EQR7;D6W5Y5;Q14011-2 | Cold-inducible RNA-binding protein | CIRBP | 2456100 | 1287200 | 2641500 | 2008500 | 3442300 | 1406400 |
| Q14019;H3BT58 | Coactosin-like protein | COTL1 | 23470000 | 8493800 | 14473000 | 0 | 0 | 0 |
| Q14137;Q14137-2;A0A075B729 | Ribosome biogenesis protein BOP1 | BOP1 | 272510 | 139910 | 231470 | 751100 | 1275000 | 493260 |
| Q15424-2;Q15424;Q15424-4;Q15424-3;Q14151 | Scaffold attachment factor B1;Scaffold attachment factor B2 | SAFB;SAFB2 | 339090 | 1302000 | 910270 | 374490 | 344590 | 141660 |
| Q14152;Q14152-2;Q96Q89-5;Q96Q89-4;Q96Q89-3;Q96Q89;Q96Q89-2 | Eukaryotic translation initiation factor 3 subunit A | EIF3A | 19914000 | 8766200 | 20967000 | 19391000 | 39781000 | 16229000 |
| Q14204;H0YJ21 | Cytoplasmic dynein 1 heavy chain 1 | DYNC1H1 | 201560 | 90096 | 238980 | 45969 | 68049 | 32089 |
| Q14232;F5H0D0;Q14232-2 | Translation initiation factor eIF-2B subunit alpha | EIF2B1 | 2609300 | 1077700 | 3167000 | 2906500 | 5113500 | 3165000 |
| Q14240;Q14240-2;E7EQG2;J3KSN7 | Eukaryotic initiation factor 4A-II;Eukaryotic initiation factor 4A-II, N-terminally processed | EIF4A2 | 1210900 | 1502100 | 2972300 | 550370 | 1258100 | 565690 |
| Q14241 | Transcription elongation factor B polypeptide 3 | TCEB3 | 175770 | 24670 | 296880 | 0 | 238880 | 158000 |
| Q14247-3;Q14247;Q14247-2;H7C314;H0YCD9;E9PKG3;E9PP90;H0YEV2 | Src substrate cortactin | CTTN | 16383000 | 4274100 | 11697000 | 668890 | 1272200 | 191730 |
| Q14258 | E3 ubiquitin/ISG15 ligase TRIM25 | TRIM25 | 1449400 | 1250100 | 3266200 | 2056300 | 4587000 | 2328800 |
| Q14315-2;Q14315 | Filamin-C | FLNC | 946550 | 334120 | 357280 | 0 | 22725 | 49961 |
| Q14444-2;Q14444;G3V153 | Caprin-1 | CAPRIN1 | 11923000 | 7132000 | 13710000 | 6527400 | 15655000 | 7450500 |
| Q14566 | DNA replication licensing factor MCM6 | MCM6 | 1078700 | 2094600 | 1603400 | 1493100 | 2530800 | 396850 |
| Q14674;Q14674-2 | Separin | ESPL1 | 386400 | 225610 | 500390 | 251850 | 660160 | 247490 |
| Q14676-2;Q14676 | Mediator of DNA damage checkpoint protein 1 | MDC1 | 34107 | 37571 | 63416 | 0 | 0 | 4360 |
| Q14677-2;Q14677;Q14677-3 | Clathrin interactor 1 | CLINT1 | 337890 | 273690 | 0 | 0 | 0 | 0 |
| Q14683;G8JLG1;H0Y7K8;V9GY57;V9GYN9 | Structural maintenance of chromosomes protein 1A;Structural maintenance of chromosomes protein | SMC1A | 142900000 | 88319000 | 105590000 | 114810000 | 90784000 | 60894000 |
| Q14684;Q14684-2 | Ribosomal RNA processing protein 1 homolog B | RRP1B | 1368400 | 764930 | 2072100 | 931120 | 3355300 | 1846700 |
| Q14690;S4R3Q4 | Protein RRP5 homolog | PDCD11 | 33063 | 5333.8 | 110590 | 84272 | 286740 | 107880 |
| Q14692 | Ribosome biogenesis protein BMS1 homolog | BMS1 | 356220 | 256490 | 87087 | 663180 | 1549400 | 1088300 |
| Q14694;Q14694-3;Q14694-2;H3BQP1;H3BNS8;H3BQC6;H3BNA1;J3KT19 | Ubiquitin carboxyl-terminal hydrolase 10 | USP10 | 454160 | 280360 | 957500 | 309880 | 1589600 | 661980 |
| Q14697;Q14697-2;F5H6X6;E9PKU7;E9PNH1;Q14697-3 | Neutral alpha-glucosidase AB | GANAB | 185570 | 142370 | 401320 | 0 | 159090 | 53906 |
| Q14847;Q14847-2;C9J9W2;Q14847-3;K7ESD6;F6S2S5 | LIM and SH3 domain protein 1 | LASP1 | 18267000 | 5245700 | 9619200 | 0 | 0 | 0 |
| Q14964;C9JB90;C9JU14;J3KR73;H7BYW1;H0YGL6;P20340-4;Q9NRW1-2;Q9NRW1;P20340-2;P20340 | Ras-related protein Rab-39A;Ras-related protein Rab-6A;Ras-related protein Rab-6B | RAB39A;RAB6B;RAB6A | 3052600 | 2612300 | 3410500 | 324950 | 1245000 | 874440 |
| Q14974;J3QR48;J3KTM9;Q14974-2;J3QKQ5 | Importin subunit beta-1 | KPNB1 | 8441500 | 5153800 | 7263700 | 4213000 | 4875600 | 1893900 |
| S4R402;S4R341;S4R349;Q14978;Q14978-3;Q14978-2 | Nucleolar and coiled-body phosphoprotein 1 | NOLC1 | 0 | 3200400 | 3561300 | 0 | 0 | 0 |
| Q14980-4;Q14980-3;Q14980-2;Q14980;A0A087WY61;F5H4J1;H0YFY6 | Nuclear mitotic apparatus protein 1 | NUMA1 | 0 | 13018 | 61707 | 11775 | 0 | 0 |
| Q14997;Q14997-2;Q14997-3;F8WBH5;Q14997-4 | Proteasome activator complex subunit 4 | PSME4 | 76994000 | 47603000 | 65190000 | 39293000 | 31712000 | 22250000 |
| Q14CX7-2;Q14CX7 | N-alpha-acetyltransferase 25, NatB auxiliary subunit | NAA25 | 1014600 | 448790 | 1510300 | 494550 | 1100200 | 491390 |
| Q15008;Q15008-4;Q15008-3;Q15008-2;C9J0E9;C9J7B7;H7C531 | 26S proteasome non-ATPase regulatory subunit 6 | PSMD6 | 515840000 | 261830000 | 419020000 | 64629000 | 48722000 | 34070000 |
| Q15018 | BRISC complex subunit Abro1 | FAM175B | 636870 | 239030 | 381370 | 0 | 0 | 41460 |
| Q15021;E7EN77 | Condensin complex subunit 1 | NCAPD2 | 693760 | 94688 | 423730 | 0 | 11414 | 0 |
| Q15024 | Exosome complex component RRP42 | EXOSC7 | 83118 | 186210 | 405140 | 2184500 | 1137300 | 1550200 |
| Q15029-2;Q15029-3;Q15029;K7EP67;K7EJ74 | 116 kDa U5 small nuclear ribonucleoprotein component | EFTUD2 | 2807000 | 1857500 | 3594600 | 705290 | 1417200 | 597260 |
| Q15046;Q15046-2;H3BVA8;J3KRL2;H3BPV7;H3BRC9 | Lysine--tRNA ligase | KARS | 4745500 | 2918800 | 6476200 | 4887800 | 7432800 | 3005500 |
| Q15050 | Ribosome biogenesis regulatory protein homolog | RRS1 | 0 | 341240 | 626590 | 4742600 | 7157200 | 2589900 |
| Q15056-2;Q15056 | Eukaryotic translation initiation factor 4H | EIF4H | 3349900 | 1926100 | 7405900 | 165510 | 792800 | 103700 |
| Q15084-3;Q15084;Q15084-4;Q15084-5;Q15084-2 | Protein disulfide-isomerase A6 | PDIA6 | 2379000 | 1449700 | 2299700 | 223520 | 434310 | 223590 |
| Q15149;Q15149-7;Q15149-8;Q15149-9;Q15149-5;Q15149-4;Q15149-6;Q15149-3;Q15149-2;H0YDN1;A0A075B730;A0A087X1U6;P58107 | Plectin | PLEC | 237820 | 149840 | 289950 | 107970 | 105200 | 47819 |
| Q15233-2;Q15233;C9IZL7;H7C367;C9JYS8;C9J4X2 | Non-POU domain-containing octamer-binding protein | NONO | 1899800 | 632100 | 4082000 | 2436200 | 7757700 | 1735900 |
| Q15269;C9J544;A0A096LPI6 | Periodic tryptophan protein 2 homolog | PWP2 | 226620 | 146120 | 422560 | 1227200 | 1568300 | 580750 |
| Q15365;F8VTZ0;H3BSP4;C9K0A2;F8WC71;C9IZV9;C9J7A9;C9JSA6;C9J5V4;C9JTY5;C9JZY3;C9J0A4;F6TJN8;P57723-2;P57723 | Poly(rC)-binding protein 1 | PCBP1 | 43077000 | 13959000 | 51881000 | 6289700 | 20994000 | 4430000 |
| Q15366-6;Q15366-3;Q15366;Q15366-2;H3BRU6;F8VZX2;Q15366-7;B4DXP5;Q15366-4;Q15366-5;F8W0G4;F8VXH9;F8W1G6;J3QT27;E9PFP8;P57721-2;P57721-3;P57721-5;P57721-4;P57721;H3BND9;H3BSS4;F8VRH0 | Poly(rC)-binding protein 2;Poly(rC)-binding protein 3 | PCBP2;PCBP3 | 12714000 | 5310700 | 20939000 | 5549500 | 14788000 | 6650200 |
| Q15386;Q15386-2;Q15386-3 | Ubiquitin-protein ligase E3C | UBE3C | 12463000 | 5447000 | 6849200 | 374820 | 441560 | 106520 |
| Q15393;Q15393-3;Q15393-2;J3QL37;J3QRB2;I3L4G7 | Splicing factor 3B subunit 3 | SF3B3 | 9566800 | 5296400 | 8242800 | 2745700 | 2374300 | 1692900 |
| Q15397 | Pumilio domain-containing protein KIAA0020 | KIAA0020 | 75055 | 96381 | 156850 | 186760 | 1850300 | 707030 |
| Q15427 | Splicing factor 3B subunit 4 | SF3B4 | 497530 | 478270 | 880770 | 600080 | 894790 | 290810 |
| Q15437;Q5QPE1;Q5QPE2 | Protein transport protein Sec23B | SEC23B | 428010 | 170910 | 172800 | 57167 | 0 | 0 |
| Q15459;Q15459-2;F8WB66 | Splicing factor 3A subunit 1 | SF3A1 | 1938300 | 990380 | 1494800 | 1289800 | 1302000 | 852370 |
| Q15477;H7C4L3 | Helicase SKI2W | SKIV2L | 1000600 | 471870 | 868220 | 441820 | 1510800 | 769770 |
| Q15633-2;Q15633;H3BV98;F8VZ57;F8VYK6;F8VZZ7;F8VTT7;F8VP94;F8VW32;F8VYK3 | RISC-loading complex subunit TARBP2 | TARBP2 | 787390 | 456250 | 1290100 | 307710 | 831420 | 435500 |
| Q15637-4;Q15637-6;Q15637-3;Q15637-2;Q15637;F8WEV5;Q15637-7;Q15637-5 | Splicing factor 1 | SF1 | 244450 | 256640 | 263790 | 116290 | 331790 | 117550 |
| Q15691;C9JB30;Q9UPY8-2;Q9UPY8 | Microtubule-associated protein RP/EB family member 1 | MAPRE1 | 3976600 | 1935100 | 3294100 | 265590 | 973450 | 130260 |
| Q15717;Q15717-2;M0QZR9;M0R055 | ELAV-like protein 1 | ELAVL1 | 6299900 | 5418700 | 10465000 | 2098400 | 13782000 | 2917200 |
| Q15773 | Myeloid leukemia factor 2 | MLF2 | 408850 | 512560 | 800210 | 0 | 0 | 0 |
| Q15819;G3V113;H0YBP9;H0YBX6;I3L0A0;Q13404-8;Q13404;Q13404-7;Q13404-2;Q13404-1;E5RIF1;D6RG00;G3V2F7;Q13404-6 | Ubiquitin-conjugating enzyme E2 variant 2;Ubiquitin-conjugating enzyme E2 variant 1 | UBE2V2;TMEM189-UBE2V1;UBE2V1 | 2905200 | 1290800 | 3359300 | 386280 | 1014300 | 68063 |
| Q16186;A0A087WX59;A0A087WUX6 | Proteasomal ubiquitin receptor ADRM1 | ADRM1 | 424820000 | 218330000 | 341920000 | 72762000 | 47243000 | 44713000 |

| | | | | | | | | |
|---|---|---|---|---|---|---|---|---|
| Q16204 | Coiled-coil domain-containing protein 6 | CCDC6 | 503640 | 98451 | 202820 | 0 | 0 | 39042 |
| Q16352;A0A087WYG8 | Alpha-internexin | INA | 87601 | 178790 | 257260 | 0 | 0 | 0 |
| Q16531;F5GY55;Q16531-2;F5H0Y5;F5H4N9;F5H7A0;F5H775;F5H581;F5H238;F8WF81;F5GWI0;F5H2L3;F5GZ34;F5H6C5;F5GYG8;F5GZY8 | DNA damage-binding protein 1 | DDB1 | 8083500 | 4779600 | 8149800 | 2842800 | 3409900 | 1822300 |
| Q16537-2;Q16537;Q16537-3;Q15173-2;Q15173;H0YN84;H7C5Q9;Q96B13;H0Y8C4;H0YJ75;E9PFR3;Q13362-2;Q13362-3;Q14738-3;Q13362;Q13362-4;Q13362-5;Q14738-2;Q14738 | Serine/threonine-protein phosphatase 2A 56 kDa regulatory subunit epsilon isoform;Serine/threonine-protein phosphatase 2A 56 kDa regulatory subunit beta isoform | PPP2R5E;PPP2R5B | 658670 | 169410 | 375970 | 0 | 72358 | 0 |
| Q16543;K7EL68;K7EQA9;K7EIU0;K7EKQ2 | Hsp90 co-chaperone Cdc37;Hsp90 co-chaperone Cdc37, N-terminally processed | CDC37 | 4552400 | 1866200 | 4929700 | 1794500 | 5691400 | 135340 |
| Q16643;Q16643-3;D6R9W4;Q16643-2;D6RCR4;D6RFI1;D6R9Q9 | Drebrin | DBN1 | 4097000 | 2029900 | 2116700 | 1541400 | 1990500 | 1291400 |
| Q16658 | Fascin | FSCN1 | 920250 | 423780 | 1063400 | 0 | 64340 | 70329 |
| Q1ED39;H3BNU8;H3BPL4 | Lysine-rich nucleolar protein 1 | KNOP1 | 98421 | 0 | 186430 | 0 | 0 | 0 |
| Q1KMD3;H3BQZ7 | Heterogeneous nuclear ribonucleoprotein U-like protein 2 | HNRNPUL2;HNRNPUL2-BSCL2 | 7501500 | 3575700 | 6965800 | 9240100 | 12255000 | 6486000 |
| Q27J81-2;Q27J81 | Inverted formin-2 | INF2 | 93785 | 103070 | 244790 | 0 | 0 | 0 |
| Q29RF7;H0Y9L9;Q29RF7-3 | Sister chromatid cohesion protein PDS5 homolog A | PDS5A | 64649 | 80856 | 380550 | 1160600 | 1906700 | 783270 |
| Q2NL82;I3L1Q5 | Pre-rRNA-processing protein TSR1 homolog | TSR1 | 6331200 | 3370200 | 8636600 | 3822400 | 4306400 | 1190300 |
| Q2TAY7;Q2TAY7-2 | WD40 repeat-containing protein SMU1;WD40 repeat-containing protein SMU1, N-terminally processed | SMU1 | 455180 | 438620 | 787850 | 178260 | 402500 | 104560 |
| Q32P28-4;Q32P28;Q32P28-3;H7C0D1;H7C2W6 | Prolyl 3-hydroxylase 1 | LEPRE1 | 630670 | 334300 | 170830 | 317960 | 225290 | 126540 |
| Q3KQU3-2;Q3KQU3-4;Q3KQU3;E9PLH3;C9JIR3;H0YF21 | MAP7 domain-containing protein 1 | MAP7D1 | 170630 | 81310 | 332130 | 622170 | 615420 | 413910 |
| Q4G0J3;Q4G0J3-3;H0YA82;D6RFF0;D6R9Z6;D6RF49;D6RAF3;D6RF22 | La-related protein 7 | LARP7 | 1848000 | 733560 | 2278900 | 1807400 | 3493400 | 2215400 |
| Q4VCS5;A6NP16;E7ERM3;Q4VCS5-2 | Angiomotin | AMOT | 600600 | 215600 | 420640 | 0 | 0 | 0 |
| Q52LJ0;Q52LJ0-2 | Protein FAM98B | FAM98B | 9621600 | 2841400 | 4344600 | 3163200 | 10804000 | 6628800 |
| Q53EL6-2;Q53EL6;Q5VZS7 | Programmed cell death protein 4 | PDCD4 | 4959700 | 3346300 | 7345400 | 10934000 | 20075000 | 9125300 |
| Q53H80 | Akirin-2 | AKIRIN2 | 1408200 | 2066800 | 2324500 | 0 | 0 | 0 |
| Q562R1 | Beta-actin-like protein 2 | ACTBL2 | 3110900 | 55306 | 3087700 | 0 | 0 | 89437 |
| Q5C9Z4 | Nucleolar MIF4G domain-containing protein 1 | NOM1 | 31779 | 0 | 43686 | 196460 | 523610 | 151360 |
| Q5F1R6;Q5F1R6-3;Q5F1R6-2 | DnaJ homolog subfamily C member 21 | DNAJC21 | 856760 | 411060 | 985090 | 354020 | 505180 | 299820 |
| Q5JR04;Q9HCE1;Q9HCE1-2 | Putative helicase MOV-10 | MOV10 | 2085400 | 1484200 | 3216200 | 2077800 | 5028900 | 2170100 |
| Q5JRC6;Q8IWS0-5;Q8IWS0-3;Q8IWS0;Q8IWS0-4;Q8IWS0-2 | PHD finger protein 6 | PHF6 | 463910 | 235740 | 282760 | 80377 | 757460 | 338480 |
| Q5JTH9-2;Q5JTH9-3;Q5JTH9;A0A087X1U8;H7C1M8;A0A087WYW2;A0A087X0G0 | RRP12-like protein | RRP12 | 2914100 | 1077200 | 3544900 | 1953800 | 3597400 | 1459400 |
| X6R5Z6;Q5JTJ3-3;Q5JTJ3;Q5JTJ3-2 | Cytochrome c oxidase assembly factor 6 homolog | COA6 | 970650 | 0 | 1303200 | 0 | 0 | 199260 |
| Q5JUP3;Q9BQI0-4;Q9BQI0-3;Q9BQI0;Q9BQI0-2 | Allograft inflammatory factor 1-like | AIF1L | 3205100 | 3327900 | 3545500 | 0 | 0 | 0 |
| Q5JVF3-3;Q5JVF3-2;Q5JVF3;Q5JVF3-4 | PCI domain-containing protein 2 | PCID2 | 0 | 56056 | 113620 | 121940 | 284790 | 90724 |
| Q5JY65;Q9BZJ0-2;Q9BZJ0-3;Q9BZJ0 | Crooked neck-like protein 1 | CRNKL1 | 122550 | 107220 | 193250 | 217890 | 361370 | 177150 |
| Q5QPM7;Q92530;H0Y555;Q5QPM9;F5H4Z3 | Proteasome inhibitor PI31 subunit | PSMF1 | 9283900 | 7484600 | 7968000 | 2097100 | 4322700 | 2584700 |
| Q5RKV6 | Exosome complex component MTR3 | EXOSC6 | 21933000 | 6313700 | 13592000 | 24735000 | 33388000 | 14882000 |
| Q5SQ08 | | C8G | 3880700 | 0 | 0 | 521920 | 507890 | 119900 |
| Q5SSJ5;X6RGJ2;Q5SSJ5-3;Q5SSJ5-2;B0QZK4;Q5SWC8;B0QZK9;B0QZK8;Q5SSJ5-5 | Heterochromatin protein 1-binding protein 3 | HP1BP3 | 1769600 | 686390 | 998540 | 1073700 | 1865800 | 1013100 |
| Q5SW79-2;Q5SW79-3;Q5SW79;H0Y2V6;E5RIH6;E5RGW7;E5RJT5;E5RFU8;E5RG47;H0Y4T4;E7EWM2;E7EMW0;Q96L14 | Centrosomal protein of 170 kDa | CEP170 | 156550 | 103360 | 245990 | 0 | 58238 | 11460 |
| Q5SYF9;Q5VY60;Q7Z4H7-2;Q7Z4H7-3;Q7Z4H7 | HAUS augmin-like complex subunit 6 | HAUS6 | 582050 | 0 | 399600 | 0 | 0 | 0 |
| Q5T1M5-2;Q5T1M5 | FK506-binding protein 15 | FKBP15 | 167270 | 71014 | 142190 | 0 | 0 | 0 |
| Q5T280;A0A087WYA7;R4GNG4 | Putative methyltransferase C9orf114 | C9orf114 | 199490 | 266510 | 286070 | 191340 | 202920 | 166590 |
| Q5T3Q7;Q9H583 | HEAT repeat-containing protein 1;HEAT repeat-containing protein 1, N-terminally processed | HEATR1 | 15796 | 23526 | 46048 | 53306 | 77573 | 55146 |
| Q5TDF0;Q9BSD7 | Cancer-related nucleoside-triphosphatase | NTPCR | 282620 | 110980 | 633110 | 0 | 389650 | 73274 |
| Q5TDH0;Q5TDH0-3;Q5TDH0-2;H0YII4;H0YI90;Q8WTU0 | Protein DDI1 homolog 2 | DDI2 | 30409000 | 9298000 | 13594000 | 522980 | 617910 | 330290 |
| Q5TZA2;Q5TZA2-2 | Rootletin | CROCC | 74400 | 20763 | 4685000 | 32630 | 29047 | 27938 |
| Q5VTE6-2;Q5VTE6 | Protein angel homolog 2 | ANGEL2 | 194460 | 0 | 94005 | 92934 | 183780 | 0 |
| Q5VTR2;C9JWJ4;C9J0A5;C9JXC9 | E3 ubiquitin-protein ligase BRE1A | RNF20 | 231530 | 236400 | 291920 | 24105 | 37040 | 31061 |
| Q5VW32-2;Q5VW32;Q5VW33;F5GXQ0 | BRO1 domain-containing protein BROX | BROX | 715130 | 81889 | 447740 | 0 | 0 | 0 |
| Q5VW36;S4R400 | Focadhesin | FOCAD | 240930 | 24520 | 314560 | 84404 | 166010 | 55483 |
| Q5VWC4;P55036-2;H7C2J9 | 26S proteasome non-ATPase regulatory subunit 4 | PSMD4 | 914310 | 526800 | 495530 | 0 | 0 | 0 |
| Q5VY93;V9GYM8;Q92974-3;Q92974-2;Q92974;V9GYF0 | Rho guanine nucleotide exchange factor 2 | ARHGEF2 | 532900 | 301520 | 370280 | 1401800 | 1058000 | 1773500 |
| Q5VYK3;J3KN16;R4GMY1;F6XAQ5;R4GMY3 | Proteasome-associated protein ECM29 homolog | ECM29;KIAA0368 | 84787000 | 43756000 | 59619000 | 8988700 | 7846000 | 4767000 |
| Q5VYS8-6;Q5VYS8;X6R3Q3;Q5VYS8-4;Q5VYT0;Q5VYS8-3;Q5VYS8-2 | Terminal uridylyltransferase 7 | ZCCHC6 | 0 | 184020 | 535060 | 5024.4 | 22390 | 0 |
| Q66GS9 | Centrosomal protein of 135 kDa | CEP135 | 36710 | 0 | 0 | 0 | 0 | 0 |
| Q66K74-2;Q66K74;M0R1M7;M0QXQ9;M0R1J2;M0QY41 | Microtubule-associated protein 1S;MAP1S heavy chain;MAP1S light chain | MAP1S | 196960 | 23026 | 367280 | 0 | 310070 | 100430 |
| Q68EM7-2;Q68EM7-6;Q68EM7-5;Q68EM7;C9IZD3;Q68EM7-4;Q68EM7-7;Q68EM7-3 | Rho GTPase-activating protein 17 | ARHGAP17 | 227500 | 80990 | 52903 | 0 | 0 | 0 |
| Q69YN2;Q69YN2-3;X6R7M0;Q69YN2-2 | CWF19-like protein 1 | CWF19L1 | 1724900 | 1185100 | 2113500 | 1020700 | 1560200 | 882740 |
| Q6IA86-4;Q6IA86-7;Q6IA86-2;Q6IA86-3;Q6IA86-5;Q6IA86;Q6IA86-6 | Elongator complex protein 2 | ELP2 | 0 | 53923 | 164170 | 0 | 0 | 0 |
| Q6IBW4-2;Q6IBW4;Q6IBW4-4;E5RJN3;H0YC55;Q6IBW4-5 | Condensin-2 complex subunit H2 | NCAPH2 | 1010800 | 807910 | 1694000 | 1820900 | 2933700 | 1142300 |
| Q6IQ49-2;Q6IQ49;Q6IQ49-3 | Protein SDE2 homolog | SDE2 | 925480 | 1215600 | 1898500 | 1754700 | 2300600 | 829470 |
| Q6N069;Q6N069-5;Q6N069-4;Q6N069-3;Q6N069-2 | N-alpha-acetyltransferase 16, NatA auxiliary subunit | NAA16 | 79980 | 14940 | 372240 | 49832 | 269880 | 232550 |
| Q6NZY4;Q6NZY4-2;F5GXQ6;F5H803;F5H6J5;F5GX80 | Zinc finger CCHC domain-containing protein 8 | ZCCHC8 | 895490 | 529450 | 554840 | 585030 | 878390 | 506230 |
| Q6P158;Q6P158-2;H7C109;A0A087WZ11;Q6P158-3;A0A087WW12 | Putative ATP-dependent RNA helicase DHX57 | DHX57 | 1411600 | 803400 | 1415100 | 454490 | 1505200 | 419580 |
| Q6P1J9 | Parafibromin | CDC73 | 1828700 | 773750 | 1407700 | 68123 | 123360 | 142800 |
| Q6P2E9;Q6P2E9-2 | Enhancer of mRNA-decapping protein 4 | EDC4 | 8912900 | 744650 | 2354900 | 6927900 | 1231200 | 558820 |
| Q6P2Q9;I3L0J9;I3L1T8 | Pre-mRNA-processing-splicing factor 8 | PRPF8 | 1469800 | 866720 | 1823000 | 742550 | 1007100 | 609510 |
| Q6P4E2;Q96J85;Q71RC2-7;Q71RC2-6;Q71RC2-3;Q71RC2;Q71RC2-4;Q8TBL5;A0A087WTL9;Q71RC2-2;Q71RC2-5;F8W1I4;F8VY40;X6RLN4 | La-related protein 4 | LARP4 | 2772000 | 1404900 | 1507200 | 592710 | 3605200 | 1621400 |
| Q6PGP7;D6RCE2 | Tetratricopeptide repeat protein 37 | TTC37 | 3029800 | 880480 | 2068600 | 2375600 | 3293800 | 1399900 |
| Q6PK04;I3L0U5 | Coiled-coil domain-containing protein 137 | CCDC137 | 125440 | 0 | 344060 | 478260 | 814680 | 261020 |
| Q6PKG0;Q6PKG0-3;E5RH50;H0YC33;H0YBW1;H0YBJ5;H0YBM7;H0YC73;H0YAN4;D6RJB0;Q659C4-9;Q659C4-2;Q659C4 | La-related protein 1 | LARP1 | 9372300 | 5231600 | 10465000 | 11850000 | 21079000 | 10668000 |
| Q6QNY0 | Biogenesis of lysosome-related organelles complex 1 subunit 3 | BLOC1S3 | 0 | 0 | 155630 | 0 | 0 | 0 |
| Q6UXN9;C9JBU3 | WD repeat-containing protein 82 | WDR82 | 5266700 | 2756400 | 3836000 | 2963200 | 2580200 | 2040800 |
| Q6WKZ4-5;Q6WKZ4-3;Q6WKZ4 | Rab11 family-interacting protein 1 | RAB11FIP1 | 20916000 | 5448400 | 13740000 | 0 | 0 | 0 |
| Q6ZN17-2;Q6ZN17 | Protein lin-28 homolog B | LIN28B | 1540600 | 0 | 1611800 | 456990 | 1438300 | 842340 |
| Q6ZSJ8;Q6ZSJ8-2 | Uncharacterized protein C1orf122 | C1orf122 | 823300 | 507960 | 1144600 | 892960 | 2181300 | 893880 |
| Q7KZ85;Q7KZ85-2;Q7KZ85-3 | Transcription elongation factor SPT6 | SUPT6H | 2216200 | 1028400 | 2259700 | 218590 | 293760 | 91363 |
| Q7KZF4;H7C597 | Staphylococcal nuclease domain-containing protein 1 | SND1 | 15888000 | 8710100 | 19926000 | 15105000 | 29735000 | 14586000 |
| Q7L1Q6-2;Q7L1Q6;Q7L1Q6-4;Q7L1Q6-3;C9IZ80;C9JFN4;H0Y503 | Basic leucine zipper and W2 domain-containing protein 1 | BZW1 | 2845800 | 1311200 | 2885100 | 357320 | 1030300 | 451480 |
| Q7L2H7;E9PRI2;E7ESM3;Q7L2H7-2 | Eukaryotic translation initiation factor 3 subunit M | EIF3M | 28173000 | 13603000 | 35028000 | 26207000 | 32032000 | 28248000 |
| Q7L2J0;Q7L2J0-2 | 7SK snRNA methylphosphate capping enzyme | MEPCE | 0 | 59830 | 41329 | 0 | 217380 | 29460 |
| Q7RTV0 | PHD finger-like domain-containing protein 5A | PHF5A | 10189000 | 5685100 | 8737400 | 5677400 | 5728800 | 2310600 |
| Q7Z2T5;Q7Z2T5-2;X6RK96 | TRMT1-like protein | TRMT1L | 1182400 | 740280 | 1596500 | 2191800 | 4149500 | 1752100 |
| Q7Z2Z2-2;Q7Z2Z2 | Elongation factor Tu GTP-binding domain-containing protein 1 | EFTUD1 | 0 | 0 | 0 | 113220 | 448650 | 157610 |
| Q7Z417 | Nuclear fragile X mental retardation-interacting protein 2 | NUFIP2 | 176580 | 48062 | 217910 | 58801 | 345450 | 244450 |
| Q8WVX6;Q7Z5K2;Q7Z5K2-2;Q7Z5K2-3 | Wings apart-like protein homolog | WAPAL | 0 | 0 | 0 | 0 | 318540 | 242670 |
| Q7Z6K5;Q7Z6K5-2 | Arpin | ARPIN | 0 | 142410 | 182590 | 0 | 0 | 0 |
| Q7Z7G8-2;Q7Z7G8-6;Q7Z7G8;Q7Z7G8-5;Q7Z7G8-4;Q7Z7G8-3 | Vacuolar protein sorting-associated protein 13B | VPS13B | 144400 | 207960 | 257880 | 78825 | 175750 | 159190 |
| Q86TB9-2;Q86TB9-4;Q86TB9 | Protein PAT1 homolog 1 | PATL1 | 374230 | 88286 | 449060 | 182230 | 239300 | 74165 |
| Q86U38;Q86U38-2 | Nucleolar protein 9 | NOP9 | 128630 | 197900 | 603610 | 345350 | 676190 | 291430 |
| Q86U42-2;Q86U42;B4DEH8;G3V4T2;Q92843-2;H0YJH9 | Polyadenylate-binding protein 2 | PABPN1 | 3806300 | 1718100 | 1946500 | 93153 | 1301900 | 258440 |
| Q86UK7-2;Q86UK7-3;Q86UK7;H3BQQ2;H3BR87;Q86UK7-4 | Zinc finger protein 598 | ZNF598 | 0 | 108550 | 298180 | 118270 | 354380 | 184350 |
| Q86V93 | | LOC100128510 | 583340 | 290560 | 649330 | 169190 | 73976 | 0 |
| Q86VP6;Q86VP6-2;H0YH27;Q86VP6-3;F5H6I6;F8WBB8;O75155-2;O75155 | Cullin-associated NEDD8-dissociated protein 1 | CAND1 | 20367000 | 2659500 | 17784000 | 2362600 | 2017200 | 1048300 |
| Q86W42-3;Q86W42-2;Q86W42 | THO complex subunit 6 homolog | THOC6 | 161450 | 0 | 412210 | 0 | 0 | 0 |
| Q86W50 | Methyltransferase-like protein 16 | METTL16 | 0 | 33456 | 62028 | 21068 | 15898 | 7892.6 |
| Q86XP3-2;Q86XP3 | ATP-dependent RNA helicase DDX42 | DDX42 | 65526 | 150730 | 123350 | 231390 | 618500 | 308310 |
| Q86XZ4;F8VS10 | Spermatogenesis-associated serine-rich protein 2 | SPATS2 | 0 | 0 | 0 | 146620 | 258360 | 113070 |
| Q86YP4-2;Q86YP4;Q86YP4-3;Q8WXI9 | Transcriptional repressor p66-alpha;Transcriptional repressor p66-beta | GATAD2A;GATAD2B | 178710 | 132970 | 445550 | 131810 | 120420 | 42621 |
| Q86YQ0;Q96MX6;Q96MX6-2;H7C389;Q8ND98 | WD repeat-containing protein 92 | HZGJ;WDR92;DKFZp434B156 | 1175900 | 372600 | 974670 | 743520 | 1105400 | 452570 |
| Q86YS7;Q86YS7-2;Q86YS7-3;Q86YS7-4;Q86YS7-5;H0YGU7 | C2 domain-containing protein 5 | C2CD5 | 142870 | 0 | 375120 | 27004 | 0 | 0 |
| X6RLX0;Q8IUD2-3;Q8IUD2;X6RM00;G8JLD3;Q8IUD2-4;O15083;Q8IUD2-2;K7EPP6;H7C4G9;Q8IUD2-5 | ELKS/Rab6-interacting/CAST family member 1;ERC protein 2 | ERC1;ERC2 | 172440 | 35343 | 62501 | 0 | 128270 | 0 |
| Q8IUF8-4;Q8IUF8;H0Y9L8;Q8IUF8-3;Q8IUF8-2 | Bifunctional lysine-specific demethylase and histidyl-hydroxylase MINA | MINA | 369250 | 327030 | 784270 | 751830 | 1404800 | 900010 |
| Q8IUI8-2;Q8IUI8;J3KST8 | Cytokine receptor-like factor 3 | CRLF3 | 1194100 | 902720 | 1991300 | 0 | 69078 | 0 |
| Q8IV48;E5RIV7 | 3-5 exoribonuclease 1 | ERI1 | 106390 | 569560 | 981230 | 522090 | 809010 | 418170 |
| Q8IVF2-3;Q8IVF2 | Protein AHNAK2 | AHNAK2 | 38031 | 5677.8 | 43175 | 0 | 0 | 0 |
| Q8IVL6-2;Q8IVL6 | Prolyl 3-hydroxylase 3 | LEPREL2 | 126020 | 321410 | 272930 | 0 | 0 | 0 |
| Q8IVM0;Q8IVM0-2 | Coiled-coil domain-containing protein 50 | CCDC50 | 0 | 0 | 0 | 0 | 0 | 0 |
| Q8IX12-2;Q8IX12 | Cell division cycle and apoptosis regulator protein 1 | CCAR1 | 125190 | 116970 | 243390 | 0 | 0 | 0 |
| Q8IXB1-2;Q8IXB1;E7EP04;Q8IXB1-3 | DnaJ homolog subfamily C member 10 | DNAJC10 | 1391000 | 711070 | 777770 | 31687 | 36326 | 0 |
| Q8IXW5-2;Q8IXW5 | Putative RNA polymerase II subunit B1 CTD phosphatase RPAP2 | RPAP2 | 165250 | 249110 | 376510 | 55448 | 97238 | 64690 |
| Q8IY37;F5H3Y4 | Probable ATP-dependent RNA helicase DHX37 | DHX37 | 508630 | 145100 | 41786 | 318220 | 688960 | 379120 |
| Q8IY81;J3KS74;J3QSE2;J3KS36 | pre-rRNA processing protein FTSJ3 | FTSJ3 | 0 | 0 | 357290 | 1079300 | 2896600 | 916330 |
| Q8IYI6 | Exocyst complex component 8 | EXOC8 | 251890 | 117090 | 375140 | 35943 | 61316 | 0 |
| Q8IZH2-2;Q8IZH2;H7C5E4;Q8IZH2-3 | 5-3 exoribonuclease 1 | XRN1 | 1527400 | 920940 | 1518600 | 683450 | 925280 | 368970 |
| Q8N0X7 | Spartin | SPG20 | 3632700 | 1895300 | 2374500 | 726860 | 654300 | 383520 |
| Q8N0Z6;H9KV81 | Tetratricopeptide repeat protein 5 | TTC5 | 362740 | 41856 | 101830 | 0 | 57857 | 0 |
| Q8N163-2;Q8N163;G3V119;H0YB24;E5RHJ4;H0YC69;E5RGU7;E5RFJ3;H0YC58;A0A087X2B6 | Cell cycle and apoptosis regulator protein 2 | CCAR2 | 456770 | 165110 | 522850 | 50221 | 120820 | 21291 |
| Q8N1G4 | Leucine-rich repeat-containing protein 47 | LRRC47 | 6786000 | 3563100 | 10601000 | 4895500 | 14549000 | 4635300 |
| Q8N3C0;Q8N3C0-4 | Activating signal cointegrator 1 complex subunit 3 | ASCC3 | 589320 | 306220 | 782720 | 283920 | 676180 | 232330 |
| Q8N3R9-2;Q8N3R9 | MAGUK p55 subfamily member 5 | MPP5 | 0 | 0 | 0 | 0 | 0 | 0 |
| Q8N3U4;Q8N3U4-2;B1AMT4;Q8WVM7-2;Q8WVM7;Q68DW7;D6W5U7;Q9UJ98-2;Q9UJ98-3;Q9UJ98 | Cohesin subunit SA-2 | STAG2 | 335640 | 290770 | 744710 | 546730 | 1311200 | 481080 |
| Q8N668-2;Q8N668 | COMM domain-containing protein 1 | COMMD1 | 0 | 0 | 338780 | 0 | 0 | 0 |
| Q8N9T8;F8WFC1;H0YFD2 | Protein KRI1 homolog | KRI1 | 289440 | 111150 | 1366000 | 860480 | 3377700 | 1431900 |
| Q8NBJ5;M0QX72 | Procollagen galactosyltransferase 1 | COLGALT1 | 192610 | 263460 | 188650 | 0 | 0 | 26901 |
| Q8NC51;Q8NC51-3;Q8NC51-2;Q8NC51-4 | Plasminogen activator inhibitor 1 RNA-binding protein | SERBP1 | 116480000 | 97666000 | 182960000 | 163760000 | 291910000 | 185480000 |
| Q8NCM8;Q8NCM8-2;Q8NCM8-3 | Cytoplasmic dynein 2 heavy chain 1 | DYNC2H1 | 63371 | 37452 | 54836 | 493060 | 706920 | 455240 |
| Q8ND04-3;Q8ND04;Q8ND04-2;J3KTD7 | Protein SMG8 | SMG8 | 0 | 0 | 176460 | 81876 | 139420 | 82169 |
| Q8NE71;Q8NE71-2;H0YGW7;Q5STZ8;F5GYK6 | ATP-binding cassette sub-family F member 1 | ABCF1 | 5494400 | 3096200 | 4273300 | 2223500 | 3863400 | 2234700 |
| Q8NEM2 | SHC SH2 domain-binding protein 1 | SHCBP1 | 461340 | 86547 | 300250 | 107570 | 130000 | 0 |
| Q8NEZ5;H3BTH6;H3BTR7;Q8NEZ5-3;H3BRE0;Q8NEZ5-2 | F-box only protein 22 | FBXO22 | 7387500 | 4334900 | 7184000 | 3014300 | 4102900 | 1946800 |
| Q8NFW8;F5GYM0;Q8NFW8-2 | N-acylneuraminate cytidylyltransferase | CMAS | 331990 | 196160 | 321780 | 120860 | 451310 | 209980 |
| Q8NI27;F2Z2V2 | THO complex subunit 2 | THOC2 | 590380 | 159050 | 446930 | 129460 | 277450 | 0 |
| Q8NI36 | WD repeat-containing protein 36 | WDR36 | 155260 | 135170 | 442480 | 643310 | 1385800 | 649870 |
| Q8TAF3;Q8TAF3-4;Q8TAF3-3;Q8TAF3-5;F8W9K4;F8W7Q3;C9JC24 | WD repeat-containing protein 48 | WDR48 | 412760 | 280680 | 426260 | 381190 | 496280 | 292610 |
| Q8TBB5-3;Q8TBB5;H3BQH3;Q8TBB5-2;H3BP99;H3BQL0;H3BQ90;H3BV82;Q9UF94 | Kelch domain-containing protein 4 | KLHDC4 | 3533600 | 1114300 | 2653300 | 2273500 | 2504400 | 612090 |
| Q8TBM3;Q9Y487 | V-type proton ATPase subunit a;V-type proton ATPase 116 kDa subunit a isoform 2 | ATP6V0A2 | 22820000 | 13376000 | 18979000 | 29539000 | 26395000 | 24492000 |
| Q8TD16;Q8TD16-2 | Protein bicaudal D homolog 2 | BICD2 | 0 | 0 | 0 | 0 | 0 | 0 |
| Q8TDD1;Q8TDD1-2;F8VRX4;H0YHH7 | ATP-dependent RNA helicase DDX54 | DDX54 | 926190 | 599040 | 1112000 | 2048200 | 4335300 | 2163900 |
| Q8TDN6 | Ribosome biogenesis protein BRX1 homolog | BRIX1 | 910330 | 727760 | 2937600 | 5808500 | 8365000 | 5050500 |
| Q8TEA8 | D-tyrosyl-tRNA(Tyr) deacylase 1 | DTD1 | 1162000 | 942400 | 2708600 | 1616800 | 2309600 | 1380100 |
| Q8TEQ6 | Gem-associated protein 5 | GEMIN5 | 3407600 | 1990200 | 3995500 | 2288900 | 4433200 | 2262100 |
| Q9NP97;Q8TF09;H3BNG9;H3BPA0;Q7Z4M1;Q9NP97-2;H3BQI1;B1AKR6 | Dynein light chain roadblock-type 1;Dynein light chain roadblock-type 2 | DYNLRB1;DYNLRB2 | 1098200 | 1166200 | 3217100 | 1795700 | 2104500 | 1238200 |
| Q8TF46-2;Q8TF46-4;Q8TF46 | DIS3-like exonuclease 1 | DIS3L | 0 | 0 | 39672 | 0 | 77424 | 48367 |
| Q8WTT2 | Nucleolar complex protein 3 homolog | NOC3L | 339140 | 93785 | 533770 | 923260 | 2807600 | 1226300 |
| Q8WU90;Q8WU90-2;H7C466 | Zinc finger CCCH domain-containing protein 15 | ZC3H15 | 5303600 | 2081600 | 7381600 | 5087500 | 5790800 | 3964400 |
| Q8WUA2 | Peptidyl-prolyl cis-trans isomerase-like 4 | PPIL4 | 2855800 | 1980500 | 2836300 | 1920400 | 2061200 | 1313000 |
| Q8WUA4;A0A087WZD8;Q8WUA4-2;H7C3X9;H0Y4Q6 | General transcription factor 3C polypeptide 2 | GTF3C2 | 277710 | 104210 | 786430 | 0 | 9657.3 | 20001 |
| Q8WUM4;Q8WUM4-2;Q8WUM4-3;C9IZF9;F8WEQ7;F8WBR8;F8WDK9 | Programmed cell death 6-interacting protein | PDCD6IP | 4100800 | 794090 | 1488700 | 9388.9 | 152380 | 38975 |
| Q8WVY7 | Ubiquitin-like domain-containing CTD phosphatase 1 | UBLCP1 | 8581200 | 3354600 | 5103100 | 189760 | 211190 | 33966 |
| Q8WW12-2;Q8WW12 | PEST proteolytic signal-containing nuclear protein | PCNP | 0 | 1071300 | 1474700 | 3984500 | 3300500 | 1773900 |
| Q8WWH5 | Probable tRNA pseudouridine synthase 1 | TRUB1 | 134480 | 42317 | 356210 | 0 | 218510 | 55690 |
| Q8WX92 | Negative elongation factor B | NELFB | 1051800 | 872610 | 1350700 | 1249200 | 1492100 | 687680 |

| | | | | | | | | |
|---|---|---|---|---|---|---|---|---|
| Q8WX93-7;Q8WX93-4;Q8WX93-8;Q8WX93-3;Q8WX93-5;Q8WX93;Q8WX93-2;Q8WX93-9;H0YA05 | Palladin | PALLD | 642510 | 526190 | 521470 | 0 | 0 | 0 |
| Q8WXE1-5;Q8WXE1-3;Q8WXE1-2;Q8WXE1 | ATR-interacting protein | ATRIP | 0 | 0 | 0 | 0 | 1081400 | 804050 |
| X6RDA4;Q8WXF1-2;Q8WXF1 | Paraspeckle component 1 | PSPC1 | 1462600 | 917040 | 2258000 | 504620 | 1643800 | 1032500 |
| Q8WXX5 | DnaJ homolog subfamily C member 9 | DNAJC9 | 2393500 | 1301000 | 3104500 | 1057900 | 3538100 | 1331100 |
| Q8WYQ5-3;Q8WYQ5 | Microprocessor complex subunit DGCR8 | DGCR8 | 0 | 0 | 74185 | 0 | 82383 | 36106 |
| Q8WZ42-13;Q8WZ42-12;Q8WZ42-11;Q8WZ42-4;Q8WZ42-7;Q8WZ42-2;Q8WZ42;Q8WZ42-8;Q8WZ42-3;Q8WZ42-10;Q8WZ42-9;Q8WZ42-5;A2TKE4;Q8WZ42-6;H0Y4J7 | Titin | TTN | 42202 | 41810 | 55548 | 85602 | 122180 | 65237 |
| Q92499;F1T0B3;A0A087X2G1;Q92499-3;Q92499-2 | ATP-dependent RNA helicase DDX1 | DDX1 | 14457000 | 9130600 | 18254000 | 15324000 | 23002000 | 12453000 |
| Q92522 | Histone H1x | H1FX | 3561500 | 2078300 | 2916400 | 1863200 | 3460900 | 2044300 |
| Q92598-2;Q92598-3;Q92598;Q92598-4 | Heat shock protein 105 kDa | HSPH1 | 328380 | 168900 | 470250 | 71331 | 18010 | 0 |
| Q92600;Q92600-2 | Cell differentiation protein RCD1 homolog | RQCD1 | 292720 | 391200 | 722570 | 299080 | 504420 | 163810 |
| Q92614-3;Q92614-4;Q92614;Q92614-2;Q92614-5;H0YEV9 | Unconventional myosin-XVIIIa | MYO18A | 248830 | 80656 | 284400 | 0 | 6449.9 | 0 |
| Q92615;H0Y4V9;A0A087X183;H0Y641;X6RCG6;V9GYI5;B5MCU2 | La-related protein 4B | LARP4B | 1404300 | 682710 | 1465200 | 1058700 | 3116000 | 1314700 |
| Q92616 | Translational activator GCN1 | GCN1L1 | 3777900 | 2285800 | 5323600 | 4091000 | 9110300 | 4120500 |
| Q92620;Q92620-2;H3BV01;J3KTG2;H3BQT9;J3KRT1 | Pre-mRNA-splicing factor ATP-dependent RNA helicase PRP16 | DHX38 | 839690 | 710840 | 878040 | 38969 | 322460 | 74633 |
| Q92688-2;Q92688 | Acidic leucine-rich nuclear phosphoprotein 32 family member B | ANP32B | 42362000 | 22395000 | 49985000 | 29346000 | 27440000 | 18219000 |
| Q92769-3;Q92769;Q13547;E5RK19;E5RFI6;E5RJ04;E5RGV4;E5RHE7;E5RH52;E5RG37;E5RFP9;H3BM24;Q5TEE2;Q96G03 | Histone deacetylase 2;Histone deacetylase 1 | HDAC2;HDAC1 | 2131400 | 1135900 | 1054200 | 0 | 210950 | 0 |
| Q92878;Q92878-2;Q92878-3;E7EN38;E7ESD9;E9PM98;C9JNH8;H7C0V2 | DNA repair protein RAD50 | RAD50 | 840190 | 656060 | 1545900 | 845320 | 1444700 | 570240 |
| Q92900-2;Q92900 | Regulator of nonsense transcripts 1 | UPF1 | 1629700 | 785370 | 1656900 | 1168000 | 1844600 | 857400 |
| Q92905 | COP9 signalosome complex subunit 5 | COPS5 | 1056900 | 450180 | 882130 | 369230 | 434670 | 76285 |
| Q92917 | G patch domain and KOW motifs-containing protein | GPKOW | 2350500 | 1843200 | 2148600 | 430810 | 734830 | 350200 |
| Q92973-2;Q92973;Q92973-3;E7EW37;A0A075B780;O14787-2;O14787 | Transportin-1 | TNPO1 | 690050 | 269780 | 450850 | 329620 | 195630 | 101710 |
| Q92979;V9GYP5;A0A087WWQ2;A0A087WVM7 | Ribosomal RNA small subunit methyltransferase NEP1 | EMG1 | 1386700 | 1531600 | 3163500 | 10375000 | 11267000 | 7645100 |
| Q93008-1;Q93008;O00507;O00507-2 | Probable ubiquitin carboxyl-terminal hydrolase FAF-X | USP9X | 2487500 | 1193400 | 2077700 | 710690 | 661050 | 375730 |
| Q93009-3;Q93009;H3BND8;F5H2X1 | Ubiquitin carboxyl-terminal hydrolase 7;Ubiquitin carboxyl-terminal hydrolase | USP7 | 4246700 | 2412400 | 3060600 | 4310800 | 2327400 | 4097700 |
| Q969Q0 | 60S ribosomal protein L36a-like | RPL36AL | 120410000 | 54300000 | 140600000 | 97243000 | 189900000 | 27369000 |
| Q96A65;Q96A65-2 | Exocyst complex component 4 | EXOC4 | 286980 | 228350 | 591890 | 135600 | 145780 | 0 |
| Q96A72;F5H6N1;F5H124 | Protein mago nashi homolog 2 | MAGOHB | 2325300 | 1625100 | 4765400 | 1283800 | 3273400 | 1445800 |
| Q96B26 | Exosome complex component RRP43 | EXOSC8 | 7067900 | 3810200 | 5495600 | 6727300 | 9770300 | 5269100 |
| Q96C19;H0Y4Y4;C9JTV4;Q8WYH2;A0A087X1Z2;Q9BUP0-2;Q9BUP0 | EF-hand domain-containing protein D2 | EFHD2 | 6431400 | 2493600 | 4356800 | 0 | 0 | 0 |
| Q96CT7;M0R2F5 | Coiled-coil domain-containing protein 124 | CCDC124 | 40581000 | 20536000 | 50522000 | 53633000 | 97901000 | 43998000 |
| Q96CW5-2;Q96CW5;A0A087WWB5;A0A087WU06;Q96CW5-3 | Gamma-tubulin complex component 3 | TUBGCP3 | 290200 | 150010 | 413430 | 0 | 0 | 0 |
| Q96CX2;Q6ZWB6 | BTB/POZ domain-containing protein KCTD12 | KCTD12 | 1773500 | 0 | 966640 | 1173700 | 340770 | 0 |
| Q96DI7;Q96DI7-2 | U5 small nuclear ribonucleoprotein 40 kDa protein | SNRNP40 | 557030 | 386590 | 669920 | 742700 | 816230 | 415200 |
| Q96F86;H3BQ37;H3BPW9;H3BTD6;H3BMB8;H3BTH0;H3BQP5;H3BU87;H3BPN4 | Enhancer of mRNA-decapping protein 3 | EDC3 | 1041700 | 457070 | 946390 | 558890 | 1098700 | 373360 |
| Q96FX7;H0Y2Q1 | tRNA (adenine(58)-N(1))-methyltransferase catalytic subunit TRMT61A | TRMT61A | 1745000 | 64083 | 1367000 | 0 | 4930600 | 64230 |
| Q96GA3;A0A075B6Q4 | Protein LTV1 homolog | LTV1 | 5628500 | 4373500 | 11331000 | 2001700 | 7410300 | 3549100 |
| Q96GM8;Q96GM8-2 | Target of EGR1 protein 1 | TOE1 | 5123500 | 1507500 | 4468200 | 1463900 | 549960 | 355800 |
| Q96H79;Q96H79-2 | Zinc finger CCCH-type antiviral protein 1-like | ZC3HAV1L | 3656500 | 1616400 | 1291400 | 1468600 | 993400 | 548670 |
| Q96HC4;Q96HC4-7;Q96HC4-4;Q96HC4-6;H0Y929;H0YBI4;H0Y8Y3;F5H7Y0;Q96HC4-3;Q96HC4-2 | PDZ and LIM domain protein 5 | PDLIM5 | 2397700 | 682970 | 640690 | 0 | 0 | 0 |
| Q96HJ9-2;Q9Y383-3;Q9Y383-2;Q9Y383 | Putative RNA-binding protein Luc7-like 2 | LUC7L2 | 1534700 | 817150 | 1525100 | 196660 | 479980 | 66651 |
| Q96HQ2-2;Q96HQ2 | CDKN2AIP N-terminal-like protein | CDKN2AIPNL | 0 | 0 | 0 | 518850 | 422670 | 0 |
| Q96IV0-5;Q96IV0-2;Q96IV0;H0Y2P2;Q96IV0-4;Q96IV0-3 | Peptide-N(4)-(N-acetyl-beta-glucosaminyl)asparagine amidase | NGLY1 | 90347 | 0 | 61656 | 0 | 0 | 0 |
| Q96IZ0 | PRKC apoptosis WT1 regulator protein | PAWR | 865740 | 479360 | 664270 | 0 | 0 | 0 |
| Q96KP1 | Exocyst complex component 2 | EXOC2 | 171460 | 40399 | 152210 | 31826 | 32281 | 17567 |
| Q96KR1;H0Y8W1 | Zinc finger RNA-binding protein | ZFR | 671600 | 596030 | 2059000 | 2406800 | 4730000 | 2544900 |
| Q96L92-3;Q96L92;Q96L92-2;Q96L92-4 | Sorting nexin-27 | SNX27 | 1326300 | 556620 | 566910 | 0 | 0 | 0 |
| Q96LB3;E5RGX6;E5RIF0;E5RH29;Q96LB3-2 | Intraflagellar transport protein 74 homolog | IFT74 | 58053 | 97406 | 359150 | 0 | 0 | 0 |
| Q96NL6;D6RBA6;D6RBP0;REV__Q9UHJ3-2;REV__Q9UHJ3;Q96NL6-3;Q96NL6-2 | Sodium channel and clathrin linker 1 | SCLT1 | 987720 | 360830 | 577950 | 0 | 0 | 0 |
| Q96P11-2;Q63ZY6-4;Q63ZY6-5;Q63ZY6;Q96P11-5;Q96P11;Q96P11-4 | Probable 28S rRNA (cytosine-C(5))-methyltransferase;Putative methyltransferase NSUN5C | NSUN5;NSUN5P2 | 303160 | 174740 | 485280 | 337860 | 926220 | 420510 |
| Q96PK6;Q96PK6-2;F8WDX3;B8ZZ74;Q96PK6-3;Q96PK6-4 | RNA-binding protein 14 | RBM14 | 605720 | 604320 | 1098100 | 539000 | 1497600 | 407010 |
| Q96QC0 | Serine/threonine-protein phosphatase 1 regulatory subunit 10 | PPP1R10 | 0 | 0 | 0 | 106100 | 140410 | 43505 |
| Q96QK1;I3L4S0;I3L4P4 | Vacuolar protein sorting-associated protein 35 | VPS35 | 5152800 | 1604400 | 3250300 | 103980 | 931960 | 280420 |
| Q96QR8 | Transcriptional activator protein Pur-beta | PURB | 1120300 | 621050 | 2244900 | 711740 | 1764800 | 846630 |
| Q96RS6-3;Q96RS6-2;Q96RS6 | NudC domain-containing protein 1 | NUDCD1 | 711610 | 167640 | 477540 | 100940 | 193850 | 0 |
| Q96S55-2;Q96S55;Q96S55-3;Q96S55-4 | ATPase WRNIP1 | WRNIP1 | 12312000 | 9006400 | 8521000 | 4217000 | 4563100 | 2506500 |
| Q96SB4;H3BLV9;Q5R363;B4DS61;Q96SB4-3;D6RBF8;H7C2I2;H7C521;A8MPP7;A8MPY5;E7ETV6;Q9UPE1-2;Q9UPE1-3;Q9UPE1-4;Q9UPE1 | SRSF protein kinase 1 | SRPK1 | 8992000 | 5705700 | 13004000 | 12239000 | 21104000 | 9063600 |
| Q96SI9-2;Q96SI9;H0YC91 | Spermatid perinuclear RNA-binding protein | STRBP | 66347 | 147320 | 188010 | 336940 | 651310 | 401950 |
| Q99436;Q5TBG5;Q99436-2 | Proteasome subunit beta type-7 | PSMB7 | 1841100000 | 1068900000 | 1171400000 | 1344300000 | 894590000 | 664840000 |
| Q99459 | Cell division cycle 5-like protein | CDC5L | 3880200 | 483100 | 5424700 | 14285000 | 15877000 | 6469200 |
| Q99460;Q99460-2;C9J9M4;F8WCE3 | 26S proteasome non-ATPase regulatory subunit 1 | PSMD1 | 305150000 | 211730000 | 227460000 | 44432000 | 41039000 | 26197000 |
| Q99536;K7EJM4;K7ERT7;Q99536-2;Q99536-3 | Synaptic vesicle membrane protein VAT-1 homolog | VAT1 | 1563900 | 0 | 0 | 0 | 0 | 0 |
| Q99543;Q99543-2;H7C3L7;C9IZ83 | DnaJ homolog subfamily C member 2;DnaJ homolog subfamily C member 2, N-terminally processed | DNAJC2 | 4593900 | 2378000 | 4318600 | 5303500 | 9873600 | 5291600 |
| Q99575;E5RK39 | Ribonucleases P/MRP protein subunit POP1 | POP1 | 879770 | 461600 | 1540300 | 1156600 | 2334100 | 1250100 |
| Q99584 | Protein S100-A13 | S100A13 | 3420200 | 705710 | 2716000 | 0 | 785230 | 826670 |
| Q99613-2;Q99613;B5ME19;H3BPE4;H3BPE3;H3BTY8 | Eukaryotic translation initiation factor 3 subunit C;Eukaryotic translation initiation factor 3 subunit C-like protein | EIF3C;EIF3CL | 19131000 | 8047500 | 21071000 | 19762000 | 31722000 | 16185000 |
| Q99614;H0YB37 | Tetratricopeptide repeat protein 1 | TTC1 | 927280 | 239540 | 854400 | 167120 | 274820 | 164070 |
| Q99714;Q5H928;Q99714-2 | 3-hydroxyacyl-CoA dehydrogenase type-2 | HSD17B10 | 314370 | 196550 | 552930 | 232070 | 1420300 | 437840 |
| Q99832-3;Q99832;F8WAM2;Q99832-4;Q99832-2 | T-complex protein 1 subunit eta | CCT7 | 4530700 | 1750300 | 2290300 | 3609900 | 1966200 | 1926100 |
| Q99996-5;Q99996-3;Q99996-2;Q99996;Q99996-6;A0A087WX84;H7BYL6;Q99996-4 | A-kinase anchor protein 9 | AKAP9 | 443540 | 355460 | 276400 | 83588 | 68851 | 0 |
| Q9BPX3 | Condensin complex subunit 3 | NCAPG | 37493 | 0 | 87927 | 0 | 0 | 0 |
| Q9BQ67;M0QX71 | Glutamate-rich WD repeat-containing protein 1 | GRWD1 | 6237600 | 2793500 | 6578800 | 4297800 | 8279400 | 3851800 |
| Q9BQA1;Q9BQA1-2;H0Y711 | Methylosome protein 50 | WDR77 | 2554700 | 1421100 | 2313000 | 599250 | 2330500 | 1120700 |
| Q9BQG0;Q9BQG0-2;I3L1L3;I3L2H8;I3L311 | Myb-binding protein 1A | MYBBP1A | 1616300 | 827620 | 2113900 | 3583800 | 6344600 | 2821700 |
| Q9BR76;A0A087WW53;F5H390;F5H0D2;A9Z1Z3-3;A9Z1Z3 | Coronin-1B | CORO1B | 15355000 | 9032400 | 9063200 | 277410 | 1309300 | 667440 |
| Q9BRP8;Q9BRP8-2 | Partner of Y14 and mago | WIBG | 25368000 | 15879000 | 35973000 | 23505000 | 42216000 | 22626000 |
| Q9BRS2;E9PFQ8 | Serine/threonine-protein kinase RIO1 | RIOK1 | 1883200 | 1118200 | 1258900 | 639550 | 1286500 | 529250 |
| Q9BRT6 | Protein LLP homolog | LLPH | 720330 | 666410 | 1319600 | 0 | 1426900 | 0 |
| Q9BRX2;REV__Q5JVM6;REV__Q96JX3 | Protein pelota homolog | PELO | 2022100 | 879380 | 2668000 | 862160 | 2112300 | 944670 |
| Q9BS26 | Endoplasmic reticulum resident protein 44 | ERP44 | 105920 | 108910 | 144410 | 0 | 0 | 0 |
| Q9BSJ8;Q9BSJ8-2;F8VZB1 | Extended synaptotagmin-1 | ESYT1 | 70810 | 79792 | 80376 | 0 | 14889 | 0 |
| Q9BTE3-2;Q9BTE3 | Mini-chromosome maintenance complex-binding protein | MCMBP | 205410 | 60082 | 428190 | 105110 | 126520 | 0 |
| Q9BTT0;E9PPH5;Q5TB19;E9PLC4;E9PI45;Q5TB25;Q9BTT0-2 | Acidic leucine-rich nuclear phosphoprotein 32 family member E | ANP32E | 6609500 | 5836200 | 8591300 | 2565200 | 2337800 | 1593800 |
| U3KQC1;Q9BV38;K7EIR0;K7EML4 | WD repeat-containing protein 18 | WDR18 | 1402800 | 775150 | 2504800 | 1406800 | 2237100 | 958120 |
| Q9BVA1 | Tubulin beta-2B chain | TUBB2B | 445220 | 130830 | 209970 | 233680 | 476730 | 0 |
| Q9BVJ6-3;Q9BVJ6;Q9BVJ6-2 | U3 small nucleolar RNA-associated protein 14 homolog A | UTP14A | 61857 | 0 | 141420 | 204000 | 291750 | 102170 |
| Q9BVK6 | Transmembrane emp24 domain-containing protein 9 | TMED9 | 292560 | 92761 | 362540 | 0 | 0 | 0 |
| Q9BVP2-2;Q9BVP2;C9JZT7;B4DMU5;C9JYH9 | Guanine nucleotide-binding protein-like 3 | GNL3 | 7936200 | 4118800 | 7597600 | 6714100 | 15600000 | 4809800 |
| Q9BVS4-2;Q9BVS4;H0Y919 | Serine/threonine-protein kinase RIO2 | RIOK2 | 270220 | 285090 | 707960 | 0 | 110200 | 88781 |
| Q9BW19 | Kinesin-like protein KIFC1 | KIFC1 | 0 | 0 | 58272 | 0 | 91388 | 13744 |
| Q9BWF3;E9PB51;Q9BQ04;Q9BWF3-4;E9PLB0;E9PM61;D6R9K7;U3KQD5;J3QRR5;Q9BWF3-3;Q9BWF3-2 | RNA-binding protein 4;RNA-binding protein 4B | RBM4;RBM4B | 7621300 | 3036100 | 8503700 | 2172400 | 6383300 | 2330100 |
| Q9BWJ5 | Splicing factor 3B subunit 5 | SF3B5 | 1515300 | 0 | 0 | 0 | 0 | 0 |
| Q9BX10-2;Q9BX10-4;Q9BX10;X6RJ09;H0Y7A5;H0Y4M5;H0Y5J1;Q9BX10-3 | GTP-binding protein 2 | GTPBP2 | 160010 | 117420 | 228090 | 90241 | 204130 | 178180 |
| Q9BXJ9;Q9BXJ9-4 | N-alpha-acetyltransferase 15, NatA auxiliary subunit | NAA15 | 1381600 | 546980 | 3033000 | 186430 | 685580 | 290960 |
| Q9BXP5-5;Q9BXP5-4;Q9BXP5-2;Q9BXP5-3;Q9BXP5;H7C3A1;H7C1K0;H7C0U8 | Serrate RNA effector molecule homolog | SRRT | 8754300 | 5791600 | 13068000 | 2252800 | 3587800 | 1489900 |
| Q9BXR0-2;Q9BXR0 | Queuine tRNA-ribosyltransferase | QTRT1 | 258570 | 350010 | 685270 | 449410 | 682780 | 338850 |
| Q9BXW9;Q9BXW9-1;Q9BXW9-3;H7BZJ7;Q9BXW9-4 | Fanconi anemia group D2 protein | FANCD2 | 882100 | 449600 | 1555200 | 1807100 | 3183600 | 1694500 |
| Q9BY44-3;Q9BY44;F8WAE5;Q9BY44-4;C9IZE1;Q9BY44-2;H7C5Q3;H7C5R5;F8WF18;F8WAT3 | Eukaryotic translation initiation factor 2A;Eukaryotic translation initiation factor 2A, N-terminally processed | EIF2A | 4097200 | 2594200 | 6220100 | 4951200 | 9975600 | 4210800 |
| Q9BYG3;H7BZL0 | MKI67 FHA domain-interacting nucleolar phosphoprotein | NIFK | 0 | 0 | 661170 | 396940 | 1082200 | 1773800 |
| Q9BZE4;Q9BZE4-2;Q9BZE4-3;Q5T3R7;H0YKC1;REV__B7ZC06;H0YLK7;REV__Q08379;REV__Q68E01-2;REV__Q68E01;C4B7M2;Q9NVE4;Q96JI7-2;Q96JI7-3;Q96JI7 | Nucleolar GTP-binding protein 1 | GTPBP4 | 3497300 | 2795100 | 5362500 | 6699000 | 14636000 | 8047100 |
| Q9BZH6;S4R3Z0 | WD repeat-containing protein 11 | WDR11 | 440440 | 246710 | 385880 | 133720 | 101100 | 50387 |
| Q9BZI7-2;Q9BZI7;A0A087WYC8 | Regulator of nonsense transcripts 3B | UPF3B | 375810 | 477210 | 2063300 | 951790 | 2642100 | 1413800 |
| Q9BZL1 | Ubiquitin-like protein 5 | UBL5 | 16571000 | 7532700 | 10427000 | 5167000 | 6653300 | 3844300 |
| Q9BZL4-2;Q9BZL4-3;Q9BZL4-4;Q9BZL4;K7EL81;Q9BZL4-5 | Protein phosphatase 1 regulatory subunit 12C | PPP1R12C | 660980 | 166320 | 1041000 | 432170 | 1086200 | 731320 |
| Q9C005 | Protein dpy-30 homolog | DPY30 | 6586900 | 3435000 | 5532700 | 2819600 | 3783200 | 2420300 |
| Q9GZL7 | Ribosome biogenesis protein WDR12 | WDR12 | 1854000 | 704210 | 1895100 | 2983300 | 4585800 | 2058800 |
| Q9GZP4-2;Q9GZP4;X6RHB9;X6R8S9 | PITH domain-containing protein 1 | PITHD1 | 1944700 | 1333300 | 2413500 | 0 | 151110 | 58169 |
| Q9GZS1-2;E7EX70;Q9GZS1 | DNA-directed RNA polymerase I subunit RPA49 | POLR1E | 6004200 | 6031400 | 9571300 | 2881400 | 2669300 | 1629000 |
| Q9GZU8;H3BP64;H3BUL4;H3BMX9;H3BTP8;H3BSY6;H3BQQ6;H3BTI2;H3BU93;Q6P4H7;H3BSF0;H3BNK9;H3BN22;H3BPH9;H3BPF9 | Protein FAM192A | FAM192A | 33053000 | 17128000 | 19410000 | 10035000 | 20879000 | 7865000 |
| Q9H0A0;Q9H0A0-2;A0A087WV29;E9PJN6;E9PMU0 | N-acetyltransferase 10 | NAT10 | 532250 | 480670 | 1101100 | 1444900 | 3306500 | 1418000 |
| Q9H0D6-2;Q9H0D6 | 5-3 exoribonuclease 2 | XRN2 | 657360 | 263640 | 1412500 | 717810 | 2553400 | 1057100 |
| Q9H0S4-2;Q9H0S4;F5H1N9 | Probable ATP-dependent RNA helicase DDX47 | DDX47 | 960240 | 1463300 | 2072100 | 495060 | 3496100 | 881760 |
| Q9H1I8;B1AH59;B1AH60;Q9H1I8-3;Q9H1I8-2 | Activating signal cointegrator 1 complex subunit 2 | ASCC2 | 590040 | 728610 | 9998100 | 259010 | 1039100 | 463460 |
| Q9H2H8;B8ZZ77;H7BZ14;Q9H2H8-2;C9K058;F8WC82;C9J4N2 | Peptidyl-prolyl cis-trans isomerase-like 3;Peptidyl-prolyl cis-trans isomerase | PPIL3 | 232980 | 645520 | 1439600 | 259190 | 708450 | 433170 |
| Q9H2U1-3;Q9H2U1-2;Q9H2U1;E7EWK3;H7C5F5 | ATP-dependent RNA helicase DHX36 | DHX36 | 3313100 | 1707200 | 1429100 | 1312500 | 1468500 | 266700 |
| Q9H3G5;H7C0X5;A0A087WVP6;C9J6L4;H7C218;C9JI22;A0A087WXJ9;C9JLV0 | Probable serine carboxypeptidase CPVL | CPVL | 0 | 60425 | 53702 | 0 | 0 | 0 |
| Q9H3P7 | Golgi resident protein GCP60 | ACBD3 | 0 | 0 | 109910 | 0 | 0 | 0 |
| Q9H3U1-2;Q9H3U1 | Protein unc-45 homolog A | UNC45A | 41858 | 0 | 125550 | 0 | 0 | 0 |
| Q9H444 | Charged multivesicular body protein 4b | CHMP4B | 2595800 | 118530 | 2680600 | 0 | 0 | 70269 |
| Q9H4L4;J3QRE4;A0A087WV50;J3QL36;A0A087X0R7 | Sentrin-specific protease 3 | SENP3;SENP3-EIF4A1 | 484600 | 144910 | 794590 | 223600 | 828010 | 355340 |
| Q9H6R4-2;Q9H6R4-4;Q9H6R4;Q9H6R4-3 | Nucleolar protein 6 | NOL6 | 0 | 0 | 0 | 128670 | 162700 | 121850 |
| Q9H6S0;D6RA70;D6R9T8;D6RF50 | Probable ATP-dependent RNA helicase YTHDC2 | YTHDC2 | 2547300 | 1306700 | 2785400 | 1883700 | 3384700 | 1818500 |
| Q9H6T3-3;Q9H6T3-2;Q9H6T3 | RNA polymerase II-associated protein 3 | RPAP3 | 1232500 | 675530 | 1170500 | 450350 | 1117600 | 598040 |
| Q9H7B2;Q5VXN0 | Ribosome production factor 2 homolog | RPF2 | 199240 | 0 | 0 | 276310 | 605980 | 0 |
| Q9H7E9;Q9H7E9-2 | UPF0488 protein C8orf33 | C8orf33 | 36034000 | 6310900 | 31316000 | 20733000 | 31895000 | 15212000 |
| Q9H814 | Phosphorylated adapter RNA export protein | PHAX | 7982200 | 3732700 | 7767700 | 312700 | 3906100 | 967810 |
| Q9H857;Q9H857-2;H7C519;Q9H857-4;Q9H857-3 | 5-nucleotidase domain-containing protein 2 | NT5DC2 | 844720 | 771140 | 1004000 | 0 | 121120 | 0 |
| Q9H9A5-4;Q9H9A5-3;Q9H9A5-2;Q9H9A5;Q9H9A5-6;H7C1X9;E9PCN5;H7C0K1;Q9H9A5-5 | CCR4-NOT transcription complex subunit 10 | CNOT10 | 248400 | 132280 | 335650 | 127710 | 217950 | 101050 |
| Q9H9Y6;Q9H9Y6-3;Q9H9Y6-5;Q9H9Y6-2;Q9H9Y6-4;H7C0D9;C9JJG2;C9JS83;F8WDS4;F8WAK7;F8WBB9;F8WCS3 | DNA-directed RNA polymerase I subunit RPA2 | POLR1B | 949410 | 515830 | 1875200 | 411540 | 807650 | 425130 |
| Q9HAU5;Q9HAU5-2 | Regulator of nonsense transcripts 2 | UPF2 | 0 | 0 | 0 | 0 | 0 | 0 |
| Q9HAV4;H0Y3Q8 | Exportin-5 | XPO5 | 186600 | 90951 | 217140 | 39976 | 66483 | 0 |
| Q9HB71;B2ZWH1;Q9HB71-2;Q9HB71-3 | Calcyclin-binding protein | CACYBP | 47743000 | 2634600 | 25278000 | 28670000 | 14962000 | 0 |
| Q9HC35-2;Q9HC35;B5MBZ0 | Echinoderm microtubule-associated protein-like 4 | EML4 | 85258 | 30179 | 189410 | 163170 | 553800 | 363240 |
| Q9HCN4;Q9HCN4-3;Q9HCN4-2;Q9HCN4-4;Q9HCN4-5;F8WD09 | GPN-loop GTPase 1 | GPN1 | 2705000 | 1043700 | 646980 | 800340 | 2998400 | 1900500 |
| Q9HCS7 | Pre-mRNA-splicing factor SYF1 | XAB2 | 437600 | 118370 | 231310 | 60473 | 225700 | 46866 |
| Q9NP77 | RNA polymerase II subunit A C-terminal domain phosphatase SSU72 | SSU72 | 208130 | 191400 | 496260 | 0 | 259780 | 133620 |
| Q9NPD3;E9PI41;E9PPI9 | Exosome complex component RRP41 | EXOSC4 | 9047000 | 4485500 | 7776100 | 4469500 | 8377800 | 3583500 |
| Q9NPI6-2;Q9NPI6 | mRNA-decapping enzyme 1A | DCP1A | 0 | 0 | 0 | 30778 | 169310 | 67111 |
| Q9NQT5-2;Q9NQT5 | Exosome complex component RRP40 | EXOSC3 | 6511100 | 5229800 | 9047800 | 4757700 | 13083000 | 5384100 |
| Q9NQZ2 | Something about silencing protein 10 | UTP3 | 0 | 0 | 0 | 392560 | 812960 | 549530 |
| Q9NR12;D6RF83;H7BYK4;D6RH06;Q9NR12-3;Q9NR12-5;Q9NR12-6;Q9NR12-4;Q9NR12-2 | PDZ and LIM domain protein 7 | PDLIM7 | 0 | 52596 | 73514 | 0 | 0 | 0 |
| Q9NR30;Q9NR30-2 | Nucleolar RNA helicase 2 | DDX21 | 30857000 | 18035000 | 48185000 | 39677000 | 72242000 | 38331000 |
| Q9NR33 | DNA polymerase epsilon subunit 4 | POLE4 | 0 | 686170 | 2489800 | 346260 | 1180100 | 577290 |
| Q9NR50-3;Q9NR50-2;Q9NR50;Q5QP88;H0Y580 | Translation initiation factor eIF-2B subunit gamma | EIF2B3 | 838740 | 371810 | 852300 | 869060 | 1219900 | 0 |
| Q9NRX1;F8WBJ6 | RNA-binding protein PNO1 | PNO1 | 6891900 | 4132800 | 9574500 | 2955500 | 5018700 | 2315400 |
| Q9NSD9;Q9NSD9-2;REV__Q6ZSS7 | Phenylalanine--tRNA ligase beta subunit | FARSB | 20062000 | 12858000 | 25203000 | 28689000 | 42616000 | 23862000 |
| Q9NTI5-2;Q9NTI5;A9IYQ1;Q9NTI5-4;Q9NTI5-3 | Sister chromatid cohesion protein PDS5 homolog B | PDS5B | 155920 | 92109 | 175100 | 330050 | 584660 | 262570 |
| Q9NU22;Q5T795 | Midasin | MDN1 | 171290 | 112060 | 395110 | 91256 | 268810 | 62841 |
| Q9NUD5 | Zinc finger CCHC domain-containing protein 3 | ZCCHC3 | 111310 | 0 | 0 | 0 | 0 | 0 |
| Q9NUQ3;Q9NUQ3-2 | Gamma-taxilin | TXLNG | 4820200 | 2290200 | 4283900 | 4089600 | 5034200 | 2375500 |
| Q9NV70-2;Q9NV70 | Exocyst complex component 1 | EXOC1 | 294890 | 121750 | 318690 | 97460 | 88335 | 47049 |

| | | | | | | | | |
|---|---|---|---|---|---|---|---|---|
| Q9NVI1;Q9NVI1-2;Q9NVI1-1;F8W7R3;H3BP78;H3BS60;H3BQE2;H3BT54;H3BMG4;Q9NVI1-4 | Fanconi anemia group I protein | FANCI | 1723800 | 2746200 | 2150900 | 1368400 | 1629300 | 807260 |
| Q9NVM9-2;Q9NVM9;H0YIA9;H0YI01;H0YH12 | Protein asunder homolog | ASUN | 305330 | 127100 | 125190 | 45499 | 41554 | 0 |
| Q9NVP1 | ATP-dependent RNA helicase DDX18 | DDX18 | 543770 | 248880 | 289560 | 832440 | 1400700 | 799610 |
| Q9NW13-2;Q9NW13;C9JAA9;H7C5G8;C9JE21 | RNA-binding protein 28 | RBM28 | 530720 | 338010 | 1664700 | 2703000 | 4230000 | 1837400 |
| Q9NWT1 | p21-activated protein kinase-interacting protein 1 | PAK1IP1 | 165070 | 231840 | 273620 | 3277700 | 4829700 | 2004700 |
| Q9NWX6 | Probable tRNA(His) guanylyltransferase | THG1L | 0 | 315210 | 791490 | 1018400 | 2085400 | 643940 |
| Q9NWZ8;H7BYE2 | Gem-associated protein 8 | GEMIN8 | 303020 | 384490 | 1402800 | 1261500 | 1724700 | 1013100 |
| Q9NX24;D6RC52;D6RCB9;H0YC83;J3QSY4 | H/ACA ribonucleoprotein complex subunit 2 | NHP2 | 1348300 | 995220 | 3016000 | 4219900 | 2527300 | 2532500 |
| Q9NX58 | Cell growth-regulating nucleolar protein | LYAR | 4095400 | 2776700 | 6507400 | 11377000 | 13993000 | 7393900 |
| Q9NXG2;H3BNW0 | THUMP domain-containing protein 1 | THUMPD1 | 127010 | 0 | 0 | 1152500 | 1694400 | 801470 |
| Q9NXV6;J3KNE1;D6RGD2 | CDKN2A-interacting protein | CDKN2AIP | 343600 | 159240 | 913810 | 1372400 | 3157500 | 1303500 |
| Q9NYB0 | Telomeric repeat-binding factor 2-interacting protein 1 | TERF2IP | 1038000 | 775150 | 1814500 | 821750 | 1084500 | 489440 |
| Q9NYU2-2;Q9NYU2;F8WCI2;F8WCE6 | UDP-glucose:glycoprotein glucosyltransferase 1 | UGGT1 | 259850 | 134550 | 105830 | 0 | 0 | 0 |
| Q9NZB2;Q9NZB2-6;Q9NZB2-4;Q9NZB2-2;Q9NZB2-5;Q5T035 | Constitutive coactivator of PPAR-gamma-like protein 1 | FAM120A | 2974200 | 1846700 | 4927000 | 2862400 | 7306900 | 3046600 |
| Q9NZI8;Q9NZI8-2;REV__M0QZ54;REV__M0R118;REV__M0R0M7;G8JLG2;REV__A0A087WWS7;REV__Q15833;Q15517 | Insulin-like growth factor 2 mRNA-binding protein 1 | IGF2BP1 | 31564000 | 18080000 | 47394000 | 37074000 | 76350000 | 39406000 |
| Q9NZM5 | Glioma tumor suppressor candidate region gene 2 protein | GLTSCR2 | 0 | 0 | 0 | 140170 | 386270 | 208100 |
| Q9NZT1 | Calmodulin-like protein 5 | CALML5 | 8015400 | 15085000 | 9297400 | 3027900 | 4819600 | 4318100 |
| Q9NZZ3-2;Q9NZZ3 | Charged multivesicular body protein 5 | CHMP5 | 355180 | 182860 | 0 | 0 | 0 | 8348400 |
| Q9P000-2;Q9P000 | COMM domain-containing protein 9 | COMMD9 | 485540 | 118260 | 373720 | 0 | 0 | 0 |
| Q9P031 | Thyroid transcription factor 1-associated protein 26 | CCDC59 | 942870 | 317920 | 648470 | 883620 | 1361700 | 210100 |
| Q9P0P0;C9J1C6;F8WBU5;C9J391;H7C426 | E3 ubiquitin-protein ligase RNF181 | RNF181 | 52583000 | 25112000 | 31832000 | 13495000 | 15370000 | 11035000 |
| Q9P258 | Protein RCC2 | RCC2 | 1673800 | 865580 | 1855900 | 285340 | 1802900 | 516810 |
| Q9P265;Q14689-3;Q14689-2;Q14689-4;Q14689-5;Q14689-6;Q14689 | Disco-interacting protein 2 homolog B | DIP2B | 343120 | 140520 | 259430 | 105350 | 214390 | 84263 |
| Q9P2J5;Q9P2J5-2;Q9P2J5-3;A0A087WXY1 | Leucine--tRNA ligase, cytoplasmic | LARS | 5769300 | 3515100 | 8048000 | 4086100 | 8076200 | 3945900 |
| Q9UBB4-2;Q9UBB4;B1AHE3;B1AHE4;H0Y7N2 | Ataxin-10 | ATXN10 | 2361500 | 1090200 | 2315100 | 546610 | 1136400 | 333770 |
| Q9UBV8 | Peflin | PEF1 | 1625100 | 1339900 | 1791200 | 0 | 0 | 0 |
| Q9UG63;Q9UG63-2;C9JHK9;C9JZV3 | ATP-binding cassette sub-family F member 2 | ABCF2 | 9731200 | 4565200 | 13240000 | 12044000 | 20174000 | 8567900 |
| Q9UGR2-2;Q9UGR2 | Zinc finger CCCH domain-containing protein 7B | ZC3H7B | 0 | 0 | 0 | 0 | 0 | 0 |
| Q9UHB9;Q9UHB9-4;Q9UHB9-2;Q9UHB9-3 | Signal recognition particle subunit SRP68 | SRP68 | 2269300 | 1150000 | 3241400 | 2979900 | 4998700 | 2383900 |
| Q9UHD2;F5GZI4;F5H206;F5H1A3 | Serine/threonine-protein kinase TBK1 | TBK1 | 57459 | 82450 | 99799 | 0 | 0 | 0 |
| Q9UHD9 | Ubiquilin-2 | UBQLN2 | 1644600 | 2189300 | 2721700 | 526000 | 690200 | 472760 |
| Q9UHI6;E9PJ60;Q9UHI6-2 | Probable ATP-dependent RNA helicase DDX20 | DDX20 | 613310 | 254540 | 873450 | 498890 | 1121400 | 665000 |
| Q9UHR4 | Brain-specific angiogenesis inhibitor 1-associated protein 2-like protein 1 | BAIAP2L1 | 216600 | 23461 | 206280 | 0 | 0 | 0 |
| Q9UHV9 | Prefoldin subunit 2 | PFDN2 | 1731200 | 1043600 | 2200800 | 586580 | 0 | 460700 |
| Q9UHX1-4;Q9UHX1-6;Q9UHX1-3;Q9UHX1-5;Q9UHX1-2;Q9UHX1;E9PQ56;H0YEM1;E9PL19;E9PMU7;E9PN18;H0YCP8 | Poly(U)-binding-splicing factor PUF60 | PUF60 | 9634900 | 4880300 | 7402900 | 6530100 | 6408300 | 2962900 |
| Q9UI10;E7ERK9;Q9UI10-3;A0A087WTA5;Q9UI10-2;F8W8L6;A0A087WW69;F8WEV6;H7C2L8 | Translation initiation factor eIF-2B subunit delta | EIF2B4 | 2064400 | 1548600 | 3632600 | 1836700 | 4306400 | 1803600 |
| Q9UII4;E9PBL0;Q14393-5;Q14393-4;Q14393-3;Q14393-2;Q14393 | E3 ISG15--protein ligase HERC5 | HERC5 | 165700 | 205030 | 785760 | 158340 | 893190 | 398660 |
| Q9UJA5;Q9UJA5-4;Q9UJA5-3;Q9UJA5-2 | tRNA (adenine(58)-N(1))-methyltransferase non-catalytic subunit TRM6 | TRMT6 | 1613700 | 1081100 | 1922200 | 2497700 | 7072000 | 3403100 |
| Q9UJX2-3;Q9UJX2 | Cell division cycle protein 23 homolog | CDC23 | 1327300 | 654580 | 957540 | 439510 | 568110 | 131440 |
| Q9UJX3-2;Q9UJX3 | Anaphase-promoting complex subunit 7 | ANAPC7 | 150480 | 62853 | 107140 | 0 | 58606 | 0 |
| Q9UK59;Q9UK59-2;F8WAY1 | Lariat debranching enzyme | DBR1 | 2923800 | 1753300 | 3030100 | 1798200 | 2356800 | 1420800 |
| Q9UKD2 | mRNA turnover protein 4 homolog | MRTO4 | 7765200 | 4566300 | 8970800 | 4827700 | 10578000 | 5374200 |
| Q9UKN8 | General transcription factor 3C polypeptide 4 | GTF3C4 | 253590 | 243390 | 532150 | 78341 | 36133 | 19740 |
| Q9UKS6;E9PIY1;E9PJ75;E9PNM9 | Protein kinase C and casein kinase substrate in neurons protein 3 | PACSIN3 | 1140000 | 274160 | 861510 | 1258100 | 2802100 | 1064800 |
| Q9UKV8;Q9UKV8-2;Q5TA58;Q9UL18 | Protein argonaute-2 | AGO2 | 936890 | 364570 | 816750 | 604880 | 951280 | 365750 |
| Q9UL25 | Ras-related protein Rab-21 | RAB21 | 383590 | 0 | 357310 | 0 | 0 | 0 |
| Q9UL42 | Paraneoplastic antigen Ma2 | PNMA2 | 995090 | 3265200 | 7164900 | 1415000 | 1131800 | 869170 |
| Q9UL54-2;Q9UL54-4;Q9UL54 | Serine/threonine-protein kinase TAO2 | TAOK2 | 0 | 0 | 0 | 0 | 0 | 0 |
| Q9ULR0;Q9ULR0-2;Q9ULR0-1 | Pre-mRNA-splicing factor ISY1 homolog | ISY1 | 0 | 103140 | 333460 | 345980 | 412470 | 235250 |
| Q9ULV4;Q9ULV4-2;Q9ULV4-3;B4E3S0;F8W1H8;F8VUX3;F8VSA4;F8VRE9;F8VTT6;F8VVB7;H0YHL7 | Coronin-1C;Coronin | CORO1C | 1962600 | 1146400 | 1477500 | 0 | 61446 | 29554 |
| Q9ULX3;H3BUR4 | RNA-binding protein NOB1 | NOB1 | 9454900 | 4865100 | 10786000 | 3720200 | 7376900 | 2629400 |
| Q9UM54-6;E7EW20;Q9UM54-5;Q9UM54-2;Q9UM54-4;Q9UM54-1;Q9UM54 | Unconventional myosin-VI | MYO6 | 561600 | 639810 | 685330 | 0 | 0 | 0 |
| Q9UMS4;H0YGF3;F5GY56;F5H2I0;H0YGZ5 | Pre-mRNA-processing factor 19 | PRPF19 | 13094000 | 6492200 | 10530000 | 9125300 | 15328000 | 8304000 |
| Q9UMX0-2;Q9UMX0 | Ubiquilin-1 | UBQLN1 | 13454000 | 6727400 | 10899000 | 1194000 | 1287700 | 671730 |
| Q9UN81 | LINE-1 retrotransposable element ORF1 protein | L1RE1 | 236800 | 102690 | 443260 | 195520 | 501420 | 228120 |
| Q9UN86-2;Q9UN86;D6R9X5;D6R9A4;D6RBM9;D6REX8;D6RBR0;D6RE13;D6RBW8;D6RGJ4;D6RAC7;D6RB17 | Ras GTPase-activating protein-binding protein 2 | G3BP2 | 5056100 | 2562000 | 7542100 | 5143600 | 10267000 | 4691900 |
| Q9UNE7-2;Q9UNE7;H3BS86;H3BTA3;H3BUD0 | E3 ubiquitin-protein ligase CHIP | STUB1 | 1297900 | 395310 | 1449700 | 394410 | 849010 | 510490 |
| Q9UNF0-2;Q9UNF0;B0QYG7;B0QYG8 | Protein kinase C and casein kinase substrate in neurons protein 2 | PACSIN2 | 2502200 | 1222500 | 2807400 | 717520 | 763260 | 383760 |
| Q9UNM6;Q9UNM6-2;J3KNQ3;A0A087WUL9;E9PL38;H0YD73;E9PQG3;E9PPD2;F8W8J6 | 26S proteasome non-ATPase regulatory subunit 13 | PSMD13 | 649330000 | 397650000 | 529090000 | 63511000 | 67175000 | 40540000 |
| Q9UNN5;B1ANM7;Q9UNN5-2;B3KT28 | FAS-associated factor 1 | FAF1 | 2331100 | 746530 | 1889800 | 406060 | 558530 | 117530 |
| Q9UNQ2;D6RCL3 | Probable dimethyladenosine transferase | DIMT1 | 1276300 | 913080 | 3138500 | 1952800 | 4740600 | 3272100 |
| Q9UNX4 | WD repeat-containing protein 3 | WDR3 | 371620 | 283160 | 524280 | 2240100 | 2899200 | 1153500 |
| Q9UQ35;I3L3Q8;Q9UQ35-2 | Serine/arginine repetitive matrix protein 2 | SRRM2 | 0 | 64567 | 101310 | 35640 | 190800 | 79958 |
| Q9UQ80;F8VR77;H0YIN7;F8W0A3;F8VZ69 | Proliferation-associated protein 2G4 | PA2G4 | 132990000 | 69916000 | 156190000 | 141480000 | 258690000 | 147030000 |
| Q9UQE7;E7EPJ2;A0A087WVF7;Q9NZM3-4;Q9NZM3-3;Q9NZM3-2;Q9NZM3 | Structural maintenance of chromosomes protein 3 | SMC3 | 2872800 | 2330600 | 5390900 | 6098200 | 11112000 | 4734600 |
| Q9Y224;G3V4C6;H0YJB9;G3V4E7;P53708 | UPF0568 protein C14orf166 | C14orf166 | 11377000 | 7317000 | 18040000 | 21661000 | 29971000 | 14934000 |
| Q9Y230;Q9Y230-2;M0R0Y3;X6R2L4;M0R0Z0;M0QXI6;M0QXZ7 | RuvB-like 2 | RUVBL2 | 55142000 | 21611000 | 36584000 | 20910000 | 21444000 | 9627400 |
| Q9Y244 | Proteasome maturation protein | POMP | 83398000 | 38772000 | 40033000 | 16038000 | 10865000 | 6924500 |
| Q9Y265;Q9Y265-2;E7ETR0;H7C4G5;J3QLR1 | RuvB-like 1 | RUVBL1 | 36424000 | 20545000 | 27936000 | 13234000 | 17036000 | 8178400 |
| Q9Y266 | Nuclear migration protein nudC | NUDC | 3532300 | 1831000 | 3468500 | 1034000 | 1506300 | 507620 |
| Q9Y281;Q9Y281-3;F8WDN3;G3V2U0 | Cofilin-2 | CFL2 | 45191000 | 7041700 | 13607000 | 0 | 530660 | 245890 |
| Q9Y285;K7ER00;Q9Y285-2;K7ER16;K7EPH2;K7EK06;K7ERS0 | Phenylalanine--tRNA ligase alpha subunit | FARSA | 8038300 | 6541500 | 17328000 | 9589500 | 23780000 | 8493600 |
| Q9Y295;J3QRI9;J3QKW7;A8MZF9;P55039 | Developmentally-regulated GTP-binding protein 1 | DRG1 | 3063100 | 1957600 | 6013200 | 2707200 | 5321800 | 2365700 |
| Q9Y2A7;Q9Y2A7-2 | Nck-associated protein 1 | NCKAP1 | 1111200 | 748530 | 1983500 | 236420 | 259840 | 94793 |
| Q9Y2L1;G3V1J5;Q9Y2L1-2;F2Z2C0 | Exosome complex exonuclease RRP44 | DIS3 | 1488300 | 1025800 | 1754500 | 1237100 | 3548300 | 1482300 |
| Q9Y2P8;Q5VYW8 | RNA 3-terminal phosphate cyclase-like protein | RCL1 | 0 | 50833 | 226220 | 105550 | 1006000 | 567350 |
| Q9Y2R4 | Probable ATP-dependent RNA helicase DDX52 | DDX52 | 420900 | 0 | 308380 | 523260 | 800770 | 84151 |
| Q9Y2S6 | Translation machinery-associated protein 7 | TMA7 | 20036000 | 14051000 | 32265000 | 16478000 | 48950000 | 21311000 |
| Q9Y2X3;F8WED0;H7BZ72 | Nucleolar protein 58 | NOP58 | 364680 | 255760 | 846880 | 661050 | 1339200 | 366400 |
| Q9Y333 | U6 snRNA-associated Sm-like protein LSm2 | LSM2 | 859990 | 844320 | 2272300 | 458960 | 1657200 | 0 |
| Q9Y3B4 | Splicing factor 3B subunit 6 | SF3B6 | 2664500 | 1682400 | 1717300 | 4599800 | 2209900 | 813520 |
| Q9Y3C1;D6RIC3;A0A087X2G6;Q6PIM0;Q9Y3C1-2;D6RC60;A0A087WU10;A0A087X1T1 | Nucleolar protein 16 | NOP16 | 1859300 | 922700 | 604770 | 4867900 | 8580400 | 3506000 |
| Q9Y3F4;Q9Y3F4-2;H0YH33 | Serine-threonine kinase receptor-associated protein | STRAP | 13713000 | 4237600 | 13028000 | 2375000 | 6306100 | 2645200 |
| Q9Y3I0 | tRNA-splicing ligase RtcB homolog | RTCB | 6533800 | 4293000 | 8160400 | 5830400 | 10839000 | 5686500 |
| Q9Y3I1-3;Q9Y3I1-2;Q9Y3I1 | F-box only protein 7 | FBXO7 | 10803000 | 8969800 | 7969200 | 1918100 | 3525600 | 2629900 |
| Q9Y3T9 | Nucleolar complex protein 2 homolog | NOC2L | 653780 | 342260 | 988560 | 1890400 | 2951600 | 1710900 |
| Q9Y3U8;J3QSB5;J3KTD3 | 60S ribosomal protein L36 | RPL36 | 103070000 | 33860000 | 65838000 | 28642000 | 65266000 | 21073000 |
| Q9Y3Z3-4;Q9Y3Z3;Q9Y3Z3-3;Q9Y3Z3-2 | Deoxynucleoside triphosphate triphosphohydrolase SAMHD1 | SAMHD1 | 991810 | 972770 | 3359700 | 344750 | 1148300 | 283860 |
| Q9Y450-4;Q9Y450;B7Z524;H0YDX7;J3QT46;H0YES5;E9PJ90;E9PN23;E9PS53;G5E991;E9PMN1;E9PLR4;E9PHZ9;Q9Y450-2 | HBS1-like protein | HBS1L | 998390 | 542210 | 1176400 | 926240 | 1216300 | 1267000 |
| Q9Y490 | Talin-1 | TLN1 | 120540 | 23354 | 70051 | 44708 | 36469 | 17231 |
| Q9Y4C2-2;Q9Y4C2;C9K0E7;C9J9F4;C9JD13 | TRPM8 channel-associated factor 1 | TCAF1 | 4954100 | 2452300 | 3539100 | 510950 | 560050 | 366070 |
| Q9Y4C8 | Probable RNA-binding protein 19 | RBM19 | 0 | 0 | 0 | 0 | 0 | 0 |
| Q9Y4E8;Q9Y4E8-2;Q9Y4E8-3;Q9Y4E8-4;F8W0H4;H0YI31;F8VVY7;F8VZG8;H0YHM4;H7C189;Q13107-2;Q13107 | Ubiquitin carboxyl-terminal hydrolase 15 | USP15 | 4605500 | 2147400 | 2973200 | 273940 | 194760 | 149720 |
| Q9Y4W2-2;Q9Y4W2;Q9Y4W2-4;Q9Y4W2-3 | Ribosomal biogenesis protein LAS1L | LAS1L | 588380 | 0 | 92825 | 307130 | 286990 | 74223 |
| Q9Y4X5;H3BNB9;A0A087WT96;H3BSK4 | E3 ubiquitin-protein ligase ARIH1 | ARIH1 | 1134800 | 704020 | 1012900 | 641670 | 956420 | 579690 |
| U3KQS7;U3KQK1;V9GZ56;Q9Y4Z0;M0QXB0 | U6 snRNA-associated Sm-like protein LSm4 | LSM4 | 2686200 | 1674100 | 3667700 | 3331500 | 4428800 | 799110 |
| Q9Y5B9;G3V5A4;G3V401;G3V2X0 | FACT complex subunit SPT16 | SUPT16H | 119990 | 84855 | 376790 | 17812 | 18202 | 45477 |
| Q9Y5J9;E9PIR3;G3XAN8 | Mitochondrial import inner membrane translocase subunit Tim8 B | TIMM8B | 4094900 | 3766900 | 7196600 | 0 | 0 | 0 |
| Q9Y5K5-2;Q9Y5K5-4;Q9Y5K5-3;Q9Y5K5;Q5LJA5;Q5LJA9;Q5LJB0;H0Y6Y4;H0Y4E0;H0Y4K0;H0Y636 | Ubiquitin carboxyl-terminal hydrolase isozyme L5;Ubiquitin carboxyl-terminal hydrolase | UCHL5 | 183240000 | 109630000 | 143290000 | 17245000 | 14600000 | 9264800 |
| Q9Y5K6 | CD2-associated protein | CD2AP | 495750 | 101060 | 614030 | 0 | 0 | 0 |
| Q9Y5S2;H0YLY0 | Serine/threonine-protein kinase MRCK beta | CDC42BPB | 82765 | 38106 | 97387 | 0 | 29814 | 0 |
| Q9Y5S9;Q9Y5S9-2 | RNA-binding protein 8A | RBM8A | 5236200 | 1264700 | 13109000 | 568060 | 2533000 | 801280 |
| Q9Y5T5-5;Q9Y5T5-2;Q9Y5T5;H9KVB6;Q9Y5T5-3 | Ubiquitin carboxyl-terminal hydrolase 16 | USP16 | 1449600 | 740870 | 1513400 | 1618200 | 1687800 | 684320 |
| Q9Y678;H0Y8X7 | Coatomer subunit gamma-1 | COPG1 | 263670 | 408940 | 287870 | 0 | 175010 | 74738 |
| Q9Y6A4 | Cilia- and flagella-associated protein 20 | CFAP20 | 691110 | 284500 | 298950 | 121110 | 448910 | 302010 |
| Q9Y6W5;Q9Y6W5-2 | Wiskott-Aldrich syndrome protein family member 2 | WASF2 | 428620 | 317540 | 603740 | 54266 | 193180 | 70417 |
| R4GNH3;E9PN50 | | PSMC3 | 606290000 | 299230000 | 269030000 | 98061000 | 36197000 | 20384000 |

### Table S2. Statistical analysis and fold-changes using normalized values

| Protein names | Gene names | Normal #1 | Normal #2 | Normal #3 | lucose-free # | glucose-free #2 | glucose-free #3 | p-value | Log2(Glucose-free/Normal) |
|---|---|---|---|---|---|---|---|---|---|
| | | Normalization with PSMB2 values for each group | | | | | | Analysis | |
| Proteasome subunit beta type-2 | PSMB2 | 1 | 1 | 1 | 1 | 1 | 1 | 0.001556127 | 0 |
| Ubiquitin-like domain-containing CTD phosphatase 1 | UBLCP1 | 0.004804165 | 0.00351157 | 0.00466079 | 0.00020401 | 0.000282865 | 6.61396E-05 | 0.012890741 | -4.513863045 |
| YTH domain-containing family protein 3;YTH domain-containing family protein 1 | YTHDF3;YTHDF1 | 0 | 0.00010189 | 0.00066644 | 0.00068636 | 0.001509891 | 0.00025423 | 0.396111582 | 1.397242861 |
| WD repeat-containing protein 3 | WDR3 | 0.000208051 | 0.00029641 | 0.00047884 | 0.00240832 | 0.003883152 | 0.00224613 | 0.015864424 | 3.022361673 |
| 40S ribosomal protein S17-like;40S ribosomal protein S17 | RPS17;RPS17L | 0.080522898 | 0.04305663 | 0.08491369 | 0.19192603 | 0.272565329 | 0.239392464 | 0.047991311 | 1.571906737 |
| Lymphoid-specific helicase | HELLS | 0.000157524 | 0.00020998 | 0.00012153 | 5.1296E-05 | 0.000115068 | 0.000218732 | 0.441831886 | -0.45308497 |
| Ribosome biogenesis protein BOP1 | BOP1 | 0.000152564 | 0.00014646 | 0.00021141 | 0.0008075 | 0.001707719 | 0.000960491 | 0.020773556 | 2.654357521 |
| Ribosome biogenesis regulatory protein homolog | RRS1 | 0 | 0.00035721 | 0.00057228 | 0.00509875 | 0.009586263 | 0.005043131 | 0.013908835 | 3.4953291 |
| Far upstream element-binding protein 2 | KHSRP | 0.000890662 | 0.00091923 | 0.00153667 | 0.00087587 | 0.00170571 | 0.001177101 | 0.770592816 | 0.356740327 |
| Pyrroline-5-carboxylate reductase;Pyrroline-5-carboxylate reductase 2 | PYCR2 | 0 | 0 | 0.00020152 | 0 | 0.0002242 | 0 | NaN | 0.153891888 |
| UBX domain-containing protein 1 | UBXN1 | 0.000165732 | 0.0006907 | 0.00082869 | 0 | 0.000193608 | 0 | 0.183237183 | -1.834929012 |
| Heterogeneous nuclear ribonucleoprotein D-like | HNRNPDL | 0.000615105 | 0.00079639 | 0.00187515 | 5.7313E-05 | 0.001175862 | 0.000796845 | 0.393678893 | 0.000825828 |
| 40S ribosomal protein S24 | RPS24 | 0.011262457 | 0.00558254 | 0.02031875 | 0.01277858 | 0.043149703 | 0.040044786 | 0.173106779 | 1.830092812 |
| Eukaryotic translation initiation factor 5B | EIF5B | 0.000961538 | 0.00201057 | 0.00309069 | 0.0005234 | 0.001451226 | 0.0039225 | 0.75727157 | -0.47033451 |
| Spectrin beta chain, non-erythrocytic 1 | SPTBN1 | 0.005023346 | 0.00359908 | 0.00709407 | 0.00054001 | 0.001095512 | 0.001035829 | 0.023678317 | -2.277862794 |
| C-Myc-binding protein | MYCBP | 0.007056321 | 0.00636858 | 0.00913325 | 0.00415761 | 0.008770442 | 0.006853081 | 0.667697881 | -0.042163453 |
| Antigen KI-67 | MKI67 | 0.000126626 | 0.00024568 | 0.00026747 | 0.00027947 | 0.000143087 | 0.000433531 | 0.583283782 | 0.185906424 |
| TAR DNA-binding protein 43 | TARDBP | 0.003234856 | 0.00209222 | 0.00447931 | 0.0046471 | 0.007098351 | 0.004564697 | 0.209072791 | 0.522629198 |
| ATP-dependent RNA helicase DDX50 | DDX50 | 0.000339945 | 0.00064903 | 0.00159503 | 0.00237026 | 0.006741003 | 0.00496641 | 0.058573022 | 2.935841852 |
| Clathrin heavy chain;Clathrin heavy chain 1 | CLTC | 0.031969544 | 0.01768764 | 0.02700703 | 0.00086123 | 0.003311233 | 0.002574822 | 0.019186445 | -3.390790548 |
| Ribosome-binding protein 1 | RRBP1 | 0.000104417 | 0.00029148 | 0.00083181 | 0.00130108 | 0.005597032 | 0.003228897 | 0.067179254 | 3.469577569 |
| DNA-directed RNA polymerases I, II, and III subunit RPABC1 | POLR2E | 0.007415183 | 0.01398723 | 0.0210832 | 0.00979799 | 0.01681333 | 0.01292766 | 0.980090783 | -0.113649019 |
| Endophilin-B1 | SH3GLB1 | 5.62087E-05 | 0 | 0 | 0 | 0 | 0 | NaN | NaN |
| | PSMD1 | 0.001541989 | 0.00096317 | 0.00075591 | 0.00014718 | 8.91844E-05 | 0 | 0.019165534 | -3.071569086 |
| Round spermatid basic protein 1 | RSBN1;RSBN1L | 0 | 0 | 6.3561E-05 | 6.2909E-05 | 0.000383788 | 0 | 0.36939974 | 1.289610265 |
| Hepatoma-derived growth factor-related protein 2 | HDGFRP2 | 0.002457508 | 0.00223898 | 0.00130185 | 0.00183132 | 0.002493939 | 0.001028021 | 0.713737239 | -0.289961244 |
| 60S ribosomal protein L17 | RPL17 | 0.111180159 | 0.13163404 | 0.2264225 | 0.21003064 | 0.481563333 | 0.39805277 | 0.10785976 | 1.596427068 |
| YTH domain-containing family protein 2 | YTHDF2 | 0.004018307 | 0.00416832 | 0.01182665 | 0.00957975 | 0.019729176 | 0.014785318 | 0.128342722 | 1.826625947 |
| Unconventional myosin-Va | MYO5A | 2.54423E-05 | 2.0108E-05 | 6.2027E-05 | 9.2847E-06 | 8.37961E-05 | 4.52653E-05 | 0.965222344 | 0.831176419 |
| Cyclin-dependent kinase 1 | CDK1 | 0.004670474 | 0.00512572 | 0.00798776 | 0.00163898 | 0.004912203 | 0.003585435 | 0.177617001 | -0.515606508 |
| AP-2 complex subunit mu | AP2M1 | 0.002765088 | 0.00200073 | 0.00437775 | 0.00227813 | 0.002599751 | 0.002873527 | 0.761303944 | -0.088951945 |
| ATP-dependent RNA helicase DHX29 | DHX29 | 0.0003116 | 0.00025353 | 0.00119618 | 0.00063613 | 0.002456169 | 0.001892085 | 0.166826446 | 2.602209763 |
| Selenoprotein H | C11orf31;SELH | 0.000746333 | 0.00131885 | 0.00122541 | 0.00130474 | 0.003846988 | 0.004443579 | 0.116066827 | 1.650465977 |
| Signal recognition particle 19 kDa protein | SRP19 | 0.003070765 | 0.0018456 | 0 | 0.00120604 | 0.004335463 | 0.002511732 | 0.985953186 | 0.07733931 |
| Prostaglandin E synthase 3 | PTGES3 | 0.003830198 | 0.00990945 | 0.00329911 | 0.00328893 | 0.006586169 | 0 | 0.899636027 | 0.281110383 |
| Sorting nexin;Sorting nexin-9 | SNX9 | 0.000467708 | 0.00055236 | 0.0006038 | 0.00027121 | 0.000403089 | 0.000445448 | 0.316616523 | -0.454513047 |
| Myristoylated alanine-rich C-kinase substrate | MARCKS | 0.003574572 | 0.0035727 | 0.00878153 | 0.00038004 | 0.001119728 | 0.000771746 | 0.031787561 | -2.211572506 |
| Heterogeneous nuclear ribonucleoprotein A/B | HNRNPAB | 0.007893293 | 0.01109599 | 0.02601973 | 0.01996022 | 0.04711429 | 0.035739461 | 0.136540405 | 1.687479354 |
| Kinesin-like protein;Kinesin-like protein KIF3A | KIF3A | 0 | 0 | 0 | 0 | 0 | 0 | NaN | NaN |
| Ribosome maturation protein SBDS | SBDS | 0.009433994 | 0.00281158 | 0.01131154 | 0.00075353 | 0.003849667 | 0.00294908 | 0.169604581 | -1.677603724 |
| | TPM3 | 0.003540813 | 0.00042245 | 0.00490538 | 0 | 0 | 0 | NaN | NaN |
| Hypoxia up-regulated protein 1 | HYOU1 | 0.00018031 | 0.00015863 | 0.00039612 | 0.00010947 | 0.00026043 | 0.000186309 | 0.553216982 | 0.047217957 |
| RNA-binding protein 26 | RBM26 | 0 | 2.9181E-05 | 0 | 0 | 0 | 0 | NaN | NaN |
| Calcium-binding protein 39;Calcium-binding protein 39-like | CAB39;CAB39L | 0.000700593 | 0.00014052 | 0.00033556 | 8.6083E-05 | 0 | 0 | 0.080967186 | -1.962754968 |
| Kanadaptin | SLC4A1AP | 8.00582E-05 | 4.4836E-05 | 9.9763E-05 | 0 | 0.000211998 | 0 | 0.031714396 | 1.40492957 |
| Chromatin target of PRMT1 protein | CHTOP | 0.00071151 | 0 | 0 | 0.00070134 | 0.003147694 | 0.002951222 | 0.14613929 | 2.05235651 |
| Ribonuclease P protein subunit p40 | RPP40 | 0.00313386 | 0.00092097 | 0.0026965 | 0.00213718 | 0.003824353 | 0.002924934 | 0.418606871 | 0.117314967 |
| Ankyrin repeat domain-containing protein 36B;Ankyrin repeat domain-containing protein 36A | ANKRD36B;ANKRD36 | 0.000122579 | 0.00014592 | 0.0002511 | 4.7345E-05 | 2.34446E-05 | 0 | 0.038038941 | -2.130899997 |
| Proteasome activator complex subunit 2 | PSME2 | 0.142212518 | 0.14030148 | 0.20858526 | 0.13591356 | 0.159721943 | 0.162620972 | 0.886602766 | 0.167514066 |
| WD repeat-containing protein 6 | WDR6 | 0 | 8.2192E-05 | 0.00013954 | 3.9051E-05 | 0.00020352 | 0.000513582 | 0.71525167 | 0.926306603 |
| C-Jun-amino-terminal kinase-interacting protein 4 | SPAG9 | 1.95336E-05 | 0 | 2.1405E-05 | 2.7866E-05 | 0 | 0 | 0.350500492 | 0.446589076 |
| Putative pre-mRNA-splicing factor ATP-dependent RNA helicase DHX16 | DHX16 | 0.000651271 | 0.00089312 | 0.00098347 | 0.00012484 | 0.000230107 | 0.00015035 | 0.029082307 | -2.570534008 |
| Mortality factor 4-like protein 2 | MORF4L2 | 0.001220076 | 0.00068081 | 0.00298 | 0.00055453 | 0.001315024 | 0.000635031 | 0.320660216 | -0.942072092 |
| p21-activated protein kinase-interacting protein 1 | PAK1IP1 | 9.24141E-05 | 0.00024269 | 0.0002499 | 0.00352384 | 0.006468839 | 0.003903612 | 0.010301914 | 4.007634984 |
| Protein AAR2 homolog | AAR2 | 0.000878345 | 0.00063191 | 0.00123573 | 0.00018373 | 0.000397048 | 0.000221653 | 0.048189383 | -1.98648483 |
| RNA-binding protein 7 | RBM7 | 0.000776061 | 0.00062613 | 0.00087653 | 0.00256851 | 0.003830514 | 0.002894168 | 0.039512488 | 1.898906709 |
| 40S ribosomal protein S6 | RPS6 | 0.001505319 | 0.00209986 | 0.00348909 | 0 | 0.005366925 | 0.003966508 | 0.157365623 | 1.1356867 |
| Prefoldin subunit 6 | PFDN6 | 0.000182656 | 0.00146676 | 0.00242534 | 0.00087248 | 0.000787359 | 0.001049031 | 0.967278843 | -0.749437836 |
| Negative elongation factor E | NELFE | 0 | 0.00015581 | 0.00020018 | 0.00028608 | 0.000359786 | 0.000384305 | 0.18152954 | 1.026562256 |
| Zinc finger CCCH domain-containing protein 11A | ZC3H11A | 0 | 0 | 1.8699E-05 | 0 | 0.000277896 | 0.000162946 | 0.010977475 | 3.508475057 |
| PEST proteolytic signal-containing nuclear protein | PCNP | 0 | 0.00112143 | 0.00134688 | 0.00428372 | 0.004420648 | 0.003454191 | 0.068329922 | 1.801383816 |
| SLIT-ROBO Rho GTPase-activating protein 2 | SRGAP2 | 0.001399899 | 0.00152067 | 0.00148032 | 0.00010472 | 0.000269351 | 0.00030441 | 0.019180579 | -2.458347801 |
| Exosome complex component RRP4 | EXOSC2 | 0 | 0.00046511 | 0.00013976 | 0.00169704 | 0.000472014 | 0.000550599 | 0.199258786 | 1.110755738 |
| CCR4-NOT transcription complex subunit 1 | CNOT1 | 7.05912E-05 | 9.1968E-05 | 0.00024156 | 6.6938E-05 | 0.001107352 | 0.000138076 | 0.541745242 | 0.586258704 |
| ADP-sugar pyrophosphatase | NUDT5 | 0 | 0 | 0.00035682 | 0 | 0 | 0 | NaN | NaN |
| Ras-related protein R-Ras2 | RRAS2 | 0.000163263 | 0 | 0.00076881 | 0 | 0 | 0 | NaN | NaN |
| 40S ribosomal protein SA | RPSA;RPSAP58 | 0.160687493 | 0.20385219 | 0.32221207 | 0.24434769 | 0.559261194 | 0.436744231 | 0.186066287 | 1.099265169 |
| Host cell factor 1;HCF N-terminal chain 1;HCF N-terminal chain 2;HCF N-terminal chain 3;HCF N-terminal chain 4;HCF N-terminal chain 5;HCF N-terminal chain 6;HCF C-terminal chain 1;HCF C-terminal chain 2;HCF C-terminal chain 3;HCF C-terminal chain 4;HCF C-terminal chain 5;HCF C-terminal chain 6 | HCFC1 | 2.24449E-05 | 6.0404E-05 | 2.3422E-05 | 0.00013358 | 0.000280682 | 9.73888E-05 | 0.051078368 | 2.511766323 |
| Small nuclear ribonucleoprotein E | SNRPE | 0.012702385 | 0.01974877 | 0.02566536 | 0.01480406 | 0.04750807 | 0.040821731 | 0.292831942 | 1.047574558 |
| Structural maintenance of chromosomes flexible hinge domain-containing protein 1 | SMCHD1 | 8.71123E-05 | 5.4407E-05 | 0.0001375 | 0.00064396 | 0.001163579 | 0.000564775 | 0.016288577 | 2.886024975 |
| Obscurin-like protein 1 | OBSL1 | 0.000166034 | 0.00017792 | 0.00034083 | 5.7074E-05 | 0.00012942 | 3.33638E-05 | 0.087465142 | -1.640357548 |
| Afadin | MLLT4 | 0.000449121 | 0.0003001 | 0.00023736 | 9.5231E-05 | 0.000185813 | 6.00428E-05 | 0.070687115 | -1.655965683 |
| Small nuclear ribonucleoprotein-associated proteins B and B;Small nuclear ribonucleoprotein-associated protein N | SNRPN;SNRPB | 0.003379353 | 0.01940752 | 0.00585944 | 0.00179584 | 0.008278218 | 0.000913621 | 0.251791752 | -1.706104528 |
| Glycogen synthase kinase-3 alpha | GSK3A | 0.00061001 | 0.00077437 | 0.00056056 | 0.00014521 | 0.000292295 | 0 | 0.075176321 | -1.566035011 |
| Carbonyl reductase [NADPH] 1;Carbonyl reductase [NADPH] 3 | CBR1;CBR3 | 0 | 0 | 0.00022217 | 0 | 0 | 0 | NaN | NaN |
| Small ubiquitin-related modifier 3 | SUMO3 | 0.0026043 | 0.00071491 | 0.00165312 | 0 | 0.000912431 | 0.279992211 | 0.323442396 | 3.273324545 |
| Protein TSSC1 | TSSC1 | 0.001165491 | 0.00140804 | 0.0029789 | 0.00093275 | 0.001686824 | 0.001325148 | 0.457666125 | -0.087533797 |
| 60S ribosomal protein L23a | RPL23A | 0.171604524 | 0.173736 | 0.37629921 | 0.24833629 | 0.758615609 | 0.498159868 | 0.181020499 | 1.519712083 |
| Suppressor of SWI4 1 homolog | PPAN | 0.001808868 | 0.00148508 | 0.0026006 | 0.00618062 | 0.014089016 | 0.009506182 | 0.030474509 | 2.39377889 |
| TFIIH basal transcription factor complex helicase XPD subunit | ERCC2 | 0.000268889 | 0.00031273 | 0.0003755 | 0.00012685 | 0.000181728 | 0.000155876 | 0.133821358 | -1.004514194 |
| M-phase phosphoprotein 6 | MPHOSPH6 | 0.000324325 | 0.00038368 | 0.00070275 | 0.00884728 | 0.015727086 | 0.01113173 | 0.008798946 | 4.858626687 |
| Matrin-3 | MATR3 | 0.000840835 | 0.00012236 | 0.00054289 | 0.00056838 | 0.000450449 | 0.001277188 | 0.420010501 | 0.066198419 |
| Putative RNA-binding protein Luc7-like 1 | LUC7L | 0 | 0 | 0.00074302 | 0 | 0 | 0 | NaN | NaN |
| Eukaryotic translation initiation factor 3 subunit L | EIF3L | 0.023656365 | 0.02503925 | 0.05129966 | 0.06288556 | 0.11626552 | 0.095591471 | 0.070552218 | 1.932690264 |
| RNA-binding protein EWS | EWSR1 | 0.005714366 | 0.00582309 | 0.01052699 | 0.00900263 | 0.020313149 | 0.015035732 | 0.145487045 | 1.368537788 |
| Coatomer subunit delta | ARCN1 | 0.000659277 | 0.00045983 | 0.00083438 | 0.00019595 | 0.000584415 | 0.000182506 | 0.143334458 | -1.750421266 |
| Secretory carrier-associated membrane protein 3 | SCAMP3 | 0.000377181 | 0.00019992 | 0.00187113 | 0 | 0 | 0 | NaN | NaN |
| Forkhead-associated domain-containing protein 1 | FHAD1 | 0.000354826 | 0 | 0 | 0 | 8.12124E-05 | 0 | NaN | -2.127339799 |
| F-actin-capping protein subunit beta | CAPZB | 0.004581066 | 0.00311222 | 0.00367029 | 0 | 0.00316082 | 0.001938721 | 0.335921278 | -0.568192018 |
| ELAV-like protein;ELAV-like protein 2;ELAV-like protein 3;ELAV-like protein 4 | ELAVL2;ELAVL4;ELAVL3 | 0 | 0.00034115 | 0.00043418 | 0 | 0.000747137 | 0.000505676 | 0.321597172 | 0.675450414 |
| ATP-dependent RNA helicase DDX19B;ATP-dependent RNA helicase DDX19A | DDX19A;DDX19B | 0.0001247 | 7.5183E-05 | 0.00077803 | 0 | 0 | 0 | NaN | NaN |
| Tudor domain-containing protein 3 | TDRD3 | 0.008650207 | 0.02032346 | 0.02157001 | 0.01600817 | 0.032592652 | 0.038095609 | 0.241726068 | 0.68140078 |
| Exosome complex component CSL4 | EXOSC1 | 0.000695051 | 0.00017406 | 0 | 0 | 0.000417407 | 0 | 0.811667682 | 0.26309888 |
| THO complex subunit 6 homolog | THOC6 | 9.03874E-05 | 0 | 0.00037648 | 0 | 0 | 0 | NaN | NaN |
| Epidermal growth factor receptor substrate 15 | EPS15 | 0.000256942 | 0.00011934 | 0.00027983 | 0 | 6.7252E-05 | 0 | 0.038494241 | -1.933793656 |
| Putative high mobility group protein B1-like 1;High mobility group protein B1 | HMGB1;HMGB1P1 | 0.000794032 | 0.00065628 | 0.00141081 | 0.00052336 | 0.000440056 | 0.000150351 | 0.11813708 | -0.851510022 |
| Bifunctional arginine demethylase and lysyl-hydroxylase JMJD6 | JMJD6 | 0 | 0.00026223 | 0 | 0 | 0 | 0 | NaN | NaN |
| Adenylyl cyclase-associated protein 1 | CAP1 | 0.001021442 | 0.00059767 | 0.00079865 | 1.7352E-05 | 4.96364E-05 | 0 | 0.012474102 | -4.766237656 |

| | | | | | | | | | |
|---|---|---|---|---|---|---|---|---|---|
| Eukaryotic translation initiation factor 3 subunit H | EIF3H | 0.013065726 | 0.01319167 | 0.02963102 | 0.0337709 | 0.078646147 | 0.082476877 | 0.065409144 | 2.575749092 |
| Double-strand break repair protein MRE11A | MRE11A | 0.000237734 | 0.00045042 | 0.00087392 | 0.00027456 | 0.000798784 | 0.000259975 | 0.776327765 | -0.714172752 |
| Nucleosome assembly protein 1-like 1 | NAP1L1 | 0 | 0 | 0 | 0.00066483 | 0 | 0 | NaN | NaN |
| Arf-GAP with GTPase, ANK repeat and PH domain-containing protein 3;Arf-GAP with GTPase, ANK repeat and PH domain-containing protein 1 | AGAP1;AGAP3 | 0.000105694 | 0.00016642 | 0.00094584 | 0 | 0 | 0 | NaN | NaN |
| Polymerase delta-interacting protein 2 | POLDIP2 | 9.07737E-05 | 0.00014472 | 0.00016592 | 0 | 6.01238E-05 | 3.91705E-05 | 0.098349604 | -1.57633057 |
| Exocyst complex component 7 | EXOC7 | 8.30702E-05 | 6.3238E-05 | 0.00024497 | 0 | 3.55795E-05 | 0 | 0.088012841 | -1.223284791 |
| Citrate synthase;Citrate synthase, mitochondrial | CS | 0.00370412 | 0.00253899 | 0.00212659 | 0.00217223 | 0.002264636 | 0.001043774 | 0.260030113 | -0.225079551 |
| Periodic tryptophan protein 1 homolog | PWP1 | 0.001179991 | 0.00043769 | 0.00277806 | 0.00174499 | 0.004492171 | 0.0007225 | 0.584457653 | 0.564440433 |
| Cysteine--tRNA ligase, cytoplasmic | CARS | 0.000222327 | 0.00020232 | 0.00045609 | 8.5592E-05 | 0.000317488 | 0.000242917 | 0.47039474 | 0.127780293 |
| Protein unc-45 homolog A | UNC45A | 2.34341E-05 | 0 | 0.00011467 | 0 | 0 | 0 | NaN | NaN |
| Cytoplasmic dynein 1 intermediate chain 2 | DYNC1I2 | 6.75792E-05 | 0 | 0.00027688 | 0 | 0 | 0 | NaN | NaN |
| T-complex protein 1 subunit gamma | CCT3 | 0.004879465 | 0.00346226 | 0.00456772 | 0.0037165 | 0.005438716 | 0.003218771 | 0.882508881 | -0.297531721 |
| Nucleoside diphosphate kinase 7 | NME7 | 0.000598197 | 0.00037459 | 0.00079321 | 8.6666E-05 | 0.000370327 | 0.00013063 | 0.095380481 | -2.195134416 |
| Fragile X mental retardation syndrome-related protein 1 | FXR1 | 0.002462714 | 0.00280237 | 0.00598676 | 0.00288921 | 0.010812472 | 0.007804303 | 0.282352856 | 1.477624402 |
| Heterogeneous nuclear ribonucleoproteins C1/C2;Heterogeneous nuclear ribonucleoprotein C-like 4;Heterogeneous nuclear ribonucleoprotein C-like 1;Heterogeneous nuclear ribonucleoprotein C-like 3;Heterogeneous nuclear ribonucleoprotein C-like 2 | HNRNPC;HNRNPCL4;HNRNP | 0.01010861 | 0.01285251 | 0.01672025 | 0.01709617 | 0.039694084 | 0.023845779 | 0.151000451 | 0.891684116 |
| Interleukin enhancer-binding factor 2 | ILF2 | 0.008104356 | 0.00711274 | 0.01920723 | 0.04280922 | 0.028795489 | 0.025345147 | 0.064194822 | 1.829073368 |
| Serine/threonine-protein kinase 26 | STK26 | 0.0019275 | 0.00160641 | 0.00234852 | 0.0002319 | 0.000526902 | 0.000492571 | 0.030073328 | -1.968327263 |
| Apoptosis inhibitor 5 | API5 | 0 | 0 | 0.00027788 | 0 | 0.000172982 | 0.000196592 | 0.246378013 | -0.591543542 |
| Condensin complex subunit 2 | NCAPH | 3.05313E-05 | 2.4055E-05 | 0.00027187 | 0 | 0 | 0 | NaN | NaN |
| GTP-binding nuclear protein Ran | RAN | 0.010119807 | 0.00788632 | 0.01991871 | 0.00531162 | 0.014618074 | 0.007024632 | 0.427912087 | -0.526687222 |
| Abl interactor 1;Abl interactor 2 | ABI1;ABI2 | 0.000335013 | 0.0002939 | 0.00041193 | 5.8931E-05 | 0.000282276 | 7.63996E-05 | 0.112228891 | -2.13258009 |
| Heterogeneous nuclear ribonucleoprotein L-like | HNRNPLL;HNRPLL | 0.000304742 | 0.00061431 | 0.00054614 | 0.00013153 | 0.000147011 | 0.000170887 | 0.059379482 | -1.893349359 |
| Heterogeneous nuclear ribonucleoprotein U-like protein 1 | HNRNPUL1 | 0.003843411 | 0.00395603 | 0.00645109 | 0.00700941 | 0.012021939 | 0.010835362 | 0.128611483 | 1.453620258 |
| Zinc finger protein 346 | ZNF346 | 0.000498197 | 0.00056607 | 0.00112412 | 0.00044726 | 0.001600836 | 0.001413708 | 0.462471311 | 1.320424146 |
| Calpastatin | CAST | 0.000343791 | 0.00022482 | 0.0004893 | 0.00154126 | 0.000578441 | 0.000619394 | 0.114013564 | 0.84932563 |
| Annexin A7 | ANXA7 | 0.000847609 | 0.00139611 | 0.0035933 | 6.6702E-05 | 0 | 0 | 0.012362458 | -4.38753249 |
| Eukaryotic peptide chain release factor subunit 1 | ETF1 | 0.001150431 | 0.0008761 | 0.00171815 | 0.00063464 | 0.002516709 | 0.001784208 | 0.743712588 | 0.633109355 |
| Rab GTPase-activating protein 1-like, isoform 10 | RABGAP1L | 0 | 0 | 0 | 0 | 0 | 0 | NaN | NaN |
| T-complex protein 1 subunit epsilon | CCT5 | 0.002880361 | 0.00222621 | 0.00431647 | 0.00616718 | 0.004601465 | 0.003150813 | 0.313914259 | 0.675843617 |
| Squamous cell carcinoma antigen recognized by T-cells 3 | SART3 | 0.001673945 | 0.00177389 | 0.00272463 | 0.00251529 | 0.004012805 | 0.00257268 | 0.315974402 | 0.53635303 |
| Chromobox protein homolog 3 | CBX3 | 0.002078323 | 0.00246143 | 0.00480236 | 0.00145987 | 0.004971136 | 0.002702366 | 0.877643132 | 0.134728715 |
| Pescadillo homolog | PES1 | 0 | 0 | 0.00015665 | 0.00208923 | 0.002213873 | 0.001220271 | 0.008322396 | 3.737319884 |
| U6 snRNA-associated Sm-like protein LSm5 | LSM5 | 0.023085881 | 0.01973307 | 0.02695223 | 0.02305757 | 0.028368224 | 0.010752994 | 0.630167356 | -0.001770249 |
| Small ubiquitin-related modifier 1 | SUMO1 | 0.005261841 | 0.00248152 | 0.00370911 | 0.00088648 | 0.001544046 | 0.001490351 | 0.06978228 | -1.315419351 |
| Prothymosin alpha;Prothymosin alpha, N-terminally processed;Thymosin alpha-1 | PTMA | 0.006687941 | 0.0049373 | 0.00757476 | 0.01292372 | 0.015226156 | 0.013905365 | 0.11480192 | 1.056007589 |
| Ankyrin repeat and zinc finger domain-containing protein 1 | ANKZF1 | 0.000138736 | 0.00018195 | 0.00043507 | 3.542E-05 | 0.000170839 | 4.64025E-05 | 0.137353458 | -1.971291398 |
| Transcription elongation factor B polypeptide 2 | TCEB2 | 0.016159445 | 0.01340103 | 0.02254452 | 0.01555126 | 0.02162307 | 0.023394022 | 0.681578306 | 0.420193719 |
| DNA-directed RNA polymerase;DNA-directed RNA polymerase I subunit RPA1 | POLR1A | 0.000298936 | 0.00054438 | 0.0010432 | 0.00045108 | 0.001000054 | 0.000365573 | 0.983794751 | -0.271247741 |
| Histone H2A;Histone H2A.V;Histone H2A.Z;Histone H2A type 1-J;Histone H2A type 1-H;Histone H2A.J;Histone H2A type 2-C;Histone H2A type 1-C;Histone H2A type 2-B;Histone H2A type 3;Histone H2A type 2-A;Histone H2A type 1-D;Histone H2A type 1;Histone H2A type 1-B/E;Histone H2A type 1-A;Histone H2AX | H2AFV;H2AFZ;HIST1H2AJ;HIS | 0.005310827 | 0.01037904 | 0.01248881 | 0.00427845 | 0.00922503 | 0.009251874 | 0.601755063 | -0.170047918 |
| Cystatin-A;Cystatin-A, N-terminally processed | CSTA | 0.001836357 | 0.0086957 | 0.00468682 | 0.00174972 | 0.006893157 | 0.004954922 | 0.912075341 | 0.080252651 |
| General transcription factor IIE subunit 1 | GTF2E1 | 3.0856E-05 | 4.7091E-05 | 0.00030258 | 0 | 0 | 0 | NaN | NaN |
| Sorcin | SRI | 0.000187263 | 0.00113242 | 0.0011676 | 0.0006607 | 0.001161892 | 0.000216902 | 0.879438322 | -0.777340611 |
| Gamma-aminobutyric acid receptor-associated protein-like 2 | GABARAPL2 | 0.001881536 | 0.00033942 | 0.00117682 | 0 | 0 | 0 | NaN | NaN |
| DNA-directed RNA polymerase;DNA-directed RNA polymerase II subunit RPB2 | POLR2B | 0.001827791 | 0.0021298 | 0.00331217 | 0.00105171 | 0.001567217 | 0.001123766 | 0.158947803 | -0.922377658 |
| Vacuolar protein sorting-associated protein 29 | VPS29 | 0.011806629 | 0.00716288 | 0.01204128 | 0.00173682 | 0.001717229 | 0.00100072 | 0.019721375 | -2.781442762 |
| Protein CMSS1 | CMSS1 | 0.001145336 | 0.00153868 | 0.00222331 | 0.00333817 | 0.007856846 | 0.005489047 | 0.05329679 | 1.834863405 |
| Heat shock protein beta-1 | HSPB1 | 0.003237543 | 0.0077562 | 0.01036168 | 0 | 0.001706647 | 0.002158894 | 0.093370095 | -2.01462183 |
| 60S ribosomal protein L37a | RPL37A | 0.026997537 | 0.02529886 | 0.07669559 | 0.06074397 | 0.150881987 | 0.151274462 | 0.113393054 | 2.48252088 |
| | HDLBP | 0.000425031 | 0 | 0 | 0 | 0 | 0 | NaN | NaN |
| Nucleosome assembly protein 1-like 4 | NAP1L4 | 0.001410928 | 0.00127363 | 0.00250516 | 0 | 0.000847069 | 0 | 0.087011432 | -0.736092971 |
| Intraflagellar transport protein 74 homolog | IFT74 | 3.25008E-05 | 0.00010196 | 0.00032802 | 0 | 0 | 0 | NaN | NaN |
| Zinc finger CCCH-type antiviral protein 1 | ZC3HAV1 | 0.000246098 | 0.00027633 | 0.00044052 | 0.00030966 | 0.00066222 | 0.000413845 | 0.363679763 | 0.582687887 |
| Junction plakoglobin | JUP | 6.9673E-05 | 0.00041224 | 0.00013412 | 0.00015486 | 0.000348616 | 0.00027933 | 0.471274941 | 1.058424626 |
| DCN1-like protein;DCN1-like protein 1 | DCUN1D1 | 0.004052122 | 0.00128891 | 0.00275148 | 0.00111272 | 0.001448681 | 0.000393379 | 0.144391023 | -1.306114148 |
| 60S ribosomal export protein NMD3 | NMD3 | 0.000513621 | 0.00053158 | 0.00070316 | 0 | 0.00123671 | 0.001078493 | 0.167962941 | 1.119397866 |
| Trifunctional purine biosynthetic protein adenosine-3;Phosphoribosylamine--glycine ligase;Phosphoribosylformylglycinamidine cyclo-ligase;Phosphoribosylglycinamide formyltransferase | GART | 0.000358459 | 0 | 0 | 0 | 0 | 0 | NaN | NaN |
| Malate dehydrogenase;Malate dehydrogenase, cytoplasmic | MDH1 | 5.79106E-05 | 8.1515E-05 | 0.00042441 | 0.00020262 | 0 | 0 | 0.483572678 | 1.313665788 |
| COP9 signalosome complex subunit 1 | GPS1 | 0.000727242 | 0.00038858 | 0.00113435 | 0.00036166 | 0.000750499 | 0.000643968 | 0.634798205 | -0.175446521 |
| Proline-, glutamic acid- and leucine-rich protein 1 | PELP1 | 4.90091E-05 | 0 | 9.2849E-05 | 9.1041E-05 | 0.000477157 | 0.000243871 | 0.169660403 | 1.854080623 |
| Golgi to ER traffic protein 4 homolog | GET4 | 0.004634028 | 0 | 0.00095607 | 0.00137548 | 0 | 0 | 0.627913543 | -0.613794688 |
| E3 ubiquitin-protein ligase RNF123 | RNF123 | 0.002716773 | 0.00243138 | 0.00549466 | 0.00367704 | 0.009955666 | 0.006014799 | 0.204491453 | 1.146622501 |
| WD repeat and FYVE domain-containing protein 1 | WDFY1 | 9.66073E-05 | 0 | 0 | 0 | 0 | 0 | NaN | NaN |
| DNA-directed RNA polymerases I, II, and III subunit RPABC3 | POLR2H | 0.004833389 | 0.00623155 | 0.01270984 | 0.00555846 | 0.008574222 | 0.005786778 | 0.776342855 | -0.106830757 |
| 60S ribosomal protein L24 | RPL24 | 0.039834285 | 0.04632995 | 0.09523244 | 0.08232758 | 0.11460334 | 0.205977996 | 0.131186387 | 1.306632091 |
| Protein SEC13 homolog | SEC13 | 0.001946199 | 0.00033645 | 0.00097936 | 0 | 0 | 0 | NaN | NaN |
| Peptidyl-prolyl cis-trans isomerase;Peptidyl-prolyl cis-trans isomerase H | PPIH | 0.000666163 | 0.00152748 | 0.00169038 | 0.00099125 | 0.001120987 | 0 | 0.765801792 | -0.535107122 |
| Uncharacterized protein C7orf50 | C7orf50 | 0.006556937 | 0.00620046 | 0.01465431 | 0.01326345 | 0.033858373 | 0.020050628 | 0.112302511 | 1.612553485 |
| CGG triplet repeat-binding protein 1 | CGGBP1 | 0.00044588 | 0.00108783 | 0.00154571 | 0.00154889 | 0.003898287 | 0.001901821 | 0.147085021 | 0.805933653 |
| Eukaryotic translation initiation factor 4 gamma 2 | EIF4G2 | 0.001010021 | 0.00090445 | 0.00213198 | 0.00101218 | 0.003350879 | 0.001427514 | 0.541017134 | 0.499119605 |
| Cytoplasmic dynein 2 heavy chain 1 | DYNC2H1 | 3.54781E-05 | 3.9204E-05 | 5.0083E-05 | 0.00053009 | 0.00094684 | 0.000886457 | 0.010518536 | 4.498963165 |
| 60S ribosomal protein L32 | RPL32 | 0.10408129 | 0.07969538 | 0.1272171 | 0.15065312 | 0.302781908 | 0.293330737 | 0.104587678 | 1.494817505 |
| Epsin-3;Epsin-1 | EPN3;EPN1 | 0.000599877 | 0 | 0.00082356 | 0 | 0 | 0.000332976 | 0.101186792 | -1.077855547 |
| Calcium/calmodulin-dependent protein kinase type II subunit delta | CAMK2D | 0.000975087 | 0.00048812 | 0.00080695 | 0.00023892 | 0.000286174 | 0.00015794 | 0.054448624 | -1.755963734 |
| Cyclin-dependent kinase 7 | CDK7 | 0.001179655 | 0.00146792 | 0.00190374 | 0.00035031 | 0.000699294 | 0.000141832 | 0.065689061 | -2.067069455 |
| 60S ribosomal protein L9 | RPL9 | 0.026650431 | 0.04072438 | 0.06669924 | 0.0648218 | 0.170370073 | 0.135141661 | 0.09634401 | 1.730507852 |
| COP9 signalosome complex subunit 4 | COPS4 | 0.000391132 | 0.00023976 | 0.00060087 | 5.7758E-05 | 0 | 0.000180136 | 0.105520593 | -1.939060966 |
| 40S ribosomal protein S23 | RPS23 | 0.117136939 | 0.09450539 | 0.16924833 | 0.0867473 | 0.257724917 | 0.117361503 | 0.781528071 | 0.002763152 |
| Ubiquitin-conjugating enzyme E2 K | UBE2K | 0 | 0 | 0 | 0 | 0 | 0 | NaN | NaN |
| PDZ and LIM domain protein 1 | PDLIM1 | 0.012652559 | 0.00395792 | 0.00704831 | 0 | 0 | 0 | NaN | NaN |
| Prefoldin subunit 1 | PFDN1 | 0.000288685 | 0.00155899 | 0.00249073 | 0 | 0.00018221 | 0.000655262 | 0.31006625 | -2.173699989 |
| | NPM1 | 0 | 0 | 0 | 0 | 0 | 0 | NaN | NaN |
| Transcription elongation factor B polypeptide 1 | TCEB1 | 0.003801086 | 0.0020313 | 0.00373742 | 0.00093476 | 0.00203252 | 0.001222101 | 0.106654567 | -1.612678792 |
| Vacuolar protein sorting-associated protein 26A | VPS26A | 0.002743142 | 0.00241191 | 0.00279414 | 0.00030513 | 0.000661577 | 0.000393379 | 0.022299669 | -2.801837457 |
| Transcription elongation factor A protein 1 | TCEA1 | 0.000810548 | 0.00080432 | 0.00105507 | 0.00104934 | 0.001509757 | 0.001201032 | 0.375033856 | 0.567305062 |
| Double-stranded RNA-binding protein Staufen homolog 2 | STAU2 | 0.000605027 | 0.00084312 | 0.00202685 | 0.00073356 | 0.002886514 | 0.001703748 | 0.481999157 | 1.014907209 |
| S-phase kinase-associated protein 1 | SKP1 | 0.013340052 | 0.00892578 | 0.00965659 | 0.0140386 | 0.007754383 | 0.005784442 | 0.602118144 | -0.316501831 |
| Protein LYRIC | MTDH | 0.00011787 | 0.00050134 | 0.00089744 | 0.00072423 | 0.002227267 | 0.001876701 | 0.150435934 | 1.904337564 |
| DNA-3-methyladenine glycosylase | MPG | 6.32908E-05 | 0 | 0.00021189 | 0 | 0 | 0 | NaN | NaN |
| Switch-associated protein 70 | SWAP70 | 0.000413145 | 0.00041636 | 0.00057153 | 0.00058181 | 0.000565476 | 0.000474112 | 0.689128551 | 0.441630855 |
| Sequestosome-1 | SQSTM1 | 0.000395404 | 0.00041169 | 0.00048879 | 0.0002634 | 0.000226624 | 0.000100019 | 0.101173088 | -0.861268265 |
| Mitogen-activated protein kinase kinase kinase kinase 4 | MAP4K4 | 0.000126542 | 0.00011535 | 6.9046E-05 | 0 | 8.97242E-05 | 8.1881E-05 | 0.715013331 | -0.428383131 |
| Ras-related protein Rab-1A;Ras-related protein Rab-1B;Putative Ras-related protein Rab-1C | RAB1A;RAB1B;RAB1C | 0.003931363 | 0.00561175 | 0.00603078 | 0.00054288 | 0.001714282 | 0.000864706 | 0.028354938 | -2.698167849 |
| Condensin complex subunit 3 | NCAPG | 2.09904E-05 | 0 | 8.0306E-05 | 0 | 0 | 0 | NaN | NaN |
| 60S ribosomal protein L14 | RPL14 | 0.081502631 | 0.05742175 | 0.0959083 | 0.07596409 | 0.194438864 | 0.162570344 | 0.194531894 | 0.996145569 |
| Septin-7 | Sep-07 | 0.00024958 | 0 | 0 | 0 | 0 | 0 | NaN | NaN |
| Protein PRRC2C | PRRC2C | 0.000360184 | 0.00046408 | 0.00110366 | 0.00157781 | 0.004297023 | 0.003045273 | 0.051646292 | 2.714110594 |

| | | | | | | | | | |
|---|---|---|---|---|---|---|---|---|---|
| Gamma-tubulin complex component 6 | TUBGCP6 | 0 | 0 | 0 | 0 | 0 | 0 | NaN | NaN |
| Ezrin | EZR | 0 | 8.9207E-05 | 8.5534E-05 | 0.00010333 | 6.95772E-05 | 0.00010208 | 0.939067468 | 0.224806593 |
| Polyadenylate-binding protein;Polyadenylate-binding protein 1;Polyadenylate-binding protein 3 | PABPC1;PABPC3 | 0.008961482 | 0.01805715 | 0.02478217 | 0.01345052 | 0.037434537 | 0.029993185 | 0.35339108 | 0.732064125 |
| Brain-specific angiogenesis inhibitor 1-associated protein 2-like protein 1 | BAIAP2L1 | 0.000121263 | 2.4559E-05 | 0.0001884 | 0 | 0 | 0 | NaN | NaN |
| Neurofilament medium polypeptide | NEFM | 0.040532415 | 0.05718727 | 0.12567358 | 0.01317314 | 0.011391891 | 0.010742284 | 0.025387036 | -2.327686814 |
| Serine/threonine-protein phosphatase;Serine/threonine-protein phosphatase 2B catalytic subunit alpha isoform;Serine/threonine-protein phosphatase 2B catalytic subunit beta isoform | PPP3CA;PPP3CB | 0.000221089 | 0 | 0.00011135 | 0 | 0.000159976 | 0 | 0.957859558 | 0.027971647 |
| Ras-related protein Rab-11A;Ras-related protein Rab-11B | RAB11A;RAB11B | 0.004750644 | 0.00593112 | 0.00691871 | 0.00210547 | 0.001456048 | 0.001677461 | 0.052023186 | -1.822025579 |
| Treacle protein | TCOF1 | 1.63739E-05 | 0.00014409 | 0.0004201 | 0.00012779 | 0.00049339 | 0.00035202 | 0.364237457 | 1.288677171 |
| Serine/threonine-protein kinase MARK2;MAP/microtubule affinity-regulating kinase 3;MAP/microtubule affinity-regulating kinase 4;Serine/threonine-protein kinase MARK1 | MARK2;MARK4;MARK3;MARK | 4.3344E-05 | 3.0403E-05 | 3.8332E-05 | 0 | 3.92682E-05 | 3.58407E-05 | 0.972232128 | -0.031080423 |
| Basic leucine zipper and W2 domain-containing protein 2 | BZW2 | 0.002927052 | 0.00341798 | 0.00627199 | 0.00120389 | 0.003428698 | 0.002114302 | 0.177341938 | -0.692964182 |
| Golgin subfamily B member 1 | GOLGB1 | 0.000753611 | 0.000397 | 0.00053634 | 0.00017764 | 0.000228687 | 0 | 0.096657014 | -1.411993248 |
| Casein kinase II subunit alpha 3;Casein kinase II subunit alpha | CSNK2A1;CSNK2A3 | 0.000995689 | 0.0012864 | 0.0019989 | 0.00145772 | 0.004512932 | 0.003372408 | 0.163082355 | 1.390437532 |
| Protein FAM193A | FAM193A | 0.000687605 | 0.00032851 | 0.0012213 | 0.00081318 | 0.00319618 | 0.000705676 | 0.350768275 | 0.241996471 |
| Microtubule-associated protein;Microtubule-associated protein 4 | MAP4 | 0.001796495 | 0.00195352 | 0.00513225 | 0.00703962 | 0.012975449 | 0.009340668 | 0.056268157 | 2.257448225 |
| Cytoplasmic FMR1-interacting protein 2 | CYFIP2 | 0.000783563 | 0.00069361 | 0.00122313 | 8.5211E-05 | 0.000420836 | 0.000117256 | 0.06147979 | -2.740387527 |
| Protein SDA1 homolog | SDAD1 | 0.000283277 | 0.00047829 | 0.00143958 | 0.00065369 | 0.0022641 | 0.001806991 | 0.233829336 | 1.917631788 |
| Cleavage stimulation factor subunit 2 | CSTF2 | 0.000344833 | 0.00047203 | 0.0006264 | 0.00034518 | 0.000508673 | 0.000218752 | 0.39461205 | -0.451525498 |
| Eukaryotic translation initiation factor 4B | EIF4B | 0.01367596 | 0.01507066 | 0.01787469 | 0.00890136 | 0.02066005 | 0.012462857 | 0.683300645 | -0.274107576 |
| Dynactin subunit 1 | DCTN1;DKFZp686E0752 | 0.00045159 | 0.00048595 | 0.00052964 | 0.0017266 | 0.002084756 | 0.000383429 | 0.20985437 | 1.829050677 |
| Chromosome transmission fidelity protein 18 homolog;Tubulin-specific chaperone E | CHTF18;TBCE | 0.000188249 | 4.7904E-05 | 0.0002023 | 4.3077E-05 | 6.82378E-05 | 7.46295E-05 | 0.235488613 | -1.463999161 |
| Nascent polypeptide-associated complex subunit alpha;Nascent polypeptide-associated complex subunit alpha, muscle-specific form | NACA | 0.028588064 | 0.04914791 | 0.11670472 | 0.01912487 | 0.107493872 | 0.056276896 | 0.872008388 | 0.195412631 |
| THO complex subunit 4 | ALYREF | 0 | 0 | 0.00116157 | 0.00023272 | 0.005127309 | 0 | 0.970615194 | -0.088653395 |
| DNA-directed RNA polymerase II subunit RPB4 | POLR2D | 0.002696675 | 0.00092361 | 0.0041833 | 0 | 0.001997027 | 0.001405433 | 0.688085975 | -0.686747402 |
| Multifunctional protein ADE2;Phosphoribosylaminoimidazole-succinocarboxamide synthase;Phosphoribosylaminoimidazole carboxylase | PAICS | 0.00041768 | 0.00025956 | 0.00047403 | 0.00012882 | 0.000316256 | 0.000236686 | 0.194965353 | -0.819423691 |
| Translation initiation factor eIF-2B subunit epsilon | EIF2B5 | 0.000570877 | 0.00113242 | 0.00154416 | 0.00158104 | 0.004901354 | 0.003061046 | 0.107990496 | 1.434616865 |
| Heterogeneous nuclear ribonucleoprotein H;Heterogeneous nuclear ribonucleoprotein H, N-terminally processed;Heterogeneous nuclear ribonucleoprotein H2 | HNRNPH1;HNRNPH2 | 0.022698466 | 0.01658118 | 0.03322769 | 0.02917701 | 0.030532674 | 0.019028722 | 0.792331574 | 0.362237443 |
| Structural maintenance of chromosomes protein;Structural maintenance of chromosomes protein 4 | SMC4 | 0.001684974 | 0.00173726 | 0.00420577 | 0.00173381 | 0.005029132 | 0.00233395 | 0.728485431 | 0.425964122 |
| Heat shock 70 kDa protein 4L | HSPA4L | 5.88456E-05 | 4.0986E-05 | 8.8866E-05 | 2.2243E-05 | 3.07215E-05 | 4.95862E-05 | 0.170633218 | -0.937685888 |
| Ankyrin repeat and KH domain-containing protein 1 | ANKHD1 | 0 | 0 | 0 | 0 | 0 | 0 | NaN | NaN |
| Signal recognition particle 9 kDa protein | SRP9 | 0 | 0 | 0.01023838 | 0 | 0.002714268 | 0 | NaN | -1.915352096 |
| Far upstream element-binding protein 1 | FUBP1 | 0.000731945 | 0.00067439 | 0.00092218 | 0.00054913 | 0.000695571 | 0.000480985 | 0.427404792 | -0.414598024 |
| Striatin-3 | STRN3 | 0.000703281 | 0.00011169 | 0.00027621 | 0 | 0 | 0 | NaN | NaN |
| Nucleolar protein 14 | NOP14 | 4.75165E-06 | 0 | 9.4374E-05 | 2.2977E-05 | 0.000365827 | 0.000274871 | 0.304336852 | 3.698238599 |
| Protein FAM98A | FAM98A | 0.000436345 | 0.00074261 | 0.00144333 | 6.5571E-05 | 0.001548198 | 0.001209756 | 0.706668091 | 0.704029733 |
| Vesicle-fusing ATPase | NSF | 0.000438428 | 9.4429E-05 | 0.00048692 | 0 | 0 | 0 | NaN | NaN |
| Syndecan-1 | SDC1 | 0 | 0 | 0.00040991 | 0 | 0 | 0 | NaN | NaN |
| Band 4.1-like protein 2 | EPB41L2 | 0.000229448 | 0.00011254 | 0.00040779 | 0 | 3.3699E-05 | 5.18022E-05 | 0.060856853 | -2.457235751 |
| BRO1 domain-containing protein BROX | BROX | 0.000400364 | 8.5721E-05 | 0.00040893 | 0 | 0 | 0 | NaN | NaN |
| ATP-dependent 6-phosphofructokinase, liver type | PFKL | 0.015211622 | 0.01806867 | 0.01838615 | 0.00124281 | 0.002575106 | 0.000917204 | 0.013229283 | -3.861812656 |
| Probable ATP-dependent RNA helicase DDX10 | DDX10 | 2.1511E-05 | 2.5307E-05 | 4.1558E-05 | 6.1274E-05 | 0.000325257 | 0.00017366 | 0.063786273 | 2.778645688 |
| | CYHR1 | 0 | 0 | 0 | 0 | 0 | 0 | NaN | NaN |
| Mitogen-activated protein kinase;Mitogen-activated protein kinase 3 | MAPK3 | 0.000662692 | 0.00051468 | 0.00056564 | 0.00016909 | 0.000163954 | 0.000473917 | 0.099448175 | -1.742086823 |
| Methylosome subunit pICln | CLNS1A | 0 | 0.0005046 | 0.00092821 | 0 | 0.000153132 | 0 | 0.026242617 | -2.160015113 |
| Transferrin receptor protein 1;Transferrin receptor protein 1, serum form | TFRC | 0.000877393 | 0.00021589 | 0.00086707 | 0 | 0 | 0 | NaN | NaN |
| Protein arginine N-methyltransferase 1 | PRMT1 | 0.003380473 | 0.00408406 | 0.00525418 | 0.00426662 | 0.006166807 | 0.005224029 | 0.582326269 | 0.355159904 |
| Serpin H1 | SERPINH1 | 0.000511695 | 0.00030286 | 0.00074703 | 0 | 0.000281151 | 0 | 0.132401517 | -0.863938958 |
| Nucleolar transcription factor 1 | UBTF | 1.40673E-05 | 0 | 0.00016814 | 3.0695E-05 | 0 | 0 | 0.737628845 | -0.663977463 |
| NEDD8 | NEDD8;NEDD8-MDP1 | 0.004326895 | 0.00828316 | 0.00924742 | 0.00496221 | 0.008516361 | 0.007042547 | 0.926751073 | -0.234083447 |
| Ubiquitin-40S ribosomal protein S27a;Ubiquitin;40S ribosomal protein S27a;Ubiquitin-60S ribosomal protein L40;Ubiquitin;60S ribosomal protein L40;Polyubiquitin-B;Ubiquitin;Polyubiquitin-C;Ubiquitin | RPS27A;UBB;UBC;UBA52 | 0.641977382 | 0.80475243 | 0.85935702 | 0.13568779 | 0.211221387 | 0.216726706 | 0.038571398 | -1.929789115 |
| Probable RNA-binding protein EIF1AD | EIF1AD | 0.002828519 | 0.00062866 | 0.00086829 | 0.0011281 | 0.001346218 | 0 | 0.92319759 | 0.505155015 |
| 60S ribosomal protein L27a | RPL27A | 0.097626246 | 0.13016853 | 0.20514202 | 0.0177079 | 0.234633878 | 0.180342712 | 0.66005464 | 0.470360389 |
| Small acidic protein | C11orf58;SMAP | 0.016196395 | 0.00941798 | 0.01507718 | 0.00042393 | 0.008182585 | 0.003155681 | 0.143252748 | -2.256342953 |
| | HSPA8 | 0.00190365 | 0.0015191 | 9.3442E-05 | 0.00214127 | 0.002297049 | 0.003661571 | 0.234228987 | 0.59656099 |
| Lysine-rich nucleolar protein 1 | KNOP1 | 5.51008E-05 | 0 | 0.00017027 | 0 | 0 | 0 | NaN | NaN |
| Mitochondrial chaperone BCS1 | BCS1L | 0 | 0.0002221 | 0.00057977 | 0 | 0 | 0 | NaN | NaN |
| Ribosomal RNA-processing protein 8 | RRP8 | 0 | 0.00051158 | 0.00010334 | 0.00133215 | 0.002896291 | 0.001805627 | 0.063977199 | 2.973230545 |
| Prefoldin subunit 4 | PFDN4 | 0 | 0 | 0.00031233 | 0.00028681 | 0.000379623 | 0.000635576 | 0.317806258 | 0.281496253 |
| 40S ribosomal protein S30 | FAU | 0.003317602 | 0.01126557 | 0.01002649 | 0.00012456 | 0.006144306 | 0.005639762 | 0.319721766 | -0.830109884 |
| WD repeat-containing protein 74 | WDR74 | 0.000754619 | 0 | 0.00098941 | 0.00159523 | 0.0048968 | 0.003292571 | 0.081693675 | 1.929987838 |
| Gamma-tubulin complex component 2 | TUBGCP2 | 0.000308353 | 0.00012757 | 0.00065747 | 2.8874E-05 | 0 | 0 | 0.026324327 | -3.416749988 |
| Ras GTPase-activating-like protein IQGAP3 | IQGAP3 | 0 | 0 | 0 | 0 | 0 | 0 | NaN | NaN |
| NSFL1 cofactor p47 | NSFL1C | 0.000608554 | 0.00040205 | 0.00073289 | 0 | 0.000313336 | 0 | 0.108161327 | -0.957674752 |
| Non-histone chromosomal protein HMG-14 | HMGN1 | 0 | 0 | 0 | 0 | 0 | 0 | NaN | NaN |
| tRNA (uracil-5-)-methyltransferase homolog A | TRMT2A | 0.000915575 | 0.00045651 | 0.00137739 | 0.00045765 | 0.002514432 | 0.002054133 | 0.496418115 | 1.16577962 |
| 60S ribosomal protein L35 | RPL35 | 0.238775053 | 0.27552601 | 0.56192346 | 0.28327689 | 1.00273235 | 1.049362282 | 0.243509932 | 1.863676134 |
| BTB/POZ domain-containing adapter for CUL3-mediated RhoA degradation protein 3 | KCTD10 | 0.00058812 | 0.00085459 | 0.00086125 | 0 | 0.000172835 | 0.0003631 | 0.079969525 | -1.77035127 |
| Chromodomain-helicase-DNA-binding protein 4 | CHD4 | 3.45146E-05 | 8.8232E-06 | 0.00015909 | 0 | 8.35282E-05 | 3.04605E-05 | 0.797319917 | 0.547394722 |
| Proteolipid protein 2 | PLP2 | 0.000694155 | 0.00021186 | 0.00093771 | 0 | 0 | 0 | NaN | NaN |
| DNA-binding protein SMUBP-2 | IGHMBP2 | 0 | 0.00021425 | 0.00019492 | 0 | 0.000147828 | 0.000191082 | 0.680139408 | -0.282030248 |
| Multifunctional methyltransferase subunit TRM112-like protein | TRMT112 | 0.006437689 | 0.00724464 | 0.00987396 | 0.00741106 | 0.008082801 | 0.008707623 | 0.913889728 | 0.157942323 |
| Diablo homolog, mitochondrial | DIABLO | 0.000333109 | 0.00089779 | 0.00162855 | 0 | 0.000671288 | 0 | 0.758139636 | -0.419447781 |
| CD2-associated protein | CD2AP | 0.000277545 | 0.00010579 | 0.00056081 | 0 | 0 | 0 | NaN | NaN |
| Dedicator of cytokinesis protein 4 | DOCK4 | 0 | 0 | 1.5762E-05 | 0 | 0 | 0 | NaN | NaN |
| Protein C10 | C12orf57 | 0.001428955 | 0.00045061 | 0.00190209 | 0 | 0.000681735 | 0 | 0.352973632 | -1.067677527 |
| Constitutive coactivator of peroxisome proliferator-activated receptor gamma | FAM120B | 0.000455235 | 0.00041471 | 0.0005168 | 0.00042365 | 0.000727662 | 0.000314225 | 0.994923429 | -0.103731801 |
| Striatin-4 | STRN4 | 0.000598589 | 0.00024166 | 0.00033444 | 0 | 0.000156373 | 0 | 0.064164889 | -1.096764603 |
| 4F2 cell-surface antigen heavy chain | SLC3A2 | 0.000192431 | 0.00012997 | 0.00069015 | 0 | 0.000170718 | 0 | 0.45555267 | -0.172726051 |
| Gephyrin;Molybdopterin adenylyltransferase;Molybdopterin molybdenumtransferase | GPHN | 0 | 0 | 0.00018297 | 0 | 0 | 0 | NaN | NaN |
| DNA polymerase;DNA polymerase epsilon catalytic subunit A | POLE | 3.65295E-05 | 3.1453E-05 | 0.00018922 | 9.1569E-06 | 4.44462E-05 | 5.34554E-05 | 0.38484533 | 0.282998072 |
| tRNA pseudouridine synthase;tRNA pseudouridine synthase A, mitochondrial | PUS1 | 0.000104977 | 9.4395E-05 | 0.00098484 | 0.00026913 | 0.000878879 | 0.000804498 | 0.302080527 | 2.93801556 |
| C-1-tetrahydrofolate synthase, cytoplasmic;Methylenetetrahydrofolate dehydrogenase;Methenyltetrahydrofolate cyclohydrolase;Formyltetrahydrofolate synthetase;C-1-tetrahydrofolate synthase, cytoplasmic, N-terminally processed | MTHFD1 | 0.00095734 | 0.00076706 | 0.00164216 | 0.00089345 | 0.002536666 | 0.001203427 | 0.523499674 | 0.330045335 |
| Elongator complex protein 1 | IKBKAP | 0.00045449 | 0.00035596 | 0.00078077 | 0.00034004 | 0.00058546 | 0.000228099 | 0.388090263 | -0.418535601 |
| ATP-dependent RNA helicase DDX55 | DDX55 | 0.000410749 | 0.00034702 | 0.00123107 | 0.00094032 | 0.002165789 | 0.001160101 | 0.160949808 | 1.497921446 |

| | | | | | | | | | |
|---|---|---|---|---|---|---|---|---|---|
| Cleavage and polyadenylation specificity factor subunit 7 | CPSF7 | 0.001033311 | 0.00189762 | 0.00216221 | 0.00102178 | 0.00160097 | 0.00093175 | 0.372489685 | -0.893108147 |
| Actin-related protein 2 | ACTR2 | 0.000120121 | 0 | 0 | 0 | 0 | 0 | NaN | NaN |
| Protein disulfide-isomerase | P4HB | 0.00373704 | 0.00253062 | 0.00437346 | 0.00257808 | 0.003801717 | 0.001946237 | 0.500708597 | -0.535599533 |
| Dual specificity tyrosine-phosphorylation-regulated kinase 1A;Dual specificity tyrosine-phosphorylation-regulated kinase 1B | DYRK1A;DYRK1B | 0.00038882 | 0 | 0.00014769 | 0 | 0 | 0 | NaN | NaN |
| Protein-L-isoaspartate O-methyltransferase;Protein-L-isoaspartate(D-aspartate) O-methyltransferase | PCMT1 | 0.000187543 | 0.00053696 | 0.00070885 | 0.0005583 | 0.000789341 | 0.000689436 | 0.303124957 | 0.360596639 |
| Ran-specific GTPase-activating protein | RANBP1 | 0.00508784 | 0.0039127 | 0.00805882 | 0.00341719 | 0.005697352 | 0.004756694 | 0.635887502 | -0.097094083 |
| Chromobox protein homolog 5 | CBX5 | 0.001024409 | 0 | 0 | 0 | 0 | 0 | NaN | NaN |
| CCR4-NOT transcription complex subunit 2 | CNOT2 | 0 | 0.00046476 | 0.000706 | 0 | 0.000195028 | 0 | 0.048705788 | -1.554404562 |
| CAD protein;Glutamine-dependent carbamoyl-phosphate synthase;Aspartate carbamoyltransferase;Dihydroorotase | CAD | 0.000146613 | 9.4633E-05 | 0.00020366 | 0.00016483 | 0.000350866 | 0.000284101 | 0.182237419 | 0.954390878 |
| Cytokine receptor-like factor 3 | CRLF3 | 0.000668514 | 0.00094496 | 0.0018187 | 0 | 9.25222E-05 | 0 | 0.010854467 | -3.352381786 |
| Serine/threonine-protein phosphatase;Serine/threonine-protein phosphatase PP1-gamma catalytic subunit | PPP1CC | 0.002444687 | 0.00108448 | 0.00465303 | 0.00262409 | 0.00417487 | 0.003396748 | 0.449163121 | 0.474504426 |
| Clathrin light chain B | CLTB | 0.091266376 | 0.05253638 | 0.09923281 | 0.00378057 | 0.006876415 | 0.002485055 | 0.010691047 | -4.593406629 |
| Lysozyme;Lysozyme C | LYZ | 0.000546299 | 0.00133393 | 0.00168435 | 0.00043063 | 0.001244237 | 0.000880732 | 0.537119331 | -0.598904726 |
| Density-regulated protein | DENR | 0.002427556 | 0.00283754 | 0.00449247 | 0.00195657 | 0.00196823 | 0 | 0.285511 | -0.532028415 |
| | PPP1R12A | 0 | 0 | 0.00035201 | 0 | 0.000269779 | 0 | NaN | -0.383854757 |
| Kinesin-like protein;Kinesin-like protein KIF19 | KIF19 | 0.000421935 | 0.00047144 | 0.00077659 | 0.00057245 | 0.001483372 | 0.000717535 | 0.256635124 | 0.605962793 |
| Protein SCAF11 | SCAF11 | 2.50705E-05 | 5.6529E-05 | 5.4547E-05 | 2.6329E-05 | 0 | 0 | 0.164117076 | -1.050828421 |
| Cullin-2 | CUL2 | 7.02441E-05 | 6.994E-05 | 0.0001284 | 1.5803E-05 | 0 | 0 | 0.013857131 | -2.152196851 |
| Keratin, type I cytoskeletal 18 | KRT18 | 0.002489363 | 0.00818947 | 0.01063202 | 0.0105927 | 0.012340445 | 0.024233278 | 0.17492543 | 0.591552603 |
| Protein canopy homolog 2 | CNPY2 | 5.77315E-05 | 8.3938E-05 | 0.00035838 | 0 | 0.00013706 | 9.01762E-05 | 0.924819652 | 0.405417113 |
| Dynactin subunit 2 | DCTN2 | 0.002162412 | 0.00090086 | 0.00096949 | 0.0011124 | 0.000637254 | 0.000837406 | 0.318770231 | -0.211306203 |
| Syntaxin-7 | STX7 | 0.00119242 | 0.00045641 | 0.00129007 | 0 | 5.2133E-05 | 0 | 0.010497694 | -4.515551786 |
| | NAP1L1 | 0 | 0 | 0 | 0 | 0.000371399 | 0 | NaN | NaN |
| Heterogeneous nuclear ribonucleoprotein A1;Heterogeneous nuclear ribonucleoprotein A1, N-terminally processed;Heterogeneous nuclear ribonucleoprotein A1-like 2 | HNRNPA1;HNRNPA1L2 | 0.023453701 | 0.02392442 | 0.05423052 | 0.04935226 | 0.096129171 | 0.081756402 | 0.104430514 | 1.772847636 |
| Ubiquitin-associated protein 2-like | UBAP2L | 6.62524E-05 | 0 | 0.0002907 | 0 | 0.000207484 | 4.87586E-05 | 0.778753981 | -0.464428462 |
| | RPL10 | 0.036794872 | 0.03211138 | 0.07439949 | 0.05107993 | 0.088253573 | 0.047356635 | 0.436617103 | 0.473251917 |
| Actin-binding LIM protein 1 | ABLIM1 | 0 | 0 | 0 | 0 | 0 | 0 | NaN | NaN |
| Reticulon;Reticulon-4 | RTN4 | 0.00221347 | 0.00351816 | 0.0042898 | 0.00068578 | 0.00167852 | 0.001186837 | 0.06778721 | -1.567700107 |
| Insulin-like growth factor 2 mRNA-binding protein 2 | IGF2BP2 | 0.003388758 | 0.00277923 | 0.0087998 | 0.00747073 | 0.01893358 | 0.01236569 | 0.116148537 | 1.867514225 |
| Probable ATP-dependent RNA helicase DDX31 | DDX31 | 0 | 2.9277E-05 | 0.00012404 | 0.00012681 | 0.000577852 | 0.000394606 | 0.138278155 | 2.711103567 |
| Methyltransferase-like protein 2B | METTL2B | 0.001878513 | 0 | 0.00322431 | 0.00247562 | 0.001722184 | 0.003731477 | 0.958306442 | 0.008499598 |
| THUMP domain-containing protein 1 | THUMPD1 | 7.11063E-05 | 0 | 0 | 0.00123905 | 0.002269458 | 0.001560646 | 0.005935174 | 4.456022131 |
| CDK5 regulatory subunit-associated protein 2 | CDK5RAP2 | 0 | 0 | 0 | 0 | 0 | 0 | NaN | NaN |
| Protein NipSnap homolog 1 | NIPSNAP1 | 0.000620927 | 0.00102285 | 0.00267897 | 0 | 0.001155315 | 0 | 0.943765012 | 0.175690259 |
| Prolyl 3-hydroxylase 3 | LEPREL2 | 7.0552E-05 | 0.00033645 | 0.00024927 | 0 | 0 | 0 | NaN | NaN |
| Probable ATP-dependent RNA helicase DDX56 | DDX56 | 0.000251209 | 0.00052883 | 0.00093926 | 0.00033141 | 0.001523285 | 0.001337338 | 0.409264441 | 1.338490941 |
| Cleavage and polyadenylation specificity factor subunit 6 | CPSF6 | 0.003241126 | 0.00436062 | 0.00573596 | 0.00029507 | 0.000659889 | 0.000770285 | 0.01726758 | -2.724238 |
| U4/U6 small nuclear ribonucleoprotein Prp31 | PRPF31 | 0.000284363 | 0.00020674 | 0.00087372 | 0 | 0 | 0 | NaN | NaN |
| Protein lin-7 homolog C;Protein lin-7 homolog A;Protein lin-7 homolog B | LIN7C;LIN7A;LIN7B | 0.001201265 | 0.00101971 | 0.00205818 | 0 | 0.001301416 | 0.001452361 | 0.97962958 | 0.1946867 |
| 60S ribosomal protein L18 | RPL18 | 0.128238719 | 0.16936041 | 0.27572381 | 0.31947535 | 0.429126318 | 0.471327037 | 0.115989511 | 1.341305716 |
| Protein PBDC1 | PBDC1 | 0.000238904 | 0.00017871 | 0.00076507 | 0 | 0 | 0 | NaN | NaN |
| DNA-(apurinic or apyrimidinic site) lyase;DNA-(apurinic or apyrimidinic site) lyase, mitochondrial | APEX1 | 0.012895532 | 0.01205381 | 0.02457393 | 0.0307531 | 0.062510548 | 0.053073703 | 0.0653693 | 2.041125918 |
| Signal recognition particle 54 kDa protein | SRP54 | 0.000568693 | 0.00042449 | 0.00042697 | 0.00059084 | 0.001622936 | 0.000753228 | 0.158296446 | 0.818951925 |
| Unconventional myosin-XVIIIa | MYO18A | 0.000139307 | 8.443E-05 | 0.00025975 | 0 | 8.63891E-06 | 0 | 0.010184608 | -4.011274649 |
| Glucosamine 6-phosphate N-acetyltransferase | GNPNAT1 | 0 | 0 | 0.00103991 | 0 | 0 | 0 | NaN | NaN |
| Receptor expression-enhancing protein 5;Receptor expression-enhancing protein | REEP5 | 0.002322752 | 0.00402167 | 0.00324094 | 0.00039959 | 0.000629592 | 0.000511576 | 0.022237455 | -2.663389598 |
| | PSMA3 | 0.047879857 | 0.0591521 | 0.05856517 | 0.03821427 | 0.044763665 | 0.063734787 | 0.715473858 | -0.387714658 |
| ATP-dependent RNA helicase DDX24 | DDX24 | 3.97777E-05 | 0.00016405 | 0.0004549 | 0.00063969 | 0.001875276 | 0.000979418 | 0.095134542 | 2.577762734 |
| HAUS augmin-like complex subunit 4 | HAUS4 | 0.000262042 | 0.00066236 | 0.0003245 | 9.5615E-05 | 0 | 0 | 0.025760448 | -1.762936364 |
| Cyclin-dependent kinase 2 | CDK2 | 0.000737599 | 0.00049749 | 0.00108887 | 0.00015917 | 0.000619011 | 0.000685892 | 0.31022077 | -0.252871653 |
| Proteasome subunit alpha type;Proteasome subunit alpha type-6 | PSMA6 | 1.188892621 | 1.3656443 | 1.40250251 | 1.14379401 | 1.19739891 | 1.209697206 | 0.782654747 | -0.189677902 |
| UBX domain-containing protein 7 | UBXN7 | 0.000361163 | 0.00056762 | 0.00142095 | 0 | 0 | 0 | NaN | NaN |
| Catenin alpha-1 | CTNNA1 | 0.000152105 | 3.5464E-05 | 9.3287E-05 | 0 | 0.000113074 | 3.65709E-05 | 0.781862192 | -0.536731863 |
| Ubiquitin carboxyl-terminal hydrolase 14;Ubiquitin carboxyl-terminal hydrolase | USP14 | 0.034126638 | 0.02709201 | 0.0279715 | 0.00604322 | 0.009417768 | 0.006152079 | 0.037891726 | -2.184811894 |
| Lamina-associated polypeptide 2, isoforms beta/gamma;Thymopoietin;Thymopentin;Lamina-associated polypeptide 2, isoform alpha;Thymopoietin;Thymopentin | TMPO | 2.85007E-05 | 0 | 0.00020149 | 0 | 0.000237272 | 5.60997E-05 | 0.756445573 | 0.606418242 |
| Zinc phosphodiesterase ELAC protein 2 | ELAC2 | 0.000354971 | 0.00038454 | 0.00067883 | 0.00022909 | 0.000823629 | 0.000656041 | 0.846747098 | 0.770656057 |
| Protein transport protein Sec24C | SEC24C | 0.000171661 | 0.00017007 | 0.00013362 | 8.9028E-05 | 7.75291E-05 | 0 | 0.188632389 | -1.033578445 |
| Negative elongation factor A | NELFA | 0.000172898 | 0.00024657 | 0.00031919 | 4.8885E-05 | 0.000155556 | 0 | 0.136042055 | -1.499565983 |
| Importin subunit alpha-4 | KPNA3 | 0.003901635 | 0.00051087 | 0.00089715 | 0.00395904 | 0.002483894 | 0.000799182 | 0.566953946 | 1.469182485 |
| Peptidyl-prolyl cis-trans isomerase NIMA-interacting 4 | PIN4 | 0.001194323 | 0.00093212 | 0.00255859 | 0.00097343 | 0.002662434 | 0.002280791 | 0.615096422 | 0.933341191 |
| Casein kinase I isoform epsilon | CSNK1E | 0.001335909 | 0.00090848 | 0.00193853 | 0.00151395 | 0.002424023 | 0.000788979 | 0.874466156 | 0.180494869 |
| Serine/threonine-protein kinase MRCK alpha | CDC42BPA | 0.000110844 | 9.6465E-05 | 0.00011004 | 0 | 0.00027144 | 0.000150523 | 0.184701562 | 0.87731993 |
| Mitochondrial import inner membrane translocase subunit Tim8 B | TIMM8B | 0.00229252 | 0.00394316 | 0.00657284 | 0 | 0 | 0 | NaN | NaN |
| Condensin-2 complex subunit G2 | NCAPG2 | 0.000479263 | 0.00038737 | 0.00078963 | 0.00039418 | 0.000678065 | 0.000446013 | 0.85892527 | -0.103731804 |
| Calmodulin | CALM2;CALM1;CALM3 | 0.043910536 | 0.05237831 | 0.07261211 | 0.01153147 | 0.019363523 | 0.01597527 | 0.046465059 | -1.713129197 |
| DNA replication licensing factor MCM2 | MCM2 | 0.000154126 | 0.00019569 | 0.0008888 | 0.00010279 | 4.98265E-05 | 5.39792E-05 | 0.096242075 | -1.858072417 |
| Heterogeneous nuclear ribonucleoprotein D0 | HNRNPD | 0.031681223 | 0.02791793 | 0.07315645 | 0.03570499 | 0.086639611 | 0.052222763 | 0.484951229 | 0.72105076 |
| Protein transport protein Sec24A | SEC24A | 0.000653174 | 0 | 0 | 0 | 0 | 0 | NaN | NaN |
| Biorientation of chromosomes in cell division protein 1-like 1 | BOD1L1 | 0 | 0 | 0 | 0.000142 | 0 | 0 | NaN | NaN |
| Activating signal cointegrator 1 complex subunit 1 | ASCC1 | 0.004936401 | 0.00062363 | 0.00959357 | 0.00155427 | 0.00230388 | 0.003482426 | 0.762693545 | -1.09939402 |
| Double-stranded RNA-specific adenosine deaminase | ADAR | 2.44178E-05 | 5.3572E-05 | 0.00027099 | 0.00015823 | 0.000451909 | 0.000333599 | 0.164371914 | 2.638571981 |
| Engulfment and cell motility protein 2;Engulfment and cell motility protein 1 | ELMO2;ELMO1 | 0.000159803 | 0.00010924 | 0 | 0.00022667 | 0.000359411 | 5.39266E-05 | 0.796075662 | 0.77869489 |
| DCN1-like protein;DCN1-like protein 5 | DCUN1D5 | 0.001544732 | 0.00107568 | 0.00198639 | 0.00030982 | 0.001570432 | 0.001663967 | 0.462795723 | 0.023804913 |
| Gem-associated protein 2 | GEMIN2 | 0.001807972 | 0.00173422 | 0.00384172 | 0.00321099 | 0.008952197 | 0.003632947 | 0.17454196 | 1.006767984 |
| ATP synthase subunit beta;ATP synthase subunit beta, mitochondrial | ATP5B | 0.002838316 | 0.00830305 | 0.00571577 | 0.00470311 | 0.00333695 | 0.003611138 | 0.449656589 | -0.662495067 |
| F-box only protein 21 | FBXO21 | 0.000522943 | 0.00167947 | 0.00349301 | 0.00098685 | 0.000714309 | 0.000190205 | 0.239159893 | -1.233387539 |
| Serine hydroxymethyltransferase;Serine hydroxymethyltransferase, mitochondrial | SHMT2 | 0.00037011 | 0.0005909 | 0.00106695 | 0.00109079 | 0.001653876 | 0.001883867 | 0.095951641 | 1.484857843 |
| | PSMA3 | 0.00169298 | 0.00433686 | 0.00242524 | 0.00062071 | 0.001844336 | 0.001442021 | 0.160837356 | -0.750037727 |
| 60S ribosomal protein L28 | RPL28 | 0.110922629 | 0.12797027 | 0.24399489 | 0.17614363 | 0.424076827 | 0.420036997 | 0.149740187 | 1.714707712 |
| SAFB-like transcription modulator | SLTM | 0.000279364 | 0.00078163 | 0.00163522 | 0.00177412 | 0.004156655 | 0.003248953 | 0.103731875 | 2.055418973 |
| Signal recognition particle 14 kDa protein | SRP14 | 0.028209047 | 0.03150633 | 0.05322952 | 0.04995861 | 0.144426139 | 0.096512511 | 0.104171167 | 1.615074122 |
| Ankyrin repeat domain-containing protein 17 | ANKRD17 | 0.000340012 | 0.00019967 | 0.00042288 | 0.00028595 | 0.000559958 | 0.000259488 | 0.749789406 | -0.249802578 |
| Brachyury protein | T | 0 | 0 | 0.00065413 | 0 | 0 | 0 | NaN | NaN |
| 40S ribosomal protein S27;40S ribosomal protein S27-like | RPS27L | 0.002284067 | 0.00316665 | 0.00823993 | 0.00075716 | 0.010708402 | 0.005289066 | 0.910101361 | 0.740056014 |
| Acidic leucine-rich nuclear phosphoprotein 32 family member A | ANP32A | 0.018525921 | 0.01489584 | 0.03059823 | 0.01456647 | 0.013190689 | 0.013395775 | 0.307880293 | -0.467767217 |
| WD repeat-containing protein 61;WD repeat-containing protein 61, N-terminally processed | WDR61 | 0.020628149 | 0.0186936 | 0.02466253 | 0.01568779 | 0.025181822 | 0.027389738 | 0.91051159 | 0.2877683 |
| Probable ATP-dependent RNA helicase DDX17 | DDX17 | 0.008790729 | 0.012349 | 0.02657594 | 0.01694673 | 0.040965163 | 0.029101353 | 0.20534253 | 1.23669201 |
| Ankyrin repeat and fibronectin type-III domain-containing protein 1 | ANKFN1 | 0.000420821 | 0.00039722 | 0.00037652 | 0.00023499 | 0.0002448 | 0.000116429 | 0.127849578 | -0.757299815 |
| 26S proteasome non-ATPase regulatory subunit 14 | PSMD14 | 0.093864069 | 0.11447713 | 0.14392182 | 0.01308069 | 0.02105383 | 0.02180703 | 0.02153858 | -2.442904774 |
| Alpha-internexin | INA | 4.90432E-05 | 0.00018716 | 0.00023496 | 0 | 0 | 0 | NaN | NaN |
| Major vault protein | MVP | 0.003967137 | 0.00582508 | 0 | 0.00861022 | 0.000195778 | 0.001980917 | 0.458186802 | -1.279019733 |
| Rab11 family-interacting protein 1 | RAB11FIP1 | 0.011709775 | 0.00570334 | 0.01254909 | 0 | 0 | 0 | NaN | NaN |
| E3 ubiquitin-protein ligase BRE1B | RNF40 | 0.00088092 | 0.00079657 | 0.0011054 | 0.00079845 | 0.002244679 | 0.001379768 | 0.317852925 | 0.647342793 |
| Microtubule-actin cross-linking factor 1, isoforms 1/2/3/5 | MACF1 | 0.00021342 | 0.00014341 | 0.00031408 | 3.1111E-06 | 1.95818E-05 | 1.2666E-05 | 0.022277666 | -4.074662491 |
| RNA-binding protein FUS | FUS | 0.011835181 | 0.01868209 | 0.02236369 | 0.03807343 | 0.077085761 | 0.059304839 | 0.054956815 | 1.666494043 |
| Hydroxyacylglutathione hydrolase, mitochondrial | HAGH | 0.000505475 | 0.00045365 | 0.00081362 | 0.00070776 | 0.000861561 | 0.000532801 | 0.632243755 | 0.485614373 |
| Adenine phosphoribosyltransferase | APRT | 0.000366342 | 0 | 0.00024146 | 0.00023661 | 0.000265279 | 0.000166449 | 0.471800351 | -0.330009843 |
| Annexin A6;Annexin | ANXA6 | 0.042256746 | 0.02096724 | 0.0329747 | 0.00174552 | 0.003272123 | 0.001751572 | 0.012165049 | -4.23463738 |

| | | | | | | | | | |
|---|---|---|---|---|---|---|---|---|---|
| Telomere length regulation protein TEL2 homolog | TELO2 | 0.00036894 | 0.00038955 | 0.00090902 | 0.00015038 | 0.000283039 | 0.000114392 | 0.085254931 | -1.373168545 |
| Zinc finger protein 768 | ZNF768 | 0.000313319 | 0.00030118 | 0.00060928 | 9.2527E-05 | 0.000376917 | 8.10632E-05 | 0.16047492 | -1.759685939 |
| Palmitoyl-protein thioesterase 1 | PPT1 | 0.001051338 | 0.00197383 | 0.00332834 | 0 | 0 | 0 | NaN | NaN |
| Pyruvate kinase;Pyruvate kinase PKM | PKM | 0.000569925 | 0.00063428 | 0.00117262 | 6.9452E-05 | 0.000707143 | 0.000535761 | 0.285905212 | -0.243534835 |
| Ataxin-2-like protein | ATXN2L | 8.64517E-05 | 0.00016679 | 0.00063433 | 0.00080345 | 0.002710384 | 0.001323649 | 0.08267743 | 2.988459155 |
| Putative ATP-dependent RNA helicase DHX30 | DHX30 | 0.001461091 | 0.00166586 | 0.00310741 | 0.00258141 | 0.004733663 | 0.003599649 | 0.182225532 | 1.111585612 |
| Tropomyosin alpha-1 chain | TPM1 | 0.004092095 | 0.00755407 | 0.01291442 | 0 | 0.000416416 | 0.000109432 | 0.020981635 | -5.145151588 |
| Elongator complex protein 2 | ELP2 | 0 | 5.6446E-05 | 0.00014994 | 0 | 0 | 0 | NaN | NaN |
| Bifunctional purine biosynthesis protein PURH;Phosphoribosylaminoimidazolecarboxamide formyltransferase;IMP cyclohydrolase | ATIC | 0.000659501 | 0.00054693 | 0.00119061 | 0 | 0.000588554 | 0 | 0.358311478 | -0.164200058 |
| UPF0769 protein C21orf59 | C21orf59 | 0.001092767 | 0.00109965 | 0.00208339 | 0.00042018 | 0.001889742 | 0.001701295 | 0.724848129 | 0.629582351 |
| Akirin-2 | AKIRIN2 | 0.000788378 | 0.00216351 | 0.00212302 | 0 | 0 | 0 | NaN | NaN |
| Nucleolar protein 7 | NOL7 | 0 | 0.00122412 | 0.00078094 | 0.00334107 | 0.004356223 | 0.005404732 | 0.044242386 | 2.155565608 |
| Probable rRNA-processing protein EBP2 | EBNA1BP2 | 0.002267215 | 0.00244572 | 0.00516906 | 0.01542654 | 0.03449994 | 0.028499659 | 0.018184892 | 3.542611147 |
| COP9 signalosome complex subunit 3 | COPS3 | 0.000791849 | 0.00077641 | 0.00131574 | 0.00066105 | 0.001131421 | 0.00070776 | 0.71257244 | -0.161965117 |
| Peroxiredoxin-4 | PRDX4 | 0.000137056 | 0.00032837 | 0.00111782 | 8.6676E-05 | 0.000529982 | 0.000505287 | 0.7852665 | 0.621789522 |
| Drebrin-like protein | DBNL | 0.000389794 | 0.00108123 | 0.00123856 | 0 | 0 | 0 | NaN | NaN |
| DNA-directed RNA polymerase II subunit GRINL1A | GCOM1;POLR2M | 4.98175E-05 | 9.2768E-05 | 9.2127E-05 | 0.00011412 | 6.6751E-05 | 7.58037E-05 | 0.791923794 | -0.281357332 |
| Annexin | ANXA6 | 0.000951853 | 0.0002497 | 0.00052466 | 0 | 0 | 0 | NaN | NaN |
| Hepatocyte growth factor-regulated tyrosine kinase substrate | HGS | 0 | 0 | 0 | 0 | 0 | 0 | NaN | NaN |
| Clustered mitochondria protein homolog | CLUH | 7.84571E-05 | 7.7457E-05 | 0.0004224 | 0 | 0.000117387 | 6.58047E-05 | 0.598312732 | 0.163791675 |
| Gem-associated protein 4 | GEMIN4 | 0.000485377 | 0.00050337 | 0.00100521 | 0.00087876 | 0.002299728 | 0.001430961 | 0.117891141 | 1.507290345 |
| Eukaryotic translation initiation factor 5A;Eukaryotic translation initiation factor 5A-1 | EIF5A | 0.059965289 | 0.05269549 | 0.13649648 | 0.02072031 | 0.045990544 | 0.029113037 | 0.105839342 | -1.04246229 |
| 40S ribosomal protein S15a | RPS15A | 0.099860038 | 0.13609337 | 0.24242397 | 0.16848895 | 0.404856619 | 0.23910038 | 0.226081462 | 0.813019595 |
| Vacuolar protein sorting-associated protein 4A | VPS4A | 0.000926044 | 0.00024123 | 0.00038056 | 0 | 0 | 0.000205783 | 0.151903476 | -0.886981171 |
| Protein kinase C alpha type | PRKCA | 7.45829E-05 | 0.00013425 | 0.00034578 | 0 | 0 | 0 | NaN | NaN |
| Protein AHNAK2 | AHNAK2 | 2.12916E-05 | 5.9435E-06 | 3.9433E-05 | 0 | 0 | 0 | NaN | NaN |
| Protein transport protein Sec16A | SEC16A | 0.000360234 | 0.00036313 | 0.00050638 | 0.00013796 | 0.000239027 | 0.000172355 | 0.110818321 | -1.075110917 |
| Cytosolic carboxypeptidase 1;Cytosolic carboxypeptidase 4 | AGTPBP1;AGBL1 | 3.55425E-05 | 0 | 0.00013117 | 4.0291E-05 | 0.000209587 | 6.93506E-05 | 0.817873586 | 0.022441839 |
| Homeodomain-interacting protein kinase 1 | HIPK1 | 0.002662188 | 0.00109275 | 0.00275916 | 0.00063936 | 0.000133521 | 0.000584558 | 0.078417191 | -2.187194374 |
| Fructose-bisphosphate aldolase;Fructose-bisphosphate aldolase A | ALDOA | 0.024474863 | 0.01799121 | 0.04180016 | 0.02011073 | 0.028701732 | 0.02898452 | 0.933995272 | 0.229837014 |
| Obg-like ATPase 1 | OLA1 | 0.004909921 | 0.00261258 | 0.00872938 | 0.0118712 | 0.023380346 | 0.014413202 | 0.072754132 | 1.553619159 |
| DnaJ homolog subfamily B member 3;DnaJ homolog subfamily B member 6 | DNAJB6;DNAJB3 | 0.000977494 | 0.00202931 | 0.00278208 | 0 | 0 | 0 | NaN | NaN |
| Isoleucine--tRNA ligase, cytoplasmic | IARS | 0.003500616 | 0.00406103 | 0.00780838 | 0.0049277 | 0.013310028 | 0.008053159 | 0.244441101 | 0.987709844 |
| Serine/threonine-protein kinase SMG1 | SMG1 | 2.12462E-05 | 2.1671E-05 | 0.00015885 | 3.802E-05 | 0.000228419 | 7.01412E-05 | 0.449671988 | 1.694516357 |
| U3 small nucleolar RNA-associated protein 18 homolog | UTP18 | 0 | 0.00023812 | 0.0005186 | 0.00114003 | 0.002358527 | 0 | 0.076770859 | 2.222247015 |
| Probable ATP-dependent RNA helicase DHX40 | DHX40 | 0.000603852 | 0.00075294 | 0.00115326 | 0.00100354 | 0.000690401 | 0.001038984 | 0.783497854 | 0.414494664 |
| Probable ATP-dependent RNA helicase DDX5 | DDX5 | 0.003009349 | 0.0053692 | 0.00793515 | 0.00889641 | 0.01884518 | 0.014429948 | 0.085983847 | 1.426286244 |
| Ribosomal protein L19;60S ribosomal protein L19 | RPL19 | 0.157417982 | 0.15968806 | 0.31771851 | 0.28197603 | 0.658938402 | 0.521059293 | 0.108011151 | 1.706191142 |
| NAD kinase | NADK | 0 | 0 | 0 | 0 | 0 | 0 | NaN | NaN |
| Serine/arginine-rich splicing factor 1 | SRSF1 | 0.000174555 | 7.791E-05 | 0.00010664 | 4.6329E-05 | 0.000287633 | 0.000214624 | 0.738732332 | 1.009062727 |
| Small nuclear ribonucleoprotein Sm D1 | SNRPD1 | 0.000666275 | 0.00230148 | 0.00236204 | 0.00253981 | 0 | 0 | 0.287663996 | 0.142158337 |
| Polycomb protein SUZ12 | SUZ12 | 7.76453E-05 | 0.00011987 | 0.00021029 | 4.5303E-05 | 0 | 0 | 0.050838901 | -1.403755335 |
| Protein 4.1 | EPB41 | 0.000173178 | 9.8772E-05 | 0.00041119 | 0 | 0 | 0 | NaN | NaN |
| BUB3-interacting and GLEBS motif-containing protein ZNF207 | ZNF207 | 0.001480741 | 0.00236512 | 0.00146269 | 0.00182487 | 0.002740654 | 0.00117048 | 0.907636954 | 0.301472004 |
| Mitotic checkpoint protein BUB3 | BUB3 | 0.011111298 | 0.01056945 | 0.01088775 | 0.00686513 | 0.006429997 | 0.007098822 | 0.249757256 | -0.665347376 |
| Cullin-4B | CUL4B | 5.22735E-05 | 0 | 0 | 0 | 6.92892E-05 | 0 | NaN | 0.40655074 |
| Trifunctional enzyme subunit alpha, mitochondrial;Long-chain enoyl-CoA hydratase;Long chain 3-hydroxyacyl-CoA dehydrogenase | HADHA | 0.000190264 | 4.1104E-05 | 0.00010526 | 0 | 0 | 0 | NaN | NaN |
| Ras-related protein Rab-5C | RAB5C | 0.004352704 | 0.00504145 | 0.00580774 | 0.00211815 | 0.001608872 | 0.001329023 | 0.06339668 | -1.647790042 |
| Asparagine--tRNA ligase, cytoplasmic | NARS | 0.000493691 | 0.00048785 | 0.00170299 | 0.00151771 | 0.003192296 | 0.001922461 | 0.121230728 | 1.961274081 |
| E3 ubiquitin-protein ligase RNF126 | RNF126 | 0.001789049 | 0.00131205 | 0.00242999 | 0 | 0.000921164 | 0.000792639 | 0.155071918 | -1.066060005 |
| Coiled-coil and C2 domain-containing protein 1A | CC2D1A | 0.00018386 | 0.00015883 | 0.00025568 | 0.00022468 | 0.000347625 | 0.000346685 | 0.278360702 | 0.915017771 |
| Eukaryotic translation initiation factor 3 subunit G | EIF3G | 0.016376666 | 0.0182215 | 0.0320769 | 0.02884589 | 0.065467915 | 0.054047318 | 0.116413877 | 1.568581222 |
| 40S ribosomal protein S15 | RPS15 | 0.039509573 | 0.02443526 | 0.06132889 | 0.05449014 | 0.128695035 | 0.019423425 | 0.682600898 | 0.463792837 |
| Ubiquitin-protein ligase E3C | UBE3C | 0.006977382 | 0.00570187 | 0.00625555 | 0.00040297 | 0.00059142 | 0.000207419 | 0.010889832 | -3.956402671 |
| Glucosidase 2 subunit beta | PRKCSH | 0.00035809 | 9.6977E-05 | 0.00042558 | 9.5325E-05 | 0.000358259 | 0.000197761 | 0.695967335 | -0.856564287 |
| cAMP-dependent protein kinase type I-alpha regulatory subunit;cAMP-dependent protein kinase type I-alpha regulatory subunit, N-terminally processed | PRKAR1A | 0.000429885 | 0 | 0.00023183 | 0 | 0.000214972 | 0 | 0.276145334 | -0.554357148 |
| Eukaryotic translation initiation factor 1 | EIF1 | 0.006118576 | 0.00627185 | 0.00874774 | 0.00587486 | 0.017055759 | 0.008141369 | 0.436358067 | 0.376379893 |
| Proteasome assembly chaperone 2 | PSMG2 | 0.10279364 | 0.07466555 | 0.09000183 | 0.08355104 | 0.070296406 | 0.069157823 | 0.647443663 | -0.356503356 |
| Eukaryotic translation initiation factor 3 subunit K | EIF3K | 0.005474359 | 0.00908667 | 0.01489086 | 0.00600484 | 0.01952291 | 0.018489534 | 0.500063507 | 1.024884639 |
| 60S ribosomal protein L22 | RPL22 | 0.177986788 | 0.2572595 | 0.4879715 | 0.29400634 | 0.880660586 | 0.683419336 | 0.206662832 | 1.409546713 |
| Programmed cell death protein 5 | PDCD5 | 0.001337196 | 0.00212614 | 0.00288693 | 0 | 0.000774568 | 0.00080849 | 0.106901494 | -1.42585242 |
| Thioredoxin-like protein 4A | TXNL4A | 0 | 0 | 0 | 0.00101325 | 0 | 0 | NaN | NaN |
| rRNA 2-O-methyltransferase fibrillarin | FBL | 0.000396758 | 0.00113064 | 0.00328624 | 0.00126313 | 0.005370943 | 0.004369195 | 0.256698388 | 1.950227853 |
| Heterogeneous nuclear ribonucleoprotein L | HNRNPL | 0.010418766 | 0.01037632 | 0.01642707 | 0.01568672 | 0.021935147 | 0.017877519 | 0.291027583 | 0.778962122 |
| Zinc finger protein 428 | ZNF428 | 0.003202553 | 0.00137653 | 0.00225619 | 0.00099221 | 0.001566146 | 0.001469049 | 0.236200441 | -0.61900477 |
| PIH1 domain-containing protein 1 | PIH1D1 | 0.000181542 | 0.00057985 | 0.0012202 | 6.3189E-05 | 0.000592786 | 0.000448622 | 0.492158685 | -0.370176883 |
| 40S ribosomal protein S11 | RPS11 | 0.250319113 | 0.23369622 | 0.5035163 | 0.40421437 | 0.907046517 | 0.620270665 | 0.146489923 | 1.309129444 |
| Small glutamine-rich tetratricopeptide repeat-containing protein alpha | SGTA | 0.002138674 | 0.00075141 | 0.00164673 | 0 | 0 | 0 | NaN | NaN |
| 40S ribosomal protein S5;40S ribosomal protein S5, N-terminally processed | RPS5 | 0.235886239 | 0.24115985 | 0.4195269 | 0.38281998 | 0.831451494 | 0.465972155 | 0.172609265 | 0.950254039 |
| Nebulin-related-anchoring protein | NRAP | 9.73687E-06 | 3.2356E-05 | 5.081E-05 | 0 | 0 | 0 | NaN | NaN |
| General transcription factor IIF subunit 1 | GTF2F1 | 0.000107911 | 0.0001686 | 0.00035571 | 0 | 0.000254765 | 0 | 0.413966399 | 0.595596783 |
| 60S ribosomal protein L18a | RPL18A | 0.088366364 | 0.08048152 | 0.13964746 | 0.13755846 | 0.246916061 | 0.202862428 | 0.159206024 | 1.198932461 |
| Epidermal growth factor receptor substrate 15-like 1 | EPS15L1 | 0.000534699 | 0.00042377 | 0.00115773 | 0.00036984 | 0.000492332 | 0 | 0.365120894 | -0.325452786 |
| 40S ribosomal protein S16 | RPS16;ZNF90 | 0.188870227 | 0.10852088 | 0.20055713 | 0.27332151 | 0.538889112 | 0.417486126 | 0.094259186 | 1.144332957 |
| DNA polymerase;DNA polymerase delta catalytic subunit | POLD1 | 3.85069E-05 | 0.00380048 | 0 | 0.00942633 | 0.003371104 | 0 | 0.306754978 | 3.881236988 |
| Splicing factor U2AF 35 kDa subunit;Splicing factor U2AF 26 kDa subunit | U2AF1L4;U2AF1 | 0 | 0.0113692 | 0.02254909 | 0.06600656 | 0.069747258 | 0.103721157 | 0.038704266 | 2.123036301 |
| Inverted formin-2 | INF2 | 5.25053E-05 | 0.00010789 | 0.00022357 | 0 | 0 | 0 | NaN | NaN |
| SURP and G-patch domain-containing protein 2 | SUGP2 | 0 | 0 | 7.2537E-05 | 0 | 0 | 0 | NaN | NaN |
| Fibronectin type III and SPRY domain-containing protein 1 | FSD1 | 0.00105117 | 0.00079728 | 0.00268646 | 0.00164608 | 0.002894818 | 0.001408198 | 0.437501264 | 0.647037569 |
| Kinesin-like protein KIF2A | KIF2A | 0.003691748 | 0.00429436 | 0.00553493 | 0.00501607 | 0.008179773 | 0.00718781 | 0.299957558 | 0.743109807 |
| Cell division control protein 42 homolog | CDC42 | 0.000976318 | 0.00115681 | 0.00040438 | 0 | 0 | 0 | NaN | NaN |
| GTP-binding protein 1 | GTPBP1 | 0.000497581 | 0.00075633 | 0.0007768 | 0.00131516 | 0.002827179 | 0.00243248 | 0.059301593 | 1.685344008 |
| AP-3 complex subunit beta-1 | AP3B1 | 2.43702E-05 | 0 | 0 | 0 | 0 | 0 | NaN | NaN |
| Procollagen-lysine,2-oxoglutarate 5-dioxygenase 2 | PLOD2 | 0.000318503 | 0.00025117 | 0.00051014 | 0 | 5.13186E-05 | 0 | 0.011735921 | -2.633753243 |
| Spectrin alpha chain, non-erythrocytic 1 | SPTAN1 | 0.008091479 | 0.00515639 | 0.01049868 | 0.00018322 | 0.000332878 | 0.000471483 | 0.009007455 | -4.603337999 |
| Transcription elongation factor SPT5 | SUPT5H | 0.001788657 | 0.0015549 | 0.00289022 | 0.00104384 | 0.001648786 | 0.000650901 | 0.156139367 | -0.7769734 |
| Huntingtin-interacting protein 1 | HIP1 | 0.000889206 | 0.0004699 | 0.00121929 | 0 | 4.55673E-05 | 0.000298705 | 0.104785204 | -2.930121593 |
| Eukaryotic translation initiation factor 3 subunit F | EIF3F | 0.012942 | 0.01372344 | 0.02588912 | 0.03689942 | 0.067829255 | 0.031019375 | 0.087283398 | 1.426956384 |
| Importin-5 | IPO5 | 0.000691804 | 0.00044233 | 0.00091844 | 0.00119991 | 0.001875276 | 0.001105949 | 0.132926425 | 0.794495748 |
| Insulin-like growth factor 2 mRNA-binding protein 3 | IGF2BP3 | 0.003605139 | 0.00434021 | 0.01172984 | 0.01143794 | 0.030133537 | 0.020810048 | 0.092407375 | 2.261444536 |
| RNA 3-terminal phosphate cyclase | RTCA | 0.000154585 | 0 | 9.5616E-05 | 0 | 8.85308E-05 | 0 | 0.339523737 | -0.457610784 |
| Zinc finger protein 593 | ZNF593 | 0.001163476 | 0.00308364 | 0.00331026 | 0 | 0.012116768 | 0.00676643 | 0.080967186 | 1.554030063 |
| Nucleolar protein 56 | NOP56 | 0.000185382 | 0.00034168 | 0.0012297 | 0.00089622 | 0.00307751 | 0.002147795 | 0.126779718 | 2.652125924 |
| ATP-dependent RNA helicase DDX3X;ATP-dependent RNA helicase DDX3Y | DDX3X;DDX3Y | 0.003244821 | 0.00365372 | 0.00670509 | 0.00274375 | 0.008748476 | 0.004689904 | 0.79538315 | 0.360191919 |
| Insulin receptor substrate 4 | IRS4 | 0.001014388 | 0.00140354 | 0.00242771 | 0.00045288 | 0.000747057 | 0.000640463 | 0.090819148 | -1.131880992 |
| Protein arginine N-methyltransferase 5;Protein arginine N-methyltransferase 5, N-terminally processed | PRMT5 | 0.002071157 | 0.00180645 | 0.00254361 | 0.00221792 | 0.00413268 | 0.002750073 | 0.394952112 | 0.409032997 |
| Transcription elongation regulator 1 | TCERG1 | 0.002276341 | 0.00091906 | 0.00159403 | 0.00198613 | 0.00286361 | 0.001609249 | 0.378162097 | 0.317284716 |
| Proteasome subunit alpha type-7 | PSMA7 | 0.885455156 | 1.40824872 | 1.28632752 | 1.04139117 | 1.241853176 | 1.517749002 | 0.86164137 | -0.050763407 |
| Proteasome subunit alpha type-7 | PSMA7 | 0.021928675 | 0.00480477 | 0.03979633 | 0.01141966 | 0.020404227 | 0.031776847 | 0.806219819 | -0.103950586 |
| Myeloid leukemia factor 2 | MLF2 | 0.000228894 | 0.00053654 | 0.00073085 | 0 | 0 | 0 | NaN | NaN |
| Protein phosphatase 1 regulatory subunit 12A | PPP1R12A | 0.000861998 | 0.00094636 | 0.00063716 | 0 | 0.000175259 | 5.62633E-05 | 0.029860503 | -3.117807217 |
| Spectrin beta chain, non-erythrocytic 2 | SPTBN2 | 0.000991938 | 0.00031566 | 0.00128459 | 0 | 4.66053E-05 | 0 | 0.015871128 | -4.411684015 |
| Thyroid receptor-interacting protein 6 | TRIP6 | 0.000486362 | 0.00023736 | 0 | 0 | 0 | 0 | NaN | NaN |
| Tripartite motif-containing protein 26 | TRIM26 | 0.000415245 | 0.00015969 | 0.00012352 | 0 | 0 | 0 | NaN | NaN |

| | | | | | | | | | |
|---|---|---|---|---|---|---|---|---|---|
| DNA-directed RNA polymerases I and III subunit RPAC1 | POLR1C | 0.000542588 | 0.00061158 | 0.00102037 | 0.00080312 | 0.001812325 | 0.001390128 | 0.178243194 | 1.184609314 |
| NF-kappa-B-repressing factor | NKRF | 0.001187997 | 0.00125667 | 0.00341027 | 0.00564995 | 0.016158369 | 0.009199299 | 0.046166068 | 2.871914636 |
| UDP-N-acetylglucosamine--peptide N-acetylglucosaminyltransferase 110 kDa subunit | OGT | 0.001041261 | 0.00087184 | 0.00094228 | 0.00043903 | 0.000443618 | 0.00045166 | 0.123963437 | -1.086834856 |
| E3 ubiquitin-protein ligase Midline-1 | MID1 | 0.000576923 | 0.0006987 | 0.00061091 | 0.00072302 | 0.00166218 | 0.000789894 | 0.26230702 | 0.370711015 |
| Protein phosphatase 1G | PPM1G | 0.002198074 | 0.00126128 | 0.00440022 | 0.00041967 | 0.000632807 | 0.000712219 | 0.051050862 | -1.796402493 |
| Eukaryotic translation initiation factor 3 subunit D | EIF3D | 0.011441048 | 0.01165393 | 0.02331354 | 0.01812611 | 0.048795221 | 0.038533736 | 0.132156388 | 1.725305394 |
| Nuclear valosin-containing protein-like | NVL | 0.000186032 | 0.00012965 | 0.00040807 | 0 | 0.000117396 | 0 | 0.169112704 | -0.664167553 |
| Nucleoporin Nup37 | NUP37 | 0.000609842 | 0.00048562 | 0.00109005 | 0 | 7.24676E-05 | 0 | 0.009022405 | -3.073027492 |
| Thymocyte selection-associated high mobility group box protein TOX;TOX high mobility group box family member 3;TOX high mobility group box family member 4 | TOX;TOX3;TOX4 | 0.000126705 | 0.00031402 | 0.00019689 | 0.00046406 | 0.000263498 | 0.000614779 | 0.133882812 | 1.236900417 |
| DNA-directed RNA polymerase I subunit RPA34 | CD3EAP | 0.000211147 | 0.00037216 | 0.00067285 | 7.2196E-05 | 0.000721273 | 0.000183349 | 0.488682376 | -1.021311222 |
| Pre-mRNA-splicing factor ATP-dependent RNA helicase DHX15 | DHX15 | 0.007679991 | 0.00675882 | 0.0088261 | 0.0024782 | 0.004221883 | 0.002956479 | 0.09468816 | -1.377224591 |
| mRNA cap guanine-N7 methyltransferase | RNMT | 0.000776789 | 0.00050551 | 0.0005784 | 0.00161028 | 0.001133483 | 0.000833083 | 0.162712134 | 0.970623296 |
| U4/U6 small nuclear ribonucleoprotein Prp4 | PRPF4 | 0.000504552 | 0.00045853 | 0.00082505 | 0.00133 | 0.000776992 | 0.000560354 | 0.334670581 | 0.622896781 |
| D-3-phosphoglycerate dehydrogenase | PHGDH | 0.004085377 | 0.00282288 | 0.00760553 | 0.00067168 | 0.002645022 | 0.001766624 | 0.119949259 | -1.209474201 |
| Phospholipase A-2-activating protein | PLAA | 0.00022933 | 0.0003223 | 0.00054021 | 3.2379E-05 | 0 | 0 | 0.0088368 | -3.315278247 |
| 26S proteasome non-ATPase regulatory subunit 3 | PSMD3 | 0.001235416 | 0.00547388 | 0.00654197 | 0.00040302 | 0.000653648 | 0.000920592 | 0.071556799 | -3.065978763 |
| Eukaryotic translation elongation factor 1 epsilon-1 | EEF1E1;EEF1E1-BLOC1S5 | 0.002060743 | 0.00520423 | 0.00712366 | 0.00514412 | 0.010225151 | 0.010481939 | 0.217603863 | 0.974365797 |
| Heterogeneous nuclear ribonucleoprotein R | HNRNPR | 0.005258594 | 0.00738962 | 0.01833044 | 0.01463742 | 0.040201712 | 0.024276117 | 0.131107104 | 1.715966377 |
| U4/U6 small nuclear ribonucleoprotein Prp3 | PRPF3 | 0.000913783 | 0.00131058 | 0.00264243 | 0.00212159 | 0.003095458 | 0.00224574 | 0.222567307 | 0.776982186 |
| Ras-related protein Rab-7a | RAB7A | 0.002444631 | 0.00170972 | 0.00196018 | 0.00057324 | 0.000465343 | 0.000396553 | 0.036386928 | -2.074618994 |
| Eukaryotic translation initiation factor 4 gamma 3 | EIF4G3 | 7.00314E-05 | 8.5545E-05 | 0.00034066 | 0.00032798 | 0.001004326 | 0.000837698 | 0.088353433 | 3.291676477 |
| Pleiotropic regulator 1 | PLRG1 | 0.000435869 | 8.5906E-05 | 0.00045072 | 0.000124 | 0.000622078 | 0.000458514 | 0.763460485 | 0.073071185 |
| Ubiquitin-protein ligase E3A | UBE3A | 0.00687381 | 0.00703988 | 0.00897991 | 0.00060267 | 0.001189952 | 0.000973479 | 0.016453467 | -2.85432969 |
| Probable 18S rRNA (guanine-N(7))-methyltransferase | WBSCR22 | 0.000460385 | 0.00039446 | 0.00159266 | 0.00059662 | 0.003050455 | 0.000504157 | 0.63337435 | 0.373981184 |
| HIV Tat-specific factor 1 | HTATSF1 | 0.000976206 | 0.0008295 | 0.00165157 | 0.00053818 | 0.000854717 | 0.000513932 | 0.1861088 | -0.859091479 |
| COP9 signalosome complex subunit 7b | COPS7B | 0.000423206 | 0 | 0.00020961 | 0 | 0 | 0 | NaN | NaN |
| Myosin light polypeptide 6 | MYL6 | 0.011743366 | 0.01348477 | 0.01820349 | 0.0018339 | 0.002907408 | 0.002208548 | 0.021769267 | -2.610160715 |
| U3 small nucleolar RNA-interacting protein 2 | RRP9 | 8.0926E-05 | 0 | 0.00035052 | 0.00143611 | 0.002802266 | 0.00160775 | 0.04076351 | 3.254893438 |
| Putative adenosylhomocysteinase 2;Adenosylhomocysteinase;Putative adenosylhomocysteinase 3 | AHCYL1;AHCYL2 | 0.000815642 | 0.00073053 | 0.00108604 | 5.0204E-05 | 0.00026256 | 0 | 0.064358242 | -2.828679695 |
| Annexin A4;Annexin | ANXA4 | 0.003620871 | 0.0070359 | 0.00680099 | 0 | 0 | 0 | NaN | NaN |
| Double-strand-break repair protein rad21 homolog | RAD21 | 0.000111908 | 0.00012757 | 0.0003871 | 0.00012595 | 0 | 0.000235615 | 0.965471518 | 0.433312358 |
| Copine-3 | CPNE3 | 0.008155862 | 0.00480917 | 0.00973879 | 0 | 0.000437497 | 0.000327057 | 0.01223063 | -4.430357851 |
| SWI/SNF-related matrix-associated actin-dependent regulator of chromatin subfamily A member 5 | SMARCA5 | 0.000288305 | 0.00015709 | 0.00067988 | 7.9573E-05 | 0.000193783 | 0.000103957 | 0.146083469 | -1.471608951 |
| Intron-binding protein aquarius | AQR | 3.4556E-05 | 5.9433E-05 | 9.3963E-05 | 5.5305E-05 | 0.000217302 | 0.00019854 | 0.200020872 | 1.740103296 |
| Heterogeneous nuclear ribonucleoprotein Q | SYNCRIP | 0.009884112 | 0.00806804 | 0.01836241 | 0.02216847 | 0.023928155 | 0.017672281 | 0.168266457 | 1.165325744 |
| Nuclear export mediator factor NEMF | NEMF | 0.001449222 | 0.00145703 | 0.00288803 | 0.00202408 | 0.004825143 | 0.004000779 | 0.170508243 | 1.457251345 |
| Alpha-adducin | ADD1 | 0.001724387 | 0.00135423 | 0.00322724 | 0 | 0 | 0 | NaN | NaN |
| Importin subunit alpha-7 | KPNA6 | 0.000461438 | 0.00109662 | 0.00080964 | 0 | 0.000435113 | 0 | 0.1313223 | -0.895899356 |
| Eukaryotic translation initiation factor 1b | EIF1B | 0.006891725 | 0 | 0.00406731 | 0 | 0.002988173 | 0 | 0.154834199 | -0.825206724 |
| Sorting nexin-2 | SNX2 | 0.000364746 | 0.00058196 | 0.00063206 | 2.9334E-05 | 8.9889E-05 | 0 | 0.026805687 | -3.502498811 |
| H/ACA ribonucleoprotein complex subunit 4 | DKC1 | 0.000759881 | 0.00121334 | 0.00287926 | 0.00184013 | 0.004972342 | 0.002676468 | 0.212009569 | 1.141351283 |
| Endothelial differentiation-related factor 1 | EDF1 | 0.007349681 | 0.01467288 | 0.02676317 | 0.01167769 | 0.028586545 | 0.023399864 | 0.515605369 | 0.67334827 |
| DNA/RNA-binding protein KIN17 | KIN | 0 | 0.00030217 | 0 | 0.00047762 | 0.001420688 | 0.00058561 | 0.090771559 | 0.954594079 |
| DnaJ homolog subfamily A member 2 | DNAJA2 | 0.006136491 | 0.00407977 | 0.00843684 | 0.0023694 | 0.002669399 | 0.001634174 | 0.076700093 | -1.372890322 |
| mRNA-capping enzyme;Polynucleotide 5-triphosphatase;mRNA guanylyltransferase | RNGTT | 0.000398623 | 0.0002985 | 0.00042812 | 9.0519E-05 | 0.000271454 | 0.000556596 | 0.460851676 | -0.554317211 |
| WD repeat-containing protein 1 | WDR1 | 0.008925092 | 0.00618476 | 0.00801443 | 0.00022579 | 0.001041052 | 0.004956285 | 0.122122256 | -2.944557965 |
| Creatine kinase U-type, mitochondrial | CKMT1A;CKMT1B | 0.000271851 | 0.00023555 | 0.00041581 | 0 | 5.26018E-05 | 0 | 0.013389728 | -2.369632062 |
| Origin recognition complex subunit 4 | ORC4 | 4.71084E-05 | 0.0001265 | 0.00016533 | 0 | 0 | 0 | NaN | NaN |
| Protein NipSnap homolog 2 | GBAS | 0.001023346 | 0.00058618 | 0.0015775 | 0 | 0.000814254 | 0.000375075 | 0.281901429 | -0.888893103 |
| Putative PIP5K1A and PSMD4-like protein | PIPSL | 5.13498E-05 | 0 | 0.00011254 | 0 | 0 | 0 | NaN | NaN |
| Filamin-B | FLNB | 0.003191916 | 0.00221375 | 0.00265805 | 0.00018651 | 0.000310216 | 0.000353753 | 0.015024045 | -3.099023783 |
| Tropomyosin3 | TPM3 | 0.07874818 | 0.14531561 | 0.16331172 | 0.00373467 | 0.004719599 | 0.001944212 | 0.007645939 | -5.282066632 |
| Barrier-to-autointegration factor;Barrier-to-autointegration factor, N-terminally processed | BANF1 | 0.001936009 | 0.00134942 | 0.0094036 | 0.0004879 | 0.009915217 | 0 | 0.855172113 | 0.184065208 |
| Splicing factor 3B subunit 1 | SF3B1 | 0.00064864 | 0.00081312 | 0.00087128 | 0.00119701 | 0.001626284 | 0.001370071 | 0.182424578 | 0.75271757 |
| Cold shock domain-containing protein E1 | CSDE1 | 0.002143265 | 0.00144792 | 0.00518906 | 0.00068367 | 0.001569896 | 0.000591101 | 0.120194389 | -1.648427664 |
| Interferon-inducible double-stranded RNA-dependent protein kinase activator A | PRKRA | 0.00152329 | 0.00178384 | 0.00262599 | 0.00241069 | 0.009303653 | 0.004803817 | 0.128498052 | 1.429196591 |
| E3 ubiquitin-protein ligase MYCBP2 | MYCBP2 | 0 | 0 | 4.9483E-06 | 3.2129E-05 | 9.93919E-06 | 0 | 0.123526694 | 1.852534834 |
| Nucleoplasmin-3 | NPM3 | 0.000378345 | 0.00573338 | 0.00415737 | 0 | 0.003770241 | 0.009896407 | 0.399811177 | 0.555111026 |
| U5 small nuclear ribonucleoprotein 200 kDa helicase | SNRNP200 | 0.000974303 | 0.00132503 | 0.00226249 | 0.00122023 | 0.001532259 | 0.001527485 | 0.983001772 | 0.205134277 |
| Surfeit locus protein 6 | SURF6 | 0.000216101 | 0.00042963 | 0.00097086 | 0.00271472 | 0.005228968 | 0.004023561 | 0.031058644 | 3.227289473 |
| 26S protease regulatory subunit 10B | PSMC6 | 0.500979734 | 0.34585994 | 0.39378939 | 0.0903779 | 0.090625628 | 0.066079252 | 0.029827304 | -2.123382353 |
| Calcium-binding and coiled-coil domain-containing protein 2 | CALCOCO2 | 6.03236E-05 | 0.00014499 | 0.00021101 | 0 | 0 | 0 | NaN | NaN |
| Eukaryotic translation initiation factor 3 subunit J | EIF3J | 0.016773038 | 0.02042814 | 0.02972052 | 0.05023491 | 0.082127215 | 0.070731185 | 0.062971629 | 1.791788722 |
| Pre-mRNA-splicing factor SPF27 | BCAS2 | 0.000277875 | 0.00028705 | 0.00027372 | 0.00154599 | 0.00023091 | 0.000754941 | 0.214128362 | 1.441927856 |
| Dynactin subunit 3 | DCTN3 | 0.002628373 | 0.00246258 | 0.00264225 | 0.00126689 | 0.001700486 | 0.002019667 | 0.257841014 | -0.628222899 |
| DnaJ homolog subfamily C member 8 | DNAJC8 | 0.000948662 | 0.00248843 | 0.00065142 | 0.00332237 | 0.003831451 | 0.001457657 | 0.186722391 | 1.808245383 |
| Survival of motor neuron-related-splicing factor 30 | SMNDC1 | 0.000294334 | 0.00022148 | 0.00026868 | 0 | 0.00026556 | 0.000215072 | 0.849031557 | -0.168970888 |
| Neurabin-2 | PPP1R9B | 5.9719E-05 | 0.00013674 | 0.00021156 | 0 | 0 | 0 | NaN | NaN |
| Glutaredoxin-3 | GLRX3 | 0.001053969 | 0.0013422 | 0.00157777 | 0.00069772 | 0.001229812 | 0.000997722 | 0.401925232 | -0.427885571 |
| Ribosomal L1 domain-containing protein 1 | RSL1D1 | 0.00384565 | 0.00430681 | 0.01313819 | 0.00888061 | 0.021214556 | 0.017332684 | 0.1534503 | 2.008801817 |
| Nebulette | NEBL | 9.15071E-05 | 0 | 0 | 0 | 0 | 0 | NaN | NaN |
| Probable cytosolic iron-sulfur protein assembly protein CIAO1 | CIAO1 | 0.000182964 | 0.00024111 | 0.00068884 | 0.00026687 | 0.000582794 | 0.000359498 | 0.691893169 | 0.576304224 |
| Signal recognition particle subunit SRP72 | SRP72 | 0.000573116 | 0.00075287 | 0.00187433 | 0.00134054 | 0.00275137 | 0.00198559 | 0.170126391 | 1.399089328 |
| Unconventional prefoldin RPB5 interactor 1 | URI1 | 0 | 0.00028195 | 0.00048886 | 0.00022909 | 0.000834425 | 0.000574238 | 0.669414059 | 0.629215201 |
| E3 ubiquitin-protein ligase listerin | LTN1 | 0.000137667 | 0.00013999 | 0.00031859 | 0.00022218 | 0.00036217 | 0.000375348 | 0.210664481 | 1.371374191 |
| Pre-mRNA-processing factor 6 | PRPF6 | 0.000105436 | 0.00014251 | 0.00042067 | 0.00040267 | 0.000707612 | 0.000469789 | 0.1182107 | 1.720949776 |
| Transmembrane emp24 domain-containing protein 9 | TMED9 | 0.000163789 | 9.7101E-05 | 0.00033112 | 0 | 0 | 0 | NaN | NaN |
| Zyxin | ZYX | 0.002238271 | 0.00121972 | 0.00089865 | 0 | 0 | 0 | NaN | NaN |
| Centrosomal protein of 152 kDa | CEP152 | 0 | 0 | 0 | 0 | 0 | 0 | NaN | NaN |
| Utrophin | UTRN | 2.13431E-05 | 5.9335E-06 | 2.4236E-05 | 0 | 0 | 0 | NaN | NaN |
| Zinc finger Ran-binding domain-containing protein 2 | ZRANB2 | 0.001214758 | 0.00142259 | 0.00142652 | 0.00011186 | 0.000812258 | 0 | 0.141060335 | -2.238606086 |
| Kinesin-like protein KIF20A | KIF20A | 0 | 0 | 8.119E-05 | 0 | 0 | 0 | NaN | NaN |
| Chromosome-associated kinesin KIF4A | KIF4A | 0.000203874 | 9.5029E-05 | 0.0002732 | 0.00022962 | 0.000299835 | 0.000300243 | 0.259410954 | 0.556491 |
| SNARE-associated protein Snapin | SNAPIN | 0.001102452 | 0.00125353 | 0.00206969 | 0.00054603 | 0.001480425 | 0.000889144 | 0.254454518 | -0.495510984 |
| Structural maintenance of chromosomes protein 2 | SMC2 | 0.001944015 | 0.00167602 | 0.00411261 | 0.00135817 | 0.003402312 | 0.002593905 | 0.932211562 | 0.416086292 |
| Cysteine and glycine-rich protein 1 | CSRP1 | 0.000817994 | 0.00045161 | 0 | 0 | 0 | 0 | NaN | NaN |
| Urotensin-2 | UTS2 | 0.024760385 | 0.05145923 | 0.06828934 | 0.03697576 | 0.047891135 | 0.040917145 | 0.875127994 | -0.330724301 |
| CD2 antigen cytoplasmic tail-binding protein 2 | CD2BP2 | 0.002774549 | 0.00410112 | 0.00481523 | 0.00256937 | 0.00341008 | 0.002967384 | 0.517128946 | -0.466826324 |
| Proteasome assembly chaperone 1 | PSMG1 | 0.057496361 | 0.06856904 | 0.05892867 | 0.0169392 | 0.021801208 | 0.027200857 | 0.070503162 | -1.434561604 |
| E3 ubiquitin-protein ligase HERC2 | HERC2 | 7.68055E-05 | 3.6419E-05 | 0.00012402 | 6.635E-05 | 0.000127022 | 0.000102637 | 0.495027429 | 0.418269377 |
| U6 snRNA-associated Sm-like protein LSm8 | LSM8 | 0.004548203 | 0.00499246 | 0.00696931 | 0.00687868 | 0.010381725 | 0.010069321 | 0.22862864 | 1.012142761 |
| AP-2 complex subunit alpha-1 | AP2A1 | 0.003164763 | 0.00141715 | 0.00335811 | 0.00205064 | 0.003307617 | 0.002532178 | 0.909160522 | -0.321718642 |
| Double-stranded RNA-binding protein Staufen homolog 1 | STAU1 | 0.003884 | 0.00379336 | 0.0080337 | 0.00700801 | 0.013306546 | 0.009811703 | 0.146483022 | 1.336960363 |
| BAG family molecular chaperone regulator 2 | BAG2 | 0.002621543 | 0.00162661 | 0.00208814 | 0.00074191 | 0.001334432 | 0.00089039 | 0.120591161 | -1.229706383 |
| DNA topoisomerase 3-beta-1;DNA topoisomerase | TOP3B | 0 | 2.5697E-05 | 6.4313E-05 | 0.00012737 | 0.000166513 | 0 | 0.089748973 | 1.840900698 |
| L-lactate dehydrogenase A chain | LDHA | 0.008893181 | 0.00528713 | 0.0084405 | 0.00155104 | 0.002208784 | 0.001603291 | 0.038317402 | -2.396291651 |
| Glutamate dehydrogenase 1, mitochondrial;Glutamate dehydrogenase 2, mitochondrial | GLUD1;GLUD2 | 0.00386155 | 0.00395645 | 0.01221938 | 0.00054188 | 0.000925731 | 0.000385104 | 0.024988654 | -2.868162314 |
| Prelamin-A/C;Lamin-A/C | LMNA | 0 | 0.00010173 | 0.00024164 | 0 | 0.000111552 | 4.65583E-05 | 0.27045315 | -1.121357559 |
| Keratin, type II cytoskeletal 6B | KRT6B | 0.00014294 | 0.00133644 | 4.9301E-05 | 6.9901E-05 | 0.000194734 | 0.000567423 | 0.957597439 | 0.446095103 |
| Glyceraldehyde-3-phosphate dehydrogenase | GAPDH | 0.002926828 | 0.00268827 | 0.00485478 | 0.00075398 | 0.001436627 | 0.001250745 | 0.063351343 | -1.226550281 |
| Cellular tumor antigen p53 | TP53 | 9.96417E-05 | 7.4881E-05 | 0 | 4.3693E-05 | 0 | 0.000250453 | 0.847403204 | 0.276258481 |
| Guanine nucleotide-binding protein G(i) subunit alpha-2 | GNAI2 | 0.000225311 | 0.00019442 | 0.00058818 | 0 | 0.000194399 | 0.00016931 | 0.32886518 | -0.312573758 |
| Sodium/potassium-transporting ATPase subunit alpha-1 | ATP1A1 | 0.000315659 | 0.00021341 | 0.00058871 | 2.3528E-05 | 0.000230053 | 7.72856E-05 | 0.132263177 | -2.030095347 |
| Protein S100-A8;Protein S100-A8, N-terminally processed | S100A8 | 0.000307334 | 0 | 0 | 0 | 0 | 0 | NaN | NaN |
| Eukaryotic translation initiation factor 2 subunit 1 | EIF2S1 | 0.016206472 | 0.01646603 | 0.02891223 | 0.01630167 | 0.038637307 | 0.030332003 | 0.427587536 | 0.881347836 |
| 60S acidic ribosomal protein P2 | RPLP2 | 0.146808868 | 0.21963781 | 0.30476756 | 0.36742461 | 0.745489613 | 0.615110505 | 0.08661412 | 1.485719183 |
| 60S acidic ribosomal protein P0;60S acidic ribosomal protein P0-like | RPLP0;RPLP0P6 | 0.083434106 | 0.08379462 | 0.15507352 | 0.11565876 | 0.263189617 | 0.178779087 | 0.207893063 | 1.093248477 |

| | | | | | | | | | |
|---|---|---|---|---|---|---|---|---|---|
| Lupus La protein | SSB | 0.005207368 | 0.00453114 | 0.00851375 | 0.00843982 | 0.021919074 | 0.012158505 | 0.113405087 | 1.223339578 |
| Eukaryotic translation initiation factor 4E | EIF4E | 0.000194178 | 0.00023027 | 0.00060263 | 0.00033373 | 0.001620793 | 0.000681005 | 0.214180139 | 1.564320129 |
| Alpha-enolase | ENO1 | 0.019057776 | 0.01015231 | 0.0246333 | 0.00787851 | 0.014682364 | 0.007186837 | 0.184401903 | -1.274384322 |
| Nucleophosmin | NPM1 | 0.316224387 | 0.36297498 | 0.7345237 | 0.7306671 | 1.705174053 | 1.355758933 | 0.085654581 | 1.901158657 |
| Ubiquitin domain-containing protein UBFD1 | UBFD1 | 0.001798511 | 0.00185083 | 0.00270253 | 0 | 0.000179639 | 0.000113547 | 0.01385506 | -3.69590695 |
| Histone H2B type 2-E;Histone H2B type 1-B;Histone H2B type 1-O;Histone H2B type 1-J;Histone H2B type 3-B | HIST2H2BE;HIST1H2BB;HIST1 | 0.006168962 | 0.01056108 | 0.01233903 | 0.01088642 | 0.014348857 | 0.018002337 | 0.288296515 | 0.442178444 |
| L-lactate dehydrogenase B chain;L-lactate dehydrogenase | LDHB | 0.003982981 | 0.00450748 | 0.00658709 | 0.00135677 | 0.003981597 | 0.002880343 | 0.153142877 | -0.646082068 |
| Neurofilament light polypeptide | NEFL | 0.01650711 | 0.02975714 | 0.05493652 | 0.00502811 | 0.003845247 | 0.004517379 | 0.021006714 | -2.719678204 |
| Annexin A2;Annexin;Putative annexin A2-like protein | ANXA2;ANXA2P2 | 0.009388646 | 0.00860191 | 0.01565166 | 0.00056178 | 0.002597072 | 0.001488911 | 0.033577897 | -2.656659593 |
| Tubulin beta chain | TUBB | 0.014467025 | 0.01732754 | 0.02261576 | 0.02974251 | 0.025319779 | 0.023931458 | 0.318818497 | 0.54719788 |
| Bifunctional glutamate/proline--tRNA ligase;Glutamate--tRNA ligase;Proline--tRNA ligase | EPRS | 0.004076587 | 0.00528242 | 0.00710722 | 0.00909649 | 0.014646201 | 0.009800993 | 0.136354118 | 0.891727819 |
| Heat shock protein HSP 90-alpha | HSP90AA1 | 0.01720748 | 0.01303988 | 0.02650927 | 0.02137075 | 0.032910087 | 0.02346607 | 0.373891848 | 0.447540412 |
| Heat shock 70 kDa protein 1A | HSPA1A | 0.145974695 | 0.14105517 | 0.18290255 | 0.13017255 | 0.201939433 | 0.174205043 | 0.865812499 | 0.255068092 |
| Ubiquitin-like protein 4A | UBL4A | 0.002585097 | 0.00438794 | 0.00463147 | 0 | 0 | 0.000446617 | 0.010195298 | -3.296434056 |
| ATP-dependent 6-phosphofructokinase, muscle type | PFKM | 0.004046635 | 0.00210196 | 0.00309855 | 0 | 0.000106894 | 6.3616E-05 | 0.009008883 | -5.231699191 |
| Heat shock protein HSP 90-beta;Putative heat shock protein HSP 90-beta-3 | HSP90AB1;HSP90AB3P | 0.070400851 | 0.06108029 | 0.10633848 | 0.07572435 | 0.114275191 | 0.084143706 | 0.68787236 | 0.257262493 |
| U2 small nuclear ribonucleoprotein B | SNRPB2 | 0.009498376 | 0.00643306 | 0.01953877 | 0.01819384 | 0.031731426 | 0.023586798 | 0.119093328 | 1.312226808 |
| U1 small nuclear ribonucleoprotein 70 kDa | SNRNP70 | 0.010272646 | 0.01359992 | 0.0210896 | 0.01802612 | 0.040681212 | 0.021193652 | 0.197298719 | 0.640034468 |
| Vimentin | VIM | 0.02389598 | 0.04002826 | 0.06102018 | 0.01478901 | 0.010203721 | 0.012535099 | 0.059437687 | -1.675045622 |
| Annexin A5;Annexin | ANXA5 | 0.00242067 | 0.0046442 | 0.00519098 | 0.00106384 | 0.00087825 | 0.000878123 | 0.038634527 | -2.402725283 |
| U1 small nuclear ribonucleoprotein A | SNRPA | 0.007517075 | 0.00946237 | 0.01280665 | 0.01837231 | 0.033283776 | 0.018281764 | 0.107028201 | 0.957259543 |
| U1 small nuclear ribonucleoprotein C | SNRPC | 0.003429739 | 0.00456883 | 0.00591935 | 0.00813363 | 0.005449297 | 0.009218577 | 0.233851624 | 0.832076258 |
| Syntaxin-12 | STX12 | 0.000500705 | 0.00018876 | 0.00047897 | 0 | 0 | 0 | NaN | NaN |
| Beta-adducin | ADD2 | 0.000225557 | 0.0002568 | 0.00019512 | 0 | 5.8846E-05 | 6.55691E-05 | 0.058771251 | -1.860439932 |
| BRISC complex subunit Abro1 | ABRAXAS2 | 0.00035655 | 0.00025021 | 0.00034831 | 0 | 0 | 8.07322E-05 | 0.020179407 | -2.109176492 |
| HAUS augmin-like complex subunit 5 | HAUS5 | 6.82958E-05 | 0.00018679 | 0.00020838 | 0 | 0 | 0 | NaN | NaN |
| U2 small nuclear ribonucleoprotein A | SNRPA1 | 0.006273094 | 0.01229561 | 0.01378117 | 0.00866387 | 0.020414942 | 0.019285172 | 0.341666121 | 0.649348233 |
| Poly [ADP-ribose] polymerase 1 | PARP1 | 0.00355089 | 0.00275683 | 0.005098 | 0.00499425 | 0.007714335 | 0.006281764 | 0.213425996 | 0.822989074 |
| Fructose-bisphosphate aldolase C | ALDOC | 0.000116532 | 0.00019531 | 0.00041626 | 0 | 3.40325E-05 | 0 | 0.025915777 | -2.520780779 |
| 60 kDa SS-A/Ro ribonucleoprotein | TROVE2 | 0.002313459 | 0.00229656 | 0.00309124 | 0.00100744 | 0.001993946 | 0.001871716 | 0.235547456 | -0.305689983 |
| Thioredoxin | TXN | 0.017949278 | 0.02269654 | 0.0353256 | 0.02392087 | 0.036956376 | 0.028010904 | 0.640533032 | 0.303516481 |
| 60 kDa heat shock protein, mitochondrial | HSPD1 | 0 | 2.1429E-05 | 6.4137E-05 | 3.7494E-05 | 0.000169312 | 0.000131138 | 0.266326398 | 1.822652059 |
| 78 kDa glucose-regulated protein | HSPA5 | 0.057171649 | 0.06221187 | 0.07601607 | 0.03930119 | 0.053548707 | 0.044642196 | 0.357103522 | -0.478781899 |
| Heat shock cognate 71 kDa protein | HSPA8 | 0.153079162 | 0.1327227 | 0.18677505 | 0.10188572 | 0.175687441 | 0.158518158 | 0.800535015 | 0.0503702 |
| SH3 domain-binding glutamic acid-rich-like protein | SH3BGRL | 0.001401187 | 0.00057186 | 0.00119591 | 0 | 0 | 0 | NaN | NaN |
| CTTNBP2 N-terminal-like protein | CTTNBP2NL | 0.00012937 | 6.2287E-05 | 0 | 0 | 0 | 0 | NaN | NaN |
| DNA topoisomerase 1 | TOP1 | 0.000712182 | 0.00068986 | 0.00167531 | 7.9537E-05 | 0.000454039 | 0.000336092 | 0.114641354 | -1.08338977 |
| DNA topoisomerase 2-alpha | TOP2A | 0.000967137 | 0.00097293 | 0.00207882 | 5.7565E-05 | 0.000275673 | 0.000101036 | 0.031239053 | -3.267466581 |
| Ubiquilin-1 | UBQLN1 | 0.007532191 | 0.00704219 | 0.00995433 | 0.00128366 | 0.001724729 | 0.001308013 | 0.025195751 | -2.525692705 |
| Proliferating cell nuclear antigen | PCNA | 0.017299854 | 0.01404376 | 0.04269979 | 0.01484277 | 0.036500984 | 0.021511051 | 0.934963809 | 0.314318152 |
| Histidine--tRNA ligase, cytoplasmic | HARS | 0.008391557 | 0.00971517 | 0.01905379 | 0.00694361 | 0.022213739 | 0.014895726 | 0.780621173 | 0.616587592 |
| Nucleoprotein TPR | TPR | 0 | 0 | 0 | 0 | 0 | 0 | NaN | NaN |
| Creatine kinase B-type | CKB | 6.85366E-05 | 0.00028872 | 0.00054187 | 0.00011988 | 9.63662E-05 | 7.96261E-05 | 0.258175637 | -1.583051939 |
| TRPM8 channel-associated factor 1 | TCAF1 | 0.002773542 | 0.00256705 | 0.00323235 | 0.00054932 | 0.000750124 | 0.000712823 | 0.036132725 | -1.96011378 |
| UV excision repair protein RAD23 homolog A | RAD23A | 0.007008174 | 0.00383063 | 0.00658252 | 0.00043359 | 0.000300532 | 0.00034538 | 0.011285254 | -4.252383333 |
| X-ray repair cross-complementing protein 6 | XRCC6 | 0.010777069 | 0.0123668 | 0.03460225 | 0.02252862 | 0.041728613 | 0.03549411 | 0.195879558 | 1.521107865 |
| X-ray repair cross-complementing protein 5 | XRCC5 | 0.010447878 | 0.01012687 | 0.02558864 | 0.02273719 | 0.045733382 | 0.034935255 | 0.11032044 | 1.741473715 |
| LIM and SH3 domain protein 1 | LASP1 | 0.010226738 | 0.00549115 | 0.00878546 | 0 | 0 | 0 | NaN | NaN |
| Elongation factor 2 | EEF2 | 0.034578435 | 0.03033707 | 0.05767193 | 0.08561415 | 0.210203453 | 0.158539577 | 0.046410196 | 2.196898552 |
| Protein disulfide-isomerase A4 | PDIA4 | 0.002514836 | 0.00194682 | 0.00490949 | 0.00333817 | 0.004825947 | 0.004337455 | 0.352614112 | 0.78638447 |
| Prolyl 4-hydroxylase subunit alpha-1 | P4HA1 | 0.000373939 | 0.00047264 | 0.000638 | 3.8449E-05 | 0.000154498 | 4.86379E-05 | 0.034213777 | -3.280579729 |
| Plastin-3 | PLS3 | 0.019662412 | 0.00434021 | 0.00589095 | 0.00024482 | 0.000824955 | 0.000481005 | 0.025997487 | -3.614376267 |
| cAMP-dependent protein kinase type II-alpha regulatory subunit | PRKAR2A | 0.001070093 | 0.00117377 | 0.00179825 | 0.00038856 | 0.000906511 | 0.000483186 | 0.101274551 | -1.280495494 |
| Endoplasmin | HSP90B1 | 0.004399507 | 0.00404669 | 0.00827473 | 0.00323227 | 0.007899573 | 0.00590478 | 0.982627503 | 0.424541441 |
| Aspartate--tRNA ligase, cytoplasmic | DARS | 0.007375994 | 0.00930483 | 0.01684994 | 0.01340751 | 0.026397986 | 0.018104566 | 0.188744759 | 0.960302516 |
| 40S ribosomal protein S2 | RPS2 | 0.108100997 | 0.14237412 | 0.19661156 | 0.17469265 | 0.427894081 | 0.313192484 | 0.146381621 | 1.137362636 |
| Desmoplakin | DSP | 0.000249793 | 0.00040413 | 0.00043812 | 6.9934E-05 | 0.000134635 | 6.99017E-05 | 0.039528422 | -2.530773434 |
| Histone H1.3;Histone H1.4 | HIST1H1D;HIST1H1E | 0 | 0 | 0 | 0.00011888 | 0 | 0 | NaN | NaN |
| Histone H1.2 | HIST1H1C | 0.022382712 | 0.02021983 | 0.02855786 | 0.01757889 | 0.03010541 | 0.021701879 | 0.918030845 | -0.044564891 |
| Y-box-binding protein 3 | YBX3 | 0.022809876 | 0.03510102 | 0.05831583 | 0.03620169 | 0.077697861 | 0.070635771 | 0.253480961 | 1.008886222 |
| Heat shock 70 kDa protein 6 | HSPA6 | 0.035607995 | 0.0338344 | 0.04550187 | 0.03162823 | 0.048918445 | 0.040280401 | 0.911101262 | 0.177876842 |
| Tropomyosin alpha-1 chain | TPM1 | 0.00035477 | 0.00018562 | 0.00052579 | 0 | 0 | 0 | NaN | NaN |
| CTP synthase 1 | CTPS1 | 0.001644553 | 0.00148582 | 0.00208129 | 0.00018162 | 0.001235866 | 0.000655866 | 0.136123765 | -1.326222511 |
| Alpha-actinin-4 | ACTN4 | 0.130438921 | 0.08750236 | 0.1303498 | 0.00112627 | 0.00119176 | 0.000382339 | 0.006269816 | -6.854691878 |
| 26S proteasome non-ATPase regulatory subunit 9 | PSMD9 | 0.000434806 | 0.00021947 | 0.00059791 | 0 | 0 | 0 | NaN | NaN |
| T-complex protein 1 subunit alpha | TCP1 | 0.002815418 | 0.0024429 | 0.00506412 | 0.00226781 | 0.003872839 | 0.001545633 | 0.411491642 | -0.312051903 |
| 60S ribosomal protein L35a | RPL35A | 0.109097525 | 0.16569664 | 0.3156544 | 0.30852013 | 0.644057808 | 0.512803038 | 0.097644086 | 1.629860461 |
| 60S ribosomal protein L7 | RPL7 | 0.107753891 | 0.15658955 | 0.26938533 | 0.15200774 | 0.41215628 | 0.356927271 | 0.26197555 | 1.188642169 |
| Vinculin | VCL | 0.000633468 | 0.00048253 | 0.00076601 | 0.00032069 | 0.000217208 | 0.000191251 | 0.085542889 | -1.544194503 |
| Phosphoglycerate mutase 1 | PGAM1 | 0.003429347 | 0.0113556 | 0.01302676 | 0 | 0.0094542 | 0.009074287 | 0.805634097 | -0.293961559 |
| Nucleolin | NCL | 0.09021946 | 0.10094944 | 0.21692392 | 0.38667957 | 0.884464446 | 0.664648038 | 0.034851276 | 2.718957661 |
| DNA-directed RNA polymerase II subunit RPB3 | POLR2C | 0.003668346 | 0.00304878 | 0.00559805 | 0.00209181 | 0.002942768 | 0.001420037 | 0.156424085 | -0.810375835 |
| Interferon-induced, double-stranded RNA-activated protein kinase | EIF2AK2 | 0.000242386 | 0.00023219 | 0.00092438 | 0.00097026 | 0.003782698 | 0.002587674 | 0.07943816 | 3.416277598 |
| Casein kinase II subunit alpha | CSNK2A2 | 0.001767663 | 0.00111431 | 0.00177395 | 0.00273494 | 0.0102064 | 0.004540941 | 0.076717464 | 1.361148037 |
| Eukaryotic translation initiation factor 2 subunit 2 | EIF2S2 | 0.015741238 | 0.01621271 | 0.02889853 | 0.01914422 | 0.040059737 | 0.026178561 | 0.381918615 | 0.691260713 |
| Clathrin light chain A | CLTA | 0.091215989 | 0.0594138 | 0.10683167 | 0.0042319 | 0.005837318 | 0.003457502 | 0.010541391 | -4.429909972 |
| Transcription factor BTF3 | BTF3 | 0.026480797 | 0.03041976 | 0.05607361 | 0.03166801 | 0.068480197 | 0.054444553 | 0.400312882 | 0.839778806 |
| Proteasome subunit beta type-1 | PSMB1 | 0.728473855 | 0.90079556 | 1.04548361 | 0.878826 | 0.822129358 | 0.960821731 | 0.991263136 | -0.035622174 |
| Filamin-A | FLNA | 0.00661684 | 0.0042282 | 0.00460371 | 0.00026133 | 0.000448922 | 4.53841E-05 | 0.027924545 | -4.138829591 |
| Neurofibromin;Neurofibromin truncated | NF1 | 6.89956E-05 | 4.1216E-05 | 0.00025438 | 4.8089E-05 | 9.31423E-05 | 0.000250745 | 0.856345984 | 0.432932147 |
| Adenosylhomocysteinase | AHCY | 0.000288932 | 0.00023184 | 0.00061914 | 0 | 0 | 0 | NaN | NaN |
| Ubiquitin-like modifier-activating enzyme 1 | UBA1 | 0.0008522 | 0.0006181 | 0.00107133 | 0.00061003 | 0.00051545 | 0.00029557 | 0.175998682 | -0.725359562 |
| Nucleoside diphosphate kinase;Nucleoside diphosphate kinase B;Putative nucleoside diphosphate kinase | NME1-NME2;NME2;NME1;NM | 0.000258599 | 0 | 0.00054245 | 0 | 0.000121263 | 0 | 0.06967181 | -1.62696534 |
| Heterogeneous nuclear ribonucleoproteins A2/B1 | HNRNPA2B1 | 0.027038406 | 0.03770334 | 0.07641885 | 0.06999946 | 0.135559395 | 0.149895823 | 0.100744207 | 1.846160904 |
| Splicing factor, proline- and glutamine-rich | SFPQ | 0.004442 | 0.00490181 | 0.0084383 | 0.00693136 | 0.018371037 | 0.01278707 | 0.133030117 | 1.383298962 |
| Peptidyl-prolyl cis-trans isomerase B | PPIB | 0.007000336 | 0.0061142 | 0.00949219 | 0.00606494 | 0.00949706 | 0.008072048 | 0.907135196 | 0.205510584 |
| 40S ribosomal protein S3 | RPS3 | 0.222735416 | 0.23142468 | 0.38785277 | 0.31849702 | 0.809338209 | 0.574880732 | 0.151563115 | 1.312719932 |
| Cell division cycle and apoptosis regulator protein 1 | CCAR1 | 7.00873E-05 | 0.00012244 | 0.00022229 | 0 | 0 | 0 | NaN | NaN |
| 26S proteasome non-ATPase regulatory subunit 3 | PSMD3 | 0.283322136 | 0.37834188 | 0.41567266 | 0.05879482 | 0.059725961 | 0.072964658 | 0.025745718 | -2.663260423 |
| Elongation factor 1-beta | EEF1B2 | 0.0380898 | 0.0360536 | 0.05388894 | 0.01450196 | 0.048011679 | 0.023495278 | 0.247645583 | -0.697033875 |
| Low molecular weight phosphotyrosine protein phosphatase | ACP1 | 0.008789049 | 0.01327227 | 0.01743356 | 7.9347E-05 | 0.000516709 | 0.000453841 | 0.017070079 | -4.870084382 |
| DNA-directed RNA polymerase II subunit RPB1;DNA-directed RNA polymerase | POLR2A | 0.000697066 | 0.00103662 | 0.00177742 | 0.00068103 | 0.001489131 | 0.001082232 | 0.89508859 | 0.062126847 |
| DNA replication licensing factor MCM3 | MCM3 | 0.000372965 | 0.00045078 | 0.00071202 | 0.00019365 | 0.0004293 | 0.000241164 | 0.178392312 | -0.902408945 |
| Zinc-alpha-2-glycoprotein | AZGP1 | 0.000320614 | 0.00036252 | 0.00046963 | 0 | 0.000137716 | 0.000148868 | 0.099523091 | -1.34021556 |
| 40S ribosomal protein S12 | RPS12 | 0.189771582 | 0.22558359 | 0.22590191 | 0.27326775 | 0.540804436 | 0.168719696 | 0.408489728 | 0.276653127 |
| Proteasome subunit alpha type-1 | PSMA1 | 1.53073564 | 2.20157019 | 2.11069504 | 2.39316239 | 2.025153695 | 2.320124623 | 0.633309236 | 0.213070094 |
| Proteasome subunit alpha type-2 | PSMA2 | 0.732840667 | 0.8483199 | 0.91707005 | 1.07160135 | 0.925811334 | 1.017194041 | 0.621374495 | 0.261914605 |
| Proteasome subunit alpha type-3 | PSMA3 | 2.213134028 | 2.19114414 | 2.56452644 | 2.86190399 | 2.58742851 | 2.622724175 | 0.698411977 | 0.244975267 |
| Proteasome subunit alpha type-4;Proteasome subunit alpha type;Proteasome subunit beta type | PSMA4 | 1.655637667 | 1.82665131 | 2.03835967 | 2.01956674 | 1.791698477 | 1.919559926 | 0.91767381 | 0.071574333 |
| Moesin | MSN | 0 | 6.5796E-05 | 0.00019456 | 0 | 0.000184474 | 0.000422744 | 0.254954975 | 1.303467454 |
| Probable ATP-dependent RNA helicase DDX6 | DDX6 | 0.000254064 | 8.8405E-05 | 0 | 0.00019898 | 0.000911547 | 0 | 0.313331408 | 1.506773141 |
| DNA (cytosine-5)-methyltransferase 1 | DNMT1 | 0.001215541 | 0.00150811 | 0.0021781 | 0.00051349 | 0.001765982 | 0.001376964 | 0.394788044 | -0.13125369 |
| 60S ribosomal protein L13 | RPL13 | 0.126010525 | 0.19475557 | 0.32908028 | 0.35824329 | 0.812860797 | 0.825703437 | 0.076942369 | 2.061343691 |
| High mobility group protein B2 | HMGB2 | 0.004093551 | 0.00155093 | 0.00762727 | 0.00111692 | 0.00284218 | 0.003023464 | 0.385052436 | -0.526354944 |
| Polypyrimidine tract-binding protein 1 | PTBP1 | 0.027612249 | 0.01557521 | 0.06909855 | 0.02595388 | 0.121353853 | 0.051342615 | 0.409764409 | 0.894848378 |
| Threonine--tRNA ligase, cytoplasmic | TARS | 0.000341591 | 0.00041415 | 0.00102073 | 0.00014148 | 0.000509637 | 0.000205413 | 0.21918062 | -1.011636351 |
| Valine--tRNA ligase | VARS | 0.004758538 | 0.00478834 | 0.01081651 | 0.00329033 | 0.010338597 | 0.006122091 | 0.906272324 | 0.354499195 |
| Elongation factor 1-gamma | EEF1G | 0.080825216 | 0.09405946 | 0.11440314 | 0.06682686 | 0.176316953 | 0.134578911 | 0.603052361 | 0.516807428 |
| 14-3-3 protein theta | YWHAQ | 0.043990595 | 0.03162043 | 0.03968764 | 0.0124023 | 0.017733489 | 0.014400351 | 0.085883742 | -1.462585908 |
| 60S ribosomal protein L10 | RPL10 | 0.004923133 | 0.01005066 | 0.01633209 | 0.00671505 | 0.019399687 | 0.017591861 | 0.517805872 | 0.80761715 |
| Replication protein A 70 kDa DNA-binding subunit;Replication protein A 70 kDa DNA-binding subunit, N-terminally processed | RPA1 | 0.000778244 | 0.00124631 | 0.00141629 | 0.00257464 | 0.003575227 | 0.001803505 | 0.108210321 | 1.04670664 |

| | | | | | | | | | |
|---|---|---|---|---|---|---|---|---|---|
| Calreticulin | CALR | 0.003173273 | 0.00221124 | 0.00423646 | 0.00343514 | 0.004738351 | 0.00537085 | 0.348047987 | 0.578413429 |
| Proteasome subunit beta type-9;Proteasome subunit beta type | PSMB9 | 0.003386799 | 0.00418874 | 0.00450927 | 0.00338053 | 0.003507588 | 0.002387499 | 0.477005546 | -0.309265854 |
| Proteasome subunit alpha type-5 | PSMA5 | 0.698466017 | 0.79196064 | 0.87069139 | 0.7076493 | 0.742181326 | 0.729841301 | 0.845710705 | -0.117845939 |
| Proteasome subunit beta type-4 | PSMB4 | 1.327230993 | 1.39097666 | 1.68243675 | 1.59275386 | 1.545385141 | 1.631253043 | 0.824144075 | 0.195425122 |
| Proteasome subunit beta type-6 | PSMB6 | 1.301589968 | 1.67067937 | 1.90802813 | 1.61221308 | 1.639276195 | 1.637678902 | 0.969626717 | -0.028782365 |
| Proteasome subunit beta type-5 | PSMB5 | 0.820904714 | 0.93051397 | 0.91834871 | 0.78497017 | 0.996664925 | 0.934865154 | 0.969864951 | 0.025716218 |
| Proteasome subunit beta type-5 | PSMB5 | 0.003943063 | 0 | 0 | 0 | 0 | 0 | NaN | NaN |
| Mitogen-activated protein kinase 1 | MAPK1 | 8.81368E-05 | 0 | 0 | 0 | 0 | 0 | NaN | NaN |
| COMM domain-containing protein 9 | COMMD9 | 0.000271828 | 0.00012379 | 0.00034133 | 0 | 0 | 0 | NaN | NaN |
| Transketolase | TKT | 0.000761001 | 0.00134125 | 0.00165558 | 0.00022519 | 0.000332436 | 0.000150387 | 0.027332748 | -2.574374212 |
| Elongation factor 1-delta | EEF1D | 0.031156086 | 0.04098398 | 0.06388803 | 0.01963662 | 0.05935763 | 0.047413105 | 0.767518472 | 0.210225642 |
| Elongation factor 1-delta | EEF1D | 0.000582018 | 0.0002804 | 0.00029402 | 0 | 0.000186644 | 0.000369214 | 0.456897242 | -0.163531286 |
| Endoplasmic reticulum resident protein 29 | ERP29 | 0.002468201 | 0.00288276 | 0.00368052 | 0.00042143 | 0.002196461 | 0.002698666 | 0.259837341 | -0.392269348 |
| 60S ribosomal protein L12 | RPL12 | 0.150251931 | 0.16207474 | 0.24849758 | 0.24970166 | 0.600956323 | 0.420406971 | 0.117956222 | 1.375127324 |
| Phosphatidylethanolamine-binding protein 1;Hippocampal cholinergic neurostimulating peptide | PEBP1 | 0.001521666 | 0.00019261 | 0.00155101 | 0.01254529 | 0.000170316 | 0 | 0.753488647 | -0.057971946 |
| Protein disulfide-isomerase A3 | PDIA3 | 0.003936289 | 0.00310007 | 0.00645548 | 0.00561275 | 0.0047713 | 0.004347386 | 0.734067454 | 0.277546254 |
| Serine/threonine-protein phosphatase 2A 65 kDa regulatory subunit A alpha isoform | PPP2R1A | 0.007606091 | 0.00554276 | 0.00819627 | 0.00482417 | 0.004533558 | 0.004820173 | 0.308214115 | -0.658070273 |
| Cell division cycle protein 27 homolog | CDC27 | 4.63128E-05 | 4.1797E-05 | 0.00010509 | 0.00028499 | 0.000385583 | 0.00035979 | 0.024211456 | 2.957672201 |
| Glycylpeptide N-tetradecanoyltransferase 1 | NMT1 | 0.000758034 | 0.00125374 | 0.0028019 | 0.00147858 | 0.005325404 | 0.003633726 | 0.224551627 | 1.535209143 |
| Alpha-2-macroglobulin receptor-associated protein | LRPAP1 | 0.001144609 | 0.00135088 | 0.00127893 | 0.00020972 | 0.000748423 | 0.000224243 | 0.062892319 | -2.511802447 |
| CAP-Gly domain-containing linker protein 1 | CLIP1 | 5.26162E-05 | 9.0062E-06 | 0 | 0 | 0 | 2.35089E-05 | 0.936279257 | 0.110959898 |
| DnaJ homolog subfamily A member 1 | DNAJA1 | 0.009723435 | 0.00923846 | 0.01031418 | 0.00029696 | 0.002557292 | 0.002054133 | 0.06699561 | -2.242936469 |
| Heterogeneous nuclear ribonucleoprotein H3 | HNRNPH3 | 0.000409131 | 0.00086331 | 0.00240789 | 0.00085905 | 0.002212132 | 0.001840756 | 0.468165112 | 1.092347799 |
| Caspase-14;Caspase-14 subunit p17, mature form;Caspase-14 subunit p10, mature form;Caspase-14 subunit p20, intermediate form;Caspase-14 subunit p8, intermediate form | CASP14 | 0 | 0 | 0 | 0 | 0 | 0 | NaN | NaN |
| 14-3-3 protein beta/alpha;14-3-3 protein beta/alpha, N-terminally processed | YWHAB | 0.043996753 | 0.03522768 | 0.04392913 | 0.01471806 | 0.020622547 | 0.015497809 | 0.096052757 | -1.503113527 |
| Stress-induced-phosphoprotein 1 | STIP1 | 0.008819281 | 0.00471004 | 0.00802977 | 0.00335656 | 0.009334191 | 0.003905949 | 0.416093282 | -1.039686271 |
| Protein S100-A11;Protein S100-A11, N-terminally processed | S100A11 | 0.030932706 | 0.10880352 | 0.13351904 | 0.01102188 | 0.019762661 | 0.01695278 | 0.060938563 | -2.682131413 |
| Peroxiredoxin-2 | PRDX2 | 0.004874482 | 0.00380006 | 0.00829299 | 0.00247164 | 0.009702924 | 0.006399961 | 0.99983658 | 0.392814199 |
| Kinesin-1 heavy chain | KIF5B | 6.73441E-05 | 3.3425E-05 | 0.00024117 | 7.5789E-05 | 0 | 0 | 0.90802949 | 0.17043501 |
| DNA replication licensing factor MCM4 | MCM4 | 0.000115121 | 4.8271E-05 | 0.00021243 | 6.0796E-05 | 0 | 3.34067E-05 | 0.233729019 | -1.353026929 |
| Very-long-chain (3R)-3-hydroxyacyl-CoA dehydratase 3 | HACD3 | 0.00016492 | 0 | 0.00030388 | 0 | 0 | 0 | NaN | NaN |
| Heat shock 70 kDa protein 1-like | HSPA1L | 8.43579E-05 | 0 | 0 | 0 | 0 | 0 | NaN | NaN |
| Replication factor C subunit 4 | RFC4 | 0.002884 | 0.00373066 | 0.00643118 | 0.00266817 | 0.006169888 | 0.004412618 | 0.965303146 | 0.242203588 |
| Replication factor C subunit 2 | RFC2 | 0.004498264 | 0.00393677 | 0.00852133 | 0.0040559 | 0.008501226 | 0.005797099 | 0.807608741 | 0.365962789 |
| Replication factor C subunit 1 | RFC1 | 0.000735864 | 0.0001456 | 0.0003578 | 0.00023121 | 0.000966288 | 0.000666848 | 0.519520161 | 0.898188438 |
| Liprin-alpha-1;Liprin-alpha-2 | PPFIA1;PPFIA2 | 0.25867764 | 0.27195645 | 0.33239565 | 0.05602537 | 0.06670417 | 0.074686009 | 0.035403932 | -2.027526799 |
| Myosin-10 | MYH10 | 0.000785914 | 0.0004075 | 0.00086446 | 0.00013355 | 0.000148873 | 0.000136244 | 0.031882362 | -2.528178755 |
| Coatomer subunit beta | COPB2 | 0.002265144 | 0.00110614 | 0.00220285 | 0.00043932 | 0.000926548 | 0.000370519 | 0.06038601 | -2.326040311 |
| Ribonucleoside-diphosphate reductase large subunit | RRM1 | 0 | 0 | 0 | 0.00025157 | 0.00141667 | 0.001181871 | NaN | NaN |
| Glia maturation factor beta | GMFB | 0.013116112 | 0.00667214 | 0.01003745 | 0.00013082 | 0 | 0.0002037 | 0.007639121 | -5.942244894 |
| Monocarboxylate transporter 8 | SLC16A2 | 0.008081962 | 0.01049409 | 0.01754133 | 0.02329194 | 0.015101593 | 0.016312336 | 0.262285408 | 0.636386879 |
| 60S ribosomal protein L4 | RPL4 | 0.032594334 | 0.03616874 | 0.08131793 | 0.06513251 | 0.191947603 | 0.143271347 | 0.117866062 | 1.985934747 |
| Guanine nucleotide-binding protein-like 1 | GNL1 | 0.000226436 | 0.00029196 | 0.00100374 | 0.0003863 | 0.001226276 | 0.000810281 | 0.352539194 | 1.472646644 |
| Cofilin-1 | CFL1 | 0.718508566 | 0.62425416 | 0.64176637 | 0.00935731 | 0.014627449 | 0.024500049 | 0.007552083 | -5.4552981 |
| RNA-binding motif protein, X chromosome;RNA-binding motif protein, X chromosome, N-terminally processed;RNA binding motif protein, X-linked-like-1 | RBMX;RBMXL1 | 0.002512821 | 0.00461708 | 0.00698721 | 0.00481815 | 0.010670229 | 0.006886379 | 0.259484613 | 0.576763604 |
| Stress-70 protein, mitochondrial | HSPA9 | 0.00194116 | 0.00089764 | 0.00344004 | 0.00011253 | 0.000801463 | 0.000114365 | 0.082958156 | -4.085201531 |
| Eukaryotic initiation factor 4A-III;Eukaryotic initiation factor 4A-III, N-terminally processed | EIF4A3 | 0.003377393 | 0.00457155 | 0.00497114 | 0.00310832 | 0.008018242 | 0.005136014 | 0.658549397 | 0.167966367 |
| 40S ribosomal protein S19 | RPS19 | 0.180119807 | 0.2608814 | 0.39724176 | 0.48529807 | 0.77483559 | 0.812793302 | 0.089050218 | 1.570496139 |
| 60S ribosomal protein L3 | RPL3 | 0.130897996 | 0.15875641 | 0.26643529 | 0.20419287 | 0.539773108 | 0.339869536 | 0.164697945 | 1.098166191 |
| Flap endonuclease 1 | FEN1 | 0.001684078 | 0.00202449 | 0.00431181 | 0.00300059 | 0.009113192 | 0.005080518 | 0.175168941 | 1.327413519 |
| Alpha-taxilin | TXLNA | 0.00133938 | 0.00168638 | 0.00483469 | 0.0006646 | 0.003951595 | 0.00294168 | 0.87890923 | 0.802709799 |
| T-complex protein 1 subunit zeta | CCT6A | 0.005189732 | 0.0062818 | 0.0095406 | 0.01020061 | 0.007202556 | 0.00843209 | 0.553282851 | 0.424713173 |
| 60S ribosomal protein L13a | RPL13A;RPL13a | 0.075915351 | 0.10067099 | 0.18697598 | 0.1498038 | 0.323649563 | 0.301294908 | 0.12881761 | 1.581528241 |
| Malate dehydrogenase, mitochondrial;Malate dehydrogenase | MDH2 | 0.000676408 | 0.00029577 | 0.00170216 | 0.00011728 | 0.000454052 | 0.000337825 | 0.215919507 | -1.001617627 |
| Replication factor C subunit 5 | RFC5 | 0.001403426 | 0.00084022 | 0.00267906 | 0.00070848 | 0.002092525 | 0.001005394 | 0.626462991 | -0.481192011 |
| Replication factor C subunit 3 | RFC3 | 0.003617008 | 0.00405496 | 0.00613508 | 0.00469397 | 0.009718997 | 0.00658339 | 0.302822616 | 0.69914405 |
| Phospholipase D3 | PLD3 | 0.000356679 | 0.00053406 | 0.00090851 | 0 | 0 | 0 | NaN | NaN |
| Eukaryotic translation initiation factor 2 subunit 3;Putative eukaryotic translation initiation factor 2 subunit 3-like protein | EIF2S3;EIF2S3L | 0.019627141 | 0.02575212 | 0.02889214 | 0.03363866 | 0.033966864 | 0.063016259 | 0.210065726 | 0.399436823 |
| Eukaryotic translation initiation factor 2D | EIF2D | 0.000370854 | 0.00032077 | 0.00076788 | 0.00068619 | 0.001907823 | 0.001016162 | 0.120685457 | 1.454207185 |
| N-alpha-acetyltransferase 10 | NAA10 | 0.000287728 | 0 | 0.00052998 | 0 | 0.000635794 | 0 | 0.197567959 | 0.703232188 |
| Tyrosine-protein kinase CSK | CSK | 0.000967809 | 0.00068115 | 0.00084028 | 0.00057599 | 0.000826148 | 0.000697128 | 0.657748719 | -0.269443138 |
| Glycine--tRNA ligase | GARS | 5.08767E-05 | 0.00027658 | 0.00026739 | 0.00011806 | 0.000112233 | 4.59488E-05 | 0.387354874 | -1.252421598 |
| Superkiller viralicidic activity 2-like 2 | SKIV2L2 | 0.002357295 | 0.00277484 | 0.00330304 | 0.00534021 | 0.00986807 | 0.006454289 | 0.0917517 | 1.217856191 |
| 40S ribosomal protein S27 | RPS27 | 0.193192252 | 0.15140793 | 0.33317198 | 0.13946138 | 0.43290339 | 0.073289845 | 0.678186138 | -0.470171604 |
| Condensin-2 complex subunit D3 | NCAPD3 | 0.000509898 | 0.00030146 | 0.00083961 | 0.00035649 | 0.000811722 | 0.000515062 | 0.908954187 | 0.014537426 |
| RNA-binding protein 34 | RBM34 | 0.000796271 | 0.00087846 | 0.00237574 | 0.00273053 | 0.00574048 | 0.005408042 | 0.067222941 | 2.622062782 |
| Leucine-rich PPR motif-containing protein, mitochondrial | LRPPRC | 4.8335E-05 | 4.7638E-05 | 0.00013192 | 4.4197E-05 | 5.72387E-05 | 2.00058E-05 | 0.263012699 | -0.129113267 |
| DNA mismatch repair protein Msh2 | MSH2 | 0.002786418 | 0.0026466 | 0.00580674 | 0.00168145 | 0.0043068 | 0.002142343 | 0.417335919 | -0.379222216 |
| Cleavage and polyadenylation specificity factor subunit 5 | NUDT21 | 0.004909585 | 0.00463163 | 0.00593141 | 0.00083061 | 0.000626619 | 0.000915471 | 0.020818564 | -2.563361409 |
| Ran GTPase-activating protein 1 | RANGAP1 | 0.00203555 | 0.00151031 | 0.00274016 | 0.0013543 | 0.001379033 | 0.001250998 | 0.279761894 | -0.587873435 |
| ATP-dependent DNA helicase Q1 | RECQL | 0.00063789 | 0.00102116 | 0.00185177 | 0.00089154 | 0.003472094 | 0.002579106 | 0.282244448 | 1.336667788 |
| Probable 28S rRNA (cytosine(4447)-C(5))-methyltransferase | NOP2 | 0.000882376 | 0.00206145 | 0.00439775 | 0.00562683 | 0.014873897 | 0.012365885 | 0.07419983 | 2.584636219 |
| Large proline-rich protein BAG6 | BAG6 | 0.002767663 | 0.00260567 | 0.00295479 | 0.00020157 | 9.13502E-05 | 0.000330309 | 0.013025845 | -3.779315445 |
| 60S ribosomal protein L5 | RPL5 | 0.083837196 | 0.12107191 | 0.25482692 | 0.2337795 | 0.549108638 | 0.433511829 | 0.102359797 | 1.840207118 |
| 60S ribosomal protein L21 | RPL21 | 0.036042436 | 0.06046896 | 0.14093525 | 0.09335806 | 0.235477693 | 0.152989972 | 0.18570304 | 1.339170349 |
| 40S ribosomal protein S9 | RPS9 | 0.179627141 | 0.19917303 | 0.40883186 | 0.22872655 | 0.608014894 | 0.473683186 | 0.265380667 | 1.249900121 |
| 40S ribosomal protein S10;Putative 40S ribosomal protein S10-like | RPS10;RPS10-NUDT3;RPS10P | 0.055838092 | 0.08240657 | 0.12480592 | 0.11236897 | 0.256037289 | 0.250511148 | 0.11064728 | 1.604043468 |
| Microtubule-associated protein 1B;MAP1B heavy chain;MAP1 light chain LC1 | MAP1B | 9.35897E-05 | 3.1635E-05 | 0.00020986 | 0.00016374 | 0.000531348 | 0.00028044 | 0.155616015 | 1.583270472 |
| Cell division cycle protein 123 homolog | CDC123 | 0.000544648 | 0.00057232 | 0.00121902 | 0 | 0 | 0 | NaN | NaN |
| Ras GTPase-activating-like protein IQGAP1 | IQGAP1 | 0.001705744 | 0.01069193 | 0.00470929 | 0.00044548 | 0.000966167 | 0.000585941 | 0.056620076 | -3.006681953 |
| Eukaryotic translation initiation factor 1A, X-chromosomal;Eukaryotic translation initiation factor 1A, Y-chromosomal | EIF1AX;EIF1AY | 0.015716045 | 0.01693185 | 0.01756325 | 0.01870236 | 0.033060098 | 0.026381073 | 0.288925771 | 0.639763291 |
| Glutamine--tRNA ligase | QARS | 0.00324846 | 0.00245535 | 0.00432907 | 0.00271548 | 0.00734587 | 0.004641028 | 0.389987631 | 0.514688461 |
| 60S ribosomal protein L29 | RPL29 | 0.163447542 | 0.17789176 | 0.35945748 | 0.3177337 | 0.800377707 | 0.717047999 | 0.101249837 | 2.011069993 |
| Phosphatidylinositol 5-phosphate 4-kinase type-2 alpha | PIP4K2A | 0.000829974 | 0.00030209 | 0.0008788 | 0.00053959 | 0.001237634 | 0.000638263 | 0.646514266 | -0.378915125 |
| Alpha-actinin-1 | ACTN1 | 0.004864797 | 0.00155941 | 0.00227053 | 2.0953E-05 | 0 | 0 | 0.004986728 | -6.759760989 |
| Protein PRRC2A | PRRC2A | 0.000194855 | 0.00021754 | 0.00041079 | 0.00036266 | 0.001408768 | 0.000901684 | 0.110621391 | 2.051314695 |
| MARCKS-related protein | MARCKSL1 | 0.005243646 | 0.00647702 | 0.01276464 | 0 | 0.001499444 | 0.000512219 | 0.039041131 | -2.885700709 |
| 60S ribosomal protein L34 | RPL34 | 0.138299183 | 0.203915 | 0.37365056 | 0.26359189 | 0.768567257 | 0.72915977 | 0.148984573 | 1.838267055 |
| Nuclear autoantigenic sperm protein | NASP | 0.000597861 | 0.00027213 | 0.00071453 | 0.00062474 | 0.001010916 | 0.000818187 | 0.232244363 | 0.452620511 |
| Fatty acid synthase;[Acyl-carrier-protein] S-acetyltransferase;[Acyl-carrier-protein] S-malonyltransferase;3-oxoacyl-[acyl-carrier-protein] synthase;3-oxoacyl-[acyl-carrier-protein] reductase;3-hydroxyacyl-[acyl-carrier-protein] dehydratase;Enoyl-[acyl-carrier-protein] reductase;Oleoyl-[acyl-carrier-protein] hydrolase | FASN | 0.000497117 | 0.00034837 | 0.0010021 | 0.0001201 | 0.000180777 | 9.55058E-05 | 0.038885801 | -2.049361302 |
| Probable histidine--tRNA ligase, mitochondrial | HARS2 | 0 | 0 | 0 | 0 | 0.000105462 | 0 | NaN | NaN |
| Proteasome subunit beta type-3 | PSMB3 | 0.499277796 | 0.54067832 | 0.54141017 | 0.74398753 | 0.289294277 | 0.193650083 | 0.343613404 | -0.902232728 |
| Translation initiation factor eIF-2B subunit beta | EIF2B2 | 0.001811611 | 0.00092571 | 0.00169166 | 0.0017652 | 0.00432823 | 0.003055788 | 0.140513288 | 0.853103969 |
| Allograft inflammatory factor 1-like | AIF1L | 0.001794368 | 0.00348362 | 0.0032382 | 0 | 0 | 0 | NaN | NaN |
| Glycogen synthase kinase-3 beta | GSK3B | 0.000853544 | 0.00098315 | 0.0005806 | 0.00030054 | 0.000594996 | 0 | 0.196369382 | -1.013239832 |
| GMP synthase [glutamine-hydrolyzing] | GMPS | 0.00130898 | 0.00114184 | 0.00159129 | 0.00048572 | 0.00136805 | 0.001035362 | 0.291910089 | -0.33830778 |

| | | | | | | | | | |
|---|---|---|---|---|---|---|---|---|---|
| Rab GDP dissociation inhibitor beta;Rab GDP dissociation inhibitor alpha | GDI2;GDI1 | 0.000179549 | 0.00023154 | 0.00217645 | 0.00036292 | 0.000593308 | 0.000120092 | 0.685182444 | 0.648390114 |
| Hsc70-interacting protein;Putative protein FAM10A5 | ST13;ST13P5 | 0.004163028 | 0.00348404 | 0.00812595 | 0.00181842 | 0.00673752 | 0.004052965 | 0.57506911 | -0.038655545 |
| Dynamin-2 | DNM2 | 0.000432975 | 0.00042993 | 0.00048956 | 0.00019954 | 0.000339749 | 0.000465427 | 0.355702062 | -0.349814421 |
| Cyclin-dependent kinase 9 | CDK9 | 0.001105867 | 0.00142249 | 0.0015395 | 0.00174897 | 0.000983271 | 0.000534281 | 0.41079913 | -0.532752441 |
| T-complex protein 1 subunit theta | CCT8 | 0.009015228 | 0.0058523 | 0.01555119 | 0.01121002 | 0.005824728 | 0.00479408 | 0.443038927 | -0.630173298 |
| T-complex protein 1 subunit delta | CCT4 | 0.002739447 | 0.00204459 | 0.00393022 | 0.00628297 | 0.005372283 | 0.005609386 | 0.135224116 | 1.033958171 |
| Annexin A11 | ANXA11 | 0.002831542 | 0.00272909 | 0.00371943 | 4.3487E-05 | 0.000195711 | 0 | 0.017015066 | -4.93983385 |
| Fragile X mental retardation syndrome-related protein 2 | FXR2 | 0 | 0 | 0.00079408 | 0.00024088 | 0.00114606 | 0.000623036 | 0.476028674 | -0.349972467 |
| 26S proteasome non-ATPase regulatory subunit 6 | PSMD6 | 0.288791849 | 0.27408144 | 0.38270162 | 0.06948234 | 0.065257631 | 0.066342128 | 0.032280468 | -2.122032829 |
| B-cell receptor-associated protein 31 | BCAP31 | 0.00038316 | 0.00043276 | 0.00102411 | 0 | 0.000479728 | 0 | 0.67709155 | 0.148632821 |
| 26S proteasome non-ATPase regulatory subunit 13 | PSMD13 | 0.363525921 | 0.41625667 | 0.48323135 | 0.06828038 | 0.089973346 | 0.078940707 | 0.027224017 | -2.398632055 |
| Heterogeneous nuclear ribonucleoprotein A3 | HNRNPA3 | 0.003901243 | 0.00391982 | 0.0067618 | 0.00711272 | 0.01716157 | 0.009577062 | 0.112345289 | 1.288797207 |
| 6-phosphogluconate dehydrogenase, decarboxylating | PGD | 6.05867E-05 | 3.9771E-05 | 0.00028541 | 0 | 0 | 5.97469E-05 | 0.559676914 | -0.020137268 |
| Heterogeneous nuclear ribonucleoprotein M | HNRNPM | 0.004309372 | 0.00500618 | 0.00601945 | 0.00644649 | 0.011727944 | 0.010823484 | 0.171652578 | 1.11238405 |
| Importin subunit alpha-1 | KPNA2 | 0.006644833 | 0.00851471 | 0.01500228 | 0.00161737 | 0.004345508 | 0.003392269 | 0.0669139 | -1.327706309 |
| Importin subunit alpha-5;Importin subunit alpha-5, N-terminally processed | KPNA1 | 0.00297951 | 0.00312038 | 0.00489241 | 0.00294608 | 0.004141654 | 0.002757862 | 0.793852032 | -0.082923628 |
| Nuclear cap-binding protein subunit 2 | NCBP2 | 0.007135259 | 0.00985188 | 0.01580784 | 0.00305521 | 0.009636356 | 0.005924837 | 0.212322655 | -0.733623445 |
| Rap1 GTPase-GDP dissociation stimulator 1 | RAP1GDS1 | 0 | 4.8997E-05 | 0.00022509 | 5.0966E-05 | 0 | 0 | 0.389780281 | -1.043026349 |
| Heterogeneous nuclear ribonucleoprotein F;Heterogeneous nuclear ribonucleoprotein F, N-terminally processed | HNRNPF | 0.018577427 | 0.02390139 | 0.04050324 | 0.01120142 | 0.035947817 | 0.015033979 | 0.42967168 | -0.668867748 |
| DNA mismatch repair protein Msh6 | MSH6 | 0.00200084 | 0.00190537 | 0.00467148 | 0.0008935 | 0.002915846 | 0.001513407 | 0.316599802 | -0.402805782 |
| F-actin-capping protein subunit alpha-1 | CAPZA1 | 0.00169298 | 0.00112907 | 0.00231473 | 0.00030404 | 0.000678185 | 0.000843715 | 0.077269208 | -1.31981415 |
| ATP-citrate synthase | ACLY | 0.001928395 | 0.00227719 | 0.00421892 | 0.00151019 | 0.003854489 | 0.00186052 | 0.664460501 | -0.29154877 |
| Methionine aminopeptidase 1 | METAP1 | 0.000604412 | 0.00050401 | 0.00097406 | 0 | 0.000739288 | 0 | 0.704606069 | 0.290604192 |
| Coatomer subunit beta | COPB1 | 0.000217546 | 9.3728E-05 | 0.00033563 | 0 | 0.000123762 | 8.92416E-05 | 0.293277706 | -1.04964214 |
| Coatomer subunit alpha;Xenin;Proxenin | COPA | 0.000598813 | 0.00049064 | 0.0013087 | 0.00010937 | 0.000287191 | 0.000164921 | 0.051725575 | -1.860330428 |
| Arginine--tRNA ligase, cytoplasmic | RARS | 0.003260105 | 0.00421313 | 0.00845456 | 0.00581369 | 0.014269833 | 0.008696135 | 0.201774803 | 1.045482932 |
| Tyrosine--tRNA ligase, cytoplasmic;Tyrosine--tRNA ligase, cytoplasmic, N-terminally processed | YARS | 0.002151383 | 0.00241108 | 0.00461613 | 0.00699887 | 0.017116031 | 0.015028527 | 0.041458448 | 2.639955179 |
| Protein DDI1 homolog 2 | DDI2 | 0.017024409 | 0.00973307 | 0.01241575 | 0.00056225 | 0.000827621 | 0.000643151 | 0.010531263 | -4.270869643 |
| 5-AMP-activated protein kinase subunit gamma-1 | PRKAG1 | 0.002087336 | 0.00259824 | 0.00632651 | 0.00080366 | 0.000356156 | 0.000197157 | 0.039646269 | -2.866954054 |
| Heat shock-related 70 kDa protein 2 | HSPA2 | 0 | 0 | 0 | 0 | 0 | 0 | NaN | NaN |
| 26S proteasome non-ATPase regulatory subunit 2 | PSMD2 | 0.217153734 | 0.18332461 | 0.22157275 | 0.04326291 | 0.055498855 | 0.051644436 | 0.038633871 | -2.072031929 |
| 26S proteasome non-ATPase regulatory subunit 1 | PSMD1 | 0.170837532 | 0.22163718 | 0.207745 | 0.04776864 | 0.054967118 | 0.051011586 | 0.042071828 | -2.025916893 |
| Eukaryotic translation initiation factor 5 | EIF5 | 0.001176688 | 0.00066555 | 0.00106183 | 0.00045831 | 0.000893519 | 0.000633317 | 0.316483817 | -0.745555814 |
| 26S proteasome non-ATPase regulatory subunit 4 | PSMD4 | 0.216851416 | 0.17166335 | 0.1536122 | 0.05546525 | 0.041215628 | 0.05900886 | 0.048221877 | -1.629926042 |
| Exportin-2 | CSE1L | 0.003608554 | 0.00188653 | 0.00194602 | 0.00070792 | 0.0016887 | 0.000953967 | 0.111025807 | -1.028516754 |
| Transitional endoplasmic reticulum ATPase | VCP | 0.052387751 | 0.041566 | 0.05162846 | 0.01472666 | 0.018071014 | 0.01863655 | 0.045253559 | -1.514489076 |
| Mesencephalic astrocyte-derived neurotrophic factor | MANF | 0.001043388 | 0.00148142 | 0.00142999 | 0.00016997 | 0.000830166 | 0.000603933 | 0.124099472 | -1.243548682 |
| NHP2-like protein 1;NHP2-like protein 1, N-terminally processed | NHP2L1 | 0.001710391 | 0.00334733 | 0.0068651 | 0.00286491 | 0.008275539 | 0.008328303 | 0.361574201 | 1.305844648 |
| Eukaryotic translation initiation factor 3 subunit B | EIF3B | 0.006046915 | 0.00645619 | 0.01700155 | 0.01226469 | 0.030939848 | 0.029695258 | 0.132805213 | 2.201477179 |
| BH3-interacting domain death agonist;BH3-interacting domain death agonist p15;BH3-interacting domain death agonist p13;BH3-interacting domain death agonist p11 | BID | 0.000196311 | 0 | 0.00030936 | 0.00018985 | 0.00031122 | 0.000577879 | 0.583092047 | 0.33671627 |
| Ribosomal RNA processing protein 1 homolog A | RRP1 | 0.000149031 | 0.00020298 | 0.00032652 | 0.00115444 | 0.002411969 | 0.002178172 | 0.016448175 | 3.423687071 |
| Methionine--tRNA ligase, cytoplasmic | MARS | 0.002528272 | 0.00292118 | 0.00548598 | 0.00377272 | 0.009306331 | 0.006704508 | 0.188075862 | 1.198581689 |
| Eukaryotic translation initiation factor 6 | EIF6 | 0.006857575 | 0.00818256 | 0.01392639 | 0.00836639 | 0.037426501 | 0.020194723 | 0.230568223 | 1.303354176 |
| Ras-related protein Rab-25 | RAB25 | 0.002745493 | 0.00253397 | 0.00105334 | 0.00258625 | 0.002908614 | 0.003615422 | 0.271606501 | 0.198933545 |
| Coronin-7;Coronin | CORO7;CORO7-PAM16 | 0.000178821 | 0.00018991 | 0.00052523 | 0 | 0.000297398 | 0 | 0.738084316 | 0.647086667 |
| Selenocysteine-specific elongation factor | EEFSEC | 0.000426111 | 0.00054104 | 0.001156 | 0.00072485 | 0.002720028 | 0.002220621 | 0.162223188 | 2.037142688 |
| Actin-related protein 2/3 complex subunit 4;Tubulin monoglycylase TTLL3 | ARPC4;ARPC4-TTLL3;TTLL3 | 6.41585E-05 | 0.00014836 | 0.00011905 | 0 | 0.000231607 | 0.000135223 | 0.246525964 | 0.571917851 |
| Triosephosphate isomerase | TPI1 | 0.0028667 | 0.00252737 | 0.00408677 | 0.00102018 | 0.000887666 | 0.001211197 | 0.065122166 | -1.490567212 |
| Eukaryotic translation initiation factor 3 subunit E | EIF3E | 0.01871235 | 0.01883597 | 0.03466618 | 0.03673171 | 0.076754932 | 0.056290527 | 0.102894679 | 1.579401978 |
| Thioredoxin-like protein 1 | TXNL1 | 0.060211622 | 0.06215116 | 0.06585807 | 0.0132484 | 0.017587495 | 0.014156752 | 0.037337897 | -2.134290944 |
| Eukaryotic initiation factor 4A-I | EIF4A1 | 0.052194043 | 0.053557 | 0.10270344 | 0.06023652 | 0.149703326 | 0.128637913 | 0.260977234 | 1.264168904 |
| 40S ribosomal protein S20 | RPS20 | 0.228053969 | 0.20196797 | 0.39104941 | 0.15911412 | 0.448493859 | 0.216454094 | 0.900389214 | -0.075314191 |
| Ribonuclease inhibitor | RNH1 | 2.24958E-05 | 0 | 0 | 0.00130936 | 0.000820268 | 0.001100867 | 0.004132909 | 5.612840698 |
| mRNA-decapping enzyme 1A | DCP1A | 0 | 0 | 0 | 3.3089E-05 | 0.000226772 | 0.000130681 | NaN | NaN |
| NEDD8-conjugating enzyme Ubc12 | UBE2M | 0.001718229 | 0.00126442 | 0.00290383 | 0.00099768 | 0.001588112 | 0.000781949 | 0.204388287 | -0.78427616 |
| Ubiquitin-conjugating enzyme E2 N;Putative ubiquitin-conjugating enzyme E2 N-like | UBE2N;UBE2NL | 0.002799295 | 0.0025592 | 0.00445639 | 0.00310262 | 0.003894938 | 0.002563528 | 0.978162825 | 0.148422549 |
| Ras-related protein Rab-14 | RAB14 | 0.000203393 | 0.00053073 | 7.578E-05 | 0.00114057 | 0 | 0 | 0.067605602 | 2.487411794 |
| PDZ and LIM domain protein 5 | PDLIM5 | 0.001342347 | 0.00071493 | 0.00058516 | 0 | 0 | 0 | NaN | NaN |
| Alpha-centractin;Beta-centractin | ACTR1A;ACTR1B | 0.003549155 | 0.00167947 | 0.00340058 | 0.00091109 | 0.000815178 | 0.001016707 | 0.065309825 | -1.900117474 |
| ATP-binding cassette sub-family E member 1 | ABCE1 | 0.003437913 | 0.00348718 | 0.00482775 | 0.00368672 | 0.012835081 | 0.008830494 | 0.179548913 | 1.340434555 |
| 40S ribosomal protein S3a | RPS3A | 0.225053186 | 0.23191668 | 0.42542698 | 0.4557437 | 0.900188854 | 0.821224808 | 0.093419162 | 1.824170655 |
| 60S ribosomal protein L26 | RPL26;RPL26L1 | 0.130601276 | 0.18778394 | 0.30599141 | 0.33096812 | 0.828665568 | 0.613747444 | 0.080803766 | 1.708571412 |
| Proteasome activator complex subunit 3 | PSME3 | 0.176144889 | 0.24510625 | 0.33461503 | 0.20606354 | 0.163177563 | 0.166848408 | 0.408985436 | -0.554869345 |
| 60S ribosomal protein L15;Ribosomal protein L15 | RPL15 | 0.080970776 | 0.09495551 | 0.18787104 | 0.15004032 | 0.346137877 | 0.246655632 | 0.137080013 | 1.377174591 |
| 60S ribosomal protein L27 | RPL27 | 0.092386071 | 0.16352978 | 0.30475843 | 0.08828576 | 0.459972409 | 0.34972252 | 0.584482732 | 1.096657306 |
| Serine/threonine-protein kinase TBK1 | TBK1 | 3.21683E-05 | 8.6308E-05 | 9.1149E-05 | 0 | 0 | 0 | NaN | NaN |
| Small ubiquitin-related modifier 2;Small ubiquitin-related modifier 4 | SUMO2;SUMO4 | 0.387862501 | 0.22816916 | 0.22896155 | 0.17880987 | 0.265292455 | 0.273488463 | 0.698426446 | 0.212478308 |
| DDB1- and CUL4-associated factor 7 | DCAF7 | 0.010900235 | 0.01178478 | 0.00794885 | 0.00118357 | 0.001953764 | 0.000182551 | 0.050119693 | -3.203138643 |
| Heterogeneous nuclear ribonucleoprotein K | HNRNPK | 0.017804277 | 0.01568827 | 0.03689835 | 0.01947643 | 0.060934089 | 0.043129199 | 0.276871014 | 1.276441071 |
| 14-3-3 protein gamma;14-3-3 protein gamma, N-terminally processed | YWHAG | 0.014360094 | 0.0172909 | 0.02625171 | 0.006786 | 0.007435609 | 0.011131535 | 0.111159382 | -1.217490401 |
| 40S ribosomal protein S7 | RPS7 | 0.186681223 | 0.22767717 | 0.52227601 | 0.33330108 | 0.871378631 | 0.694985883 | 0.178016651 | 1.609994045 |
| Serine/threonine-protein phosphatase PP1-alpha catalytic subunit;Serine/threonine-protein phosphatase | PPP1CA;PPP1CB | 0.000274157 | 0.00022821 | 0.00101836 | 0.00057649 | 0 | 0.000421166 | 0.761030224 | 0.845837466 |
| Serine/threonine-protein phosphatase PP1-beta catalytic subunit;Serine/threonine-protein phosphatase | PPP1CB | 0.001892845 | 0.00093027 | 0.00092958 | 0.00025885 | 0.000785886 | 0.000586895 | 0.129181664 | -0.664551738 |
| 26S protease regulatory subunit 8 | PSMC5 | 0.50584481 | 0.5201298 | 0.56933053 | 0.10932645 | 0.14114464 | 0.112964658 | 0.035226966 | -2.20300022 |
| Tropomyosin alpha-4 chain | TPM4 | 0.020494905 | 0.02594264 | 0.02913508 | 0.00019576 | 0.00036213 | 0 | 0.007012406 | -6.606369675 |
| 40S ribosomal protein S8 | RPS8 | 0.120787146 | 0.17855124 | 0.30416476 | 0.20562275 | 0.568007393 | 0.465757959 | 0.177934941 | 1.383242269 |
| 14-3-3 protein epsilon | YWHAE | 0.270635987 | 0.22748875 | 0.24496301 | 0.06968876 | 0.116095418 | 0.107161912 | 0.085967934 | -1.192771688 |
| 40S ribosomal protein S14 | RPS14 | 0.309422237 | 0.32165812 | 0.63247785 | 0.81455679 | 1.613827835 | 1.324019083 | 0.064165098 | 2.041323911 |
| 40S ribosomal protein S18 | RPS18 | 0.201461203 | 0.2385638 | 0.46686455 | 0.48007311 | 0.951782055 | 0.735351962 | 0.094249043 | 1.624059778 |
| 40S ribosomal protein S13 | RPS13 | 0.095308476 | 0.13493144 | 0.19513198 | 0.17454174 | 0.329958077 | 0.355875767 | 0.141457817 | 1.290056241 |
| U6 snRNA-associated Sm-like protein LSm3 | LSM3 | 0.023553353 | 0.0188098 | 0.04361677 | 0.01450949 | 0.032193515 | 0.017888229 | 0.465566914 | -0.396921891 |
| U6 snRNA-associated Sm-like protein LSm6 | LSM6 | 0.005622551 | 0.00496713 | 0.0098493 | 0.00603924 | 0.013357844 | 0.007826891 | 0.468994358 | 0.477214511 |
| Small nuclear ribonucleoprotein Sm D2 | SNRPD2 | 0.006077147 | 0.01090652 | 0.01533017 | 0.00653024 | 0.015136417 | 0.007723104 | 0.818209324 | -0.497938407 |
| Small nuclear ribonucleoprotein Sm D3 | SNRPD3 | 0.017517075 | 0.0233759 | 0.03026852 | 0.01958286 | 0.046768728 | 0.034641223 | 0.41211681 | 0.567467768 |
| 60S ribosomal protein L7a | RPL7A | 0.14908185 | 0.16323668 | 0.30558955 | 0.23259689 | 0.632646228 | 0.513173011 | 0.138867113 | 1.652480024 |
| DNA-directed RNA polymerase II subunit RPB7 | POLR2G | 0.000739839 | 0.00137664 | 0.002411 | 0 | 0.000437779 | 0 | 0.059202431 | -1.652872487 |
| Cellular nucleic acid-binding protein | CNBP | 0.000626134 | 0.0002311 | 0.00133364 | 0 | 0 | 0.000372194 | 0.429818111 | -0.750416645 |
| 40S ribosomal protein S4, X isoform | RPS4X | 0.203963722 | 0.22724798 | 0.45581332 | 0.48726549 | 0.915846292 | 0.650959011 | 0.099094089 | 1.518299202 |
| Actin, gamma-enteric smooth muscle;Actin, alpha skeletal muscle;Actin, alpha cardiac muscle 1;Actin, aortic smooth muscle | ACTG2;ACTA1;ACTC1;ACTA2 | 0.001184414 | 0.00309986 | 0.00305526 | 0.00021574 | 0.000152744 | 5.62204E-05 | 0.019616028 | -4.322105653 |
| 40S ribosomal protein S6 | RPS6 | 0.125859366 | 0.15852612 | 0.31803818 | 0.19376445 | 0.611966087 | 0.3543764 | 0.228644401 | 1.160561987 |
| Histone H4 | HIST1H4A | 0.011743926 | 0.01561394 | 0.02869668 | 0.01137451 | 0.020886407 | 0.018927855 | 0.893368027 | 0.277676019 |
| 60S ribosomal protein L23 | RPL23 | 0.049136715 | 0.05550717 | 0.05743356 | 0.14360049 | 0.344035038 | 0.289728361 | 0.034392981 | 2.383954843 |
| 40S ribosomal protein S25 | RPS25 | 0.298387639 | 0.35890296 | 0.62631291 | 0.57708972 | 1.238424345 | 0.981851816 | 0.117721822 | 1.451911475 |
| 40S ribosomal protein S26;Putative 40S ribosomal protein S26-like 1 | RPS26;RPS26P11 | 0.247749412 | 0.17882341 | 0.31338935 | 0.67525668 | 0.901514847 | 0.501119657 | 0.072332953 | 1.446554371 |
| 40S ribosomal protein S28 | RPS28 | 0.160687493 | 0.28060295 | 0.50960818 | 1.12659249 | 0.730140234 | 1.245604128 | 0.055225345 | 2.005363662 |
| Guanine nucleotide-binding protein G(I)/G(S)/G(T) subunit beta-1;Guanine nucleotide-binding protein G(I)/G(S)/G(T) subunit beta-3;Guanine nucleotide-binding protein subunit beta-4;Guanine nucleotide-binding protein G(I)/G(S)/G(T) subunit beta-2 | GNB1;GNB4;GNB2;GNB3 | 0.001121319 | 0.00094936 | 0.00231537 | 0 | 0.000534697 | 0.000398131 | 0.085040523 | -1.281142463 |
| E3 ubiquitin-protein ligase RBX1;E3 ubiquitin-protein ligase RBX1, N-terminally processed | RBX1 | 0.012162692 | 0.01023249 | 0.00753311 | 0.0058683 | 0.006340258 | 0 | 0.283528181 | -0.746343423 |
| 60S ribosomal protein L30 | RPL30 | 0.088276789 | 0.09632576 | 0.17111152 | 0.23609095 | 0.523606702 | 0.437172622 | 0.052946522 | 2.182209456 |
| 60S ribosomal protein L31 | RPL31 | 0.226514388 | 0.19589658 | 0.40291351 | 0.37048863 | 0.835616989 | 0.62640444 | 0.115647167 | 1.467491747 |
| 60S ribosomal protein L10a | RPL10A | 0.088814243 | 0.13596776 | 0.21699699 | 0.23919798 | 0.478897952 | 0.354103787 | 0.092815395 | 1.380907676 |

| | | | | | | | | | |
|---|---|---|---|---|---|---|---|---|---|
| 60S ribosomal protein L11 | RPL11 | 0.135466353 | 0.13696221 | 0.28667458 | 0.23473633 | 0.561283669 | 0.408275728 | 0.125685129 | 1.575765905 |
| 60S ribosomal protein L8 | RPL8 | 0.133366924 | 0.13533968 | 0.3043109 | 0.26189324 | 0.647459852 | 0.416804595 | 0.117712351 | 1.622786257 |
| Peptidyl-prolyl cis-trans isomerase A;Peptidyl-prolyl cis-trans isomerase A, N-terminally processed;Peptidyl-prolyl cis-trans isomerase | PPIA | 0.00371487 | 0.00541579 | 0.01371267 | 0.00138515 | 0.007066742 | 0.002605589 | 0.262481989 | -1.055561199 |
| UV excision repair protein RAD23 homolog B | RAD23B | 0.177421341 | 0.15117764 | 0.20600055 | 0.01379132 | 0.019822933 | 0.016442411 | 0.013868743 | -3.431685787 |
| Growth factor receptor-bound protein 2 | GRB2 | 0.003710503 | 0.00192515 | 0.00328304 | 8.5186E-05 | 0.000395039 | 0.000305384 | 0.02518767 | -3.426335946 |
| AP-2 complex subunit beta | AP2B1 | 0.001410144 | 0.00093117 | 0.00227473 | 0.0005693 | 0.001016916 | 0.000818693 | 0.17168488 | -0.784448029 |
| 14-3-3 protein zeta/delta | YWHAZ | 0.084996081 | 0.05919188 | 0.09810028 | 0.03247111 | 0.045044936 | 0.03637231 | 0.128755104 | -1.224555769 |
| Serine/threonine-protein phosphatase 2A 55 kDa regulatory subunit B alpha isoform;Serine/threonine-protein phosphatase 2A 55 kDa regulatory subunit B gamma isoform | PPP2R2A;PPP2R2C | 0.000644105 | 0.00049641 | 0.00097561 | 0.00019423 | 0.00022562 | 0.000446734 | 0.090831725 | -1.513400931 |
| Dynein light chain 2, cytoplasmic;Dynein light chain 1, cytoplasmic | DYNLL2;DYNLL1 | 0.014363453 | 0.00079762 | 0.00835684 | 0 | 0.001636196 | 0 | 0.302579685 | -2.352611245 |
| 60S ribosomal protein L38 | RPL38 | 0.044493338 | 0.06451586 | 0.12911681 | 0.0814245 | 0.305380319 | 0.260383604 | 0.162198108 | 2.012912861 |
| 40S ribosomal protein S21 | RPS21 | 0.034684246 | 0.02956349 | 0.05342132 | 0.08281352 | 0.10288638 | 0.06248856 | 0.123935066 | 1.255585884 |
| Guanine nucleotide-binding protein subunit beta-2-like 1;Guanine nucleotide-binding protein subunit beta-2-like 1, N-terminally processed | GNB2L1 | 0.121895644 | 0.14840364 | 0.24294456 | 0.23352148 | 0.481563333 | 0.417213514 | 0.116348097 | 1.491259382 |
| Serine/threonine-protein phosphatase 2A catalytic subunit alpha isoform;Serine/threonine-protein phosphatase;Serine/threonine-protein phosphatase 2A catalytic subunit beta isoform | PPP2CA;PPP2CB | 0.00086922 | 0.00129708 | 0.00316367 | 0.00081619 | 0.002050468 | 0.001108558 | 0.675876419 | -0.226582097 |
| Nuclease-sensitive element-binding protein 1 | YBX1 | 0.078378681 | 0.06843295 | 0.17277377 | 0.14638499 | 0.265854998 | 0.237698374 | 0.12173255 | 1.600598835 |
| Casein kinase II subunit beta | CSNK2B-LY6G5B-1181;CSNK2 | 0.005292744 | 0.00523291 | 0.00725582 | 0.00609687 | 0.012271199 | 0.007568689 | 0.366550326 | 0.516027552 |
| Elongation factor Tu GTP-binding domain-containing protein 1 | EFTUD1 | 0 | 0 | 0 | 0.00012172 | 0.000600916 | 0.000306903 | NaN | NaN |
| Putative elongation factor 1-alpha-like 3;Elongation factor 1-alpha 1;Elongation factor 1-alpha | EEF1A1P5;EEF1A1 | 0.240129885 | 0.25632786 | 0.4690474 | 0.14524539 | 0.423340164 | 0.269262973 | 0.657597682 | 0.071025591 |
| Tubulin alpha-1B chain;Tubulin alpha-1A chain;Tubulin alpha-1C chain;Tubulin alpha-4A chain;Tubulin alpha-3C/D chain;Tubulin alpha-3E chain | TUBA1B;TUBA1A;TUBA1C;TUE | 0.11419214 | 0.08213231 | 0.09814595 | 0.20853626 | 0.126100642 | 0.088359459 | 0.418582912 | 0.361574994 |
| Tubulin beta-4B chain;Tubulin beta-4A chain | TUBB4B;TUBB4A | 0.026495353 | 0.02712865 | 0.04944378 | 0.03577165 | 0.051803485 | 0.040434232 | 0.516766669 | 0.5757599 |
| Immunoglobulin-binding protein 1 | IGBP1 | 0.00020884 | 0.00012275 | 0 | 4.9895E-05 | 0 | 0 | 0.042591688 | -1.682069977 |
| Ribonuclease P protein subunit p38 | RPP38 | 0.000400224 | 0.00057322 | 0.00112622 | 0.00047883 | 0.002030913 | 0.001492591 | 0.330111053 | 1.38065049 |
| Ribonuclease P protein subunit p30 | RPP30 | 0.00186099 | 0.00103099 | 0.00234524 | 0.00475214 | 0.006867307 | 0.003939246 | 0.06298165 | 1.352506125 |
| General transcription factor II-I | GTF2I | 0.003715093 | 0.00334408 | 0.00644351 | 0.00239133 | 0.003984276 | 0.002162204 | 0.260470821 | -0.635582076 |
| SRSF protein kinase 2;SRSF protein kinase 2 N-terminal;SRSF protein kinase 2 C-terminal | SRPK2 | 0.000622999 | 0.00071075 | 0.00113691 | 0.00159963 | 0.002568543 | 0.00225976 | 0.082795277 | 1.668753429 |
| T-complex protein 1 subunit beta | CCT2 | 0.006009965 | 0.00477934 | 0.00727856 | 0.00845262 | 0.007998018 | 0.006946938 | 0.480615873 | 0.412285939 |
| DNA-dependent protein kinase catalytic subunit | PRKDC | 0.002807468 | 0.00276416 | 0.00495123 | 0.00321443 | 0.007969355 | 0.00745536 | 0.210893137 | 1.40900844 |
| Brain acid soluble protein 1 | BASP1 | 0.000476766 | 0.00091656 | 0.00269449 | 0.00014927 | 0.000577062 | 0.000635517 | 0.22078812 | -0.667502994 |
| Dermcidin;Survival-promoting peptide;DCD-1 | DCD | 0.006515508 | 0.00746781 | 0.01250434 | 0.01702844 | 0.010008438 | 0.016312141 | 0.227224221 | 1.12718883 |
| SAP domain-containing ribonucleoprotein | SARNP | 0.000495314 | 0.00025244 | 0.00098941 | 0.00177165 | 0.005681279 | 0.003137572 | 0.049614872 | 2.663233262 |
| Enhancer of rudimentary homolog | ERH | 0.022950397 | 0.02553648 | 0.03632843 | 0.01942912 | 0.036652335 | 0.027056762 | 0.931663493 | 0.083429465 |
| RNA-binding protein 10 | RBM10 | 7.35472E-05 | 7.7551E-05 | 9.9991E-05 | 0.00011048 | 0.000193689 | 0.000126987 | 0.217603698 | 0.711472824 |
| Vigilin | HDLBP | 0.003109562 | 0.00326526 | 0.007647 | 0.0057924 | 0.01985106 | 0.011684938 | 0.136123765 | 1.839381627 |
| Transcriptional activator protein Pur-alpha | PURA | 0.002077763 | 0.00306898 | 0.01057357 | 0.00343923 | 0.018525067 | 0.008767988 | 0.354993726 | 1.514484699 |
| Peptidyl-prolyl cis-trans isomerase FKBP3 | FKBP3 | 0.008100437 | 0.00963833 | 0.02273541 | 0.03078428 | 0.074743172 | 0.05570441 | 0.053292624 | 2.530935846 |
| Actin, cytoplasmic 1;Actin, cytoplasmic 1, N-terminally processed;Actin, cytoplasmic 2;Actin, cytoplasmic 2, N-terminally processed | ACTB;ACTG1 | 1.290337028 | 1.04259395 | 1.65092703 | 0.04881256 | 0.069677609 | 0.049048778 | 0.009105929 | -4.717386934 |
| Heterogeneous nuclear ribonucleoprotein U | HNRNPU | 0.008848953 | 0.00910562 | 0.02821719 | 0.02723002 | 0.081827192 | 0.049597897 | 0.093925074 | 2.445449633 |
| Nucleolysin TIAR;Nucleolysin TIA-1 isoform p40 | TIAL1;TIA1 | 0.000541418 | 0.00036811 | 0.00059617 | 0.00026101 | 0.000697607 | 8.2105E-05 | 0.326203542 | -1.052626716 |
| Protein SET;Protein SETSIP | SET;SETSIP | 0.03132404 | 0.02460693 | 0.04506713 | 0.03335699 | 0.040341008 | 0.029780937 | 0.901160455 | 0.0907186 |
| Fatty acid-binding protein, epidermal | FABP5 | 0.000581962 | 0.00448603 | 0.00087012 | 0.00080271 | 0.001644768 | 0.001084568 | 0.833955056 | 0.317840857 |
| 26S protease regulatory subunit 4 | PSMC1 | 0.364891949 | 0.35990788 | 0.43208512 | 0.08211579 | 0.096919409 | 0.094341349 | 0.037208597 | -1.951507184 |
| Exosome component 10 | EXOSC10 | 0.002880137 | 0.00243306 | 0.00467778 | 0.00868989 | 0.017544635 | 0.011518255 | 0.04377718 | 1.999712824 |
| ATP-dependent 6-phosphofructokinase, platelet type | PFKP | 0.003528272 | 0.00299445 | 0.00387871 | 0 | 0.000736127 | 6.13845E-05 | 0.071304389 | -4.052939377 |
| Peptidyl-prolyl cis-trans isomerase FKBP4;Peptidyl-prolyl cis-trans isomerase FKBP4, N-terminally processed | FKBP4 | 0.001395141 | 0.00118884 | 0.00311252 | 0.00176015 | 0.002383306 | 0.001481199 | 0.873355141 | 0.335284167 |
| Gamma-tubulin complex component 3 | TUBGCP3 | 0.000162468 | 0.00015703 | 0.0003776 | 0 | 0 | 0 | NaN | NaN |
| 60S ribosomal protein L6 | RPL6 | 0.114466465 | 0.11779546 | 0.25317381 | 0.20026877 | 0.587401722 | 0.405471716 | 0.127816057 | 1.783317382 |
| DNA topoisomerase 2-beta;DNA topoisomerase 2 | TOP2B | 0.000281536 | 0.00011306 | 0.00030029 | 7.9917E-05 | 0.000269177 | 9.7085E-05 | 0.34958955 | -1.535999095 |
| Eukaryotic translation initiation factor 4 gamma 1 | EIF4G1 | 0.002646232 | 0.00267581 | 0.0079031 | 0.00531269 | 0.013474237 | 0.00885094 | 0.167836088 | 1.725855543 |
| Single-stranded DNA-binding protein, mitochondrial | SSBP1 | 0.008083641 | 0.02104365 | 0.01787287 | 0.01675751 | 0.031908225 | 0.025060851 | 0.236525762 | 0.48766449 |
| 14-3-3 protein eta | YWHAH | 0.025100213 | 0.0184989 | 0.02689835 | 0.00959877 | 0.013566655 | 0.010129491 | 0.127310394 | -1.309137925 |
| Katanin p60 ATPase-containing subunit A-like 2 | KATNAL2 | 0.00372142 | 0.00638187 | 0.00629747 | 0 | 0 | 0 | NaN | NaN |
| Cleavage stimulation factor subunit 1 | CSTF1 | 0.000460307 | 0.00010332 | 0.00145959 | 0.00092451 | 0.002348616 | 0.003383507 | 0.164809588 | 2.351142564 |
| Inosine-5-monophosphate dehydrogenase;Inosine-5-monophosphate dehydrogenase 1;Inosine-5-monophosphate dehydrogenase 2 | IMPDH1;IMPDH2 | 0.010400851 | 0.01070658 | 0.01458398 | 0.00081689 | 0.001039418 | 0.001043034 | 0.013108233 | -3.3646504 |
| Dynamin-1 | DNM1 | 9.42616E-05 | 3.456E-05 | 5.6933E-05 | 0 | 3.07845E-05 | 6.63188E-05 | 0.65697879 | -0.333456637 |
| Elongation factor 1-alpha 2 | EEF1A2 | 0.003553018 | 0.00215168 | 0.00619162 | 0.00300349 | 0.009841417 | 0.005508714 | 0.414446141 | 0.632670566 |
| Exosome complex component RRP45 | EXOSC9 | 0.003237655 | 0.00421826 | 0.00638351 | 0.01257862 | 0.016195872 | 0.008786292 | 0.076373015 | 1.576254636 |
| Proteasome activator complex subunit 1 | PSME1 | 0.050694771 | 0.07871559 | 0.09915974 | 0.05392034 | 0.062598947 | 0.063432967 | 0.595762843 | -0.330510952 |
| Fragile X mental retardation protein 1 | FMR1 | 0.000400252 | 0.0005065 | 0.00162134 | 0.00017633 | 0.003165508 | 0.002371726 | 0.681980388 | 2.227300326 |
| Peroxiredoxin-1 | PRDX1 | 0.013702833 | 0.01511776 | 0.02127957 | 0.00715551 | 0.027300733 | 0.019583293 | 0.919815934 | 0.373378607 |
| KH domain-containing, RNA-binding, signal transduction-associated protein 1 | KHDRBS1 | 0.001531519 | 0.00130179 | 0.00154471 | 0.00098887 | 0.000830447 | 0.001330484 | 0.366426223 | -0.631106111 |
| ATP-dependent RNA helicase A | DHX9 | 0.009821968 | 0.01060295 | 0.02046214 | 0.02889534 | 0.070151753 | 0.046441437 | 0.047591685 | 2.130946627 |
| Galectin-3-binding protein | LGALS3BP | 0.000972791 | 0.00057847 | 0.00062629 | 0.00035967 | 0 | 0.000135473 | 0.084699142 | -1.504496755 |
| Desmocollin-1 | DSC1 | 9.85668E-05 | 7.9302E-05 | 0.00027607 | 7.3466E-05 | 0.000102226 | 0 | 0.478523657 | -0.185722218 |
| Protein VAC14 homolog | VAC14 | 0.000315194 | 5.2836E-05 | 0.00042977 | 6.1266E-05 | 0.000192149 | 0.000114988 | 0.491217 | -1.454756761 |
| tRNA (cytosine(34)-C(5))-methyltransferase | NSUN2 | 0.000994681 | 0.0012863 | 0.00360115 | 0.00638725 | 0.011563333 | 0.007142635 | 0.047280073 | 2.473232606 |
| Histone-binding protein RBBP4 | RBBP4 | 0.005731161 | 0.00629551 | 0.00724925 | 0.00380595 | 0.004504895 | 0.003868562 | 0.256726769 | -0.702525631 |
| Nuclear cap-binding protein subunit 1 | NCBP1 | 0.004727354 | 0.00384099 | 0.00509599 | 0.00218062 | 0.00320127 | 0.002877227 | 0.225261716 | -0.716353855 |
| Heat shock 70 kDa protein 14 | HSPA14 | 0.001577259 | 0.00059992 | 0.00139885 | 0.00224544 | 0.00400142 | 0.003098432 | 0.075330015 | 1.147298069 |
| AP-1 complex subunit beta-1 | AP1B1 | 0.001690348 | 0.00113514 | 0.00226897 | 0.00026978 | 0.000479849 | 0.000784558 | 0.055509307 | -1.8166679 |
| Cleavage and polyadenylation specificity factor subunit 1 | CPSF1 | 4.30719E-05 | 5.1664E-05 | 4.4068E-05 | 8.007E-06 | 2.34393E-05 | 2.5055E-05 | 0.08749103 | -0.910801937 |
| Transducin beta-like protein 3 | TBL3 | 0.000284274 | 0.00034144 | 0.00041052 | 0.00129173 | 0.003219351 | 0.002733132 | 0.020649845 | 3.000842701 |
| General transcription factor 3C polypeptide 1 | GTF3C1 | 2.6852E-05 | 0.00011484 | 0.00027096 | 4.5351E-05 | 0.00017783 | 0 | 0.965045171 | -0.354827759 |
| | UBA52 | 0 | 0.00026452 | 0.00014979 | 0 | 0 | 0 | NaN | NaN |
| Splicing factor 3A subunit 3 | SF3A3 | 0.000907849 | 0.00112907 | 0.00127473 | 0.00085068 | 0.001371934 | 0.001046675 | 0.953082147 | -0.10932011 |
| Aminoacyl tRNA synthase complex-interacting multifunctional protein 1;Endothelial monocyte-activating polypeptide 2 | AIMP1 | 0.001480797 | 0.00195834 | 0.00421783 | 0.00232618 | 0.00769103 | 0.004288774 | 0.258030825 | 1.130935505 |
| Interleukin enhancer-binding factor 3 | ILF3 | 0.018796887 | 0.02007328 | 0.04906841 | 0.05486319 | 0.102580999 | 0.064317009 | 0.086199044 | 1.679924302 |
| Actin-related protein 3 | ACTR3 | 0.000952973 | 0.00044428 | 0.00046519 | 0 | 0 | 0 | NaN | NaN |
| Cell division cycle protein 16 homolog | CDC16 | 0.000314287 | 0.00058812 | 0.00111362 | 0.00030195 | 0.000864293 | 0.000379204 | 0.651099726 | -0.633133917 |
| Protein HEXIM1 | HEXIM1 | 0.000697906 | 0.00081971 | 0.00145182 | 0 | 0 | 0 | NaN | NaN |
| Acetyl-CoA carboxylase 1;Biotin carboxylase | ACACA | 6.73609E-05 | 8.2203E-05 | 9.7023E-05 | 1.6491E-05 | 3.21667E-05 | 5.91432E-05 | 0.102456878 | -1.353631885 |
| Chromatin assembly factor 1 subunit B | CHAF1B | 0.000201719 | 0.00028617 | 0.00037359 | 0.00042009 | 0.000913208 | 0.000428585 | 0.148380286 | 0.582698837 |
| Unconventional myosin-VI | MYO6 | 0.00031441 | 0.00066975 | 0.00062593 | 0 | 0 | 0 | NaN | NaN |
| Heterogeneous nuclear ribonucleoprotein A0 | HNRNPA0 | 0.024921621 | 0.02160997 | 0.04211983 | 0.02793205 | 0.084836796 | 0.06092104 | 0.198555758 | 1.289542661 |
| Aminoacyl tRNA synthase complex-interacting multifunctional protein 2 | AIMP2 | 0.000913615 | 0.00063953 | 0.001019 | 0.0021331 | 0.004257377 | 0.002942849 | 0.047088397 | 1.687555276 |
| Coronin-1C;Coronin | CORO1C | 0.001098757 | 0.00120004 | 0.00134944 | 0 | 8.23E-05 | 5.75484E-05 | 0.012617039 | -4.124106965 |
| Serine/arginine-rich splicing factor 9 | SRSF9 | 0.000620927 | 0.0008591 | 0.00145073 | 0.00030302 | 0.000786328 | 0.00047861 | 0.176888084 | -0.84397889 |
| Transcription intermediary factor 1-beta | TRIM28 | 0.003924253 | 0.00488213 | 0.00587451 | 0.00202892 | 0.004435381 | 0.004325187 | 0.354837202 | -0.174748498 |
| Ras GTPase-activating protein-binding protein 1 | G3BP1 | 0.005925428 | 0.00720004 | 0.01173075 | 0.00876945 | 0.022074443 | 0.015544153 | 0.177450562 | 1.110294779 |
| Polyadenylate-binding protein 4;Polyadenylate-binding protein | PABPC4 | 0.002547755 | 0.00400764 | 0.00698292 | 0.0044109 | 0.011688566 | 0.011223639 | 0.193717078 | 1.485714967 |
| Serine-protein kinase ATM | ATM | 0 | 0 | 0 | 0 | 0 | 0 | NaN | NaN |
| Eukaryotic translation initiation factor 3 subunit I | EIF3I | 0.013423469 | 0.01501727 | 0.0301644 | 0.02705478 | 0.05225218 | 0.045469769 | 0.116954616 | 1.598284914 |
| Probable methyltransferase TARBP1 | TARBP1 | 9.34218E-05 | 8.5084E-05 | 0.00022318 | 3.1159E-05 | 6.43335E-05 | 6.16357E-05 | 0.114976284 | -0.599993026 |
| Vacuolar fusion protein CCZ1 homolog;Vacuolar fusion protein CCZ1 homolog B | CCZ1B;CCZ1 | 0.000536771 | 0 | 0.00032816 | 0 | 0 | 0 | NaN | NaN |
| DNA repair protein XRCC4 | XRCC4 | 0.00013541 | 0.00012709 | 0.00014656 | 3.6443E-05 | 0.00022195 | 0 | 0.582508751 | -0.590370012 |
| Splicing factor 3B subunit 2 | SF3B2 | 0.001319897 | 0.00134942 | 0.0019484 | 0.00211536 | 0.002553006 | 0.001896583 | 0.346490828 | 0.648563463 |
| 28 kDa heat- and acid-stable phosphoprotein | PDAP1 | 0.007386071 | 0.01207055 | 0.01231985 | 0.02183841 | 0.065758562 | 0.036790965 | 0.048373044 | 1.607859625 |

| | | | | | | | | | |
|---|---|---|---|---|---|---|---|---|---|
| Peptidyl-prolyl cis-trans isomerase FKBP5;Peptidyl-prolyl cis-trans isomerase FKBP5, N-terminally processed | FKBP5 | 0.000713526 | 0.00075634 | 0.00121993 | 0.00196248 | 0.003055678 | 0.002279428 | 0.075949983 | 1.59156883 |
| Serine/threonine-protein kinase ATR | ATR | 0 | 1.6136E-05 | 0 | 0 | 2.10927E-05 | 0 | NaN | 0.38643397 |
| KRR1 small subunit processome component homolog | KRR1 | 0.000292627 | 0.00057578 | 0.0009863 | 0.00296253 | 0.006730957 | 0.005561484 | 0.024378661 | 3.271887831 |
| Cullin-1 | CUL1 | 0.00827063 | 0.0076309 | 0.00935337 | 0.00636865 | 0.00705817 | 0.004294421 | 0.332123883 | -0.37700964 |
| Histone deacetylase 2;Histone deacetylase 1 | HDAC2;HDAC1 | 0.001193259 | 0.00118905 | 0.00096283 | 0 | 0.000282544 | 0 | 0.019825457 | -2.073263137 |
| Cullin-3 | CUL3 | 0.000288825 | 0.00010704 | 0.00015845 | 0.00010884 | 0 | 0.000122876 | 0.367978724 | -0.454329082 |
| Procollagen-lysine,2-oxoglutarate 5-dioxygenase 1 | PLOD1 | 0.000357026 | 0.00017745 | 0.00015417 | 0 | 0 | 0 | NaN | NaN |
| Four and a half LIM domains protein 1 | FHL1 | 0.001590415 | 0.00202753 | 0.00341876 | 0.00125087 | 0.002302139 | 0.002074189 | 0.596555278 | 0.032823386 |
| 26S proteasome non-ATPase regulatory subunit 12 | PSMD12 | 0.241568693 | 0.26513137 | 0.25852589 | 0.05606193 | 0.066153681 | 0.052873138 | 0.035332603 | -2.205215595 |
| Nucleolar GTP-binding protein 2 | GNL2 | 7.50476E-05 | 0.00014233 | 6.4291E-05 | 0.00095645 | 0.001753124 | 0.000725246 | 0.013563836 | 3.671808694 |
| Spliceosome RNA helicase DDX39B;ATP-dependent RNA helicase DDX39A | DDX39B;DDX39A | 0.001315866 | 0.00143641 | 0.00283314 | 0.00264678 | 0.006672694 | 0.005655146 | 0.093671046 | 1.977099682 |
| Tubulin beta-2A chain | TUBB2A | 0 | 8.0594E-05 | 0.00011194 | 0.00012029 | 0.000456048 | 0.000429131 | 0.151569355 | 2.175706933 |
| Bystin | BYSL | 0.006815026 | 0.00667256 | 0.00928943 | 0.0079255 | 0.011341932 | 0.007726025 | 0.667156175 | 0.217782065 |
| Cytoskeleton-associated protein 5 | CKAP5 | 0.00047212 | 0.00051138 | 0.00057136 | 0.00136516 | 0.003548439 | 0.002317593 | 0.035654915 | 2.18016241 |
| Cold-inducible RNA-binding protein | CIRBP | 0.001375042 | 0.00134743 | 0.00241255 | 0.00215933 | 0.004610573 | 0.002738584 | 0.175233782 | 0.993954447 |
| Rho GTPase-activating protein 17 | ARHGAP17 | 0.000127365 | 8.478E-05 | 4.8318E-05 | 0 | 0 | 0 | NaN | NaN |
| Actin-related protein 2/3 complex subunit 2 | ARPC2 | 0.000567014 | 0.00025514 | 0.00041255 | 0 | 0 | 0 | NaN | NaN |
| Scaffold attachment factor B1;Scaffold attachment factor B2 | SAFB;SAFB2 | 0.000189839 | 0.00136292 | 0.00083137 | 0.00040261 | 0.000461539 | 0.000275845 | 0.478152322 | -1.046105722 |
| Eukaryotic translation initiation factor 3 subunit A | EIF3A | 0.011148808 | 0.00917638 | 0.01914969 | 0.02084718 | 0.053282169 | 0.031601597 | 0.085499251 | 1.503107998 |
| Cytoplasmic dynein 1 heavy chain 1 | DYNC1H1 | 0.000112843 | 9.4312E-05 | 0.00021827 | 4.9421E-05 | 9.1144E-05 | 6.24847E-05 | 0.140358543 | -0.852742047 |
| Translation initiation factor eIF-2B subunit alpha | EIF2B1 | 0.001460811 | 0.00112813 | 0.0028925 | 0.00312476 | 0.006848957 | 0.006162983 | 0.072571296 | 2.076859278 |
| Eukaryotic initiation factor 4A-II;Eukaryotic initiation factor 4A-II, N-terminally processed | EIF4A2 | 0.00067792 | 0.00157239 | 0.00271468 | 0.0005917 | 0.001685083 | 0.001101529 | 0.567762145 | -0.513447946 |
| Transcription elongation factor B polypeptide 3 | TCEB3 | 9.84044E-05 | 2.5824E-05 | 0.00027115 | 0 | 0.000319953 | 0.000307662 | 0.23034251 | 1.672808399 |
| Src substrate cortactin | CTTN | 0.009171985 | 0.00447409 | 0.01068317 | 0.00071912 | 0.001703969 | 0.000373342 | 0.03194289 | -3.67292755 |
| E3 ubiquitin/ISG15 ligase TRIM25 | TRIM25 | 0.000811443 | 0.00130859 | 0.0029831 | 0.00221072 | 0.00614377 | 0.004534709 | 0.139104153 | 1.792992413 |
| Filamin-C | FLNC | 0.000529924 | 0.00034975 | 0.00032631 | 0 | 3.04376E-05 | 9.72856E-05 | 0.037128635 | -2.68422821 |
| Caprin-1 | CAPRIN1 | 0.006675064 | 0.00746572 | 0.01252169 | 0.00701758 | 0.020968109 | 0.014507838 | 0.32383072 | 0.95847962 |
| DNA replication licensing factor MCM6 | MCM6 | 0.000603908 | 0.00219261 | 0.00146443 | 0.00160522 | 0.003389722 | 0.000772758 | 0.686008441 | 0.132440237 |
| Separin | ESPL1 | 0.000216325 | 0.00023617 | 0.00045702 | 0.00027076 | 0.00088421 | 0.00048192 | 0.265552177 | 1.028986286 |
| Mediator of DNA damage checkpoint protein 1 | MDC1 | 1.90947E-05 | 3.9329E-05 | 5.7919E-05 | 0 | 0 | 8.48992E-06 | 0.033741316 | -2.211770639 |
| Coactosin-like protein | COTL1 | 0.013139626 | 0.00889124 | 0.01321856 | 0 | 0 | 0 | NaN | NaN |
| Structural maintenance of chromosomes protein 1A;Structural maintenance of chromosomes protein | SMC1A | 0.080002239 | 0.09245159 | 0.09643803 | 0.1234317 | 0.121594942 | 0.118574628 | 0.423790729 | 0.395313244 |
| Ribosomal RNA processing protein 1 homolog B | RRP1B | 0.000766096 | 0.00080072 | 0.0018925 | 0.00100104 | 0.004494046 | 0.00359595 | 0.199867161 | 2.166999611 |
| Protein RRP5 homolog | PDCD11 | 1.85102E-05 | 5.5834E-06 | 0.000101 | 9.06E-05 | 0.000384056 | 0.000210067 | 0.112555731 | 3.504457154 |
| Ribosome biogenesis protein BMS1 homolog | BMS1 | 0.000199429 | 0.00026849 | 7.9539E-05 | 0.00071298 | 0.002075247 | 0.00211917 | 0.035024404 | 3.379335939 |
| Ubiquitin carboxyl-terminal hydrolase 10 | USP10 | 0.00025426 | 0.00029348 | 0.00087451 | 0.00033315 | 0.00212909 | 0.001289027 | 0.261194857 | 2.134958219 |
| Neutral alpha-glucosidase AB | GANAB | 0.000103891 | 0.00014903 | 0.00036654 | 0 | 0.000213083 | 0.000104967 | 0.772840129 | 0.00505353 |
| Centrosomal protein of 85 kDa | CEP85 | 0.000108135 | 0.00024364 | 0.00021458 | 0 | 0 | 0 | NaN | NaN |
| Ras-related protein Rab-39A;Ras-related protein Rab-6A;Ras-related protein Rab-6B | RAB39A;RAB6B;RAB6A | 0.001708991 | 0.00273453 | 0.0031149 | 0.00034935 | 0.001667537 | 0.001702736 | 0.194381274 | -0.713576227 |
| Importin subunit beta-1 | KPNB1 | 0.004725955 | 0.00539495 | 0.00663412 | 0.00452938 | 0.006530317 | 0.003687859 | 0.703899247 | -0.252298032 |
| Carboxypeptidase D | CPD | 3.40057E-05 | 2.5458E-05 | 6.8616E-05 | 0 | 0 | 0 | NaN | NaN |
| Nuclear mitotic apparatus protein 1 | NUMA1 | 0 | 1.3627E-05 | 5.6359E-05 | 1.2659E-05 | 0 | 0 | 0.320968448 | -1.130370937 |
| Proteasome activator complex subunit 4 | PSME4 | 0.043104915 | 0.04983042 | 0.05953968 | 0.04224372 | 0.042474652 | 0.043325869 | 0.667013731 | -0.230424607 |
| N-alpha-acetyltransferase 25, NatB auxiliary subunit | NAA25 | 0.000568021 | 0.00046979 | 0.0013794 | 0.00053169 | 0.001473594 | 0.000956849 | 0.631501495 | 0.752347004 |
| DnaJ homolog subfamily C member 10 | DNAJC10 | 0.000778748 | 0.00074434 | 0.00071036 | 3.4067E-05 | 4.86546E-05 | 0 | 0.01246023 | -4.192428767 |
| Glioma tumor suppressor candidate region gene 2 protein | GLTSCR2 | 0 | 0 | 0 | 0.0001507 | 0.000517365 | 0.000405219 | NaN | NaN |
| Condensin complex subunit 1 | NCAPD2 | 0.0003884 | 9.9119E-05 | 0.000387 | 0 | 1.52878E-05 | 0 | 0.016779645 | -4.661892035 |
| Exosome complex component RRP42 | EXOSC7 | 4.65334E-05 | 0.00019492 | 0.00037002 | 0.00234855 | 0.001523285 | 0.003018596 | 0.036435315 | 3.590791614 |
| 116 kDa U5 small nuclear ribonucleoprotein component | EFTUD2 | 0.001571493 | 0.00194442 | 0.00328304 | 0.00075825 | 0.00189818 | 0.001163003 | 0.177884808 | -0.741481351 |
| Lysine--tRNA ligase | KARS | 0.002656757 | 0.00305538 | 0.00591488 | 0.00525485 | 0.009955398 | 0.0058524 | 0.16882286 | 0.937678919 |
| Thioredoxin reductase 1, cytoplasmic | TXNRD1 | 0 | 0.00011827 | 0.00019478 | 0 | 0 | 0 | NaN | NaN |
| Eukaryotic translation initiation factor 4H | EIF4H | 0.001875434 | 0.00201623 | 0.006764 | 0.00017794 | 0.001061866 | 0.000201928 | 0.068766529 | -3.319743766 |
| Protein disulfide-isomerase A6 | PDIA6 | 0.001331878 | 0.00151753 | 0.00210037 | 0.00024031 | 0.000581709 | 0.000435381 | 0.0388109 | -1.801378485 |
| Plectin | PLEC | 0.000133143 | 0.00015685 | 0.00026482 | 0.00011608 | 0.000140904 | 9.31146E-05 | 0.277928176 | -0.43430016 |
| Non-POU domain-containing octamer-binding protein | NONO | 0.001063599 | 0.00066168 | 0.00372819 | 0.00261915 | 0.010390565 | 0.003380197 | 0.169112704 | 1.668153004 |
| Periodic tryptophan protein 2 homolog | PWP2 | 0.000126873 | 0.00015296 | 0.00038593 | 0.00131936 | 0.002100561 | 0.001130854 | 0.023487943 | 3.108636953 |
| Poly(rC)-binding protein 1 | PCBP1 | 0.02411656 | 0.01461216 | 0.04738424 | 0.00676203 | 0.028119098 | 0.008626229 | 0.240862047 | -1.483222214 |
| Poly(rC)-binding protein 2;Poly(rC)-binding protein 3 | PCBP2;PCBP3 | 0.007117904 | 0.0055592 | 0.01912412 | 0.00596624 | 0.01980686 | 0.012949469 | 0.683522358 | 0.86336856 |
| Myosin-9 | MYH9 | 0.00208868 | 0.00180624 | 0.00238926 | 0.00011789 | 0.000163378 | 0.000130242 | 0.01136379 | -4.003324815 |
| Splicing factor 3B subunit 3 | SF3B3 | 0.005355951 | 0.00554423 | 0.00752836 | 0.00295189 | 0.003180107 | 0.003296466 | 0.155411882 | -0.801911019 |
| Pumilio domain-containing protein KIAA0020 | KIAA0020 | 4.20194E-05 | 0.00010089 | 0.00014326 | 0.00020078 | 0.002478268 | 0.00137675 | 0.084037373 | 3.770397219 |
| Splicing factor 3B subunit 4 | SF3B4 | 0.000278541 | 0.00050065 | 0.00080443 | 0.00064514 | 0.00119847 | 0.000566274 | 0.294727449 | 0.36581948 |
| Protein transport protein Sec23B | SEC23B | 0.00023962 | 0.00017891 | 0.00015782 | 6.146E-05 | 0 | 0 | 0.030719738 | -1.541490164 |
| Splicing factor 3A subunit 1 | SF3A1 | 0.001085153 | 0.00103672 | 0.00136524 | 0.00138666 | 0.001743882 | 0.00165976 | 0.400519022 | 0.613076176 |
| Helicase SKI2W | SKIV2L | 0.000560184 | 0.00049395 | 0.00079297 | 0.000475 | 0.002023546 | 0.001498919 | 0.249471515 | 1.419949741 |
| RISC-loading complex subunit TARBP2 | TARBP2 | 0.000440818 | 0.0004776 | 0.00117828 | 0.00033082 | 0.001113593 | 0.000848019 | 0.885979964 | 0.828296773 |
| Splicing factor 1 | SF1 | 0.000136855 | 0.00026865 | 0.00024093 | 0.00012502 | 0.000444395 | 0.000228897 | 0.796157372 | -0.073891539 |
| Microtubule-associated protein RP/EB family member 1 | MAPRE1 | 0.00222629 | 0.00202565 | 0.00300859 | 0.00028553 | 0.001303827 | 0.000253646 | 0.06812094 | -2.96290203 |
| ELAV-like protein 1 | ELAVL1 | 0.003526985 | 0.00567225 | 0.00955795 | 0.00225598 | 0.018459437 | 0.00568046 | 0.925137592 | 0.002086643 |
| Protein flightless-1 homolog | FLII | 1.15665E-05 | 2.8825E-05 | 2.88E-05 | 0 | 0 | 0 | NaN | NaN |
| Ubiquitin-conjugating enzyme E2 variant 2;Ubiquitin-conjugating enzyme E2 variant 1 | UBE2V2;TMEM189-UBE2V1;U | 0.00162647 | 0.0013512 | 0.00306813 | 0.00041529 | 0.001358541 | 0.000132534 | 0.132938699 | -1.969560122 |
| Proteasomal ubiquitin receptor ADRM1 | ADRM1 | 0.237834509 | 0.22854601 | 0.31228423 | 0.07822609 | 0.063276677 | 0.087066498 | 0.048583915 | -1.604236244 |
| Coiled-coil domain-containing protein 6 | CCDC6 | 0.000281962 | 0.00010306 | 0.00018524 | 0 | 0 | 7.60238E-05 | 0.078672838 | -1.284880406 |
| Lysosome-associated membrane glycoprotein 1 | LAMP1 | 0.000842123 | 0.00053611 | 0.00104156 | 0 | 0 | 0 | NaN | NaN |
| DNA damage-binding protein 1 | DDB1 | 0.004525529 | 0.00500325 | 0.00744342 | 0.00305628 | 0.004567177 | 0.003548437 | 0.295985636 | -0.495680408 |
| Serine/threonine-protein phosphatase 2A 56 kDa regulatory subunit epsilon isoform;Serine/threonine-protein phosphatase 2A 56 kDa regulatory subunit beta isoform | PPP2R5E;PPP2R5B | 0.000368755 | 0.00017734 | 0.00034338 | 0 | 9.69154E-05 | 0 | 0.034214051 | -1.825020782 |
| Hsp90 co-chaperone Cdc37;Hsp90 co-chaperone Cdc37, N-terminally processed | CDC37 | 0.002548651 | 0.00195352 | 0.00450242 | 0.00192926 | 0.007622989 | 0.000263538 | 0.600678273 | -0.401686995 |
| Drebrin | DBN1 | 0.002293696 | 0.00212488 | 0.00193324 | 0.00165715 | 0.002666051 | 0.002514653 | 0.886456592 | 0.242976608 |
| Fascin | FSCN1 | 0.0005152 | 0.00044361 | 0.00097123 | 0 | 8.61762E-05 | 0.000136947 | 0.032311261 | -2.245643068 |
| Protein enabled homolog;Ena/VASP-like protein | ENAH;EVL | 0.000468928 | 0.00043943 | 0.00083651 | 0 | 0 | 0 | NaN | NaN |
| Heterogeneous nuclear ribonucleoprotein U-like protein 2 | HNRNPUL2;HNRNPUL2-BSCL | 0.004199698 | 0.00374301 | 0.00636204 | 0.00993399 | 0.016414192 | 0.012629734 | 0.076188364 | 1.588466762 |
| Actin-related protein 2/3 complex subunit 3 | ARPC3 | 0.003364349 | 0.00218382 | 0.0019916 | 0 | 0 | 0 | NaN | NaN |
| Sister chromatid cohesion protein PDS5 homolog A | PDS5A | 3.61936E-05 | 8.4639E-05 | 0.00034757 | 0.00124776 | 0.00255381 | 0.001525207 | 0.03926199 | 4.171531847 |
| Pre-rRNA-processing protein TSR1 homolog | TSR1 | 0.003544508 | 0.0035279 | 0.00788803 | 0.00410944 | 0.005767938 | 0.002317788 | 0.643577565 | 0.213358178 |
| WD40 repeat-containing protein SMU1;WD40 repeat-containing protein SMU1, N-terminally processed | SMU1 | 0.000254831 | 0.00045914 | 0.00071956 | 0.00019165 | 0.000539103 | 0.000203602 | 0.351711579 | -1.17319496 |
| Prolyl 3-hydroxylase 1 | LEPRE1 | 0.000353079 | 0.00034994 | 0.00015602 | 0.00034184 | 0.000301751 | 0.000246402 | 0.810704987 | -0.213757275 |
| MAP7 domain-containing protein 1 | MAP7D1 | 9.55268E-05 | 8.5115E-05 | 0.00030334 | 0.00066889 | 0.000824286 | 0.000805978 | 0.035708825 | 3.076763017 |
| La-related protein 7 | LARP7 | 0.001034599 | 0.00076788 | 0.00208138 | 0.00194313 | 0.004679016 | 0.004313893 | 0.094333624 | 2.059918691 |
| Endoplasmic reticulum resident protein 44 | ERP44 | 5.92991E-05 | 0.00011401 | 0.00013189 | 0 | 0 | 0 | NaN | NaN |
| Protein FAM98B | FAM98B | 0.005386631 | 0.00297435 | 0.00396803 | 0.00340074 | 0.014470741 | 0.012907799 | 0.207999314 | 1.701746725 |
| Programmed cell death protein 4 | PDCD4 | 0.002776677 | 0.00350288 | 0.00670874 | 0.01175509 | 0.026888201 | 0.017769059 | 0.036801795 | 2.342754223 |
| Peflin | PEF1 | 0.000909809 | 0.0014026 | 0.00163595 | 0 | 0 | 0 | NaN | NaN |
| Beta-actin-like protein 2 | ACTBL2 | 0.00174163 | 5.7894E-05 | 0.00282007 | 0 | 0 | 0.000174154 | 0.326481841 | -3.322002649 |
| Nucleolar MIF4G domain-containing protein 1 | NOM1 | 1.77914E-05 | 0 | 3.99E-05 | 0.00021121 | 0.000701317 | 0.000294733 | 0.029223004 | 3.467561343 |
| DnaJ homolog subfamily C member 21 | DNAJC21 | 0.000479655 | 0.00043029 | 0.00089971 | 0.00038061 | 0.000676632 | 0.000583819 | 0.856210501 | 0.283524067 |
| Putative helicase MOV-10 | MOV10 | 0.001167506 | 0.00155365 | 0.00293744 | 0.00223383 | 0.006735645 | 0.004225684 | 0.1463108 | 1.443525207 |
| PHD finger protein 6 | PHF6 | 0.000259719 | 0.00024677 | 0.00025825 | 8.6413E-05 | 0.001014532 | 0.000659098 | 0.621136479 | 1.351713471 |
| RRP12-like protein | RRP12 | 0.001631452 | 0.0011276 | 0.00323765 | 0.00210052 | 0.004818312 | 0.002841788 | 0.252829709 | 0.800642391 |
| Cytochrome c oxidase assembly factor 6 homolog | COA6 | 0.000543416 | 0 | 0.00119025 | 0 | 0 | 0.000388005 | 0.159587438 | -1.051547216 |
| Syntaxin-binding protein 1 | STXBP1 | 6.71369E-05 | 2.6914E-05 | 4.1742E-05 | 0 | 0 | 0 | NaN | NaN |
| PCI domain-containing protein 2 | PCID2 | 0 | 5.8679E-05 | 0.00010377 | 0.0001311 | 0.000381444 | 0.000176661 | 0.13157552 | 1.178818105 |
| Crooked neck-like protein 1 | CRNKL1 | 6.86093E-05 | 0.00011224 | 0.0001765 | 0.00023425 | 0.000484014 | 0.000344952 | 0.062822938 | 1.619847273 |
| Proteasome inhibitor PI31 subunit | PSMF1 | 0.00519757 | 0.00783482 | 0.00727738 | 0.00225458 | 0.00578977 | 0.005033006 | 0.229778111 | -0.531998233 |
| Exosome complex component MTR3 | EXOSC6 | 0.01227914 | 0.00660913 | 0.01241392 | 0.02659249 | 0.044719465 | 0.028978678 | 0.055693776 | 1.238782259 |
| | C8G | 0.002172601 | 0 | 0 | 0.00056111 | 0.000680261 | 0.000233473 | 0.027478782 | -1.953057433 |
| Heterochromatin protein 1-binding protein 3 | HP1BP3 | 0.000990707 | 0.00071851 | 0.00091199 | 0.00115433 | 0.002499029 | 0.001972739 | 0.133878364 | 1.113107021 |
| Centrosomal protein of 170 kDa | CEP170 | 8.76442E-05 | 0.0001082 | 0.00022467 | 0 | 7.80032E-05 | 2.23153E-05 | 0.155779435 | -1.374792003 |

| | | | | | | | | | |
|---|---|---|---|---|---|---|---|---|---|
| Mitotic-spindle organizing protein 2A;Mitotic-spindle organizing protein 2B | MZT2B;MZT2A | 0.000266588 | 0.0003078 | 0.00052567 | 0 | 0 | 0 | NaN | NaN |
| Ras-related protein Rab-21 | RAB21 | 0.000214752 | 0 | 0.00032634 | 0 | 0 | 0 | NaN | NaN |
| Putative methyltransferase C9orf114 | C9orf114 | 0.000111684 | 0.00027898 | 0.00026128 | 0.00020571 | 0.000271788 | 0.000324389 | 0.484652901 | 0.056912672 |
| HEAT repeat-containing protein 1;HEAT repeat-containing protein 1, N-terminally processed | HEATR1 | 8.84335E-06 | 2.4627E-05 | 4.2057E-05 | 5.7309E-05 | 0.0001039 | 0.000107382 | 0.081246294 | 2.076894573 |
| Cancer-related nucleoside-triphosphatase | NTPCR | 0.000158224 | 0.00011617 | 0.00057824 | 0 | 0.000521892 | 0.000142681 | 0.807094357 | 0.78630381 |
| Probable ATP-dependent RNA helicase DDX27 | DDX27 | 0 | 0 | 0 | 0.00140407 | 0.003258997 | 0.00294168 | NaN | NaN |
| Rootletin | CROCC | 4.16527E-05 | 2.1735E-05 | 0.00427893 | 3.508E-05 | 3.89052E-05 | 5.44017E-05 | 0.473000556 | -0.09844702 |
| Protein angel homolog 2 | ANGEL2 | 0.000108868 | 0 | 8.5857E-05 | 9.9913E-05 | 0.000246153 | 0 | 0.347129355 | 0.6978536 |
| E3 ubiquitin-protein ligase BRE1A | RNF20 | 0.000129622 | 0.00024746 | 0.00026662 | 2.5915E-05 | 4.96109E-05 | 6.04829E-05 | 0.031321891 | -2.318477969 |
| Hypoxia-inducible factor 1-alpha inhibitor | HIF1AN | 0 | 0.00063766 | 0.00091725 | 0 | 0 | 0 | NaN | NaN |
| Focadhesin | FOCAD | 0.000134884 | 2.5667E-05 | 0.0002873 | 9.0742E-05 | 0.000222352 | 0.000108038 | 0.758767427 | -0.320180387 |
| Clathrin interactor 1 | CLINT1 | 0.000189167 | 0.0002865 | 0 | 0 | 0 | 0 | NaN | NaN |
| Rho guanine nucleotide exchange factor 2 | ARHGEF2 | 0.000298343 | 0.00031563 | 0.00033819 | 0.00150707 | 0.001417072 | 0.003453413 | 0.022280892 | 2.2554438 |
| Something about silencing protein 10 | UTP3 | 0 | 0 | 0 | 0.00042204 | 0.001088868 | 0.001070061 | NaN | NaN |
| Terminal uridylyltransferase 7 | ZCCHC6 | 0 | 0.00019263 | 0.00048868 | 5.4017E-06 | 2.99889E-05 | 0 | 0.061364102 | -4.591340628 |
| Centrosomal protein of 135 kDa | CEP135 | 2.0552E-05 | 0 | 0 | 0 | 0 | 0 | NaN | NaN |
| Microtubule-associated protein 1S;MAP1S heavy chain;MAP1S light chain | MAP1S | 0.000110268 | 2.4103E-05 | 0.00033545 | 0 | 0.000415304 | 0.00019556 | 0.337604898 | 1.36987536 |
| Twinfilin-1 | TWF1 | 0.000461421 | 0.00032232 | 0.00061799 | 0 | 0 | 0 | NaN | NaN |
| CWF19-like protein 1 | CWF19L1 | 0.000965681 | 0.00124055 | 0.00193031 | 0.00109735 | 0.002089712 | 0.001718898 | 0.64645338 | 0.470500565 |
| AP-2 complex subunit alpha-2 | AP2A2 | 5.47492E-05 | 0 | 8.5492E-05 | 0 | 0 | 0 | NaN | NaN |
| Condensin-2 complex subunit H2 | NCAPH2 | 0.000565894 | 0.00084571 | 0.00154717 | 0.00195764 | 0.003929361 | 0.002224321 | 0.07716161 | 1.395124943 |
| Protein SDE2 homolog | SDE2 | 0.000518128 | 0.00127248 | 0.00173395 | 0.00188647 | 0.003081395 | 0.001615169 | 0.179720423 | 0.568046176 |
| N-alpha-acetyltransferase 16, NatA auxiliary subunit | NAA16 | 4.47766E-05 | 1.5639E-05 | 0.00033998 | 5.3574E-05 | 0.000361474 | 0.000452828 | 0.335373653 | 3.013074991 |
| Zinc finger CCHC domain-containing protein 8 | ZCCHC8 | 0.000501338 | 0.00055422 | 0.00050675 | 0.00062896 | 0.001176504 | 0.000985746 | 0.201837047 | 0.959944614 |
| Putative ATP-dependent RNA helicase DHX57 | DHX57 | 0.000790281 | 0.00084099 | 0.00129245 | 0.00048862 | 0.002016046 | 0.000817019 | 0.963856738 | -0.041722448 |
| Parafibromin | CDC73 | 0.001023794 | 0.00080995 | 0.00128569 | 7.3239E-05 | 0.000165227 | 0.000278064 | 0.029386424 | -2.631404092 |
| Enhancer of mRNA-decapping protein 4 | EDC4 | 0.004989867 | 0.00077949 | 0.00215079 | 0.00744815 | 0.001649054 | 0.001088151 | 0.859611142 | -0.383228027 |
| Pre-mRNA-processing-splicing factor 8 | PRPF8 | 0.000822864 | 0.00090728 | 0.00166499 | 0.00079831 | 0.001348897 | 0.001186856 | 0.979114331 | 0.387533094 |
| La-related protein 4 | LARP4 | 0.001551898 | 0.00147064 | 0.00137656 | 0.00063722 | 0.004828759 | 0.003157239 | 0.584564442 | 1.102222288 |
| Tetratricopeptide repeat protein 37 | TTC37 | 0.001696227 | 0.00092168 | 0.0018893 | 0.002554 | 0.004411674 | 0.002725927 | 0.118054448 | 0.684417674 |
| Coiled-coil domain-containing protein 137 | CCDC137 | 7.02273E-05 | 0 | 0.00031424 | 0.00051418 | 0.001091172 | 0.000508266 | 0.114413214 | 1.791276407 |
| La-related protein 1 | LARP1 | 0.005247061 | 0.00547639 | 0.00955795 | 0.01273988 | 0.028232946 | 0.02077305 | 0.064426904 | 1.923414641 |
| Biogenesis of lysosome-related organelles complex 1 subunit 3 | BLOC1S3 | 0 | 0 | 0.00014214 | 0 | 0 | 0 | NaN | NaN |
| WD repeat-containing protein 82 | WDR82 | 0.00294855 | 0.00288538 | 0.00350352 | 0.00318572 | 0.003455887 | 0.003973907 | 0.739202117 | 0.229050386 |
| Profilin;Profilin-2 | PFN2 | 0.004239503 | 0.0032774 | 0 | 0 | 0 | 0 | NaN | NaN |
| Protein lin-28 homolog B | LIN28B | 0.000862501 | 0 | 0.0014721 | 0.00049131 | 0.001926441 | 0.00164023 | 0.96577247 | 0.541662254 |
| Uncharacterized protein C1orf122 | C1orf122 | 0.000460923 | 0.00053173 | 0.00104539 | 0.00096002 | 0.002921606 | 0.00174059 | 0.107465449 | 1.710816069 |
| Transcription elongation factor SPT6 | SUPT6H | 0.001240735 | 0.00107652 | 0.00206384 | 0.00023501 | 0.000393458 | 0.000177905 | 0.025989166 | -2.400431656 |
| Staphylococcal nuclease domain-containing protein 1 | SND1 | 0.008894861 | 0.00911766 | 0.01819892 | 0.01623932 | 0.039826683 | 0.028402298 | 0.108617477 | 1.639272303 |
| Basic leucine zipper and W2 domain-containing protein 1 | BZW1 | 0.001593215 | 0.00137255 | 0.00263504 | 0.00038415 | 0.001379971 | 0.000879135 | 0.145345653 | -0.85778434 |
| Eukaryotic translation initiation factor 3 subunit M | EIF3M | 0.01577259 | 0.01423951 | 0.03199196 | 0.02817503 | 0.042903256 | 0.055005355 | 0.129251715 | 1.443667557 |
| 7SK snRNA methylphosphate capping enzyme | MEPCE | 0 | 6.263E-05 | 3.7747E-05 | 0 | 0.000291156 | 5.73654E-05 | 0.304833583 | 1.410352287 |
| PHD finger-like domain-containing protein 5A | PHF5A | 0.005704288 | 0.00595111 | 0.00798009 | 0.00610375 | 0.007673082 | 0.00449927 | 0.828455679 | 0.036535167 |
| TRMT1-like protein | TRMT1L | 0.000661964 | 0.00077492 | 0.00145812 | 0.00235639 | 0.005557788 | 0.003411742 | 0.041515283 | 2.13839113 |
| 26S proteasome non-ATPase regulatory subunit 7 | PSMD7 | 0.208839996 | 0.21952266 | 0.25016897 | 0.06227705 | 0.063943692 | 0.064638302 | 0.047371008 | -1.779495937 |
| Nuclear fragile X mental retardation-interacting protein 2 | NUFIP2 | 9.88579E-05 | 5.0311E-05 | 0.00019902 | 6.3217E-05 | 0.000462691 | 0.000476 | 0.310643882 | 2.226620871 |
| 26S protease regulatory subunit 6B | PSMC4 | 0.356511029 | 0.37209254 | 0.40648461 | 0.08846208 | 0.092157887 | 0.078220232 | 0.035321373 | -2.072530447 |
| Myosin regulatory light chain 12A;Myosin regulatory light chain 12B;Myosin regulatory light polypeptide 9 | MYL12A;MYL12B;MYL9 | 0.001430579 | 0.00089066 | 0.00139392 | 0 | 0 | 0 | NaN | NaN |
| Vacuolar protein sorting-associated protein 13B | VPS13B | 8.0842E-05 | 0.00021769 | 0.00023553 | 8.4744E-05 | 0.000235397 | 0.00030998 | 0.818692303 | 0.112814171 |
| Protein PAT1 homolog 1 | PATL1 | 0.000209512 | 9.2417E-05 | 0.00041014 | 0.00019591 | 0.000320515 | 0.000144416 | 0.934826278 | -0.096805018 |
| Nucleolar protein 9 | NOP9 | 7.20132E-05 | 0.00020716 | 0.00055129 | 0.00037128 | 0.00090568 | 0.000567481 | 0.190436549 | 1.453826626 |
| Polyadenylate-binding protein 2 | PABPN1 | 0.002130948 | 0.00179849 | 0.00177779 | 0.00010015 | 0.001743748 | 0.000503242 | 0.173426337 | -1.837464306 |
| Zinc finger protein 598 | ZNF598 | 0 | 0.00011363 | 0.00027234 | 0.00012715 | 0.000474652 | 0.000358972 | 0.494829222 | 1.029014573 |
| | LOC100128510 | 0.000326582 | 0.00030416 | 0.00059305 | 0.0001819 | 9.90825E-05 | 0 | 0.085974274 | -1.282541246 |
| Cullin-associated NEDD8-dissociated protein 1 | CAND1 | 0.011402419 | 0.00278394 | 0.01624258 | 0.00254002 | 0.002701812 | 0.002041281 | 0.122232281 | -2.16642816 |
| EF-hand domain-containing protein D2 | EFHD2 | 0.003600605 | 0.00261028 | 0.00397918 | 0 | 0 | 0 | NaN | NaN |
| Methyltransferase-like protein 16 | METTL16 | 0 | 3.5021E-05 | 5.6652E-05 | 2.265E-05 | 2.12936E-05 | 1.53687E-05 | 0.129939695 | -1.064760183 |
| ATP-dependent RNA helicase DDX42 | DDX42 | 3.66846E-05 | 0.00015778 | 0.00011266 | 0.00024877 | 0.000828411 | 0.000600351 | 0.067687312 | 2.41384366 |
| Ubiquitin carboxyl-terminal hydrolase isozyme L5;Ubiquitin carboxyl-terminal hydrolase | UCHL5 | 0.102586496 | 0.11475976 | 0.1308704 | 0.01854002 | 0.019555056 | 0.018040697 | 0.022231095 | -2.629902163 |
| Transcriptional repressor p66-alpha;Transcriptional repressor p66-beta | GATAD2A;GATAD2B | 0.00010005 | 0.00013919 | 0.00040693 | 0.00014171 | 0.000161289 | 8.29929E-05 | 0.481183676 | 0.025844911 |
| WD repeat-containing protein 92 | HZGJ;WDR92;DKFZp434B156 | 0.000658325 | 0.00039003 | 0.00089019 | 0.00079935 | 0.001480559 | 0.000881258 | 0.227442477 | 0.420764464 |
| C2 domain-containing protein 5 | C2CD5 | 7.99854E-05 | 0 | 0.00034261 | 2.9032E-05 | 0 | 0 | 0.108470238 | -2.511470711 |
| ELKS/Rab6-interacting/CAST family member 1;ERC protein 2 | ERC1;ERC2 | 9.65401E-05 | 3.6997E-05 | 5.7084E-05 | 0 | 0.000171803 | 0 | 0.042821627 | 1.589601947 |
| Bifunctional lysine-specific demethylase and histidyl-hydroxylase MINA | MINA | 0.000206724 | 0.00034233 | 0.00071629 | 0.00080829 | 0.001881571 | 0.001752527 | 0.075222417 | 2.355968609 |
| Proteasome-associated protein ECM29 homolog | ECM29;KIAA0368 | 0.047467809 | 0.04580341 | 0.05445155 | 0.00966371 | 0.010508833 | 0.009282446 | 0.029855714 | -2.296300394 |
| 3-5 exoribonuclease 1 | ERI1 | 5.95622E-05 | 0.00059621 | 0.00089618 | 0.0005613 | 0.001083578 | 0.000814273 | 0.338790095 | 0.449689573 |
| Phosphatidylinositol 4-kinase alpha | PI4KA | 4.96165E-05 | 4.3063E-05 | 8.6235E-05 | 0 | 0 | 0 | NaN | NaN |
| WASH complex subunit FAM21B;WASH complex subunit FAM21C;WASH complex subunit FAM21A | FAM21C;FAM21A;FAM21B | 2.78009E-05 | 0 | 4.323E-05 | 0 | 0 | 0 | NaN | NaN |
| Coiled-coil domain-containing protein 50 | CCDC50 | 0 | 0 | 0 | 0 | 0 | 0 | NaN | NaN |
| Dynein assembly factor 5, axonemal | DNAAF5 | 0.000105011 | 0.00010589 | 0.00018543 | 0 | 0 | 0 | NaN | NaN |
| 26S proteasome non-ATPase regulatory subunit 8 | PSMD8 | 0.27044004 | 0.27629017 | 0.2692666 | 0.0473773 | 0.055778787 | 0.05661182 | 0.029071035 | -2.2775203 |
| Putative RNA polymerase II subunit B1 CTD phosphatase RPAP2 | RPAP2 | 9.25148E-05 | 0.00026077 | 0.00034388 | 5.9612E-05 | 0.000130239 | 0.000125966 | 0.216291651 | -1.049721393 |
| Probable ATP-dependent RNA helicase DHX37 | DHX37 | 0.000284755 | 0.00015189 | 3.8164E-05 | 0.00034212 | 0.000922784 | 0.000738234 | 0.101413257 | 2.281060792 |
| pre-rRNA processing protein FTSJ3 | FTSJ3 | 0 | 0 | 0.00032632 | 0.00116035 | 0.003879669 | 0.001784305 | 0.023569653 | 2.450994084 |
| Exocyst complex component 8 | EXOC8 | 0.00014102 | 0.00012257 | 0.00034262 | 3.8642E-05 | 8.21259E-05 | 0 | 0.112864772 | -1.323820626 |
| 5-3 exoribonuclease 1 | XRN1 | 0.000855111 | 0.00096403 | 0.00138698 | 0.00073477 | 0.001239308 | 0.000718469 | 0.629511796 | -0.391780458 |
| Spartin | SPG20 | 0.002033759 | 0.00198398 | 0.00216869 | 0.00078144 | 0.000876361 | 0.000746802 | 0.086719928 | -1.379934335 |
| Tetratricopeptide repeat protein 5 | TTC5 | 0.000203079 | 4.3815E-05 | 9.3004E-05 | 0 | 7.74929E-05 | 0 | 0.695166417 | -0.26322708 |
| Cell cycle and apoptosis regulator protein 2 | CCAR2 | 0.000255722 | 0.00017284 | 0.00047753 | 5.3992E-05 | 0.000161825 | 4.14585E-05 | 0.091613737 | -2.24374803 |
| Leucine-rich repeat-containing protein 47 | LRRC47 | 0.003799127 | 0.00372982 | 0.00968216 | 0.00526313 | 0.019486747 | 0.009025996 | 0.264035286 | 1.248418199 |
| Activating signal cointegrator 1 complex subunit 3 | ASCC3 | 0.000329929 | 0.00032055 | 0.00071488 | 0.00030524 | 0.000905667 | 0.0004524 | 0.712735654 | 0.455443336 |
| MAGUK p55 subfamily member 5 | MPP5 | 0 | 0 | 0 | 0 | 0 | 0 | NaN | NaN |
| Cohesin subunit SA-2 | STAG2 | 0.000187907 | 0.00030438 | 0.00068016 | 0.00058779 | 0.001756205 | 0.000936773 | 0.124774186 | 1.621844888 |
| COMM domain-containing protein 1 | COMMD1 | 0 | 0 | 0.00030942 | 0 | 0 | 0 | NaN | NaN |
| Protein KRI1 homolog | KRI1 | 0.000162042 | 0.00011635 | 0.0012476 | 0.0009251 | 0.004524049 | 0.002788239 | 0.110874611 | 4.104914528 |
| Procollagen galactosyltransferase 1 | COLGALT1 | 0.000107832 | 0.00027579 | 0.0001723 | 0 | 0 | 5.23824E-05 | 0.03289752 | -1.717760262 |
| Plasminogen activator inhibitor 1 RNA-binding protein | SERBP1 | 0.065211063 | 0.10223595 | 0.16710202 | 0.17605763 | 0.390980565 | 0.361172232 | 0.083938674 | 1.820784429 |
| Angiomotin | AMOT | 0.000336245 | 0.00022569 | 0.00038418 | 0 | 0 | 0 | NaN | NaN |
| Protein SMG8 | SMG8 | 0 | 0 | 0.00016117 | 8.8025E-05 | 0.000186737 | 0.000160002 | 0.571755754 | -0.010448531 |
| ATP-binding cassette sub-family F member 1 | ABCF1 | 0.003076027 | 0.00324108 | 0.00390291 | 0.00239047 | 0.005174589 | 0.004351475 | 0.778311282 | 0.425031657 |
| SHC SH2 domain-binding protein 1 | SHCBP1 | 0.00025828 | 9.0597E-05 | 0.00027423 | 0.00011565 | 0.00017412 | 0 | 0.618971572 | -0.864024811 |
| F-box only protein 22 | FBXO22 | 0.004135875 | 0.00453774 | 0.00656133 | 0.00324066 | 0.005495372 | 0.003790867 | 0.602918131 | -0.259445154 |
| N-acylneuraminate cytidylyltransferase | CMAS | 0.000185864 | 0.00020534 | 0.00029389 | 0.00012994 | 0.000604479 | 0.000408879 | 0.510345196 | 0.993666304 |
| THO complex subunit 2 | THOC2 | 0.000330523 | 0.00016649 | 0.00040819 | 0.00013918 | 0.000371613 | 0 | 0.700345689 | -0.53936386 |
| WD repeat-containing protein 36 | WDR36 | 8.6922E-05 | 0.00014149 | 0.00040413 | 0.00069162 | 0.001856123 | 0.001265446 | 0.048219332 | 3.160822966 |
| WD repeat-containing protein 48 | WDR48 | 0.000231083 | 0.00029381 | 0.00038931 | 0.00040982 | 0.000664711 | 0.000569779 | 0.180952377 | 0.95550422 |
| Kelch domain-containing protein 4 | KLHDC4 | 0.001978278 | 0.00116644 | 0.00242333 | 0.00244423 | 0.003354362 | 0.00119188 | 0.656551634 | 0.305134281 |
| V-type proton ATPase subunit a;V-type proton ATPase 116 kDa subunit a isoform 2 | ATP6V0A2 | 0.012775725 | 0.01400188 | 0.017334 | 0.03175724 | 0.035353129 | 0.047691559 | 0.086338029 | 1.33621695 |
| Protein bicaudal D homolog 2 | BICD2 | 0 | 0 | 0 | 0 | 0 | 0 | NaN | NaN |
| ATP-dependent RNA helicase DDX54 | DDX54 | 0.000518525 | 0.00062707 | 0.00101562 | 0.00220201 | 0.005806646 | 0.004213611 | 0.026642267 | 2.748358725 |
| Ribosome biogenesis protein BRX1 homolog | BRIX1 | 0.000509646 | 0.00076181 | 0.00268298 | 0.00624469 | 0.011203975 | 0.009834485 | 0.038752315 | 3.690340693 |
| D-tyrosyl-tRNA(Tyr) deacylase 1 | DTD1 | 0.000650543 | 0.0009865 | 0.00247383 | 0.00173821 | 0.003093449 | 0.002687372 | 0.179467203 | 1.445810934 |
| Gem-associated protein 5 | GEMIN5 | 0.001907737 | 0.00208332 | 0.00364919 | 0.00246079 | 0.005937772 | 0.004404829 | 0.232299681 | 1.080198095 |
| Dynein light chain roadblock-type 1;Dynein light chain roadblock-type 2 | DYNLRB1;DYNLRB2 | 0.000614825 | 0.00122077 | 0.00293826 | 0.00193055 | 0.002818741 | 0.00241106 | 0.267717892 | 0.981878503 |
| DIS3-like exonuclease 1 | DIS3L | 0 | 0 | 3.6233E-05 | 0 | 0.000103701 | 9.41817E-05 | 0.057805432 | 1.447582144 |
| Nucleolar complex protein 3 homolog | NOC3L | 0.000189867 | 9.8173E-05 | 0.00048751 | 0.00099259 | 0.003760464 | 0.002387888 | 0.045084424 | 3.652674087 |
| Zinc finger CCCH domain-containing protein 15 | ZC3H15 | 0.002969208 | 0.002179 | 0.0067418 | 0.00546955 | 0.007756124 | 0.007719599 | 0.156603815 | 1.378447747 |
| Peptidyl-prolyl cis-trans isomerase-like 4 | PPIL4 | 0.001598813 | 0.00207317 | 0.00259046 | 0.00206461 | 0.002760745 | 0.002556713 | 0.648122269 | 0.302451104 |

| | | | | | | | | | |
|---|---|---|---|---|---|---|---|---|---|
| General transcription factor 3C polypeptide 2 | GTF3C2 | 0.000155475 | 0.00010909 | 0.00071827 | 0 | 1.29349E-05 | 3.89465E-05 | 0.086600312 | -2.79222942 |
| Programmed cell death 6-interacting protein | PDCD6IP | 0.002295824 | 0.00083125 | 0.00135967 | 1.0094E-05 | 0.000204096 | 7.58933E-05 | 0.052891356 | -4.163138083 |
| Ubiquitin carboxyl-terminal hydrolase 15 | USP15 | 0.002578379 | 0.00224788 | 0.0027155 | 0.00029451 | 0.000260859 | 0.000291539 | 0.015894364 | -3.144703548 |
| 26S protease regulatory subunit 6A | PSMC3 | 0.000609002 | 0.00094452 | 0.00066804 | 0 | 0 | 0 | NaN | NaN |
| Probable tRNA pseudouridine synthase 1 | TRUB1 | 7.52883E-05 | 4.4297E-05 | 0.00032534 | 0 | 0.00029267 | 0.000108441 | 0.564743734 | 1.242595008 |
| Negative elongation factor B | NELFB | 0.000588848 | 0.00091344 | 0.00123363 | 0.00134301 | 0.0019985 | 0.001339071 | 0.193026949 | 0.556085521 |
| Mitotic spindle assembly checkpoint protein MAD1 | MAD1L1 | 0.000165485 | 0 | 0.00012045 | 0 | 0 | 0 | NaN | NaN |
| Laforin, isoform 9 | EPM2A | 0 | 0 | 0 | 0.00041241 | 0.000896385 | 0.000536773 | NaN | NaN |
| Paraspeckle component 1 | PSPC1 | 0.000818833 | 0.00095995 | 0.00206229 | 0.00054251 | 0.002201685 | 0.002010515 | 0.825432152 | 1.066533931 |
| DnaJ homolog subfamily C member 9 | DNAJC9 | 0.001339996 | 0.00136188 | 0.00283542 | 0.00113734 | 0.004738886 | 0.002591958 | 0.536955911 | 0.92844699 |
| Microprocessor complex subunit DGCR8 | DGCR8 | 0 | 0 | 6.7755E-05 | 0 | 0.000110343 | 7.03067E-05 | 0.426694203 | 0.378465278 |
| Titin | TTN | 2.36267E-05 | 4.3766E-05 | 5.0733E-05 | 9.203E-05 | 0.000163646 | 0.000127031 | 0.053437541 | 1.537284983 |
| ATP-dependent RNA helicase DDX1 | DDX1 | 0.008093719 | 0.00955784 | 0.01667184 | 0.01647476 | 0.030808588 | 0.024248856 | 0.126074371 | 1.343160919 |
| Histone H1x | H1FX | 0.001993898 | 0.00217555 | 0.00266362 | 0.00200312 | 0.004635486 | 0.003980722 | 0.316833259 | 0.871651937 |
| Heat shock protein 105 kDa | HSPH1 | 0.000183843 | 0.0001768 | 0.00042949 | 7.6688E-05 | 2.41224E-05 | 0 | 0.070398299 | -2.095718955 |
| Cell differentiation protein RCD1 homolog | RQCD1 | 0.000163879 | 0.0004095 | 0.00065994 | 0.00032154 | 0.000675614 | 0.000318976 | 0.778239452 | -0.348882845 |
| | PSMC1 | 0.00527774 | 0.00435308 | 0.00606768 | 0.00016712 | 0 | 0 | 0.005031315 | -4.980929447 |
| La-related protein 4B | LARP4B | 0.000786194 | 0.00071466 | 0.0013382 | 0.0011382 | 0.004173531 | 0.002560023 | 0.12491738 | 1.703199513 |
| Translational activator GCN1 | GCN1L1 | 0.002115049 | 0.00239276 | 0.00486218 | 0.00439822 | 0.012202221 | 0.008023561 | 0.106089326 | 1.745569386 |
| Pre-mRNA-splicing factor ATP-dependent RNA helicase PRP16 | DHX38 | 0.000470099 | 0.0007441 | 0.00080194 | 4.1895E-05 | 0.000431899 | 0.000145328 | 0.118756781 | -2.356185767 |
| Acidic leucine-rich nuclear phosphoprotein 32 family member B | ANP32B | 0.023716269 | 0.0234429 | 0.04565257 | 0.03154975 | 0.036752789 | 0.035476585 | 0.677121627 | 0.580990076 |
| Spermatogenesis-associated serine-rich protein 2 | SPATS2 | 0 | 0 | 0 | 0.00015763 | 0.000346044 | 0.000220173 | NaN | NaN |
| DNA repair protein RAD50 | RAD50 | 0.000470378 | 0.00068676 | 0.00141191 | 0.0009088 | 0.001935013 | 0.001110388 | 0.279969517 | 0.693190165 |
| Regulator of nonsense transcripts 1 | UPF1 | 0.000912384 | 0.00082212 | 0.00151329 | 0.00125571 | 0.002470634 | 0.001669555 | 0.227438606 | 0.871750569 |
| COP9 signalosome complex subunit 5 | COPS5 | 0.000591703 | 0.00047124 | 0.00080567 | 0.00039696 | 0.000582192 | 0.000148544 | 0.231170935 | -0.575890473 |
| G patch domain and KOW motifs-containing protein | GPKOW | 0.001315922 | 0.00192945 | 0.00196237 | 0.00046316 | 0.000984222 | 0.00068192 | 0.090586895 | -1.500512264 |
| Transportin-1 | TNPO1 | 0.000386323 | 0.0002824 | 0.00041177 | 0.00035437 | 0.000262024 | 0.000198053 | 0.433641791 | -0.560108611 |
| Ribosomal RNA small subunit methyltransferase NEP1 | EMG1 | 0.000776341 | 0.00160327 | 0.0028893 | 0.01115411 | 0.015090877 | 0.014886769 | 0.019779448 | 3.214944958 |
| Probable ubiquitin carboxyl-terminal hydrolase FAF-X | USP9X | 0.001392621 | 0.00124924 | 0.00189762 | 0.00076406 | 0.000885402 | 0.000731633 | 0.163010707 | -0.866044844 |
| Ubiquitin carboxyl-terminal hydrolase 7;Ubiquitin carboxyl-terminal hydrolase | USP7 | 0.002377505 | 0.00252528 | 0.00279532 | 0.00463452 | 0.00311729 | 0.007979165 | 0.160752507 | 0.875976876 |
| 60S ribosomal protein L36a-like | RPL36AL | 0.067411264 | 0.05684078 | 0.12841355 | 0.1045455 | 0.25434966 | 0.05329374 | 0.539480018 | 0.633069424 |
| Exocyst complex component 4 | EXOC4 | 0.000160665 | 0.00023903 | 0.00054059 | 0.00014578 | 0.000195256 | 0 | 0.350404221 | -0.502627188 |
| Protein mago nashi homolog 2 | MAGOHB | 0.001301814 | 0.00170114 | 0.00435236 | 0.00138021 | 0.004384351 | 0.002815305 | 0.729900387 | 0.726788503 |
| Exosome complex component RRP43 | EXOSC8 | 0.003956948 | 0.00398849 | 0.00501927 | 0.00723249 | 0.013086216 | 0.01026015 | 0.107044675 | 1.363139066 |
| FK506-binding protein 15 | FKBP15 | 9.36457E-05 | 7.4337E-05 | 0.00012987 | 0 | 0 | 0 | NaN | NaN |
| Coiled-coil domain-containing protein 124 | CCDC124 | 0.02271918 | 0.02149691 | 0.04614303 | 0.05766059 | 0.131127362 | 0.085674228 | 0.062895896 | 1.914950522 |
| DDB1- and CUL4-associated factor 8 | DCAF8 | 0.000171437 | 0.00012811 | 0.00021617 | 0 | 0 | 0 | NaN | NaN |
| BTB/POZ domain-containing protein KCTD12 | KCTD12 | 0.00099289 | 0 | 0.00088286 | 0.00126184 | 0.000456423 | 0 | 0.710710706 | -0.302993118 |
| U5 small nuclear ribonucleoprotein 40 kDa protein | SNRNP40 | 0.000311852 | 0.00040468 | 0.00061185 | 0.00079847 | 0.001093248 | 0.00080849 | 0.130196925 | 0.998451944 |
| Enhancer of mRNA-decapping protein 3 | EDC3 | 0.000583193 | 0.00047846 | 0.00086436 | 0.00060086 | 0.001471585 | 0.000727018 | 0.392892074 | 0.318017681 |
| tRNA (adenine(58)-N(1))-methyltransferase catalytic subunit TRMT61A | TRMT61A | 0.000976934 | 6.7082E-05 | 0.00124852 | 0 | 0.006603983 | 0.000125071 | 0.725852917 | -0.104255192 |
| Protein LTV1 homolog | LTV1 | 0.003151103 | 0.00457814 | 0.01034889 | 0.00215202 | 0.009925262 | 0.006910914 | 0.995343347 | 0.594114 |
| Target of EGR1 protein 1 | TOE1 | 0.00286838 | 0.00157804 | 0.00408092 | 0.00157383 | 0.000736609 | 0.000692824 | 0.086682022 | -1.961265234 |
| Zinc finger CCCH-type antiviral protein 1-like | ZC3HAV1L | 0.002047083 | 0.00169203 | 0.00117947 | 0.00157889 | 0.001330547 | 0.001068387 | 0.603838862 | -0.346739086 |
| Alpha-soluble NSF attachment protein | NAPA | 0 | 8.8509E-05 | 0.00012176 | 0 | 0 | 0 | NaN | NaN |
| Putative RNA-binding protein Luc7-like 2 | LUC7L2 | 0.000859198 | 0.00085539 | 0.00139291 | 0.00021143 | 0.000642879 | 0.000129785 | 0.087857511 | -2.022824186 |
| Serine/threonine-protein phosphatase 1 regulatory subunit 10 | PPP1R10 | 0 | 0 | 0 | 0.00011407 | 0.000188063 | 8.47142E-05 | NaN | NaN |
| Importin-7 | IPO7 | 4.31822E-05 | 2.3171E-05 | 3.1312E-05 | 0 | 0 | 0 | NaN | NaN |
| Palladin | PALLD | 0.000359708 | 0.00055081 | 0.00047627 | 0 | 0 | 0 | NaN | NaN |
| Exocyst complex component 2 | EXOC2 | 9.59915E-05 | 4.2289E-05 | 0.00013902 | 3.4216E-05 | 4.32368E-05 | 3.4207E-05 | 0.119475989 | -1.488235549 |
| Zinc finger RNA-binding protein | ZFR | 0.000375994 | 0.00062392 | 0.00188054 | 0.00258754 | 0.006335302 | 0.004955506 | 0.064031402 | 2.989601727 |
| Sodium channel and clathrin linker 1 | SCLT1 | 0.000552973 | 0.00037771 | 0.00052786 | 0 | 0 | 0 | NaN | NaN |
| E3 SUMO-protein ligase RanBP2 | RANBP2 | 0 | 1.1198E-05 | 1.5916E-05 | 0 | 0 | 0 | NaN | NaN |
| DNA replication licensing factor MCM7 | MCM7 | 4.34632E-05 | 0 | 3.2196E-05 | 0 | 0 | 0 | NaN | NaN |
| Probable 28S rRNA (cytosine-C(5))-methyltransferase;Putative methyltransferase NSUN5C | NSUN5;NSUN5P2 | 0.000169723 | 0.00018292 | 0.00044322 | 0.00036323 | 0.001240567 | 0.00081883 | 0.11395936 | 2.162382684 |
| RNA-binding protein 14 | RBM14 | 0.000339111 | 0.0006326 | 0.00100292 | 0.00057948 | 0.002005867 | 0.000792542 | 0.366871589 | 0.325200677 |
| Transgelin-2 | TAGLN2 | 0.016187437 | 0.01219198 | 0.01417207 | 0.00046641 | 0.000505498 | 0.000508227 | 0.008774896 | -4.809201404 |
| Vacuolar protein sorting-associated protein 35 | VPS35 | 0.002884783 | 0.00167947 | 0.00296858 | 0.00011179 | 0.001248255 | 0.000546042 | 0.109089938 | -2.401378972 |
| Transcriptional activator protein Pur-beta | PURB | 0.000627197 | 0.00065011 | 0.00205032 | 0.00076519 | 0.002363751 | 0.001648583 | 0.45201973 | 1.342470772 |
| NudC domain-containing protein 1 | NUDCD1 | 0.000398393 | 0.00017548 | 0.00043615 | 0.00010852 | 0.00025964 | 0 | 0.273042786 | -1.246955492 |
| ATPase WRNIP1 | WRNIP1 | 0.006892845 | 0.00942782 | 0.00778245 | 0.00453368 | 0.006111758 | 0.004880732 | 0.269600196 | -0.673126125 |
| SRSF protein kinase 1 | SRPK1 | 0.005034151 | 0.00597268 | 0.01187688 | 0.01315809 | 0.028266431 | 0.017648914 | 0.082867759 | 1.56312932 |
| Spermatid perinuclear RNA-binding protein | STRBP | 3.71442E-05 | 0.00015421 | 0.00017171 | 0.00036224 | 0.000872356 | 0.000782689 | 0.061535611 | 2.343514781 |
| Proteasome subunit beta type-7 | PSMB7 | 1.03073564 | 1.11891552 | 1.06986939 | 1.44525077 | 1.198202542 | 1.294596437 | 0.605187296 | 0.27506775 |
| Cell division cycle 5-like protein | CDC5L | 0.002172321 | 0.00050571 | 0.00495452 | 0.01535774 | 0.021265453 | 0.012597021 | 0.062747506 | 2.821656533 |
| 26S protease regulatory subunit 7 | PSMC2 | 0.595342067 | 0.57106668 | 0.59384419 | 0.12275439 | 0.137099691 | 0.131981307 | 0.034639846 | -2.169750836 |
| Synaptic vesicle membrane protein VAT-1 homolog | VAT1 | 0.000875546 | 0 | 0 | 0 | 0 | 0 | NaN | NaN |
| DnaJ homolog subfamily C member 2;DnaJ homolog subfamily C member 2, N-terminally processed | DNAJC2 | 0.002571884 | 0.00248927 | 0.00394429 | 0.00570177 | 0.013224575 | 0.010303963 | 0.059511086 | 2.002301839 |
| Ribonucleases P/MRP protein subunit POP1 | POP1 | 0.000492537 | 0.0004832 | 0.0014068 | 0.00124346 | 0.003126264 | 0.002434232 | 0.101105834 | 2.305162663 |
| Protein S100-A13 | S100A13 | 0.001914791 | 0.00073873 | 0.00248059 | 0 | 0.001051727 | 0.001609717 | 0.77943193 | -0.557403259 |
| Eukaryotic translation initiation factor 3 subunit C;Eukaryotic translation initiation factor 3 subunit C-like protein | EIF3C;EIF3CL | 0.010710447 | 0.00842406 | 0.01924468 | 0.02124604 | 0.042488046 | 0.031515919 | 0.089037401 | 1.55706204 |
| Tetratricopeptide repeat protein 1 | TTC1 | 0.000519136 | 0.00025075 | 0.00078035 | 0.00017967 | 0.00036809 | 0.000319482 | 0.257551083 | -0.700377886 |
| 3-hydroxyacyl-CoA dehydrogenase type-2 | HSD17B10 | 0.000175999 | 0.00020575 | 0.00050501 | 0.0002495 | 0.001902332 | 0.000852575 | 0.211211887 | 2.050955357 |
| T-complex protein 1 subunit eta | CCT7 | 0.002536502 | 0.0018322 | 0.00209179 | 0.00388099 | 0.002633503 | 0.00375056 | 0.256654294 | 0.842368689 |
| A-kinase anchor protein 9 | AKAP9 | 0.000248315 | 0.00037209 | 0.00025244 | 8.9865E-05 | 9.22182E-05 | 0 | 0.073870511 | -1.471479545 |
| UDP-glucose:glycoprotein glucosyltransferase 1 | UGGT1 | 0.000145476 | 0.00014085 | 9.6657E-05 | 0 | 0 | 0 | NaN | NaN |
| Glutamate-rich WD repeat-containing protein 1 | GRWD1 | 0.003492106 | 0.00292421 | 0.00600859 | 0.00462055 | 0.011089324 | 0.007500341 | 0.171919045 | 1.102858837 |
| Methylosome protein 50 | WDR77 | 0.001430243 | 0.0014876 | 0.00211252 | 0.00064425 | 0.003121442 | 0.002182261 | 0.987121394 | 0.552840886 |
| Myb-binding protein 1A | MYBBP1A | 0.000904882 | 0.00086635 | 0.00193068 | 0.00385293 | 0.008497877 | 0.005494499 | 0.030391439 | 2.602186362 |
| Coronin-1B | CORO1B | 0.008596462 | 0.00945504 | 0.00827765 | 0.00029824 | 0.00175366 | 0.001299659 | 0.036197466 | -2.725609876 |
| Partner of Y14 and mago | WIBG | 0.014202217 | 0.016622 | 0.03285506 | 0.02527012 | 0.056543577 | 0.044058027 | 0.144828697 | 1.406310562 |
| Serine/threonine-protein kinase RIO1 | RIOK1 | 0.001054305 | 0.00117052 | 0.00114979 | 0.00068758 | 0.001723122 | 0.001030572 | 0.897526078 | -0.157918814 |
| Protein LLP homolog | LLPH | 0.000403275 | 0.00069759 | 0.00120522 | 0 | 0.001911172 | 0 | 0.063867983 | 1.454002222 |
| Protein pelota homolog | PELO | 0.001132068 | 0.00092053 | 0.00243675 | 0.0009269 | 0.002829188 | 0.00183949 | 0.662064217 | 0.700345214 |
| Sorting nexin-27 | SNX27 | 0.000742526 | 0.00058267 | 0.00051777 | 0 | 0 | 0 | NaN | NaN |
| Extended synaptotagmin-1 | ESYT1 | 3.96428E-05 | 8.3526E-05 | 7.3409E-05 | 0 | 1.99421E-05 | 0 | 0.029801397 | -1.88014747 |
| Mini-chromosome maintenance complex-binding protein | MCMBP | 0.000114998 | 6.2893E-05 | 0.00039108 | 0.000113 | 0.000169459 | 0 | 0.976854268 | 0.267039897 |
| Acidic leucine-rich nuclear phosphoprotein 32 family member E | ANP32E | 0.003700314 | 0.00610929 | 0.00784665 | 0.00275783 | 0.00313122 | 0.003103495 | 0.163890635 | -0.977109723 |
| WD repeat-containing protein 18 | WDR18 | 0.000785354 | 0.00081142 | 0.0022877 | 0.00151244 | 0.002996343 | 0.00186568 | 0.229713101 | 1.20118079 |
| Tubulin beta-2B chain | TUBB2B | 0.000249255 | 0.00013695 | 0.00019177 | 0.00025123 | 0.000638526 | 0 | 0.161175522 | 1.062486746 |
| U3 small nucleolar RNA-associated protein 14 homolog A | UTP14A | 3.46305E-05 | 0 | 0.00012916 | 0.00021932 | 0.000390766 | 0.000198948 | 0.107539069 | 1.713382397 |
| Oxysterol-binding protein 1 | OSBP | 8.20849E-05 | 0 | 0.00010604 | 0 | 0 | 0 | NaN | NaN |
| Guanine nucleotide-binding protein-like 3 | GNL3 | 0.004443063 | 0.00431153 | 0.00693908 | 0.0072183 | 0.020894443 | 0.009365787 | 0.1209767 | 1.07584563 |
| Serine/threonine-protein kinase RIO2 | RIOK2 | 0.000151282 | 0.00029843 | 0.0006466 | 0 | 0.0001476 | 0.000172877 | 0.287926115 | -0.90167324 |
| Kinesin-like protein KIFC1 | KIFC1 | 0 | 0 | 5.3221E-05 | 0 | 0.000122404 | 2.67627E-05 | 0.931827283 | 0.104897485 |
| RNA-binding protein 4;RNA-binding protein 4B | RBM4;RBM4B | 0.004266767 | 0.00317816 | 0.00776665 | 0.00233554 | 0.008549711 | 0.004537241 | 0.921703351 | 0.088671962 |
| Splicing factor 3B subunit 5 | SF3B5 | 0.000848337 | 0 | 0 | 0 | 0 | 0 | NaN | NaN |
| GTP-binding protein 2 | GTPBP2 | 8.95812E-05 | 0.00012291 | 0.00020832 | 9.7018E-05 | 0.000273409 | 0.000346957 | 0.362564427 | 1.153411485 |
| N-alpha-acetyltransferase 15, NatA auxiliary subunit | NAA15 | 0.000773486 | 0.00057257 | 0.00277012 | 0.00020043 | 0.000918257 | 0.000566566 | 0.275672064 | -0.449131152 |
| Serrate RNA effector molecule homolog | SRRT | 0.004901075 | 0.0060626 | 0.01193534 | 0.00242197 | 0.004805454 | 0.002901178 | 0.117466578 | -1.063297349 |
| Queuine tRNA-ribosyltransferase | QTRT1 | 0.00014476 | 0.00036639 | 0.00062587 | 0.00048316 | 0.000914507 | 0.000659819 | 0.206352982 | 0.848698069 |
| Fanconi anemia group D2 protein | FANCD2 | 0.000493842 | 0.00047064 | 0.0014204 | 0.0019428 | 0.004264074 | 0.003299581 | 0.054061991 | 2.74016139 |
| Eukaryotic translation initiation factor 2A;Eukaryotic translation initiation factor 2A, N-terminally processed | EIF2A | 0.002293808 | 0.00271559 | 0.00568098 | 0.00532301 | 0.013361193 | 0.008199396 | 0.105704238 | 1.594253555 |
| MKI67 FHA domain-interacting nucleolar phosphoprotein | NIFK | 0 | 0 | 0.00060386 | 0.00042675 | 0.001449485 | 0.003453997 | 0.274504664 | 1.26324722 |
| Nucleolar GTP-binding protein 1 | GTPBP4 | 0.001957955 | 0.00292589 | 0.00489771 | 0.00720206 | 0.019603273 | 0.015669555 | 0.047843143 | 2.421018252 |
| WD repeat-containing protein 11 | WDR11 | 0.000246579 | 0.00025825 | 0.00035243 | 0.00014376 | 0.000135412 | 9.81151E-05 | 0.110400772 | -0.931435101 |
| Regulator of nonsense transcripts 3B | UPF3B | 0.000210396 | 0.00049954 | 0.00188446 | 0.00102327 | 0.003538795 | 0.002752994 | 0.172045359 | 2.462332244 |
| Ubiquitin-like protein 5 | UBL5 | 0.009277237 | 0.00788517 | 0.00952324 | 0.00555502 | 0.008911346 | 0.007485737 | 0.586542698 | -0.309550835 |
| Protein phosphatase 1 regulatory subunit 12C | PPP1R12C | 0.000370048 | 0.0001741 | 0.00095077 | 0.00046462 | 0.001454843 | 0.001424048 | 0.221442608 | 1.944213452 |

| | | | | | | | | | |
|---|---|---|---|---|---|---|---|---|---|
| Protein dpy-30 homolog | DPY30 | 0.003687661 | 0.00359573 | 0.00505316 | 0.00303134 | 0.00506717 | 0.004712881 | 0.945924794 | 0.353903216 |
| Ribosome biogenesis protein WDR12 | WDR12 | 0.001037958 | 0.00073716 | 0.00173084 | 0.00320733 | 0.006142163 | 0.004008957 | 0.041526588 | 1.949478876 |
| PITH domain-containing protein 1 | PITHD1 | 0.001088736 | 0.00139569 | 0.00220431 | 0 | 0.000202395 | 0.000113268 | 0.018063592 | -3.204446538 |
| DNA-directed RNA polymerase I subunit RPA49 | POLR1E | 0.003361438 | 0.00631362 | 0.00874171 | 0.00309778 | 0.003575227 | 0.003172038 | 0.197326435 | -0.993057149 |
| Protein FAM192A | FAM192A | 0.018504647 | 0.01792945 | 0.01772765 | 0.01078858 | 0.027965069 | 0.015314964 | 0.836015825 | -0.227388936 |
| N-acetyltransferase 10 | NAT10 | 0.000297979 | 0.00050316 | 0.00100566 | 0.00155341 | 0.004428684 | 0.002761172 | 0.053047494 | 2.456188751 |
| 5-3 exoribonuclease 2 | XRN2 | 0.000368021 | 0.00027598 | 0.00129007 | 0.00077171 | 0.003419992 | 0.002058417 | 0.151414025 | 2.48367528 |
| Probable ATP-dependent RNA helicase DDX47 | DDX47 | 0.000537588 | 0.00153177 | 0.0018925 | 0.00053224 | 0.004682632 | 0.00171699 | 0.68326833 | 0.164681948 |
| Activating signal cointegrator 1 complex subunit 2 | ASCC2 | 0.000330333 | 0.0007627 | 0.00913152 | 0.00027846 | 0.001391757 | 0.000902463 | 0.609321719 | 0.24274641 |
| Peptidyl-prolyl cis-trans isomerase-like 3;Peptidyl-prolyl cis-trans isomerase | PPIL3 | 0.000130433 | 0.00067572 | 0.00131482 | 0.00027865 | 0.000948889 | 0.000843482 | 0.802876187 | 0.31992105 |
| ATP-dependent RNA helicase DHX36 | DHX36 | 0.001854831 | 0.00178708 | 0.00130523 | 0.00141106 | 0.00196689 | 0.000519326 | 0.421933514 | -0.340824239 |
| 5-AMP-activated protein kinase catalytic subunit alpha-1 | PRKAA1 | 0.000229179 | 0.00016801 | 0.00024009 | 0 | 0 | 0 | NaN | NaN |
| Golgi resident protein GCP60 | ACBD3 | 0 | 0 | 0.00010038 | 0 | 0 | 0 | NaN | NaN |
| Probable serine carboxypeptidase CPVL | CPVL | 0 | 6.3252E-05 | 4.9047E-05 | 0 | 0 | 0 | NaN | NaN |
| Charged multivesicular body protein 4b | CHMP4B | 0.001453253 | 0.00012408 | 0.00244826 | 0 | 0 | 0.00013683 | 0.158327812 | -3.408829406 |
| Sentrin-specific protease 3 | SENP3;SENP3-EIF4A1 | 0.000271302 | 0.00015169 | 0.00072572 | 0.00024039 | 0.001109026 | 0.000691929 | 0.391482435 | 1.350724325 |
| Protein LSM12 homolog | LSM12 | 0 | 0 | 0 | 0 | 0.000831652 | 0.000593068 | NaN | NaN |
| Probable ATP-dependent RNA helicase YTHDC2 | YTHDC2 | 0.0014261 | 0.00136784 | 0.00254398 | 0.00202516 | 0.004533424 | 0.003541038 | 0.165717492 | 1.312097177 |
| RNA polymerase II-associated protein 3 | RPAP3 | 0.000690012 | 0.00070714 | 0.00106905 | 0.00048417 | 0.001496899 | 0.001164521 | 0.695421255 | 0.719670922 |
| Ribosome production factor 2 homolog | RPF2 | 0.000111544 | 0 | 0 | 0.00029706 | 0.000811642 | 0 | 0.073398912 | 2.138186026 |
| UPF0488 protein C8orf33 | C8orf33 | 0.020173553 | 0.0066062 | 0.0286017 | 0.02228995 | 0.04271976 | 0.029621264 | 0.24116138 | 0.554168008 |
| Phosphorylated adapter RNA export protein | PHAX | 0.004468816 | 0.00390736 | 0.00709444 | 0.00033618 | 0.005231781 | 0.001884549 | 0.22494076 | -1.245673337 |
| 5-nucleotidase domain-containing protein 2 | NT5DC2 | 0.000472915 | 0.00080722 | 0.00091698 | 0 | 0.000162227 | 0 | 0.017732656 | -2.314953329 |
| CCR4-NOT transcription complex subunit 10 | CNOT10 | 0.000139066 | 0.00013847 | 0.00030656 | 0.0001373 | 0.000291919 | 0.000196768 | 0.801469075 | 0.500725876 |
| DNA-directed RNA polymerase I subunit RPA2 | POLR1B | 0.000531525 | 0.00053997 | 0.00171267 | 0.00044244 | 0.001081756 | 0.000827826 | 0.890382588 | 0.616456321 |
| Regulator of nonsense transcripts 2 | UPF2 | 0 | 0 | 0 | 0 | 0 | 0 | NaN | NaN |
| Exportin-5 | XPO5 | 0.000104468 | 9.5207E-05 | 0.00019832 | 4.2978E-05 | 8.90465E-05 | 0 | 0.172741107 | -0.75591056 |
| Calcyclin-binding protein | CACYBP | 0.02672881 | 0.00275788 | 0.02308704 | 0.03082299 | 0.020039914 | 0 | 0.488912943 | 0.106358258 |
| Echinoderm microtubule-associated protein-like 4 | EML4 | 4.77315E-05 | 3.1591E-05 | 0.00017299 | 0.00017542 | 0.000741753 | 0.000707312 | 0.082348163 | 3.889333158 |
| GPN-loop GTPase 1 | GPN1 | 0.001514388 | 0.00109254 | 0.0005909 | 0.00086044 | 0.004016019 | 0.003700711 | 0.217550468 | 1.760121657 |
| Pre-mRNA-splicing factor SYF1 | XAB2 | 0.000244989 | 0.00012391 | 0.00021126 | 6.5014E-05 | 0.0003023 | 9.12589E-05 | 0.448735965 | -1.210989293 |
| RNA polymerase II subunit A C-terminal domain phosphatase SSU72 | SSU72 | 0.000116521 | 0.00020036 | 0.00045325 | 0 | 0.000347946 | 0.000260189 | 0.587522986 | 0.586645982 |
| Exosome complex component RRP41 | EXOSC4 | 0.005064942 | 0.00469538 | 0.00710211 | 0.00480514 | 0.011221119 | 0.006977899 | 0.484727707 | 0.462246967 |
| Small subunit processome component 20 homolog | UTP20 | 0 | 0 | 0 | 0 | 9.39701E-05 | 6.39801E-05 | NaN | NaN |
| Exosome complex component RRP40 | EXOSC3 | 0.003645224 | 0.00547451 | 0.00826359 | 0.00511498 | 0.017523205 | 0.010484081 | 0.248965885 | 0.937398395 |
| Nucleolar protein 6 | NOL6 | 0 | 0 | 0 | 0.00013833 | 0.000217918 | 0.00023727 | NaN | NaN |
| Procollagen-lysine,2-oxoglutarate 5-dioxygenase 3 | PLOD3 | 0.000144821 | 0.00017542 | 0.00021235 | 0 | 0 | 0 | NaN | NaN |
| Nucleolar RNA helicase 2 | DDX21 | 0.017275221 | 0.01887889 | 0.04400859 | 0.04265656 | 0.096760022 | 0.074639276 | 0.090598432 | 1.983161355 |
| DNA polymerase epsilon subunit 4 | POLE4 | 0 | 0.00071828 | 0.002274 | 0.00037226 | 0.001580611 | 0.001124116 | 0.626606185 | -0.185130687 |
| Translation initiation factor eIF-2B subunit gamma | EIF2B3 | 0.000469567 | 0.00038921 | 0.00077843 | 0.00093432 | 0.001633919 | 0 | 0.120717983 | 1.395761176 |
| RNA-binding protein PNO1 | PNO1 | 0.003858415 | 0.00432618 | 0.00874463 | 0.00317744 | 0.006721983 | 0.004508616 | 0.717472764 | 0.059590948 |
| Phenylalanine--tRNA ligase beta subunit | FARSB | 0.011231665 | 0.01345965 | 0.02301854 | 0.03084341 | 0.057079332 | 0.046464804 | 0.06997851 | 1.787497856 |
| Sister chromatid cohesion protein PDS5 homolog B | PDS5B | 8.72915E-05 | 9.6419E-05 | 0.00015992 | 0.00035484 | 0.000783086 | 0.000511284 | 0.031536577 | 2.406737003 |
| Midasin | MDN1 | 9.58963E-05 | 0.0001173 | 0.00036086 | 9.8109E-05 | 0.000360041 | 0.000122366 | 0.973426499 | 0.060962843 |
| Zinc finger CCHC domain-containing protein 3 | ZCCHC3 | 6.23166E-05 | 0 | 0 | 0 | 0 | 0 | NaN | NaN |
| Gamma-taxilin | TXLNG | 0.002698578 | 0.00239736 | 0.00391259 | 0.00439671 | 0.006742744 | 0.004625645 | 0.192877584 | 0.777455161 |
| Exocyst complex component 1 | EXOC1 | 0.000165093 | 0.00012745 | 0.00029107 | 0.00010478 | 0.000118315 | 9.16152E-05 | 0.193717918 | -0.655929353 |
| Fanconi anemia group I protein | FANCI | 0.000965066 | 0.0028747 | 0.00196447 | 0.00147116 | 0.002182264 | 0.001571921 | 0.94637609 | -0.321612891 |
| Protein asunder homolog | ASUN | 0.000170938 | 0.00013305 | 0.00011434 | 4.8916E-05 | 5.56569E-05 | 0 | 0.100550433 | -1.350433529 |
| ATP-dependent RNA helicase DDX18 | DDX18 | 0.000304428 | 0.00026053 | 0.00026446 | 0.00089495 | 0.00187608 | 0.001557025 | 0.029935092 | 2.557654309 |
| RNA-binding protein 28 | RBM28 | 0.000297122 | 0.00035383 | 0.00152041 | 0.00290598 | 0.005665609 | 0.003577841 | 0.049956273 | 3.337977305 |
| PDZ and LIM domain protein 7 | PDLIM7 | 0 | 5.5057E-05 | 6.7142E-05 | 0 | 0 | 0 | NaN | NaN |
| Probable tRNA(His) guanylyltransferase | THG1L | 0 | 0.00032996 | 0.00072289 | 0.00109488 | 0.002793158 | 0.001253899 | 0.104531985 | 1.360319785 |
| Gem-associated protein 8 | GEMIN8 | 0.000169645 | 0.00040248 | 0.00128121 | 0.00135623 | 0.002310041 | 0.001972739 | 0.111822769 | 2.293207511 |
| H/ACA ribonucleoprotein complex subunit 2 | NHP2 | 0.000754843 | 0.00104179 | 0.00275459 | 0.0045368 | 0.003385034 | 0.00493136 | 0.075588897 | 2.122611747 |
| Cell growth-regulating nucleolar protein | LYAR | 0.0022928 | 0.00290663 | 0.00594337 | 0.01223136 | 0.018742047 | 0.014397624 | 0.03541741 | 2.308413393 |
| | MROH2A | 6.16896E-05 | 6.4492E-05 | 4.6865E-05 | 0 | 0 | 0 | NaN | NaN |
| CDKN2A-interacting protein | CDKN2AIP | 0.000192364 | 0.00016669 | 0.00083461 | 0.00147546 | 0.004229116 | 0.002538214 | 0.049847866 | 3.721902974 |
| Telomeric repeat-binding factor 2-interacting protein 1 | TERF2IP | 0.000581122 | 0.00081142 | 0.00165723 | 0.00088346 | 0.001452566 | 0.000953052 | 0.6971064 | 0.232106069 |
| Cyclin-dependent kinase inhibitor 2A | CDKN2A | 0 | 0.00040685 | 0.00034841 | 0 | 0 | 0 | NaN | NaN |
| Constitutive coactivator of PPAR-gamma-like protein 1 | FAM120A | 0.001665099 | 0.00193311 | 0.00449995 | 0.00307735 | 0.00978677 | 0.005932431 | 0.16170704 | 1.617699682 |
| Insulin-like growth factor 2 mRNA-binding protein 1 | IGF2BP1 | 0.017671033 | 0.01892599 | 0.04328614 | 0.03985809 | 0.102262225 | 0.076732548 | 0.096589948 | 2.019469742 |
| 26S proteasome non-ATPase regulatory subunit 11 | PSMD11 | 0.366364349 | 0.37153774 | 0.38395287 | 0.0628533 | 0.06956644 | 0.065424983 | 0.025157691 | -2.505595203 |
| Calmodulin-like protein 5 | CALML5 | 0.004487403 | 0.01579085 | 0.00849155 | 0.00325528 | 0.006455311 | 0.008408334 | 0.420218416 | -0.395541657 |
| PRKC apoptosis WT1 regulator protein | PAWR | 0.000484683 | 0.00050179 | 0.00060669 | 0 | 0 | 0 | NaN | NaN |
| Akirin-1 | AKIRIN1 | 0.000991378 | 0.00082212 | 0.00093442 | 0 | 0 | 0 | NaN | NaN |
| Thyroid transcription factor 1-associated protein 26 | CCDC59 | 0.000527864 | 0.0003328 | 0.00059226 | 0.00094998 | 0.001823844 | 0.000409113 | 0.245512228 | 0.847724788 |
| E3 ubiquitin-protein ligase RNF181 | RNF181 | 0.029438473 | 0.02628703 | 0.02907297 | 0.01450841 | 0.020586384 | 0.021487684 | 0.287671713 | -0.497988265 |
| Protein RCC2 | RCC2 | 0.000937073 | 0.00090608 | 0.00169504 | 0.00030677 | 0.002414781 | 0.001006348 | 0.756054013 | 0.102895936 |
| Disco-interacting protein 2 homolog B | DIP2B | 0.000192095 | 0.0001471 | 0.00023694 | 0.00011326 | 0.000287151 | 0.000164079 | 0.846621217 | -0.227429364 |
| Leucine--tRNA ligase, cytoplasmic | LARS | 0.003229929 | 0.00367958 | 0.00735044 | 0.00439295 | 0.01081716 | 0.007683575 | 0.260558976 | 1.062237796 |
| Ataxin-10 | ATXN10 | 0.00132208 | 0.00114121 | 0.00211444 | 0.00058766 | 0.00152208 | 0.000649927 | 0.196360109 | -1.02445989 |
| 26S proteasome non-ATPase regulatory subunit 4 | PSMD4 | 0.000511874 | 0.00055145 | 0.00045258 | 0 | 0 | 0 | NaN | NaN |
| ATP-binding cassette sub-family F member 2 | ABCF2 | 0.00544799 | 0.00477881 | 0.01209243 | 0.01294845 | 0.027020801 | 0.016683672 | 0.088608271 | 1.614640894 |
| Zinc finger CCCH domain-containing protein 7B | ZC3H7B | 0 | 0 | 0 | 0 | 0 | 0 | NaN | NaN |
| Signal recognition particle subunit SRP68 | SRP68 | 0.001270462 | 0.00120381 | 0.00296045 | 0.00320368 | 0.006695196 | 0.004642002 | 0.079182513 | 1.86939392 |
| HAUS augmin-like complex subunit 6 | HAUS6 | 0.000325859 | 0 | 0.00036496 | 0 | 0 | 0 | NaN | NaN |
| Ubiquilin-2 | UBQLN2 | 0.000920726 | 0.00229174 | 0.0024858 | 0.0005655 | 0.000924445 | 0.000920572 | 0.119721119 | -1.31584154 |
| Probable ATP-dependent RNA helicase DDX20 | DDX20 | 0.00034336 | 0.00026645 | 0.00079774 | 0.00053635 | 0.001501989 | 0.001294908 | 0.152286137 | 1.915055715 |
| Actin-related protein 2/3 complex subunit 1A | ARPC1A | 0.000519298 | 0.0004431 | 0.00044301 | 0 | 0 | 0 | NaN | NaN |
| Prefoldin subunit 2 | PFDN2 | 0.000969208 | 0.00109243 | 0.00201005 | 0.00063063 | 0 | 0.000897089 | 0.242788542 | -0.538450256 |
| Poly(U)-binding-splicing factor PUF60 | PUF60 | 0.005394077 | 0.00510866 | 0.00676126 | 0.00702048 | 0.008583196 | 0.005769448 | 0.591267776 | 0.380193762 |
| Translation initiation factor eIF-2B subunit delta | EIF2B4 | 0.00115575 | 0.00162106 | 0.00331775 | 0.00197463 | 0.005767938 | 0.003512024 | 0.235919005 | 1.115364324 |
| E3 ISG15--protein ligase HERC5 | HERC5 | 9.27668E-05 | 0.00021462 | 0.00071765 | 0.00017023 | 0.001196327 | 0.000776283 | 0.384711843 | 1.854771282 |
| tRNA (adenine(58)-N(1))-methyltransferase non-catalytic subunit TRM6 | TRMT6 | 0.000903426 | 0.00113169 | 0.00175559 | 0.00268527 | 0.009472147 | 0.006626619 | 0.048683703 | 2.549799253 |
| Cell division cycle protein 23 homolog | CDC23 | 0.000743086 | 0.00068521 | 0.00087455 | 0.00047252 | 0.000760919 | 0.000255944 | 0.213778412 | -0.65316906 |
| Anaphase-promoting complex subunit 7 | ANAPC7 | 8.42459E-05 | 6.5794E-05 | 9.7854E-05 | 0 | 7.84961E-05 | 0 | 0.888562112 | -0.101985499 |
| Lariat debranching enzyme | DBR1 | 0.001636883 | 0.00183534 | 0.00276747 | 0.00193324 | 0.003156668 | 0.002766624 | 0.5408381 | 0.592079237 |
| mRNA turnover protein 4 homolog | MRTO4 | 0.00434733 | 0.00477996 | 0.00819326 | 0.00519024 | 0.01416804 | 0.010464804 | 0.237108248 | 1.130473633 |
| General transcription factor 3C polypeptide 4 | GTF3C4 | 0.000141972 | 0.00025478 | 0.00048603 | 8.4224E-05 | 4.83961E-05 | 3.84383E-05 | 0.046376411 | -2.396283672 |
| Protein kinase C and casein kinase substrate in neurons protein 3 | PACSIN3 | 0.000638226 | 0.00028699 | 0.00078684 | 0.00135258 | 0.003753097 | 0.002073411 | 0.057339284 | 1.699866834 |
| Protein argonaute-2 | AGO2 | 0.000524516 | 0.00038163 | 0.00074596 | 0.0006503 | 0.001274132 | 0.000712199 | 0.256339639 | 0.441293628 |
| Arpin | ARPIN | 0 | 0.00014907 | 0.00016676 | 0 | 0 | 0 | NaN | NaN |
| Paraneoplastic antigen Ma2 | PNMA2 | 0.000557099 | 0.00341798 | 0.00654389 | 0.00152126 | 0.001515919 | 0.001692474 | 0.638485666 | -1.167878897 |
| Serine/threonine-protein kinase TAO2 | TAOK2 | 0 | 0 | 0 | 0 | 0 | 0 | NaN | NaN |
| Pre-mRNA-splicing factor ISY1 homolog | ISY1 | 0 | 0.00010797 | 0.00030456 | 0.00037196 | 0.000552457 | 0.000458086 | 0.120194389 | 1.33697381 |
| Destrin | DSTN | 0.030259209 | 0.02260023 | 0.02712485 | 0 | 0.000235759 | 0 | 0.003112254 | -6.846158609 |
| RNA-binding protein NOB1 | NOB1 | 0.005293304 | 0.00509275 | 0.00985113 | 0.00399957 | 0.009880527 | 0.005120047 | 0.813667592 | -0.048011458 |
| Peptide-N(4)-(N-acetyl-beta-glucosaminyl)asparagine amidase | NGLY1 | 5.05806E-05 | 0 | 5.6312E-05 | 0 | 0 | 0 | NaN | NaN |
| Pre-mRNA-processing factor 19 | PRPF19 | 0.007330646 | 0.00679598 | 0.00961732 | 0.00981057 | 0.02053013 | 0.016169798 | 0.161203432 | 1.141289412 |
| Ataxin-2 | ATXN2 | 0 | 0 | 0 | 9.1066E-05 | 0.000115761 | 7.49547E-05 | NaN | NaN |
| LINE-1 retrotransposable element ORF1 protein | L1RE1 | 0.000132572 | 0.0001075 | 0.00040484 | 0.0002102 | 0.000671596 | 0.000444202 | 0.219808411 | 1.744439787 |
| Ras GTPase-activating protein-binding protein 2 | G3BP2 | 0.002830646 | 0.00268188 | 0.00688839 | 0.00552986 | 0.01375149 | 0.009136209 | 0.1293556 | 1.690464317 |
| E3 ubiquitin-protein ligase CHIP | STUB1 | 0.000726626 | 0.00041381 | 0.00132405 | 0.00042403 | 0.001137153 | 0.000994041 | 0.902872061 | 0.452092369 |
| Protein kinase C and casein kinase substrate in neurons protein 2 | PACSIN2 | 0.001400851 | 0.0012797 | 0.00256407 | 0.0007714 | 0.001022301 | 0.000747269 | 0.145892969 | -0.860748722 |
| Wings apart-like protein homolog | WAPAL | 0 | 0 | 0 | 0 | 0.000426648 | 0.000472534 | NaN | NaN |
| FAS-associated factor 1 | FAF1 | 0.001305061 | 0.00078146 | 0.001726 | 0.00043655 | 0.000748088 | 0.000228858 | 0.097277606 | -1.579888521 |
| Probable dimethyladenosine transferase | DIMT1 | 0.000714534 | 0.0009558 | 0.00286647 | 0.00209945 | 0.0063495 | 0.006371531 | 0.118867615 | 2.731856279 |
| Charged multivesicular body protein 5 | CHMP5 | 0.000198847 | 0.00019142 | 0 | 0 | 0 | 0.016256255 | 0.004548267 | 6.380666016 |
| Serine/arginine repetitive matrix protein 2 | SRRM2 | 0 | 6.7588E-05 | 9.2529E-05 | 3.8316E-05 | 0.000255555 | 0.000155697 | 0.64962733 | 0.977330745 |
| Proliferation-associated protein 2G4 | PA2G4 | 0.074454148 | 0.07318748 | 0.1426523 | 0.1521045 | 0.346486117 | 0.286301236 | 0.079785428 | 1.943109767 |
| Structural maintenance of chromosomes protein 3 | SMC3 | 0.001608331 | 0.00243965 | 0.00492365 | 0.00655615 | 0.014883272 | 0.009219355 | 0.064798343 | 1.917990449 |
| UPF0568 protein C14orf166 | C14orf166 | 0.006369388 | 0.00765937 | 0.01647639 | 0.02328764 | 0.040142779 | 0.029079934 | 0.055944798 | 1.924725604 |
| RuvB-like 2 | RUVBL2 | 0.030871123 | 0.02262221 | 0.0334131 | 0.02248025 | 0.028721823 | 0.018746763 | 0.572498055 | -0.457600206 |
| Proteasome maturation protein | POMP | 0.04669018 | 0.0405862 | 0.03656316 | 0.01724238 | 0.014552444 | 0.013483595 | 0.076486162 | -1.479727911 |
| RuvB-like 1 | RUVBL1 | 0.020391893 | 0.02150633 | 0.02551466 | 0.01422781 | 0.022817803 | 0.015925226 | 0.502054177 | -0.433447708 |

| | | | | | | | | | |
|---|---|---|---|---|---|---|---|---|---|
| Nuclear migration protein nudC | NUDC | 0.00197755 | 0.00191668 | 0.00316787 | 0.00111165 | 0.002017519 | 0.000988453 | 0.193052133 | -0.831012839 |
| Cofilin-2 | CFL2 | 0.025300078 | 0.00737119 | 0.01242762 | 0 | 0.000710759 | 0.000478804 | 0.015308867 | -4.413008274 |
| Phenylalanine--tRNA ligase alpha subunit | FARSA | 0.004500224 | 0.00684759 | 0.0158261 | 0.01030963 | 0.031850632 | 0.016538993 | 0.192220413 | 1.272203801 |
| Developmentally-regulated GTP-binding protein 1 | DRG1 | 0.00171487 | 0.0020492 | 0.00549201 | 0.0029105 | 0.007127952 | 0.004606562 | 0.289419545 | 1.168630338 |
| Nck-associated protein 1 | NCKAP1 | 0.000622103 | 0.00078355 | 0.00181158 | 0.00025417 | 0.000348026 | 0.000184584 | 0.049412619 | -1.624218084 |
| Exosome complex exonuclease RRP44 | DIS3 | 0.000833221 | 0.0010738 | 0.00160243 | 0.00133 | 0.004752548 | 0.002886379 | 0.139022443 | 1.426536781 |
| RNA 3-terminal phosphate cyclase-like protein | RCL1 | 0 | 5.3212E-05 | 0.00020661 | 0.00011348 | 0.001347424 | 0.001104761 | 0.22826983 | 3.397294022 |
| Probable ATP-dependent RNA helicase DDX52 | DDX52 | 0.00023564 | 0 | 0.00028165 | 0.00056255 | 0.001072541 | 0.000163861 | 0.457602698 | 1.126745062 |
| Translation machinery-associated protein 7 | TMA7 | 0.011217109 | 0.01470847 | 0.02946844 | 0.01771542 | 0.065563011 | 0.04149742 | 0.198104331 | 1.496374559 |
| Nucleolar protein 58 | NOP58 | 0.000204165 | 0.00026773 | 0.00077348 | 0.00071069 | 0.001793708 | 0.000713465 | 0.143015709 | 1.414080017 |
| U6 snRNA-associated Sm-like protein LSm2 | LSM2 | 0.000481463 | 0.00088383 | 0.00207535 | 0.00049343 | 0.002219633 | 0 | 0.920311857 | 0.24377749 |
| Splicing factor 3B subunit 6 | SF3B6 | 0.001491714 | 0.00176112 | 0.00156845 | 0.00494522 | 0.002959912 | 0.001584111 | 0.186904082 | 0.916211066 |
| Nucleolar protein 16 | NOP16 | 0.001040925 | 0.00096587 | 0.00055235 | 0.00523346 | 0.011492479 | 0.006826989 | 0.015857461 | 2.821341029 |
| Serine-threonine kinase receptor-associated protein | STRAP | 0.007677192 | 0.00443588 | 0.0118988 | 0.00255335 | 0.008446311 | 0.005150813 | 0.383179581 | -0.575778565 |
| tRNA-splicing ligase RtcB homolog | RTCB | 0.003657933 | 0.00449388 | 0.0074531 | 0.00626824 | 0.01451762 | 0.011072924 | 0.140620604 | 1.30100402 |
| F-box only protein 7 | FBXO7 | 0.006048035 | 0.00938951 | 0.00727847 | 0.00206214 | 0.004722144 | 0.00512102 | 0.139756251 | -0.624193773 |
| Nucleolar complex protein 2 homolog | NOC2L | 0.000366017 | 0.00035827 | 0.00090288 | 0.00203236 | 0.003953336 | 0.003331516 | 0.025515318 | 3.18619626 |
| 60S ribosomal protein L36 | RPL36 | 0.057703505 | 0.03544436 | 0.06013152 | 0.03079288 | 0.087416456 | 0.041033979 | 0.928348547 | -0.491839896 |
| Deoxynucleoside triphosphate triphosphohydrolase SAMHD1 | SAMHD1 | 0.000555263 | 0.00101829 | 0.0030685 | 0.00037064 | 0.001538019 | 0.000552741 | 0.429289828 | -0.881468702 |
| HBS1-like protein | HBS1L | 0.000558946 | 0.00056758 | 0.00107444 | 0.0009958 | 0.001629097 | 0.00246714 | 0.110420893 | 1.521174307 |
| Talin-1 | TLN1 | 6.7484E-05 | 2.4447E-05 | 6.3979E-05 | 4.8065E-05 | 4.88461E-05 | 3.35527E-05 | 0.80238431 | -0.412608723 |
| ATR-interacting protein | ATRIP | 0 | 0 | 0 | 0 | 0.001448413 | 0.00156567 | NaN | NaN |
| Probable RNA-binding protein 19 | RBM19 | 0 | 0 | 0 | 0 | 0 | 0 | NaN | NaN |
| Myotubularin-related protein 5 | SBF1 | 0 | 0 | 0 | 0 | 6.36691E-05 | 6.73508E-05 | NaN | NaN |
| Ribosomal biogenesis protein LAS1L | LAS1L | 0.000329403 | 0 | 8.4779E-05 | 0.00033019 | 0.000384391 | 0.000144529 | 0.519282312 | 0.982494227 |
| E3 ubiquitin-protein ligase ARIH1 | ARIH1 | 0.000635315 | 0.00073696 | 0.00092511 | 0.00068986 | 0.001281017 | 0.00112879 | 0.467572424 | 0.615114976 |
| U6 snRNA-associated Sm-like protein LSm4 | LSM4 | 0.001503863 | 0.00175243 | 0.0033498 | 0.00358168 | 0.005931879 | 0.001556051 | 0.385720677 | 1.031276337 |
| FACT complex subunit SPT16 | SUPT16H | 6.71761E-05 | 8.8825E-05 | 0.00034413 | 1.915E-05 | 2.43795E-05 | 8.85542E-05 | 0.157485634 | -1.865305367 |
| Protein FAM107B | FAM107B | 0.000584817 | 0.00053383 | 0.00055399 | 0 | 0 | 0 | NaN | NaN |
| CDKN2AIP N-terminal-like protein | CDKN2AIPNL | 0 | 0 | 0 | 0.00055781 | 0.000566119 | 0 | NaN | NaN |
| Nucleolar and coiled-body phosphoprotein 1 | NOLC1 | 0 | 0.00335015 | 0.00325263 | 0 | 0 | 0 | NaN | NaN |
| Serine/threonine-protein kinase MRCK beta | CDC42BPB | 4.63358E-05 | 3.9889E-05 | 8.8946E-05 | 0 | 3.99325E-05 | 0 | 0.26510075 | -0.214563885 |
| RNA-binding protein 8A | RBM8A | 0.002931475 | 0.00132388 | 0.01197278 | 0.00061072 | 0.003392668 | 0.001560277 | 0.323575882 | -0.909824576 |
| Ubiquitin carboxyl-terminal hydrolase 16 | USP16 | 0.000811555 | 0.00077554 | 0.00138223 | 0.00173972 | 0.002260618 | 0.001332528 | 0.176977251 | 1.100093524 |
| Coatomer subunit gamma-1 | COPG1 | 0.000147615 | 0.00042807 | 0.00026292 | 0 | 0.000234406 | 0.000145532 | 0.501273784 | -0.509445448 |
| Cilia- and flagella-associated protein 20 | CFAP20 | 0.000386916 | 0.00029781 | 0.00027304 | 0.0001302 | 0.000601264 | 0.000588083 | 0.824981534 | 0.9816179 |
| Wiskott-Aldrich syndrome protein family member 2 | WASF2 | 0.000239962 | 0.0003324 | 0.00055141 | 5.8341E-05 | 0.000258743 | 0.000137118 | 0.136410963 | -1.27749373 |
| | PSMC3 | 0.339430075 | 0.31323145 | 0.24571194 | 0.10542493 | 0.048481804 | 0.039692338 | 0.029543906 | -2.691713768 |

### Table S3. >2x fold decreased DAPs (Glucose-free / Normal)

| Protein names | Gene names | Normal #1 | Normal #2 | Normal #3 | glucose-free #1 | glucose-free #2 | glucose-free #3 | p-value | Log2(Glucose-free/Normal) |
|---|---|---|---|---|---|---|---|---|---|
| Protein AAR2 homolog | AAR2 | 0.00087835 | 0.00063191 | 0.00123573 | 0.000183734 | 0.000397048 | 0.000221653 | 0.04819 | -1.98648483 |
| BRISC complex subunit Abro1 | ABRAXAS2 | 0.00035655 | 0.00025021 | 0.00034831 | 0 | 0 | 8.07322E-05 | 0.02018 | -2.109176492 |
| Low molecular weight phosphotyrosine protein phosphatase | ACP1 | 0.00878905 | 0.01327227 | 0.01743356 | 7.93474E-05 | 0.000516709 | 0.000453841 | 0.01707 | -4.870084382 |
| Actin, cytoplasmic 1;Actin, cytoplasmic 1, N-terminally processed;Actin, cytoplasmic 2;Actin, cytoplasmic 2, N-terminally processed | ACTB;ACTG1 | 1.29033703 | 1.04259395 | 1.65092703 | 0.048812557 | 0.069677609 | 0.049048778 | 0.00911 | -4.717386934 |
| Actin, gamma-enteric smooth muscle;Actin, alpha skeletal muscle;Actin, alpha cardiac muscle 1;Actin, aortic smooth muscle | ACTG2;ACTA1;ACTC1;ACTA2 | 0.00118441 | 0.00309986 | 0.00305526 | 0.000215739 | 0.000152744 | 5.62204E-05 | 0.01962 | -4.322105653 |
| Alpha-actinin-1 | ACTN1 | 0.0048648 | 0.00155941 | 0.00227053 | 2.09525E-05 | 0 | 0 | 0.00499 | -6.759760989 |
| Alpha-actinin-4 | ACTN4 | 0.13043892 | 0.08750236 | 0.1303498 | 0.00112627 | 0.00119176 | 0.000382339 | 0.00627 | -6.854691878 |
| Proteasomal ubiquitin receptor ADRM1 | ADRM1 | 0.23783451 | 0.22854601 | 0.31228423 | 0.078226093 | 0.063276677 | 0.087066498 | 0.04858 | -1.604236244 |
| Fructose-bisphosphate aldolase C | ALDOC | 0.00011653 | 0.00019531 | 0.00041626 | 0 | 3.40325E-05 | 0 | 0.02592 | -2.520780779 |
| Ankyrin repeat domain-containing protein 36B;Ankyrin repeat domain-containing protein 36A | ANKRD36B;ANKRD36 | 0.00012258 | 0.00014592 | 0.0002511 | 4.73451E-05 | 2.34446E-05 | 0 | 0.03804 | -2.130899997 |
| Annexin A11 | ANXA11 | 0.00283154 | 0.00272909 | 0.00371943 | 4.34865E-05 | 0.000195711 | 0 | 0.01702 | -4.93983385 |
| Annexin A2;Annexin;Putative annexin A2-like protein | ANXA2;ANXA2P2 | 0.00938865 | 0.00860191 | 0.01565166 | 0.00056178 | 0.002597072 | 0.001488911 | 0.03358 | -2.656659593 |
| Annexin A5;Annexin | ANXA5 | 0.00242067 | 0.0046442 | 0.00519098 | 0.001063839 | 0.00087825 | 0.000878123 | 0.03863 | -2.402725283 |
| Annexin A6;Annexin | ANXA6 | 0.04225675 | 0.02096724 | 0.0329747 | 0.001745525 | 0.003272123 | 0.001751572 | 0.01217 | -4.23463738 |
| Annexin A7 | ANXA7 | 0.00084761 | 0.00139611 | 0.0035933 | 6.67021E-05 | 0 | 0 | 0.01236 | -4.38753249 |
| Large proline-rich protein BAG6 | BAG6 | 0.00276766 | 0.00260567 | 0.00295479 | 0.00020157 | 9.13502E-05 | 0.000330309 | 0.01303 | -3.779315445 |
|  | C8G | 0.0021726 | 0 | 0 | 0.000561114 | 0.000680261 | 0.000233473 | 0.02748 | -1.953057433 |
| Calmodulin | CALM2;CALM1;CALM3 | 0.04391054 | 0.05237831 | 0.07261211 | 0.011531473 | 0.019363523 | 0.01597527 | 0.04647 | -1.713129197 |
| Adenylyl cyclase-associated protein 1 | CAP1 | 0.00102144 | 0.00059767 | 0.00079865 | 1.7352E-05 | 4.96364E-05 | 0 | 0.01247 | -4.766237656 |
| Parafibromin | CDC73 | 0.00102379 | 0.00080995 | 0.00128569 | 7.32387E-05 | 0.000165227 | 0.000278064 | 0.02939 | -2.631404092 |
| Cofilin-1 | CFL1 | 0.71850857 | 0.62425416 | 0.64176637 | 0.009357308 | 0.014627449 | 0.024500049 | 0.00755 | -5.4552981 |
| Cofilin-2 | CFL2 | 0.02530008 | 0.00737119 | 0.01242762 | 0 | 0.000710759 | 0.000478804 | 0.01531 | -4.413008274 |
| Creatine kinase U-type, mitochondrial | CKMT1A;CKMT1B | 0.00027185 | 0.00023555 | 0.00041581 | 0 | 5.26018E-05 | 0 | 0.01339 | -2.369632062 |
| Methylosome subunit pICln | CLNS1A | 0 | 0.0005046 | 0.00092821 | 0 | 0.000153132 | 0 | 0.02624 | -2.160015113 |
| Clathrin light chain A | CLTA | 0.09121599 | 0.0594138 | 0.10683167 | 0.004231898 | 0.005837318 | 0.003457502 | 0.01054 | -4.429909972 |
| Clathrin light chain B | CLTB | 0.09126638 | 0.05253638 | 0.09923281 | 0.003780573 | 0.006876415 | 0.002485055 | 0.01069 | -4.593406629 |
| Clathrin heavy chain;Clathrin heavy chain 1 | CLTC | 0.03196954 | 0.01768764 | 0.02700703 | 0.000861227 | 0.003311233 | 0.002574822 | 0.01919 | -3.390790548 |
| CCR4-NOT transcription complex subunit 2 | CNOT2 | 0 | 0.00046476 | 0.000706 | 0 | 0.000195028 | 0 | 0.04871 | -1.554404562 |
| Procollagen galactosyltransferase 1 | COLGALT1 | 0.00010783 | 0.00027579 | 0.0001723 | 0 | 0 | 5.23824E-05 | 0.0329 | -1.717760262 |
| Coronin-1B | CORO1B | 0.00859646 | 0.00945504 | 0.00827765 | 0.000298242 | 0.00175366 | 0.001299659 | 0.0362 | -2.725609876 |
| Coronin-1C;Coronin | CORO1C | 0.00109876 | 0.00120004 | 0.00134944 | 0 | 8.23E-05 | 5.75484E-05 | 0.01262 | -4.124106965 |
| Copine-3 | CPNE3 | 0.00815586 | 0.00480917 | 0.00973879 | 0 | 0.000437497 | 0.000327057 | 0.01223 | -4.430357851 |
| Cleavage and polyadenylation specificity factor subunit 6 | CPSF6 | 0.00324113 | 0.00436062 | 0.00573596 | 0.000295071 | 0.000659889 | 0.000770285 | 0.01727 | -2.724238 |
| Cytokine receptor-like factor 3 | CRLF3 | 0.00066851 | 0.00094496 | 0.0018187 | 0 | 9.25222E-05 | 0 | 0.01085 | -3.352381786 |
| Src substrate cortactin | CTTN | 0.00917199 | 0.00447409 | 0.01068317 | 0.000719121 | 0.001703969 | 0.000373342 | 0.03194 | -3.67292755 |
| Cullin-2 | CUL2 | 7.0244E-05 | 6.994E-05 | 0.0001284 | 1.58028E-05 | 0 | 0 | 0.01386 | -2.152196851 |
| Protein DDI1 homolog 2 | DDI2 | 0.01702441 | 0.00973307 | 0.01241575 | 0.000562253 | 0.000827621 | 0.000643151 | 0.01053 | -4.270869643 |
| Putative pre-mRNA-splicing factor ATP-dependent RNA helicase DHX16 | DHX16 | 0.00065127 | 0.00089312 | 0.00098347 | 0.00012484 | 0.000230107 | 0.00015035 | 0.02908 | -2.570534008 |
| DnaJ homolog subfamily C member 10 | DNAJC10 | 0.00077875 | 0.00074434 | 0.00071036 | 3.40665E-05 | 4.86546E-05 | 0 | 0.01246 | -4.192428767 |
| Desmoplakin | DSP | 0.00024979 | 0.00040413 | 0.00043812 | 6.99339E-05 | 0.000134635 | 6.99017E-05 | 0.03953 | -2.530773434 |
| Destrin | DSTN | 0.03025921 | 0.02260023 | 0.02712485 | 0 | 0.000235759 | 0 | 0.00311 | -6.846158609 |
| Proteasome-associated protein ECM29 homolog | ECM29;KIAA0368 | 0.04746781 | 0.04580341 | 0.05445155 | 0.00966371 | 0.010508833 | 0.009282446 | 0.02986 | -2.296300394 |
| Epidermal growth factor receptor substrate 15 | EPS15 | 0.00025694 | 0.00011934 | 0.00027983 | 0 | 6.7252E-05 | 0 | 0.03849 | -1.933793656 |
| Extended synaptotagmin-1 | ESYT1 | 3.9643E-05 | 8.3526E-05 | 7.3409E-05 | 0 | 1.99421E-05 | 0 | 0.0298 | -1.88014747 |
| Fatty acid synthase;[Acyl-carrier-protein] S-acetyltransferase;[Acyl-carrier-protein] S-malonyltransferase;3-oxoacyl-[acyl-carrier-protein] synthase;3-oxoacyl-[acyl-carrier-protein] reductase;3-hydroxyacyl-[acyl-carrier-protein] dehydratase;Enoyl-[acyl-carrier-protein] reductase;Oleoyl-[acyl-carrier-protein] hydrolase | FASN | 0.00049712 | 0.00034837 | 0.0010021 | 0.000120099 | 0.000180777 | 9.55058E-05 | 0.03889 | -2.049361302 |
| Filamin-A | FLNA | 0.00661684 | 0.0042282 | 0.00460371 | 0.000261334 | 0.000448922 | 4.53841E-05 | 0.02792 | -4.138829591 |
| Filamin-B | FLNB | 0.00319192 | 0.00221375 | 0.00265805 | 0.000186508 | 0.000310216 | 0.000353753 | 0.01502 | -3.099023783 |
| Filamin-C | FLNC | 0.00052992 | 0.00034975 | 0.00032631 | 0 | 3.04376E-05 | 9.72856E-05 | 0.03713 | -2.68422821 |
| Fascin | FSCN1 | 0.0005152 | 0.00044361 | 0.00097123 | 0 | 8.61762E-05 | 0.000136947 | 0.03231 | -2.245643068 |
| Glutamate dehydrogenase 1, mitochondrial;Glutamate dehydrogenase 2, mitochondrial | GLUD1;GLUD2 | 0.00386155 | 0.00395645 | 0.01221938 | 0.00054188 | 0.000925731 | 0.000385104 | 0.02499 | -2.868162314 |
| Glia maturation factor beta | GMFB | 0.01311611 | 0.00667214 | 0.01003745 | 0.000130818 | 0 | 0.0002037 | 0.00764 | -5.942244894 |
| Growth factor receptor-bound protein 2 | GRB2 | 0.0037105 | 0.00192515 | 0.00328304 | 8.51863E-05 | 0.000395039 | 0.000305384 | 0.02519 | -3.426335946 |
| General transcription factor 3C polypeptide 4 | GTF3C4 | 0.00014197 | 0.00025478 | 0.00048603 | 8.4224E-05 | 4.83961E-05 | 3.84383E-05 | 0.04638 | -2.396283672 |
| HAUS augmin-like complex subunit 4 | HAUS4 | 0.00026204 | 0.00066236 | 0.0003245 | 9.56147E-05 | 0 | 0 | 0.02576 | -1.762936364 |
| Histone deacetylase 2;Histone deacetylase 1 | HDAC2;HDAC1 | 0.00119326 | 0.00118905 | 0.00096283 | 0 | 0.000282544 | 0 | 0.01983 | -2.073263137 |
| Immunoglobulin-binding protein 1 | IGBP1 | 0.00020884 | 0.00012275 | 0 | 4.98952E-05 | 0 | 0 | 0.04259 | -1.682069977 |
| Inosine-5-monophosphate dehydrogenase;Inosine-5-monophosphate dehydrogenase 1;Inosine-5-monophosphate dehydrogenase 2 | IMPDH1;IMPDH2 | 0.01040085 | 0.01070658 | 0.01458398 | 0.00081689 | 0.001039418 | 0.001043034 | 0.01311 | -3.3646504 |
| L-lactate dehydrogenase A chain | LDHA | 0.00889318 | 0.00528713 | 0.0084405 | 0.00155104 | 0.002208784 | 0.001603291 | 0.03832 | -2.396291651 |
| Microtubule-actin cross-linking factor 1, isoforms 1/2/3/5 | MACF1 | 0.00021342 | 0.00014341 | 0.00031408 | 3.11111E-06 | 1.95818E-05 | 1.2666E-05 | 0.02228 | -4.074662491 |
| Myristoylated alanine-rich C-kinase substrate | MARCKS | 0.00357457 | 0.0035727 | 0.00878153 | 0.000380035 | 0.001119728 | 0.000771746 | 0.03179 | -2.211572506 |
| MARCKS-related protein | MARCKSL1 | 0.00524365 | 0.00647702 | 0.01276464 | 0 | 0.001499444 | 0.000512219 | 0.03904 | -2.885700709 |
| Mediator of DNA damage checkpoint protein 1 | MDC1 | 1.9095E-05 | 3.9329E-05 | 5.7919E-05 | 0 | 0 | 8.48992E-06 | 0.03374 | -2.211770639 |
| Myosin-10 | MYH10 | 0.00078591 | 0.0004075 | 0.00086446 | 0.000133548 | 0.000148873 | 0.000136244 | 0.03188 | -2.528178755 |
| Myosin-9 | MYH9 | 0.00208868 | 0.00180624 | 0.00238926 | 0.000117895 | 0.000163378 | 0.000130242 | 0.01136 | -4.003324815 |
| Myosin light polypeptide 6 | MYL6 | 0.01174337 | 0.01348477 | 0.01820349 | 0.001833898 | 0.002907408 | 0.002208548 | 0.02177 | -2.610160715 |
| Unconventional myosin-XVIIIa | MYO18A | 0.00013931 | 8.443E-05 | 0.00025975 | 0 | 8.63891E-06 | 0 | 0.01018 | -4.011274649 |
| Condensin complex subunit 1 | NCAPD2 | 0.0003884 | 9.9119E-05 | 0.000387 | 0 | 1.52878E-05 | 0 | 0.01678 | -4.661892035 |
| Nck-associated protein 1 | NCKAP1 | 0.0006221 | 0.00078355 | 0.00181158 | 0.000254174 | 0.000348026 | 0.000184584 | 0.04941 | -1.624218084 |
| Neurofilament light polypeptide | NEFL | 0.01650711 | 0.02975714 | 0.05493652 | 0.005028114 | 0.003845247 | 0.004517379 | 0.02101 | -2.719678204 |
| Neurofilament medium polypeptide | NEFM | 0.04053242 | 0.05718727 | 0.12567358 | 0.013173144 | 0.011391891 | 0.010742284 | 0.02539 | -2.327686814 |
| 5-nucleotidase domain-containing protein 2 | NT5DC2 | 0.00047291 | 0.00080722 | 0.00091698 | 0 | 0.000162227 | 0 | 0.01773 | -2.314953329 |
| Cleavage and polyadenylation specificity factor subunit 5 | NUDT21 | 0.00490958 | 0.00463163 | 0.00593141 | 0.000830608 | 0.000626619 | 0.000915471 | 0.02082 | -2.563361409 |
| Nucleoporin Nup37 | NUP37 | 0.00060984 | 0.00048562 | 0.00109005 | 0 | 7.24676E-05 | 0 | 0.00902 | -3.073027492 |
| Prolyl 4-hydroxylase subunit alpha-1 | P4HA1 | 0.00037394 | 0.00047264 | 0.000638 | 3.84486E-05 | 0.000154498 | 4.86379E-05 | 0.03421 | -3.280579729 |
| Protein disulfide-isomerase A6 | PDIA6 | 0.00133188 | 0.00151753 | 0.00210037 | 0.000240305 | 0.000581709 | 0.000435381 | 0.03881 | -1.801378485 |
| ATP-dependent 6-phosphofructokinase, liver type | PFKL | 0.01521162 | 0.01806867 | 0.01838615 | 0.00124281 | 0.002575106 | 0.000917204 | 0.01323 | -3.861812656 |
| ATP-dependent 6-phosphofructokinase, muscle type | PFKM | 0.00404664 | 0.00210196 | 0.00309855 | 0 | 0.000106894 | 6.3616E-05 | 0.00901 | -5.231699191 |
| PITH domain-containing protein 1 | PITHD1 | 0.00108874 | 0.00139569 | 0.00220431 | 0 | 0.000202395 | 0.000113268 | 0.01806 | -3.204446538 |
| Phospholipase A-2-activating protein | PLAA | 0.00022933 | 0.0003223 | 0.00054021 | 3.23786E-05 | 0 | 0 | 0.00884 | -3.315278247 |
| Procollagen-lysine,2-oxoglutarate 5-dioxygenase 2 | PLOD2 | 0.0003185 | 0.00025117 | 0.00051014 | 0 | 5.13186E-05 | 0 | 0.01174 | -2.633753243 |
| Plastin-3 | PLS3 | 0.01966241 | 0.00434021 | 0.00589095 | 0.000244821 | 0.000824955 | 0.000481005 | 0.026 | -3.614376267 |
| Liprin-alpha-1;Liprin-alpha-2 | PPFIA1;PPFIA2 | 0.25867764 | 0.27195645 | 0.33239565 | 0.056025372 | 0.06670417 | 0.074686009 | 0.0354 | -2.027526799 |
| Protein phosphatase 1 regulatory subunit 12A | PPP1R12A | 0.000862 | 0.00094636 | 0.00063716 | 0 | 0.000175259 | 5.62633E-05 | 0.02986 | -3.117807217 |
| Serine/threonine-protein phosphatase 2A 56 kDa regulatory subunit epsilon isoform;Serine/threonine-protein phosphatase 2A 56 kDa regulatory subunit beta isoform | PPP2R5E;PPP2R5B | 0.00036875 | 0.00017734 | 0.00034338 | 0 | 9.69154E-05 | 0 | 0.03421 | -1.825020782 |
| 5-AMP-activated protein kinase subunit gamma-1 | PRKAG1 | 0.00208734 | 0.00259824 | 0.00632651 | 0.000803655 | 0.000356156 | 0.000197157 | 0.03965 | -2.866954054 |
| 26S protease regulatory subunit 4 | PSMC1 | 0.36489195 | 0.35990788 | 0.43208512 | 0.082115788 | 0.096919409 | 0.094341349 | 0.03721 | -1.951507184 |
|  | PSMC1 | 0.00527774 | 0.00435308 | 0.00606768 | 0.000167124 | 0 | 0 | 0.00503 | -4.980929447 |
| 26S protease regulatory subunit 7 | PSMC2 | 0.59534207 | 0.57106668 | 0.59384419 | 0.122754394 | 0.137099691 | 0.131981307 | 0.03464 | -2.169750836 |
|  | PSMC3 | 0.33943008 | 0.31323145 | 0.24571194 | 0.105424931 | 0.048481804 | 0.039692338 | 0.02954 | -2.691713768 |
| 26S protease regulatory subunit 6B | PSMC4 | 0.35651103 | 0.37209254 | 0.40648461 | 0.088462076 | 0.092157887 | 0.078220232 | 0.03532 | -2.072530447 |
| 26S protease regulatory subunit 8 | PSMC5 | 0.50584481 | 0.5201298 | 0.56933053 | 0.109326453 | 0.14114464 | 0.112964658 | 0.03523 | -2.20300022 |
| 26S protease regulatory subunit 10B | PSMC6 | 0.50097973 | 0.34585994 | 0.39378939 | 0.090377896 | 0.090625628 | 0.066079252 | 0.02983 | -2.123382353 |
| 26S proteasome non-ATPase regulatory subunit 1 | PSMD1 | 0.17083753 | 0.22163718 | 0.207745 | 0.047768639 | 0.054967118 | 0.051011586 | 0.04207 | -2.025916893 |
|  | PSMD1 | 0.00154199 | 0.00096317 | 0.00075591 | 0.000147181 | 8.91844E-05 | 0 | 0.01917 | -3.071569086 |
| 26S proteasome non-ATPase regulatory subunit 11 | PSMD11 | 0.36636435 | 0.37153774 | 0.38395287 | 0.062853303 | 0.06956644 | 0.065424983 | 0.02516 | -2.505595203 |
| 26S proteasome non-ATPase regulatory subunit 12 | PSMD12 | 0.24156869 | 0.26513137 | 0.25852589 | 0.056061925 | 0.066153681 | 0.052873138 | 0.03533 | -2.205215595 |
| 26S proteasome non-ATPase regulatory subunit 13 | PSMD13 | 0.36352592 | 0.41625667 | 0.48323135 | 0.068280385 | 0.089973346 | 0.078940707 | 0.02722 | -2.398632055 |
| 26S proteasome non-ATPase regulatory subunit 14 | PSMD14 | 0.09386407 | 0.11447713 | 0.14392182 | 0.013080686 | 0.02105383 | 0.02180703 | 0.02154 | -2.442904774 |
| 26S proteasome non-ATPase regulatory subunit 2 | PSMD2 | 0.21715373 | 0.18332461 | 0.22157275 | 0.043262915 | 0.055498855 | 0.051644436 | 0.03863 | -2.072031929 |
| 26S proteasome non-ATPase regulatory subunit 3 | PSMD3 | 0.28332214 | 0.37834188 | 0.41567266 | 0.058794818 | 0.059725961 | 0.072964658 | 0.02575 | -2.663260423 |
| 26S proteasome non-ATPase regulatory subunit 4 | PSMD4 | 0.21685142 | 0.17166335 | 0.1536122 | 0.055465248 | 0.041215628 | 0.05900886 | 0.04822 | -1.629926042 |
| 26S proteasome non-ATPase regulatory subunit 6 | PSMD6 | 0.28879185 | 0.27408144 | 0.38270162 | 0.069482342 | 0.065257631 | 0.066342128 | 0.03228 | -2.122032829 |
| 26S proteasome non-ATPase regulatory subunit 7 | PSMD7 | 0.20884 | 0.21952266 | 0.25016897 | 0.062277052 | 0.063943692 | 0.064638302 | 0.04737 | -1.779495937 |

| | | | | | | | | | |
|---|---|---|---|---|---|---|---|---|---|
| 26S proteasome non-ATPase regulatory subunit 8 | PSMD8 | 0.27044004 | 0.27629017 | 0.2692666 | 0.047377305 | 0.055778787 | 0.05661182 | 0.02907 | -2.2775203 |
| Ras-related protein Rab-1A;Ras-related protein Rab-1B;Putative Ras | RAB1A;RAB1B;RAB1C | 0.00393136 | 0.00561175 | 0.00603078 | 0.00054288 | 0.001714282 | 0.000864706 | 0.02835 | -2.698167849 |
| Ras-related protein Rab-7a | RAB7A | 0.00244463 | 0.00170972 | 0.00196018 | 0.000573241 | 0.000465343 | 0.000396553 | 0.03639 | -2.074618994 |
| UV excision repair protein RAD23 homolog A | RAD23A | 0.00700817 | 0.00383063 | 0.00658252 | 0.000433586 | 0.000300532 | 0.00034538 | 0.01129 | -4.252383333 |
| UV excision repair protein RAD23 homolog B | RAD23B | 0.17742134 | 0.15117764 | 0.20600055 | 0.013791324 | 0.019822933 | 0.016442411 | 0.01387 | -3.431685787 |
| Receptor expression-enhancing protein 5;Receptor expression-enha | REEP5 | 0.00232275 | 0.00402167 | 0.00324094 | 0.000399591 | 0.000629592 | 0.000511576 | 0.02224 | -2.663389598 |
| E3 ubiquitin-protein ligase BRE1A | RNF20 | 0.00012962 | 0.00024746 | 0.00026662 | 2.59152E-05 | 4.96109E-05 | 6.04829E-05 | 0.03132 | -2.318477969 |
| Ubiquitin-40S ribosomal protein S27a;Ubiquitin;40S ribosomal prot | RPS27A;UBB;UBC;UBA52 | 0.64197738 | 0.80475243 | 0.85935702 | 0.135687792 | 0.211221387 | 0.216726706 | 0.03857 | -1.929789115 |
| Protein transport protein Sec23B | SEC23B | 0.00023962 | 0.00017891 | 0.00015782 | 6.146E-05 | 0 | 0 | 0.03072 | -1.541490164 |
| Sorting nexin-2 | SNX2 | 0.00036475 | 0.00058196 | 0.00063206 | 2.9334E-05 | 8.9889E-05 | 0 | 0.02681 | -3.502498811 |
| Spectrin alpha chain, non-erythrocytic 1 | SPTAN1 | 0.00809148 | 0.00515639 | 0.01049868 | 0.000183218 | 0.000332878 | 0.000471483 | 0.00901 | -4.603337999 |
| Spectrin beta chain, non-erythrocytic 1 | SPTBN1 | 0.00502335 | 0.00359908 | 0.00709407 | 0.00054001 | 0.001095512 | 0.001035829 | 0.02368 | -2.277862794 |
| Spectrin beta chain, non-erythrocytic 2 | SPTBN2 | 0.00099194 | 0.00031566 | 0.00128459 | 0 | 4.66053E-05 | 0 | 0.01587 | -4.411684015 |
| SLIT-ROBO Rho GTPase-activating protein 2 | SRGAP2 | 0.0013999 | 0.00152067 | 0.00148032 | 0.000104715 | 0.000269351 | 0.00030441 | 0.01918 | -2.458347801 |
| Serine/threonine-protein kinase 26 | STK26 | 0.0019275 | 0.00160641 | 0.00234852 | 0.000231898 | 0.000526902 | 0.000492571 | 0.03007 | -1.968327263 |
| Syntaxin-7 | STX7 | 0.00119242 | 0.00045641 | 0.00129007 | 0 | 5.2133E-05 | 0 | 0.0105 | -4.515551786 |
| Transcription elongation factor SPT6 | SUPT6H | 0.00124073 | 0.00107652 | 0.00206384 | 0.000235005 | 0.000393458 | 0.000177905 | 0.02599 | -2.400431656 |
| Transgelin-2 | TAGLN2 | 0.01618744 | 0.01219198 | 0.01417207 | 0.000466409 | 0.000505498 | 0.000508227 | 0.00877 | -4.809201404 |
| TRPM8 channel-associated factor 1 | TCAF1 | 0.00277354 | 0.00256705 | 0.00323235 | 0.00054932 | 0.000750124 | 0.000712823 | 0.03613 | -1.96011378 |
| Transketolase | TKT | 0.000761 | 0.00134125 | 0.00165558 | 0.000225189 | 0.000332436 | 0.000150387 | 0.02733 | -2.574374212 |
| DNA topoisomerase 2-alpha | TOP2A | 0.00096714 | 0.00097293 | 0.00207882 | 5.75649E-05 | 0.000275673 | 0.000101036 | 0.03124 | -3.267466581 |
| Tropomyosin alpha-1 chain | TPM1 | 0.00409209 | 0.00755407 | 0.01291442 | 0 | 0.000416416 | 0.000109432 | 0.02098 | -5.145151588 |
| Tropomyosin3 | TPM3 | 0.07874818 | 0.14531561 | 0.16331172 | 0.003734666 | 0.004719599 | 0.001944212 | 0.00765 | -5.282066632 |
| Tropomyosin alpha-4 chain | TPM4 | 0.02049491 | 0.02594264 | 0.02913508 | 0.000195764 | 0.00036213 | 0 | 0.00701 | -6.606369675 |
| Gamma-tubulin complex component 2 | TUBGCP2 | 0.00030835 | 0.00012757 | 0.00065747 | 2.88738E-05 | 0 | 0 | 0.02632 | -3.416749988 |
| Thioredoxin-like protein 1 | TXNL1 | 0.06021162 | 0.06215116 | 0.06585807 | 0.013248401 | 0.017587495 | 0.014156752 | 0.03734 | -2.134290944 |
| Ubiquitin-protein ligase E3A | UBE3A | 0.00687381 | 0.00703988 | 0.00897991 | 0.000602666 | 0.001189952 | 0.000973479 | 0.01645 | -2.85432969 |
| Ubiquitin-protein ligase E3C | UBE3C | 0.00697738 | 0.00570187 | 0.00625555 | 0.000402967 | 0.00059142 | 0.000207419 | 0.01089 | -3.956402671 |
| Ubiquitin domain-containing protein UBFD1 | UBFD1 | 0.00179851 | 0.00185083 | 0.00270253 | 0 | 0.000179639 | 0.000113547 | 0.01386 | -3.69590695 |
| Ubiquitin-like protein 4A | UBL4A | 0.0025851 | 0.00438794 | 0.00463147 | 0 | 0 | 0.000446617 | 0.0102 | -3.296434056 |
| Ubiquitin-like domain-containing CTD phosphatase 1 | UBLCP1 | 0.00480417 | 0.00351157 | 0.00466079 | 0.00020401 | 0.000282865 | 6.61396E-05 | 0.01289 | -4.513863045 |
| Ubiquilin-1 | UBQLN1 | 0.00753219 | 0.00704219 | 0.00995433 | 0.001283664 | 0.001724729 | 0.001308013 | 0.0252 | -2.525692705 |
| Ubiquitin carboxyl-terminal hydrolase isozyme L5;Ubiquitin carboxy | UCHL5 | 0.1025865 | 0.11475976 | 0.1308704 | 0.01854002 | 0.019555056 | 0.018040697 | 0.02223 | -2.629902163 |
| Ubiquitin carboxyl-terminal hydrolase 14;Ubiquitin carboxyl-termina | USP14 | 0.03412664 | 0.02709201 | 0.0279715 | 0.006043219 | 0.009417768 | 0.006152079 | 0.03789 | -2.184811894 |
| Ubiquitin carboxyl-terminal hydrolase 15 | USP15 | 0.00257838 | 0.00224788 | 0.0027155 | 0.000294512 | 0.000260859 | 0.000291539 | 0.01589 | -3.144703548 |
| Transitional endoplasmic reticulum ATPase | VCP | 0.05238775 | 0.041566 | 0.05162846 | 0.014726657 | 0.018071014 | 0.01863655 | 0.04525 | -1.514489076 |
| Vacuolar protein sorting-associated protein 26A | VPS26A | 0.00274314 | 0.00241191 | 0.00279414 | 0.000305134 | 0.000661577 | 0.000393379 | 0.0223 | -2.801837457 |
| Vacuolar protein sorting-associated protein 29 | VPS29 | 0.01180663 | 0.00716288 | 0.01204128 | 0.001736817 | 0.001717229 | 0.00100072 | 0.01972 | -2.781442762 |

**Table S4. >2x fold increased DAPs (Glucose-free / Normal)**

| Protein names | Gene names | Normal #1 | Normal #2 | Normal #3 | glucose-free #1 | glucose-free #2 | glucose-free #3 | p-value | Log2(Glucose-free/Normal) |
|---|---|---|---|---|---|---|---|---|---|
| Aminoacyl tRNA synthase complex-interacting multifunctional protein 2 | AIMP2 | 0.00091362 | 0.00063953 | 0.001019 | 0.002133097 | 0.004257377 | 0.002942849 | 0.04709 | 1.687555276 |
| Rho guanine nucleotide exchange factor 2 | ARHGEF2 | 0.00029834 | 0.00031563 | 0.00033819 | 0.001507069 | 0.001417072 | 0.003453413 | 0.02228 | 2.2554438 |
| Ribosome biogenesis protein BMS1 homolog | BMS1 | 0.00019943 | 0.00026849 | 7.9539E-05 | 0.000712982 | 0.002075247 | 0.00211917 | 0.03502 | 3.379335939 |
| Ribosome biogenesis protein BOP1 | BOP1 | 0.00015256 | 0.00014646 | 0.00021141 | 0.000807504 | 0.001707719 | 0.000960491 | 0.02077 | 2.654357521 |
| Ribosome biogenesis protein BRX1 homolog | BRIX1 | 0.00050965 | 0.00076181 | 0.00268298 | 0.006244692 | 0.011203975 | 0.009834485 | 0.03875 | 3.690340693 |
| Cell division cycle protein 27 homolog | CDC27 | 4.6313E-05 | 4.1797E-05 | 0.00010509 | 0.000284986 | 0.000385583 | 0.00035979 | 0.02421 | 2.957672201 |
| CDKN2A-interacting protein | CDKN2AIP | 0.00019236 | 0.00016669 | 0.00083461 | 0.001475461 | 0.004229116 | 0.002538214 | 0.04985 | 3.721902974 |
| Charged multivesicular body protein 5 | CHMP5 | 0.00019885 | 0.00019142 | 0 | 0 | 0 | 0.016256255 | 0.00455 | 6.380666016 |
| Cytoskeleton-associated protein 5 | CKAP5 | 0.00047212 | 0.00051138 | 0.00057136 | 0.001365156 | 0.003548439 | 0.002317593 | 0.03565 | 2.18016241 |
| ATP-dependent RNA helicase DDX18 | DDX18 | 0.00030443 | 0.00026053 | 0.00026446 | 0.000894952 | 0.00187608 | 0.001557025 | 0.02994 | 2.557654309 |
| ATP-dependent RNA helicase DDX54 | DDX54 | 0.00051853 | 0.00062707 | 0.00101562 | 0.00220201 | 0.005806646 | 0.004213611 | 0.02664 | 2.748358725 |
| ATP-dependent RNA helicase A | DHX9 | 0.00982197 | 0.01060295 | 0.02046214 | 0.028895339 | 0.070151753 | 0.046441437 | 0.04759 | 2.130946627 |
| Cytoplasmic dynein 2 heavy chain 1 | DYNC2H1 | 3.5478E-05 | 3.9204E-05 | 5.0083E-05 | 0.000530087 | 0.00094684 | 0.000886457 | 0.01052 | 4.498963165 |
| Probable rRNA-processing protein EBP2 | EBNA1BP2 | 0.00226722 | 0.00244572 | 0.00516906 | 0.015426544 | 0.03449994 | 0.028499659 | 0.01818 | 3.542611147 |
| Elongation factor 2 | EEF2 | 0.03457843 | 0.03033707 | 0.05767193 | 0.085614148 | 0.210203453 | 0.158539577 | 0.04641 | 2.196898552 |
| Ribosomal RNA small subunit methyltransferase NEP1 | EMG1 | 0.00077634 | 0.00160327 | 0.0028893 | 0.011154115 | 0.015090877 | 0.014886769 | 0.01978 | 3.214944958 |
| ELKS/Rab6-interacting/CAST family member 1;ERC protein 2 | ERC1;ERC2 | 9.654E-05 | 3.6997E-05 | 5.7084E-05 | 0 | 0.000171803 | 0 | 0.04282 | 1.589601947 |
| Exosome component 10 | EXOSC10 | 0.00288014 | 0.00243306 | 0.00467778 | 0.008689889 | 0.017544635 | 0.011518255 | 0.04378 | 1.999712824 |
| Exosome complex component RRP42 | EXOSC7 | 4.6533E-05 | 0.00019492 | 0.00037002 | 0.002348546 | 0.001523285 | 0.003018596 | 0.03644 | 3.590791614 |
| pre-rRNA processing protein FTSJ3 | FTSJ3 | 0 | 0 | 0.00032632 | 0.00116035 | 0.003879669 | 0.001784305 | 0.02357 | 2.450994084 |
| Nucleolar GTP-binding protein 2 | GNL2 | 7.5048E-05 | 0.00014233 | 6.4291E-05 | 0.000956448 | 0.001753124 | 0.000725246 | 0.01356 | 3.671808694 |
| Nucleolar GTP-binding protein 1 | GTPBP4 | 0.00195796 | 0.00292589 | 0.00489771 | 0.007202064 | 0.019603273 | 0.015669555 | 0.04784 | 2.421018252 |
| KRR1 small subunit processome component homolog | KRR1 | 0.00029263 | 0.00057578 | 0.0009863 | 0.002962533 | 0.006730957 | 0.005561484 | 0.02438 | 3.271887831 |
| Cell growth-regulating nucleolar protein | LYAR | 0.0022928 | 0.00290663 | 0.00594337 | 0.012231361 | 0.018742047 | 0.014397624 | 0.03542 | 2.308413393 |
| MAP7 domain-containing protein 1 | MAP7D1 | 9.5527E-05 | 8.5115E-05 | 0.00030334 | 0.000668892 | 0.000824286 | 0.000805978 | 0.03571 | 3.076763017 |
| M-phase phosphoprotein 6 | MPHOSPH6 | 0.00032433 | 0.00038368 | 0.00070275 | 0.008847283 | 0.015727086 | 0.01113173 | 0.0088 | 4.858626687 |
| Myb-binding protein 1A | MYBBP1A | 0.00090488 | 0.00086635 | 0.00193068 | 0.003852927 | 0.008497877 | 0.005494499 | 0.03039 | 2.602186362 |
| Nucleolin | NCL | 0.09021946 | 0.10094944 | 0.21692392 | 0.386679568 | 0.884464446 | 0.664648038 | 0.03485 | 2.718957661 |
| NF-kappa-B-repressing factor | NKRF | 0.001188 | 0.00125667 | 0.00341027 | 0.005649949 | 0.016158369 | 0.009199299 | 0.04617 | 2.871914636 |
| Nucleolar complex protein 2 homolog | NOC2L | 0.00036602 | 0.00035827 | 0.00090288 | 0.00203236 | 0.003953336 | 0.003331516 | 0.02552 | 3.18619626 |
| Nucleolar complex protein 3 homolog | NOC3L | 0.00018987 | 9.8173E-05 | 0.00048751 | 0.000992593 | 0.003760464 | 0.002387888 | 0.04508 | 3.652674087 |
| Nucleolar protein 7 | NOL7 | 0 | 0.00122412 | 0.00078094 | 0.003341074 | 0.004356223 | 0.005404732 | 0.04424 | 2.155565608 |
| Nucleolar MIF4G domain-containing protein 1 | NOM1 | 1.7791E-05 | 0 | 3.99E-05 | 0.000211213 | 0.000701317 | 0.000294733 | 0.02922 | 3.467561343 |
| Nucleolar protein 16 | NOP16 | 0.00104092 | 0.00096587 | 0.00055235 | 0.005233457 | 0.011492479 | 0.006826989 | 0.01586 | 2.821341029 |
| tRNA (cytosine(34)-C(5))-methyltransferase | NSUN2 | 0.00099468 | 0.0012863 | 0.00360115 | 0.006387249 | 0.011563333 | 0.007142635 | 0.04728 | 2.473232606 |
| p21-activated protein kinase-interacting protein 1 | PAK1IP1 | 9.2414E-05 | 0.00024269 | 0.0002499 | 0.00352384 | 0.006468839 | 0.003903612 | 0.0103 | 4.007634984 |
| 28 kDa heat- and acid-stable phosphoprotein | PDAP1 | 0.00738607 | 0.01207055 | 0.01231985 | 0.021838413 | 0.065758562 | 0.036790965 | 0.04837 | 1.607859625 |
| Programmed cell death protein 4 | PDCD4 | 0.00277668 | 0.00350288 | 0.00670874 | 0.011755093 | 0.026888201 | 0.017769059 | 0.0368 | 2.342754223 |
| Sister chromatid cohesion protein PDS5 homolog A | PDS5A | 3.6194E-05 | 8.4639E-05 | 0.00034757 | 0.001247756 | 0.00255381 | 0.001525207 | 0.03926 | 4.171531847 |
| Sister chromatid cohesion protein PDS5 homolog B | PDS5B | 8.7291E-05 | 9.6419E-05 | 0.00015992 | 0.000354835 | 0.000783086 | 0.000511284 | 0.03154 | 2.406737003 |
| Pescadillo homolog | PES1 | 0 | 0 | 0.00015665 | 0.002089233 | 0.002213873 | 0.001220271 | 0.00832 | 3.737319884 |
| Suppressor of SWI4 1 homolog | PPAN | 0.00180887 | 0.00148508 | 0.0026006 | 0.006180616 | 0.014089016 | 0.009506182 | 0.03047 | 2.39377889 |
| Periodic tryptophan protein 2 homolog | PWP2 | 0.00012687 | 0.00015296 | 0.00038593 | 0.001319357 | 0.002100561 | 0.001130854 | 0.02349 | 3.108636953 |
| RNA-binding protein 28 | RBM28 | 0.00029712 | 0.00035383 | 0.00152041 | 0.002905983 | 0.005665609 | 0.003577841 | 0.04996 | 3.337977305 |
| RNA-binding protein 7 | RBM7 | 0.00077606 | 0.00062613 | 0.00087653 | 0.00256851 | 0.003830514 | 0.002894168 | 0.03951 | 1.898906709 |
| Ribonuclease inhibitor | RNH1 | 2.2496E-05 | 0 | 0 | 0.001309359 | 0.000820268 | 0.001100867 | 0.00413 | 5.612840698 |
| 60S ribosomal protein L23 | RPL23 | 0.04913671 | 0.05550717 | 0.05743356 | 0.143600495 | 0.344035038 | 0.289728361 | 0.03439 | 2.383954843 |
| 40S ribosomal protein S17-like;40S ribosomal protein S17 | RPS17;RPS17L | 0.0805229 | 0.04305663 | 0.08491369 | 0.191926033 | 0.272565329 | 0.239392464 | 0.04799 | 1.571906737 |
| Ribosomal RNA processing protein 1 homolog A | RRP1 | 0.00014903 | 0.00020298 | 0.00032652 | 0.001154437 | 0.002411969 | 0.002178172 | 0.01645 | 3.423687071 |
| U3 small nucleolar RNA-interacting protein 2 | RRP9 | 8.0926E-05 | 0 | 0.00035052 | 0.001436112 | 0.002802266 | 0.00160775 | 0.04076 | 3.254893438 |
| Ribosome biogenesis regulatory protein homolog | RRS1 | 0 | 0.00035721 | 0.00057228 | 0.005098748 | 0.009586263 | 0.005043131 | 0.01391 | 3.4953291 |
| SAP domain-containing ribonucleoprotein | SARNP | 0.00049531 | 0.00025244 | 0.00098941 | 0.00177165 | 0.005681279 | 0.003137572 | 0.04961 | 2.663233262 |
| Kanadaptin | SLC4A1AP | 8.0058E-05 | 4.4836E-05 | 9.9763E-05 | 0 | 0.000211998 | 0 | 0.03171 | 1.40492957 |
| Structural maintenance of chromosomes flexible hinge domain-containing protein 1 | SMCHD1 | 8.7112E-05 | 5.4407E-05 | 0.0001375 | 0.000643961 | 0.001163579 | 0.000564775 | 0.01629 | 2.886024975 |
| Surfeit locus protein 6 | SURF6 | 0.0002161 | 0.00042963 | 0.00097086 | 0.002714723 | 0.005228968 | 0.004023561 | 0.03106 | 3.227289473 |
| Transducin beta-like protein 3 | TBL3 | 0.00028427 | 0.00034144 | 0.00041052 | 0.001291727 | 0.003219351 | 0.002733132 | 0.02065 | 3.000842701 |
| THUMP domain-containing protein 1 | THUMPD1 | 7.1106E-05 | 0 | 0 | 0.001239047 | 0.002269458 | 0.001560646 | 0.00594 | 4.456022131 |
| TRMT1-like protein | TRMT1L | 0.00066196 | 0.00077492 | 0.00145812 | 0.002356394 | 0.005557788 | 0.003411742 | 0.04152 | 2.13839113 |
| tRNA (adenine(58)-N(1))-methyltransferase non-catalytic subunit TRM6 | TRMT6 | 0.00090343 | 0.00113169 | 0.00175559 | 0.002685266 | 0.009472147 | 0.006626619 | 0.04868 | 2.549799253 |
| Splicing factor U2AF 35 kDa subunit;Splicing factor U2AF 26 kDa subunit | U2AF1L4;U2AF1 | 0 | 0.0113692 | 0.02254909 | 0.066006558 | 0.069747258 | 0.103721157 | 0.0387 | 2.123036301 |
| Ribosome biogenesis protein WDR12 | WDR12 | 0.00103796 | 0.00073716 | 0.00173084 | 0.003207332 | 0.006142163 | 0.004008957 | 0.04153 | 1.949478876 |
| WD repeat-containing protein 3 | WDR3 | 0.00020805 | 0.00029641 | 0.00047884 | 0.002408321 | 0.003883152 | 0.00224613 | 0.01586 | 3.022361673 |
| WD repeat-containing protein 36 | WDR36 | 8.6922E-05 | 0.00014149 | 0.00040413 | 0.00069162 | 0.001856123 | 0.001265446 | 0.04822 | 3.160822966 |
| Tyrosine--tRNA ligase, cytoplasmic;Tyrosine--tRNA ligase, cytoplasmic, N-terminally processed | YARS | 0.00215138 | 0.00241108 | 0.00461613 | 0.006998871 | 0.017116031 | 0.015028527 | 0.04146 | 2.639955179 |
| Zinc finger CCCH domain-containing protein 11A | ZC3H11A | 0 | 0 | 1.8699E-05 | 0 | 0.000277896 | 0.000162946 | 0.01098 | 3.508475057 |
